# Regulation of multiple signaling pathways promotes the consistent expansion of human pancreatic progenitors in defined conditions

**DOI:** 10.1101/2023.09.06.556505

**Authors:** Luka Jarc, Manuj Bandral, Elisa Zanfrini, Mathias Lesche, Vida Kufrin, Raquel Sendra, Daniela Pezzolla, Ioannis Giannios, Shahryar Khattak, Katrin Neumann, Barbara Ludwig, Anthony Gavalas

## Abstract

The unlimited expansion of human progenitor cells in vitro could unlock many prospects for regenerative medicine. However, it remains an important challenge as it requires the decoupling of the mechanisms supporting progenitor self-renewal and expansion from those mechanisms promoting their differentiation. This study focuses on the expansion of human pluripotent stem (hPS) cell derived pancreatic progenitors (PP) to advance novel therapies for diabetes.

We obtained mechanistic insights into PP expansion requirements and, through a hypothesis-driven iterative approach, identified conditions for the robust and unlimited expansion of hPS cell derived PP cells under GMP-compliant conditions. We show that the combined stimulation of specific mitogenic pathways, suppression of retinoic acid signaling and inhibition of selected branches of the TGFβ and Wnt signaling pathways are necessary for the effective decoupling of PP proliferation from differentiation. This enabled the reproducible, 2000-fold, over ten passages and 40-45 days, expansion of PDX1^+^/SOX9^+^/NKX6-1^+^ PP cells. Transcriptome analyses confirmed the stabilisation of PP identity and the effective suppression of differentiation. Using these conditions, PDX1^+^/SOX9^+^/NKX6-1^+^ PP cells, derived from different, both XY and XX, hPS cells lines, were enriched to nearly 90% homogeneity and expanded with very similar kinetics and efficiency. Furthermore, non-expanded and expanded PP cells, from different hPS cell lines, were differentiated in microwells into homogeneous islet-like clusters (SC-islets) with very similar efficiency. These clusters contained abundant β-cells of comparable functionality as assessed by glucose-stimulated insulin secretion assays.

These findings established the signaling requirements to decouple PP proliferation from differentiation and allowed the consistent expansion of hPS cell derived PP cells. They will enable the establishment of large banks of GMP-derived PP cells derived from diverse hPS cell lines. This approach will streamline SC-islet production for further development of the differentiation process, diabetes research, personalized medicine and cell therapies.

## INTRODUCTION

Diabetes is a global epidemic affecting almost 10% of the world’s population. Its two main forms result from the immune destruction (type 1) or malfunction (type 2) of insulin-producing β cells residing in the pancreatic islets of Langerhans. Severe cases of diabetes often necessitate whole pancreas transplantation or pancreatic islet transplantation to restore metabolic control and insulin independence. However, both approaches face critical limitations due to the scarcity of tissue donors and the requirement for life-long immunosuppression which hinders, the already limited β-cell self-renewal, and is associated with severe side effects and morbidity (Nir et al., 2007).

The remarkable progress over the last decade in the differentiation of human pluripotent stem (hPS) cells into pancreatic islet cells (SC-islet cells) suggests that this approach could provide an unlimited source of β-cells for transplantations and personalized medicine (Amin et al., 2018; Balboa et al., 2022; Du et al., 2022; Hogrebe et al., 2020; Millman et al., 2016; Ramzy et al., 2021; Shapiro et al., 2021). The underlying strategy recapitulates in vitro the stepwise differentiation of epiblast cells initially to definitive endoderm and then to pancreatic progenitor-containing (PP) cells, pancreatic endocrine progenitors and, finally, to pancreatic islet cells (Kroon et al., 2008; Pagliuca et al., 2014; Rezania et al., 2014; Russ et al., 2015; Zeng et al., 2016; Zhu et al., 2016b). Clinical trials employing either GMP-grade PP cells in a non-immunoprotective macroencapsulation device (Shapiro et al., 2021) (NCT03163511) or GMP-grade SC-islet cells in an immunoprotective macroencapsulation device (NCT02239354) suggested that diabetes cell therapies can become a reality. Obstacles to be overcome include the limited maturation of the resulting β-cells, incomplete conversion of hPS cells into endocrine cells and the large number of SC-islet cells required for a single transplantation. Elucidating the mechanisms that maintain the self-renewal of pure PP cells, while inhibiting their differentiation, would allow the establishment of expandable populations of PP cells and address the need for large numbers of pancreatic endocrine cells.

During development, the intersection of several signals induces the pancreatic anlage at the posterior foregut. Repression of posteriorly derived Wnt signaling is initially essential to define the foregut region (McLin et al., 2007). There, the pancreas and liver develop from a common progenitor (Cerda-Esteban et al., 2017; Deutsch et al., 2001). Bipotent progenitors, located distant to the cardiac mesoderm, which secretes the pro-hepatic signals FGF10 and BMPs, remain competent to adopt pancreatic fate (Jung et al., 1999; Rossi et al., 2001). The combination of RA signaling, derived from the somitic mesoderm (Martin et al., 2005; Molotkov et al., 2005), and endothelial signals, derived from the dorsal aorta (Lammert et al., 2001), induces the formation of pancreatic progenitors. A complex, stage-specific expression of several transcription factors (TFs) guides pancreatic cell development (Duvall et al., 2022). The epithelial pancreatic progenitors are characterized by the combined expression of several TFs, most notably Pdx1, Nkx6-1 and Sox9, which are essential for progenitor self-renewal and subsequent endocrine differentiation. Pdx1 expression is induced at the boundary between the Sox2-expressing anterior endoderm and the Cdx2-expressing posterior endoderm and it is necessary to maintain the pancreatic identity (Sherwood et al., 2009). Its function is reinforced by Sox9 which cooperates with Pdx1 to repress Cdx2 expression in the pancreatic anlage (Shih et al., 2015). Pdx1 (Jonsson et al., 1994; Offield et al., 1996) and Sox9 (Seymour et al., 2007) loss-of-function experiments resulted in pancreatic agenesis, establishing their key role in maintaining pancreatic identity. Sox9 is subsequently essential for the induction of Neurog3, the TF which is necessary and sufficient for the induction of the pancreatic endocrine lineage (Gradwohl et al., 2000; Grapin-Botton et al., 2001). Nkx6-1 is another key contributor to the maintenance and self-renewal of pancreatic progenitors (Sander et al., 2000). It is also required for β-cell specification since ectopic Nkx6-1 expression directed endocrine precursors into β-cells, whereas β-cell specific ablation of Nkx6-1 diverted these cells into the other endocrine lineages (Schaffer et al., 2013). Given the importance of these three TFs, the identification of conditions to stabilize PDX1^+^/SOX9^+^/NKX6-1^+^ cells will set the stage for unlimited PP cell expansion and their efficient differentiation into SC-islets PDX1^+^/SOX9^+^/NKX6-1^+^ PP cells undergo self-renewal *in vivo* for a limited amount of time. During development, feed-forward loops, that steer cells toward differentiation, operate in parallel with maintenance and self-renewal mechanisms. To achieve unlimited expansion of PP cells *in vitro* it will be necessary to disentangle differentiation signals from proliferation and maintenance signals. Several pathways have been implicated in these processes. Notch signaling mediates progenitor self-renewal as well as lineage segregation (Afelik et al., 2012; Murtaugh et al., 2003; Shih et al., 2012) and its downregulation is necessary for endocrine lineage specification (Apelqvist et al., 1999; Jensen et al., 2000). Notch signaling is itself regulated by the extracellular signal sphingosine-1-phosphate (S1p) (Serafimidis et al., 2017). Canonical Wnt signaling has also been implicated in PP maintenance (Afelik et al., 2015). Low levels of endogenous RA signaling are involved in subsequent differentiation steps of PP cells (Kobayashi et al., 2002; Lorberbaum et al., 2020; Tulachan et al., 2003; Vinckier et al., 2020). Finally, expression of TGFβ ligands and receptors is widespread during pancreas development (Crisera et al., 1999; Tulachan et al., 2007) and it has been suggested that the TGFβ pathway activity regulates the pancreatic lineage allocation (Miralles et al., 1998; Sanvito et al., 1994). TEAD, and its coactivator YAP, act as integrators of extracellular signals to activate key pancreatic signaling mediators and transcription factors which promote the expansion of pancreatic progenitors and their competence to differentiate into endocrine cells (Cebola et al., 2015; Serafimidis et al., 2017). Cell confinement acts as a mechanic signal to downregulate YAP and trigger endocrine differentiation (Mamidi et al., 2018) and this finding has been applied to promote the differentiation of hPS cell-derived PPs into endocrine cells (Rosado-Olivieri et al., 2019).

Culture conditions to expand hPS cell-derived PDX1^+^/SOX9^+^ cells have been previously reported but these methods relied either on feeder layers or limiting 3D conditions in an agarose hydrogel matrix and did not maintain the expression of the crucial TF NKX6-1 (Konagaya and Iwata, 2019; Trott et al., 2017). A recent study claimed the expansion of a hPS cell-derived cell population, containing PDX1^+^/NKX6-1^+^ cells, on fibronectin and in a defined medium using SB431542, a broad (ALK4/5/7) TGFβ inhibitor (Inman et al., 2002; Nakamura et al., 2022). Unfortunately, the reproducibility of the expansion was not documented and the reported percentage of PDX1^+^/NKX6-1^+^ PP cells in the expanded cells varied widely among the three different hPS cell lines used, from 65% to 35% and 20%. Furthermore, RNA Seq data showed that *NKX6-1* expression actually decreased during expansion, in two out of the three samples presented, (E-MTAB-9992) suggesting that NKX6-1 expression was not reliably maintained under those culture conditions. In another study, PP cells, generated through genetic reprogramming of human fibroblasts into endoderm progenitor cells, could be expanded in a chemically defined medium containing epithelial growth factor (EGF), basic fibroblast growth factor (bFGF), and SB431542, another general (ALK4/5/7) TGFβ inhibitor. However, the percentage of PDX1^+^/NKX6-1^+^ cells was less than 20% (Zhu et al., 2016a). The authors then identified a BET bromodomain inhibitor that could mediate the unlimited expansion of PDX1^+^/NKX6-1^+^ PP cells on feeder layers of mouse embryonic fibroblasts (Ma et al., 2022). Unfortunately, the use of feeder layers precluded the use of this method in a clinical setting and it did not address the mechanistic requirements for the expansion.

PP cells express the components of a large number of signaling pathways and we hypothesized that a longitudinal transcriptome analysis of non-expanding and occasionally expanding PP cells would provide candidate signaling pathways reasoning that upregulated pathways in expanding cells would promote expansion, whereas downregulated pathways would be blocking expansion and/or favoring differentiation. We leveraged these findings and employed a hypothesis-driven iterative process to define conditions that allowed robust, unlimited expansion, of hPS cell-derived PP cells. We found that the combined stimulation of specific mitogenic pathways, suppression of retinoic acid signaling and inhibition of selected branches of the TGFβ and Wnt signaling pathways enabled the 2000-fold expansion of PP cells over ten passages and 40-45 days. Expansion conditions are GMP-compliant and enable the robust, reproducible expansion, as well as intermediate cryopreservation, of PP cells derived from diverse hPS cell lines with essentially identical growth kinetics. These conditions select PDX1^+^/SOX9^+^/NKX6-1^+^ PP cells, suggesting that they will be advantageous for hPS cell lines that may differentiate less efficiently into PP cells. Expanded PP cells differentiated, with very similar efficiency to non-expanded cells, into SC-islet clusters that contained functional β-cells as shown by glucose-stimulated insulin secretion (GSIS) assays.

Our findings will allow the establishment of large banks of PP cells derived under GMP conditions from diverse hPS cell lines, streamlining the generation of SC-islet clusters for further development of the differentiation procedure, diabetes research, personalized medicine and cell therapies.

## RESULTS

### Identification of candidate signaling pathways implicated in PP expansion

H1 hPS cells were differentiated into PP cells and PP cell expansion was attempted using the initial conditions (CINI) (Table S1) that relied on EGF and FGF2 and A83-01, a broad TGFβ inhibitor of ALK4/5/7 (Tojo et al., 2005). The CINI formulation was similar, but not identical, to that used previously in the attempted expansion of PP cells derived through the reprogramming of human fibroblasts into endoderm progenitor cells, where SB431542, another brοad TGFβ inhibitor of similar specificity (Inman et al., 2002), was employed (Zhu et al., 2016a). PP cells of comparable quality as assessed by qPCR for the expression of the PP markers *PDX1*, *NKX6-1* and *SOX9*, as well as for the expression of the liver and gut markers, *AFP* and *CDX2*, respectively, were plated at high density on Matrigel-coated plates and passaged every 4-6 days. In most instances, cell numbers remained either constant or decreased, resulting in growth arrest. Occasionally (one out of five attempts), PP-containing cells (referred to hereafter just as PP cells for simplicity) expanded and could then be maintained for at least up to ten passages (Figure 1A). This was not correlated with initial favorable expression patterns of *PDX1*, *NKX6-1*, *SOX9*, or *AFP* and *CDX2* (Figure 1B). Immunofluorescence experiments and qPCR analyses suggested the maintenance of the PP identity in these expanding PP (ePP) cells since expression of *PDX1*, *NKX6-1* and *SOX9* remained stable at both the protein (compare Figure S1A, B and S1C) and transcript levels (Figure S1D). Immunofluorescence analysis of cryosections showed that these ePP cells could subsequently differentiate into endocrine cells (Figure S1E). These results suggested that EGF, FGF2 and broad TGFβ inhibition were not sufficient for the reproducible expansion of hPS cell-derived PP cells.

**Figure 1.**
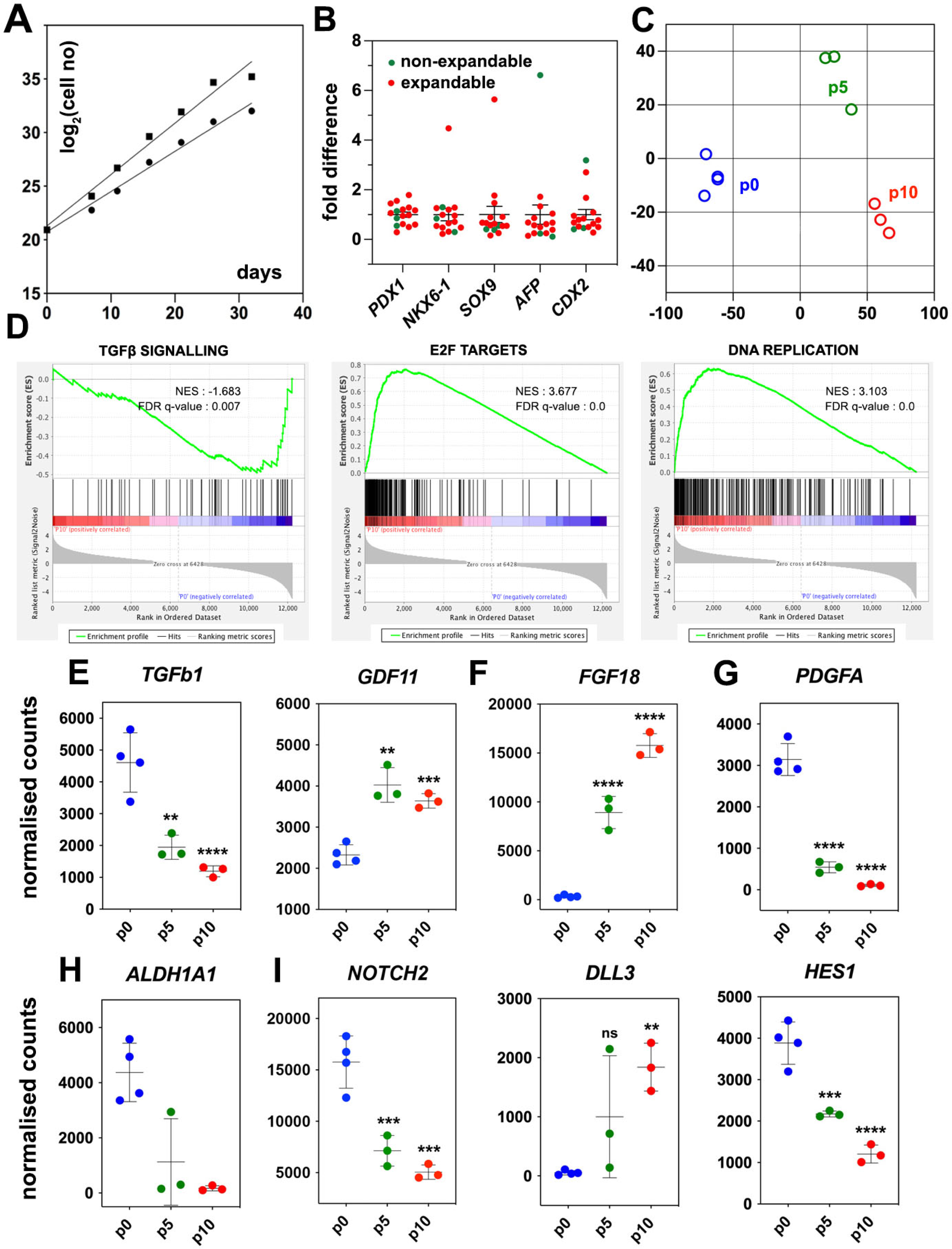
Regulated genes and signaling pathways during the expansion of PP-containing cells (PP cells) under the initial condition (CINI) **(A)** Growth curves of two samples showing exponential expansion of PP cells for 32 d. **(B)** Expression of pancreatic (*PDX1*, *NKX6-1*, *SOX9*) as well as liver and gut markers (*AFP*, *CDX2*, respectively) at the PP stage of representative differentiations before expansion. Expression at each sample is shown as a fold difference from the average expression level. In red are shown samples that could not be expanded and in green samples that could be expanded. **(C)** MDS plot representing the Euclidian distance of samples at p0 (n=4), p5 (n=3) and p10 (n=3). **(D)** Enrichment plots for regulated genes (p0 vs p10) of the TGFβ signaling pathway, E2F target genes and DNA replication show a negative correlation of the TGFβ pathway but a positive correlation of E2F target genes and DNA replication with expansion. **(D-H)** Transcript levels expressed in normalized RNA-Seq counts of p0 (n=4), p5 (n=3) and p10 (n=3) PP cells for genes encoding signals or receptors of the TGFβ (D), FGF (E) and PDGF (F) signaling pathways, the RA producing enzyme ALDH1A1 (G) as well as components of the NOTCH (H) signaling pathway. Horizontal lines represent the mean ± SD. Statistical tests were one-way ANOVA using p0 as the control condition for the comparison with *p* ≤ 0.033 (*), *p* ≤ 0.002 (**), ≤ 0.0002 (***) and ≤ 0.0001 (****).

PP cells express the components of numerous signaling pathways (Table S2 and S3) and we hypothesized that transcriptome comparison of non-expanding and CINI ePP cells could identify pathways implicated in the expansion. Upregulated pathways in ePP cells would be promoting expansion, whereas downregulated pathways would be blocking expansion and/or favoring differentiation. Thus, we compared the RNA Seq profiles of cells directly after their differentiation into pancreatic progenitors (dPP) (p0) (n=4), and CINI ePP cells at passage 5 (p5) (n=3) and p10 (n=3). Of the analyzed dPP samples, one was subsequently successfully expanded. Analysis for differentially expressed genes (DEG) identified groups of genes that were continuously up or down-regulated genes and genes showing changes only within the first five passages (Figure S1F and Table S2). Metric multidimensional scaling (MDS) plot of the Euclidian distances of the samples demonstrated clustering of the samples strictly according to their passage number. The single p0 sample, which subsequently expanded successfully, clustered with the other p0 samples suggesting that it was not fundamentally different from samples that failed to expand. These findings suggested a reproducible adaptation of ePP cells to the culture conditions primarily during the first five passages (Figure 1C). Gene Set Enrichment Analysis (GSEA) suggested that components of the TGFβ signaling pathway were negatively correlated with expansion whereas E2F target genes and DNA replication were positively correlated (Figure 1D). E2F TFs are indirectly activated by growth signals to regulate multiple cell cycle genes and promote cell proliferation (Ertosun et al., 2016; Rubin et al., 2020). This raised the possibility that expanding cells upregulate their own growth factors engaging themselves in an autocrine growth loop. On the other hand, all key components of the Hippo pathway were expressed but were not transcriptionally regulated, with the exception of *SHANK2*, a negative regulator and *WWTR1(TAZ)*, a transcriptional regulator (Meng et al., 2016) (Table S3). To gain a better mechanistic insight into potentially involved signaling pathways, we then examined the expression kinetics of all individual ligands and receptors expressed in dPP and ePP cells.

High expression of TGFβ ligands as well as Type I and Type II receptors suggested that all three branches of this signaling pathway were active at p0 (Table S3). The negative correlation of the TGFβ signaling pathway with the expansion appeared to retrospectively justify the use of the broad TGFβ signal inhibitor A83-01 in the CINI expansion medium (Tojo et al., 2005). However, since this pathway has several branches often with cell-dependent opposing functions, we assessed the expression kinetics of all expressed TGFβ ligands and receptors. The most highly expressed TGFβ ligands, in dPP cells, were *TGFb1* and *BMP2*, the expression of which was repressed nearly four-fold during expansion (Figure 1E, S1G). TGFb1 and BMP2 act through the ALK1/5 and ALK3/6/2 receptors, respectively (Brown and Schneyer, 2010), so A83-01 would block TGFB1 but not BMP2 signaling. On the other hand, *ALK4* and its ligand, *GDF11*, remained strongly expressed throughout expansion; *GDF11* was even upregulated (Figure 1E, Table S3), suggesting a possible positive role of this TGFβ signaling branch in the expansion of pancreas progenitors. These findings suggested that the use of broad TGFβ inhibitors such as A83-01 or SB431542 may not be optimal because they do not inhibit BMP2 signaling whereas they block ALK4 signaling, which might promote PP expansion through GDF11. This provided a mechanistic rationale for the subsequent use of more specific TGFβ inhibitors.

To account for the positive correlation of the expression of E2F targets with expansion, we then examined the gene expression kinetics of growth factors during expansion. There was a striking, nearly 30-fold, upregulation of *FGF18* expression (Figure 1F and Table S3) suggesting a strong requirement for the activation of the MAPK pathway. Incidentally, FGF18 has a selective affinity for FGFR3 and 4 (Zhang et al., 2006), the two most highly expressed *FGFRs* in dPP and ePP cells. Other *FGFs* were weakly expressed and not upregulated during expansion (Table S3). Expression of *FGF18*, as well as of other *FGFs* and *FGFRs*, was detected at several stages of pancreas development in whole pancreata (Dichmann et al., 2003). Another potentially interesting growth factor signaling pathway was the PDGF signaling pathway. PDGFR signaling has not been so far implicated in pancreas development or PP expansion but it promotes the expansion of young beta cells (Chen et al., 2011). *PDGF* receptors *A* and *B* were stably and strongly expressed during expansion but the initially strong expression of two of the expressed ligands, PDGFA and B was downregulated by 30- and 15-fold, respectively (Figure 1G, S1H, Table S3). These findings provided a mechanistic rationale to provide FGF18 during expansion and/or inhibit PDGF signaling in subsequent experiments.

RA promotes the differentiation of pancreatic progenitors during development (Lorberbaum et al., 2020; Martin et al., 2005; Ostrom et al., 2008; Vinckier et al., 2020). PP cells highly expressed the retinol dehydrogenase 11 (*RDH11*) and aldehyde dehydrogenase 1a1 (*ALDH1A1*) genes, which encode enzymes that convert vitamin A into retinoic acid, as well as the RA nuclear receptors *RARA*, *RARG*, *RXRA* and *RXRB*. Therefore, in the presence of vitamin A, this pathway could act in an autocrine manner to promote differentiation. Strikingly, successful expansion was accompanied by a dramatic, 25-fold, downregulation in the expression of the RA-producing enzyme *ALDH1A1* (Figure 1H, Table S3). Thus, we speculated that PP cells are poised to initiate RA-mediated differentiation in an autocrine feed-forward loop and, therefore, eliminating vitamin A in the medium might stabilize the PP state.

The Notch signaling pathway is implicated in multiple aspects of pancreatic development including the expansion of pancreatic progenitors and lineage selection (Afelik et al., 2012; Apelqvist et al., 1999; Murtaugh et al., 2003; Qu et al., 2013; Seymour et al., 2020; Shih et al., 2012). The expression of Notch receptors, ligands and effectors, such as HES1, displayed a complex pattern of changes during expansion. (Figure 1I, S1I, Table S3) that suggested an overall attenuation, but not silencing, of the pathway.

In summary, the findings suggested that several mechanisms may contribute to stabilizing the PP state. Alone or in combination, inhibition of selected branches of the TGFβ pathway, attenuation of the Notch pathway, additional mitogenic stimulation with FGF18, and suppression of RA and PDGF signaling may lead to reproducible expansion of the PP cells.

### Elimination of RA and selective TGFβ inhibition allow reproducible expansion of PP cells

We assessed several expansion culture conditions (summarized in Table S1), based on the mechanistic insights discussed above, using PP cells generated with an adaptation of published procedures (Balboa et al., 2022; Mahaddalkar et al., 2020; Rezania et al., 2014; Shi et al., 2017) (Table S4). Our strategy to achieve reproducible and robust PP expansion involved activation or inhibition of specific signaling pathways, either individually or in combination. We first suppressed RA signaling by substituting the vitamin A-containing B27 supplement with a Vitmain A-free B27 formulation. Additionally, we substituted A83-01 with the ALK5 II inhibitor (ALK5i II) that targets primarily ALK5, and to a lesser extent ALK3/6, but not ALK4 (Gellibert et al., 2004). This was named condition 0 (C0). C0 was further elaborated by replacing FGF2 with FGF18 (C1) as well as adding the Notch inhibitor XXI (Seiffert et al., 2000) (C2), or the PDGFR inhibitor CP673451 (Roberts et al., 2005) (C3) or adding both Notch and PDGFR inhibitors (C4). Since FGF2 and FGF18 belong to different FGF subfamilies and have overlapping, but not identical, FGFR specificities (Zhang et al., 2006), we also addressed a possible synergistic effect of these FGFs in PP expansion by combining them in C5. In contrast to CINI, three of these conditions, C0, C1 and C5, resulted in the reproducible expansion of PP cells for at least 10 (C0, C1, n=3) or 11 (C5, n=5) passages with doubling times (T_d_) of 3.9, 3.6 and 2.3 days, respectively (Figure 2A, Table S1).

**Figure 2.**
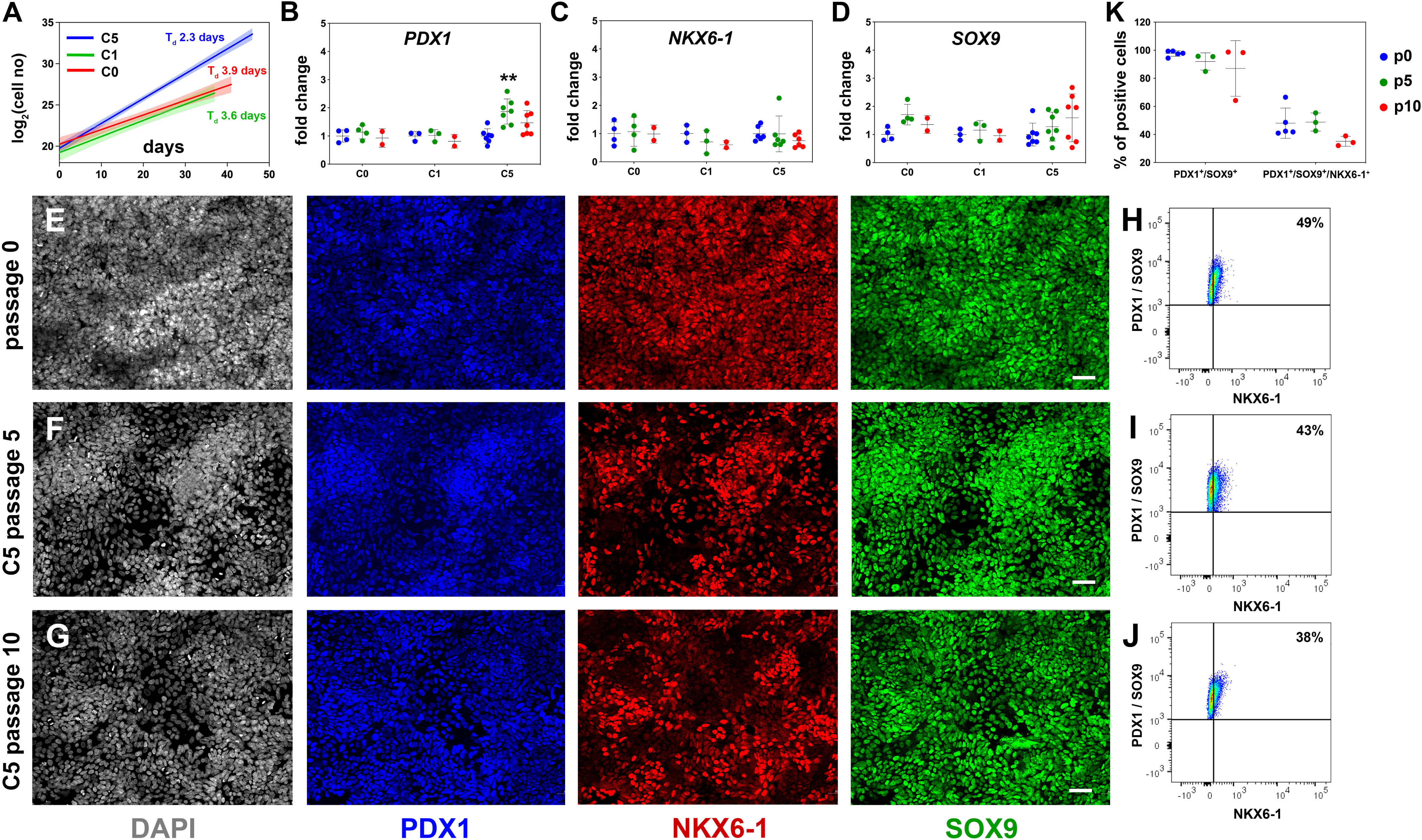
Reproducible expansion of PP cells under condition 5 (C5) **(A)** Growth curves and regression analysis for PP cells expanded under C0, C1 and C5 for at least 10 passages. The doubling time (T_d_) of C5-expanded cells (n=7) was 2.3 days with a 95% confidence interval (CI) of 2.13-2.51 days. This was clearly increased as compared to C0-(n=2, Td=3.92 days, 95% CI = 3.22-4.98 days) and C1-expanded cells (n=2, Td=3.55 days, 95% CI = 2.88-4.62 days). The translucent shading represents the 95% CI of the growth rate at the different conditions. **(B-D)** Gene expression profile of C0-, C1- and C5-expanded cells as shown by qPCR for expression of the key pancreas progenitor markers *PDX1* (B), *NKX6.1* (C) and *SOX9* (D) during the expansion. Expression is normalized against the expression of each marker at p0. **(E-G)** Representative images of immunofluorescent staining of p0 PP cells (E) as well as C5-expanded cells at p5 (F) and p10 (G) for the PP transcription factors PDX1, NKX6.1 and SOX9. **(H-J)** Flow cytometry analysis of p0 PP cells (H), as well as C5-expanded cells at p5 (I) and P10 (J) for PDX1, NKX6.1 and SOX9. **(K)** Cumulative results of the flow cytometry analyses for PDX1^+^/SOX9^+^ and PDX1^+^/SOX9^+^/NKX6.1^+^ C5-expanded PP cells at p0, p5 and p10. Horizontal lines represent the mean ± SD. Statistical tests were two-way ANOVA with Tukey’s test, using p0 as the control condition for the comparison with *p* ≤ 0.033 (*), *p* ≤ 0.002 (**), ≤ 0.0002 (***) and ≤ 0.0001 (****). Scale bar corresponds to 50 μm.

Assessment of the expression of key PP genes by qPCR at p5 and p10 suggested that, in all three conditions, initial high levels of *PDX1, NKX6-1* and *SOX9* expression were maintained (*NKX6-1*, *SOX9*) or even transiently increased at p5 (C5, *PDX1*) (Figure 2B-D). Expression of *PTF1A* was dramatically decreased in all three conditions suggesting a shift to a bipotent endocrine/duct progenitor identity (Figure S2A). Expression of *FOXA2*, involved in PP cell maintenance and endocrine lineage development (Gao et al., 2010; Gao et al., 2008; Lee et al., 2005; Lee et al., 2019), was upregulated in C5, indicating that cells expanded in this condition might be more amenable to terminal endocrine differentiation (Figure S2B). On the other hand, there was a statistically significant increase in the expression of the liver and gut markers *AFP* and *CDX2* in C0 and C1 and a similar, but weaker, trend in C5 (Figure S2C, D). This suggested that these conditions (C0, C1, C5) allowed cells to express aspects of liver and gut programs. Since C5 had a significantly lower T_d_ and a less pronounced increase in *AFP* and *CDX2* expression, we concentrated on the analysis and further improvement of C5.

To first confirm the C5 qPCR results at the protein level, we assessed the expression of several markers by immunofluorescence for C5 ePP cells. Immunofluorescence suggested that PDX1 and SOX9 were uniformly expressed at p0, p5 as well as p10 and that a large number of PP cells were NKX6-1^+^ at all three different time points even though expansion appeared to somewhat reduce the number of NKX6-1^+^ cells. (Figure 2E-G). Similarly, FOXA2 remained widely expressed at p0, p5 and p10 (Figure S2E-G). Expression of both AFP and CDX2 appeared to increase transiently upon expansion, at p5 (Figure S2H-J). We then quantified the expression of the key PP markers PDX1, SOX9 and NKX6-1 by flow cytometry at p0, p5 and p10. In these analyses, PDX1^+^/SOX9^+^ cells were first identified based on the gates set by the corresponding control stainings and, then, the number of NKX6-1^+^ cells was determined in these pre-gated cells based on the gate set by the corresponding control staining (Figure S2K-N). These experiments confirmed the immunofluorescence experiments showing that at least 90% of all cells were PDX1^+^/SOX9^+^ at all passages examined (Figure S2K-N), whereas nearly 50% of the cells at p0 and p5 and nearly 40% at p10 were PDX1^+^/SOX9^+^/NKX6-1^+^. There was a small, apparent progressive drop in the number of PDX1^+^/SOX9^+^ as well as the number of PDX1^+^/SOX9^+^/NKX6-1^+^ cells which did not reach statistical significance (Figure 2H-K, S2K-N).

These experiments established that C5 allowed reproducible and robust PP expansion, the basis of which was the elimination of RA signaling and selective TGFβ inhibition. Additionally, EGF, FGF2 and FGF18 synergized to promote a more robust expansion.

### Expansion conditions promote primarily the proliferation of PP cells

To understand the mechanism behind the successful and reproducible expansion, we asked whether it was due to enhanced cell survival, proliferation, or a combination of both. De novo generated PP cells were expanded in either C5 or CINI and assayed at p3 for EdU incorporation and apoptosis. The EdU incorporation analysis revealed that C5 cultures contained significantly more EdU^+^ expanding cells (10.4 ± 1.2%) than CINI (4.2 ± 0.6%) cultures (*n=4*) (Figure 3A-C, S3A). The apoptosis assays showed that even though the percentage of 7-AAD^+^/Annexin V^+^ cells appeared higher in CINI-expanded cells (22.1 ± 5.0%) than in C5-expanded cells (17.2 ± 4.7%) this difference did not reach statistical significance (*n=6*) (Figure 3C and Figure S3B, C). Therefore, C5, as compared to CINI, promotes primarily proliferation rather than survival of PP cells but it is important to note that cell death also appears higher in CINI cells. It is likely that the combination of these effects results in reproducible expansion under C5.

**Figure 3.**
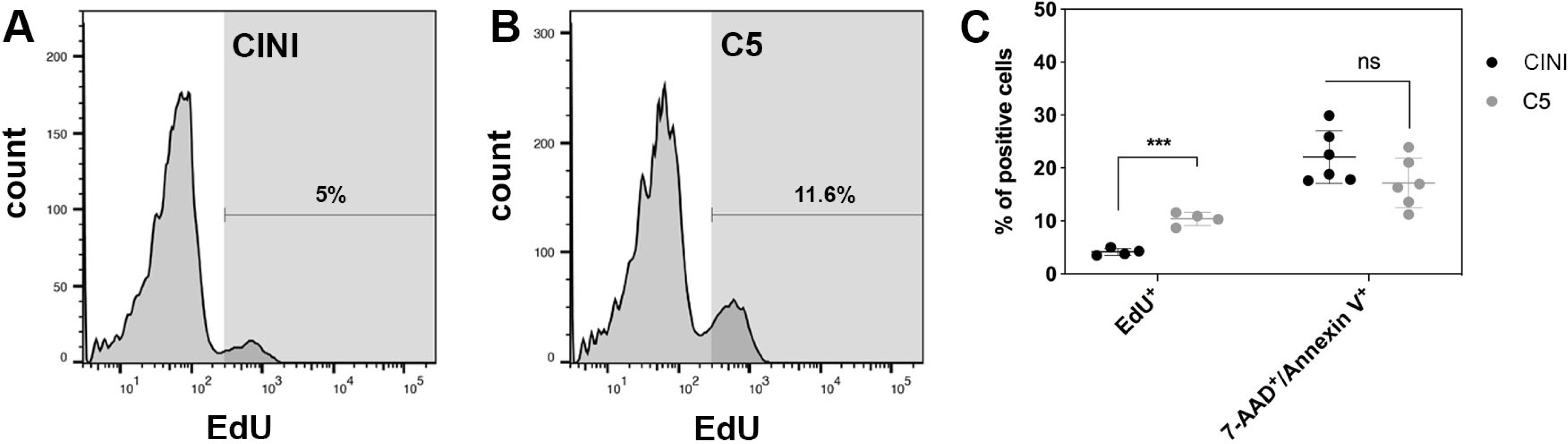
Expansion of PP cells promotes primarily their proliferation rather than their survival. **(A, B)** Histogram plots showing the % of PP cells that had incorporated EdU during expansion under CINI (A) and C5 (B). **(C)** Summary of flow cytometry data comparing proliferation, measured by EdU incorporation (n=4), and cell death, measured by Annexin V/7-AAD staining (n=6), of PP cells expanded under CINI and C5. Horizontal lines represent mean ± SD. Means were compared with multiple t-tests and significance is p ≤ 0.033 (*), p ≤ 0.0021 (**), p ≤ 0.0002 (***) or p ≤ 0.0001 (****).

### Canonical Wnt inhibition restricts upregulation of hepatic fate and promotes PP identity

The upregulation of *AFP* and *CDX2* in C5 ePP cells suggested a drift towards hepatic and intestinal fates that would hinder the efficiency of differentiation into endocrine cells and subsequent maturation (Nair et al., 2019). During pancreas development, non-canonical Wnt signaling specifies bipotent liver/pancreas progenitors to pancreatic fates whereas canonical Wnt signaling leads to liver specification and the emergence of gastrointestinal identity (Ober et al., 2006; So et al., 2013; Muñoz-Bravo et al., 2016; Rodríguez-Seguel et al., 2013). PP cells, at p0 as well as subsequent passages, strongly and stably expressed several *WNT* receptors, co-receptors as well as canonical and non-canonical signals (Table S3). Thus, to suppress the upregulation of hepatic and/or intestinal fates, we supplemented C5 with the canonical Wnt inhibitor IWR-1 to selectively inactivate the canonical Wnt signaling (Chen et al., 2009) (condition 6; C6, Table S1).

C6 ePP cells retained a growth rate similar to C5, showing that IWR-1 supplementation did not significantly affect their growth; their T_d_ was 2.5 days as opposed to 2.3 days for C5 ePP cells (Figure 4A). C6 expansions using vitronectin-N (VTN-N) showed no change in expansion efficiency or promotion of the PP identity (*n=3*). VTN-N is a defined peptide that can be produced under GMP-conditions. Since all other media components can also be produced under GMP-conditions, this finding established that this approach is also GMP-compliant (Figure 4A).

**Figure 4.**
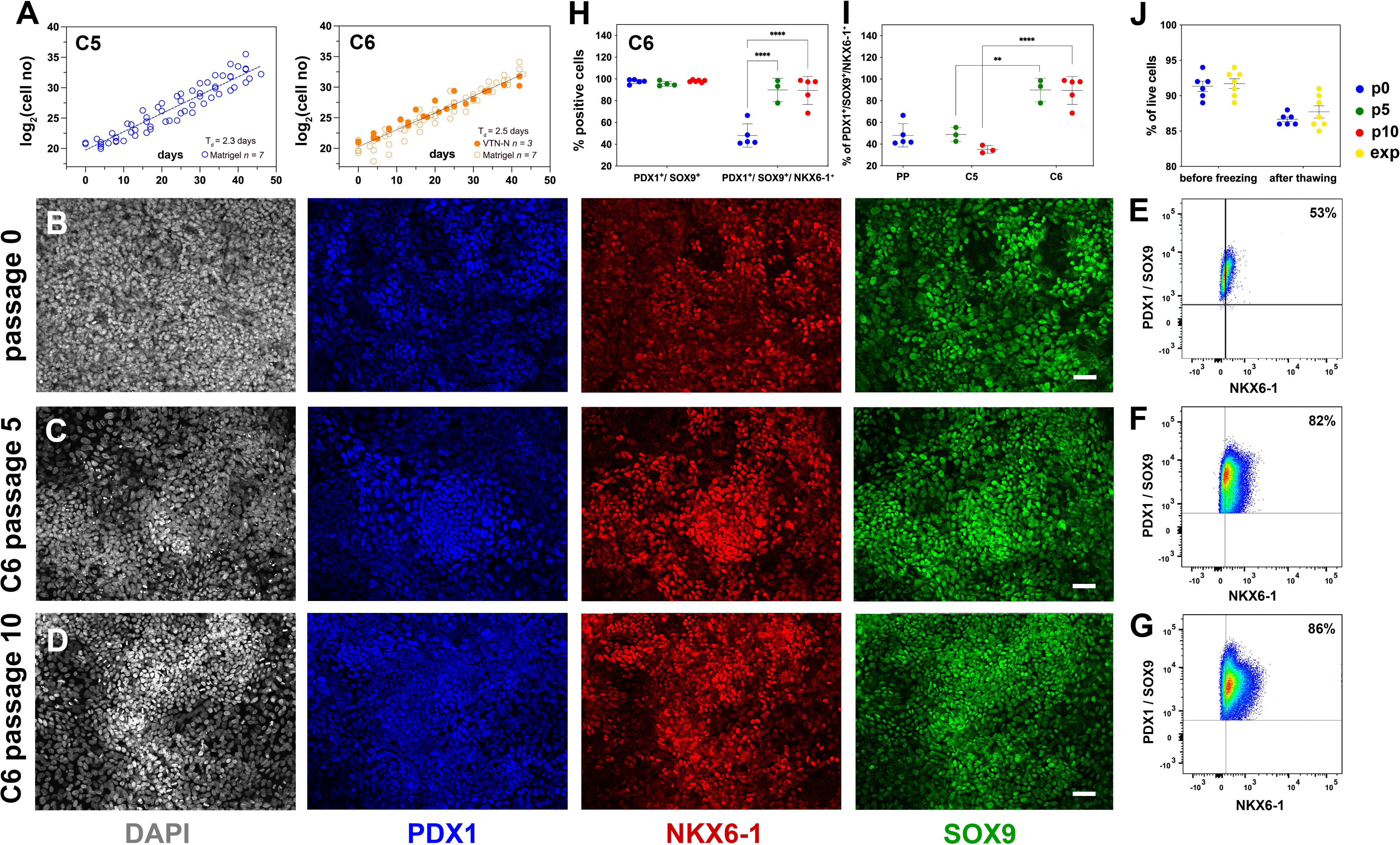
Reproducible expansion under condition 6 promotes PP identity. **(A)** Growth curves and regression analysis for PP cells expanded under C5 and C6 for ten passages. The regression line for C5 showed a doubling time of 2.3 d (n=7) as compared to 2.5 d (n=10) for C6. **(B-D)** Representative images of immunofluorescent staining of p0 PP cells (B) as well as C6-expanded cells at p5 (C) and p10 (D) for the PP transcription factors PDX1, NKX6.1 and SOX9. **(E-G)** Flow cytometry analysis of non-expanded p0 PP cells (E) and C6-expanded cells at p5 (F) and p10 for PDX1^+^/SOX9^+^/ NKX6.1^+^ cells (G). **(H, I)** Cumulative results of the flow cytometry analyses for PDX1^+^/SOX9^+^ and PDX1^+^/SOX9^+^/NKX6.1^+^ C6-expanded cells at p0, p5 and p10 (H) and comparison of the % of C5- and C6-expanded PDX1^+^/SOX9^+^/NKX6.1^+^ cells at p5 and p10 (I). **(J)** Survival rates of PP cells before freezing and after thawing at p0 or during expansion. Horizontal lines represent the mean ± SD. Statistical tests were two-way ANOVA with Tukey’s test, using p0 as the control condition for the comparison with *p* ≤ 0.033 (*), *p* ≤ 0.002 (**), ≤ 0.0002 (***) and ≤ 0.0001 (****). Scale bar corresponds to 50 μm.

Relative to C5 ePP cells of the same passage number, C6 ePP cells retained strong expression of *PDX1, NKX6-1* and *SOX9* and similar expression levels of *PTF1A* and *FOXA2* (Figure S4A). Importantly, the expression of the liver markers examined, *AFP, HHEX* and *TTR,* was significantly lower, by p10, in the C6 ePP cells as compared to C5 ePP cells (Figure S4B). However, expression of *CDX2* was only marginally reduced (Figure S4A). The expression of *PDX1, NKX6-1, SOX9* and *FOXA2* was also examined at the protein level by immunofluorescence which suggested that their expression was stable and persisted at high levels (Figure 4B-D, S4C-E). Similarly to C5 ePP cells, expression of AFP and CDX2 in C6 ePP cells was detectable by immunofluorescence at p5 but significantly reduced at p10 (Figure S4F-H). Flow cytometry quantification of the cells expressing key PP markers at p0, p5 and p10 (Figure 4E-I, S4I-L) showed a significant increase in the number of PDX1^+^/SOX9^+^/NKX6-1^+^ C6 ePP cells from 48%±11% (n=5) at p0 to 90%±10% (n=3) at p5 and 95%±5% (n=5) at p10 (Figure 4H). This, along with the reduction in the expression of liver markers, was an additional significant improvement over C5 ePP cells at both p5 and p10 (Figure 4I). C6 ePP cells could be cryopreserved and recovered with high survival rates (>85%) with no apparent loss of proliferative capacity (Figure 4J). Chromosomal stability was also assessed after 16 passages analyzing the G-banding of at least 20 metaphases and no alterations were found (Figure S4M).

To further repress the expression of liver and gut markers, we reconsidered BMP inhibition. ALK5i II is considered a relatively weak inhibitor of ALK3 (Gellibert et al., 2004). However, BMP2, a ligand of ALK3, was significantly downregulated in the CINI ePP cells (Figure S1G, Table S3). Attempting to effectively inhibit ALK3, we substituted ALK5i II with LDN-193189, which inhibits ALK3 with higher potency (Sanvitale et al., 2013). LDN-193189 was used alone (C7) or in combination with IWR-1 (C8), however, none of these conditions was efficient in PP cell expansion (Table S1).

In summary, C6, featuring inhibition of the canonical Wnt signaling by IWR-1, promoted the selection of PDX1^+^/SOX9^+^/NKX6-1^+^ in the expanding cell population and mitigated the upregulation of liver markers at both the gene and protein expression levels. C6 is a robust, highly reproducible, GMP-compliant procedure suitable for application in cell therapies.

### Expansion under C6 stabilizes PP cell identity by repressing differentiation and alternative cell fates

To understand how the C6 expansion procedure affects the transcriptome of PP cells, we performed RNA-Seq analyses on dPP cells (p0) derived from independent differentiations and the corresponding p5 and p10 ePP cells. Principal component analyses (PCA) showed that the main component, PC1, represented most (81%) of the variance among samples and clearly separated p0 PP cells from either p5 and p10 ePP cells. All expanded cells were clustered remarkably close together on the PC1 axis (Figure S5A), suggesting rapid stabilization of the ePP transcriptome after just five passages. This was confirmed by correlation analyses of the transcriptome profiles showing that all major changes occurred between p0 and p5 (Figure 5A, B, S5B). Results from Gene Ontology (GO) and Kyoto Encyclopedia of Genes and Genomes (KEGG) analyses of DEGs between p0 and p10 were consistent with a cell adaptation to culture conditions, the signaling molecules used for the expansion and an effect on the differentiation process (Figure 5C, S5C). We then compared the transcriptome of our VTN-N ePP cells with that of ePP cells expanded on feeders (Ma et al., 2022), on fibronectin (FN) (Nakamura et al., 2022) as well as with that of human fetal PPs (Ramond et al., 2018). Initial PCA analyses suggested that all *in vitro* derived PP cells clustered away from fetal cells (Figure S5D). Thus, we subsequently restricted subsequent comparisons only among *in vitro* derived PP cells.

**Figure 5.**
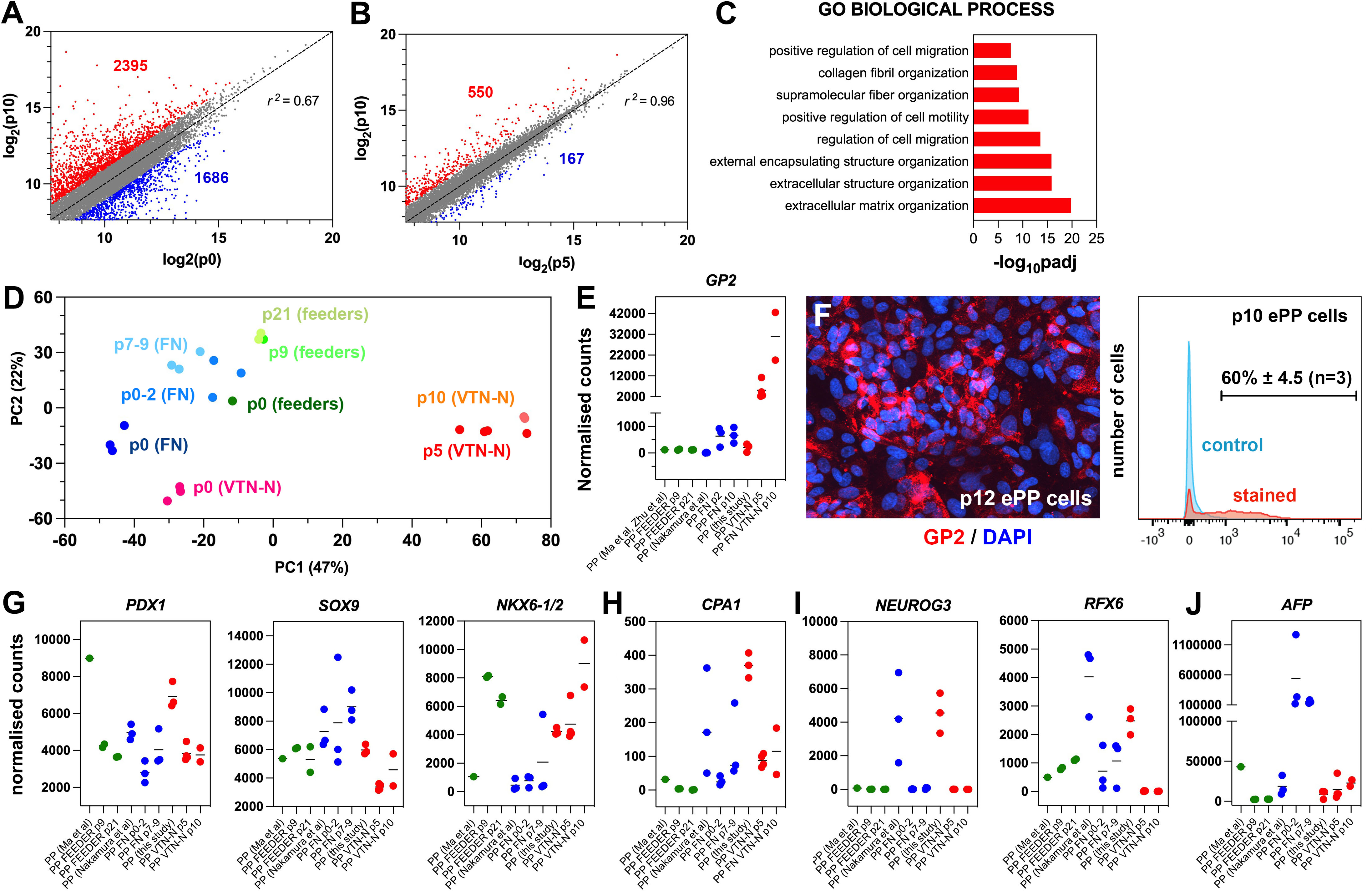
Expansion under C6 stabilizes PP cell identity by repressing differentiation and alternative cell fates. **(A, B)** Correlation analyses of the transcriptome profiles of non-expanded (p0) and p10 expanded PP cells (A) and the transcriptome profiles of p5 and p10 expanded cells (B). The numbers of upregulated and downregulated genes (normalized counts ≥ 200 and 0.5 ≥ FC ≥ 2) are shown in red and blue, respectively and r is the correlation coefficient. **(C)** Mostly affected biological processes between p0 and p10. **(D)** PCA of feeder expanded cells and corresponding p0 cells (shades of green), fibronectin (FN) expanded cells and corresponding p0 cells (shades of blue), as well as vitronectin-N (VTN-N) expanded cells and corresponding p0 cells (shades of red). Darker shades correspond to earlier passages. **(E)** Comparative expression levels of *GP2* in normalized RNA-Seq counts. **(F)** GP2 immunofluorescence of p12 expanded PP cells and FC of p10 ePP cells. **(G-J)** Comparative expression levels of progenitor (G), MPC (H), endocrine (I) and liver (J) markers in normalized RNA-Seq counts of ePP cells and their corresponding p0 cells expanded on feeders (green), FN (blue) and VTN-N (red).

In comparative PCA analysis, with the other *in vitro* derived ePP cells (Ma et al., 2022; Nakamura et al., 2022), our ePP cells clustered separately with very high similarity among p5 and p10 ePP cells (Figure 5D). To identify the molecular basis of this difference, we used a variance stabilizing transformation (vst) function (Love et al., 2014) and hierarchical clustering to identify genes that are differentially regulated in our ePP cells (Figure S5E). Interestingly, these consisted only of upregulated genes and GO analyses showed that affected categories referred to epithelial maintenance and differentiation, cellular signaling response and metabolic processes (GO BP), membrane components (GO CC) and metabolism (GO MF and KEGG) (Figure S5F). An interesting upregulated gene in our ePP cells was *GP2*, a gene encoding a zymogen granule membrane glycoprotein that has been identified as a unique marker of human fetal pancreatic progenitors (Cogger et al., 2017; Ramond et al., 2017). Importantly, GP2^+^ enriched hPS cell-derived PP cells are more efficiently differentiating into pancreatic endocrine cells (Aghazadeh et al., 2022; Ameri et al., 2017). Direct comparison of *GP2* expression among ePP cells showed that our expansion method resulted in a progressive strong *GP2* upregulation and expression. This expression was nearly 50-fold higher than expression in the FN ePP cells at p10 whereas feeder ePP cells did not express this marker to any appreciable extent (Figure 5E, Table S5). Robust GP2 expression in our ePP cells was confirmed by immunofluorescence and flow cytometry where 60% ± 4.5% (n=3) of the ePP (p10-12) cells scored positive (Figure 5F).

To further understand the differences among ePP cells we compared the expression of several additional pancreatic gene markers. *PDX1* expression was generally reduced after expansion but remained at comparable levels among the three procedures (Figure 5G, Table S5). *SOX9* expression remained stable in the feeder ePP cells and increased in the FN PP cells, whereas it was downregulated in our ePP cells (Figure 5G, Table S5). Because SOX9 is also a major driver of the ductal pancreatic program we assessed whether higher SOX9 levels might be associated with higher levels of this program. We examined the collective expression of TFs driving the duct program, such as *PROX1*, *HES1*, *GLIS3* and *ONECUT1*. During development, these genes are initially expressed in bipotent progenitors and then to duct progenitors and differentiated duct cells but they are excluded from the endocrine compartment (Bastidas-Ponce et al., 2017). Consistent with the stronger *SOX9* expression, overall expression levels of these TFs were higher in FN ePP cells suggesting that the ductal program was more active in these cells (Figure S5G, Table S5). During pancreas development, both *NKX6-1* and *NKX6-2* are expressed in progenitor cells, acting in concert to define bipotent progenitors and to subsequently specify endocrine cells (Binot et al., 2010; Henseleit et al., 2005; Nelson et al., 2007; Pedersen et al., 2005; Schaffer et al., 2010). The antibody we used most likely recognizes both proteins, given the very high similarity of NKX6-1 and NKX6-2 at the region of the antigen (Figure S5H). Consistent with the flow cytometry experiments, our expanded cells showed that the combined *NKX6-1/2* expression was strongly upregulated and similar to that of feeder ePP cells (Figure 5G, Table S5). Interestingly, that was due to a strong, expansion-dependent upregulation of *NKX6-2* (Figure S5I, Table S5). Here it is important to note that whereas *NKX6-1/2* expression in feeder and our ePP cells is strong in all samples analysed, it is very low in all FN ePP samples with the exception of a single sample (Figure 5G, Table S5). This suggests a lack of reproducibility in the expansion of PDX1^+^/NKX6-1^+^ FN ePP cells. The expression of other progenitor TF genes such as *FOXA2* and *RBPJ* remained comparable in all expanded cells, although *FOXA2* retained higher levels in the FN ePP cells and *RBPJ* retained higher levels in our ePP cells (Figure S5I, Table S5).

Gene expression analyses, during expansion in the C5 medium, showed complete repression of *PTF1A* expression during the expansion (Figure S2A). This was also observed in the C6 expansion medium (Figure S4A). During development, Nkx6-1/Nkx6-2 act antagonistically to Ptf1a to downregulate its expression and promote the conversion of multipotent progenitor cells (MPCs) into bipotent cells (BP) cells (Schaffer et al., 2010). Thus, we asked whether PP expansion promoted BP identity (PDX1^+^/SOX9^+^/NKX6-1^+^/PTF1A^-^ cells) to the expense of MPC (PDX1^+^/SOX9^+^/NKX6-1^+^/PTF1A^+^) identity. The RNA Seq analyses showed that *PTF1A* repression was a common feature of all expansion procedures (Figure S5J, Table S5). Expression of *CPA1*, a marker of MPCs (Zhou et al., 2007) was also reduced during expansion, further supporting a transition to BP identity. This downregulation was notable in the FN and VTN-N expanded cells because *CPA1* expression of the corresponding p0 cells was substantially higher (Figure 5H, Table S5). We also found that *DCDC2A*, a BP progenitor marker (Scavuzzo et al., 2018) was expressed in both FN ePP and VTN-N ePP cells, but not in feeder ePP cells (Figure S5J, Table S5). These data suggested that ePP cells, particularly FN and VTN-N ePP cells, resemble to BPs rather than MPCs.

NEUROG3^+^ pancreatic endocrine progenitors, divide very rarely, if at all and employ a feed-forward mechanism for their differentiation (Azzarelli et al., 2017; Krentz et al., 2017; Wang et al., 2008). Therefore, it is expected that an efficient PP expansion procedure would efficiently repress the endocrine program and loss of PP cells by differentiation towards the endocrine lineage. Indeed, a common feature of all expansion procedures was the repression of the endocrine differentiation program. Key TFs such as *NEUROG3* and its downstream effectors *NEUROD1*, *NKX2-2*, and *INSM1* were essentially switched off with very similar efficiency in all PP cells, however, it should be noted that expression of these genes was already very low in p0 cells in the Ma et al protocol (Ma et al., 2022). On the other hand, expression levels of *RFX3* and *RFX6* was generally higher in the feeder and FN ePP cells (Figure 5I, S5K, Table S5). Expression of terminal endocrine differentiation markers in all ePP cells was also negligible, particularly at late passages (Table S5).

Expression of acinar TFs such as *BHLHA15* and *RBPJL* was virtually undetectable in all expanded cells (Table S5). Finally, we compared the expression of liver and gut markers. Feeder ePP cells retained only negligible expression of the liver markers *AFP* and *HHEX*. However, expression of these markers was nearly 7-fold and 10-fold higher, respectively, in the FN ePP cells, as compared to our ePP cells, suggesting an overall decreased propensity to endocrine differentiation (Figure 5J, S5L, Table S5). Finally, expression of the gut marker *CDX2* was slightly higher in the feeder ePP cells but generally comparable in all three methods (Figure S5L, Table S5).

In summary, the comparative transcriptome analyses suggested that ePP cells transition from the MPC to the BP state and that our C6 expansion procedure is more efficient at strengthening the PP identity and efficiently repressing the initiation of endocrine differentiation and alternative liver fate.

### Expansion under C6 selects PDX1^+^/SOX9^+^/NKX6-1^+^ cells derived from either XX or XY hPS cells

Different hPS cell lines may vary in their differentiation efficiency and this complicates the clinical implementation of this technology. C6 expansion resulted in the selection of PDX1^+^/SOX9^+^/NKX6-1^+^ cells and this would be advantageous for iPS cell lines that may not differentiate as efficiently. To assess whether the C6 expansion condition, established using the male H1 human embryonic stem (hES) cell line, was similarly applicable to other hPS cell lines of either sex. To address this, we used the female H9 hES cell line, a line that has a preference for neural rather than endoderm differentiation, and a male iPS cell line derived in the CRTD (CRTD1, hPSCreg: CRTDi004-A) with unknown lineage preference. Cells were differentiated into PP cells in monolayer culture (Table S4) and PP cells were then expanded in C6 for at least ten passages. The comparison of the growth curves of H1-(H1-PP), H9-(H9-PP) and CRTD1-(CRTD1-PP) derived ePP cells did not suggest statistically significant differences. A T_d_ of 2.3 and 2.2 days for H9-PP and CRTD1-PP derived ePP cells, respectively, was calculated. Expansion in VTN-N coated cell culture surface appeared at least equally efficient as the expansion on Matrigel for H9- and CRTD1-PP cells (Figure 6A, B).

**Figure 6.**
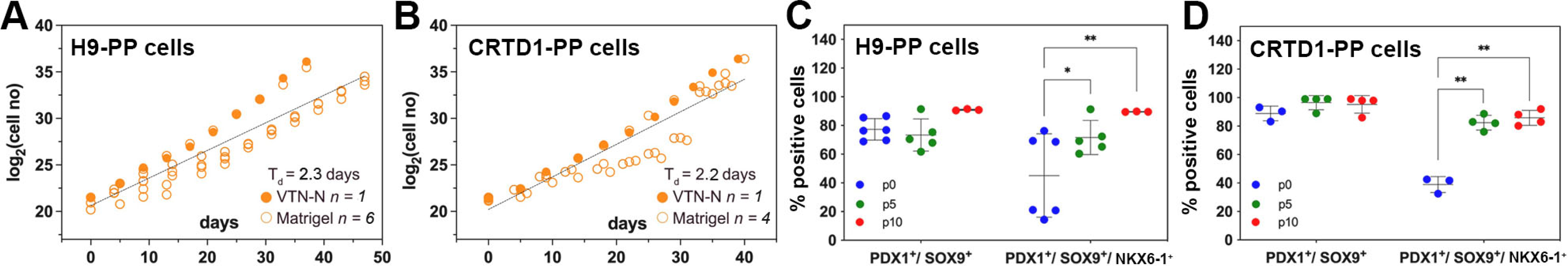
Expansion of H9-derived and CRTD1-derived PP cells under C6. **(A, B)** Growth curve and regression analysis of the expansion of H9-derived PP cells (A) and CRTD1-derived PP cells (B). **(C, D)** Flow cytometry analysis for PDX1^+^/SOX9^+^ and PDX1^+^/SOX9^+^/NKX6.1^+^ cells during the expansion under C6 of H9-derived PP cells (C) and CRTD1-derived PP cells (D) at p0, p5 and p10.

We then compared the maintenance of the PP identity during the expansion of H9-PP and CRTD1-PP cells to that of H1-PP cells using qPCR to quantify gene expression levels at p0, p5 and p10. Expression of *PDX1*, *NKX6-1* and *SOX9* in H9-derived PP cells was strikingly similar to that of H1-derived PP cells at p0 and also following expansion. Expression of these markers in CRTD1-PP cells diverged with decreased *PDX1* expression, at p0 and p10, but increased *NKX6-1* expression at p5 and p10 (Figure S6A-C). Expression of *FOXA2* and *PTF1A* in p0 H9-PP and CRTD1-PP ePPs was also very similar to that in H1-PP cells with the exception of a transient increase of *FOXA2* expression at p5 (Figure S6D, E). Expression of *AFP* and *CDX2* was also very similar in H9-PP and CRTD1-PP cells as compared to H1-PP cells except for a transient *CDX2* upregulation in CRTD1-PP cells at p5 (Figure S6F, G).

We then evaluated the presence of PDX1^+^/SOX9^+^ and PDX1^+^/SOX9^+^/NKX6-1^+^ cells in H9-PP and CRTD1-PP cells at p0, p5 and p10 using flow cytometry. H9 cells were less efficiently differentiated into PDX1^+^/SOX9^+^ cells in comparison to either H1 or CRTD1 cells giving rise to 77%±8% (n=6) as opposed to 98%±2% (n=5) and 89%±5% (n=3) PDX1^+^/SOX9^+^ PP cells for H1- and CRTD1-PP cells, respectively. However, this percentage increased to 91% (n=3) by p10, similar to that for H1-(98%±1%) (n=5) and CRTD1-(95%±6%) (n=4) p10 ePP cells (Figure 4H, 6C, D). Importantly, the percentage of PDX1^+^/SOX9^+^/NKX6-1^+^ H9-PP and CRTD1-PP cells increased from 45%±29% and 39%±6% at p0 to 90% and 85%±6% at p10, respectively. This was a very similar selection to that observed for H1-derived PP cells, which was from 48%±11% at p0 to 89%±13% at p10 (Figure 4H, 6C, D). Chromosomal stability, following expansion, was confirmed also for these lines, since no alterations were found after analysis of the G-banding of at least 20 metaphases for each line at p12 (H9-PP cells) or p13 (CRTD1-PP cells) (Figure S6H, I).

### All C6 ePP cells differentiate into SC-islets containing functional β-cells

Having established that our PP expansion procedure is efficient across PP cells derived from different hPS cell lines, we asked whether ePP cells can be differentiated equally efficiently into SC-islets. H1 dPP cells as well as H1, H9 and CRTD1 ePP cells, expanded for at least ten passages, were clustered in microwells and differentiated using an adaptation of published media (Mahaddalkar et al., 2020; Rezania et al., 2014; Shi et al., 2017) (Table S4) to generate SC-islets. Both H1 dPP and ePP cells gave rise to similar clusters containing INS^+^/NKX6-1^+^, INS^+^/MAFA^+^, GCG^+^ and SST^+^ endocrine cells (Figure 7A, B and S7A, B) as well as similar percentages of INS^+^, INS^+^/GCG^+^ and GCG^+^ cells as determined by flow cytometry (Figure 7C, S7C, D). The total number of INS^+^ and GCG^+^ cells was between 50% and 55% in both H1 dPP- and ePP-derived SC-islets (Figure 7C). We also compared the differentiation propensity of H1 dpp and ePP cells after their differentiation into SC-islets by evaluating the expression of the duct marker *KRT19*. Its expression was significantly lower in ePP cells but it should be noted that in both cases expression wase extremely weak (Figure S7E).

**Figure 7.**
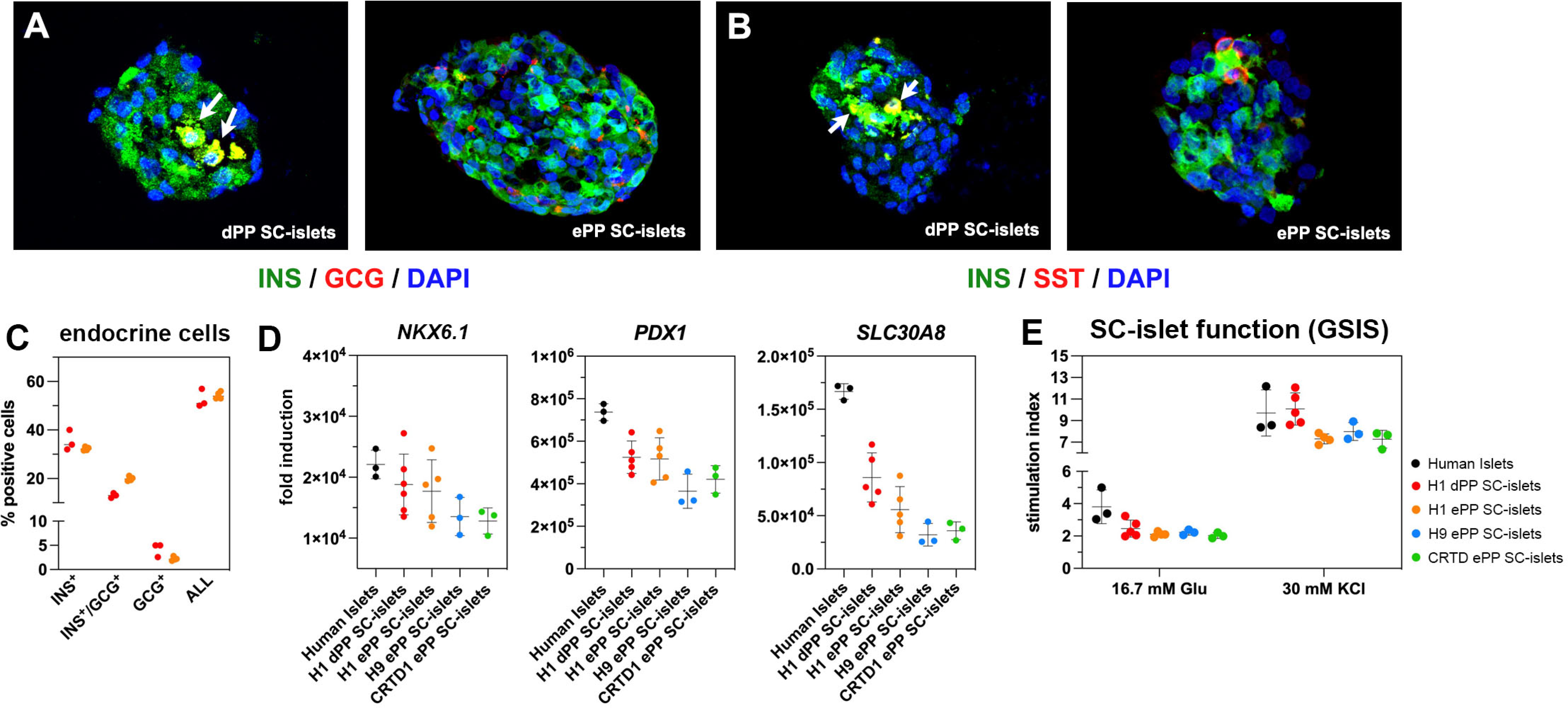
Differentiation of ePP cells into SC-islets containing functional β-cells. **(A, B)** Immunofluorescence analysis of SC-islets derived from p0 PP cells (dPP) or expanded PP cells for at least ten passages (ePP) for INS and GCG expression (A) or INS and SST expression (B). **(C)** Percentages of INS^+^, INS^+^/GCG^+^ as well as GCG^+^ cells in SC-islets derived from dPP or ePP cells as determined by flow cytometry. **(D)** Expression levels of *NKX6-1*, *PDX1* and *SLC30A8* as determined by qPCR and expressed as fold induction relative to expression levels in hPS cells. **(E)** Secretion of C-peptide following sequential stimulation by 16.7 mM glucose and 16.7 mM glucose / 30 mM KCl (30 mM KCl) after exposure in basal conditions with 2.8 mM glucose. Stimulation index is the ratio of secretion under these conditions to secretion in basal conditions. Horizontal lines represent the mean ± SD. Statistical tests were two-way ANOVA with Tukey’s test, using p0 as the control condition for the comparison with *p* ≤ 0.033 (*), *p* ≤ 0.002 (**), ≤ 0.0002 (***) and ≤ 0.0001 (****). Scale bar corresponds to 50 μm. Scale bar corresponds to 100 μm (E, F).

Gene expression levels for differentiated endocrine markers including β-cell markers such as *PDX1*, *NKX6-1*, *SLC30A8* (Figure 7D), *INS*, *MAFA* as well as *GCG* and *SST* (Figure S7F) were very similar between H1 dPP derived SC-islets and SC-islets derived from either H1, H9 or CRTD1 ePP cells. Expression of *PDX1*, *NKX6-1*, *SLC30A8* and *SST* was comparable between human islets and dPP or ePP SC-islets (Figure 7D, S7F) but expression of *INS*, *GCG* and *MAFA* was substantially higher in human islets reflecting the less advanced maturation of SC-islets (Figure S7F).

Finally, we assessed the functionality of the β-cells in H1 dPP-as well as H1, H9 and CRTD1 ePP-derived SC-islets in static GSIS assays where SC-islets were sequentially incubated in Kreb’s buffer containing low glucose levels (2.8mM), Kreb’s buffer containing high glucose levels (16.7 mM) and finally a depolarizing Kreb’s buffer containing high glucose levels (16.7 mM) and KCl (30 mM). As a comparison, human islets were processed in a similar manner. Human C-pep levels were measured from the supernatant of successive incubations and used to calculate the fold stimulation. These experiments showed that dPP- and ePP-derived SC-islets contained β-cells of very similar functionality (Figure 7E) and, as expected, close, but not similar, functionality to human islets. Importantly, the amount of secreted human C-peptide under all different conditions was very similar among dPP and ePP derived SC-islets (Figure S7G).

In summary, both dPP and ePP cells, derived from different hPS cell lines, differentiate into SC-islets with essentially the same efficiency and contain β-cells of similar functionality.

## DISCUSSION

The unlimited expansion of progenitor cells holds significant therapeutic promise but remains an important challenge in regenerative medicine. Progenitor cells are dynamic entities and the key objective in these efforts is to uncouple survival and proliferation from widely employed feed-forward mechanisms that promote their differentiation. The latter are not exclusively regulated from extrinsic signals but rely to a large extent on internal regulators as well as autocrine signals. Thus, the hallmarks of efficient progenitor expansion would be the maintenance of key progenitor features, the efficient suppression of differentiation programs and alternative lineages and the capacity to efficiently differentiate under appropriate conditions. For therapeutic applications, such expansion should also be efficient, reproducible, applicable across different cell lines and compatible with chemically defined culture media.

Regarding pancreatic development in particular, several feed-forward networks have been documented with exquisite detail (Arda et al., 2013). In the early pancreatic progenitors, a Sox9/Fgf feed-forward loop is essential to escape liver fate and promote pancreas identity and expansion (Seymour et al., 2012). Fgf10 was identified as a possible extrinsic signal but it was shown that it also promotes liver identity (Hart et al., 2003; Jung et al., 1999; Norgaard et al., 2003; Rossi et al., 2001). Ptf1a initially forms heterodimers with Rbpj to promote expansion of MPCs and activate transcription of Rbpjl (Masui et al., 2010). Expression of PTF1A in the MPCs is essential to set in motion an epigenetic cascade that is required for subsequent duct and endocrine differentiation (Miguel-Escalada et al., 2022). Rbpjl eventually replaces Rbpj in its complexes with Ptf1a and the Ptf1a. Rbpjl complexes promote the specification of acinar progenitors (Masui et al., 2010). Their mutual cross-repression with Nkx6-1/2 define the acinar progenitors and BPs, respectively (Schaffer et al., 2010). We found that PP cells at p0 resemble MPCs because they express high levels of *PTF1A* but also *CPA1*, a marker of MPCs (Zhou et al., 2007). Expansion promotes the switch to BP identity as documented by the strong downregulation of both *PTF1A* and *CPA1* and the corresponding upregulation of *NKX6-1/2* as well as *DCDC2A* (Scavuzzo et al., 2018) suggesting that ePP cells may resemble bipotent progenitors as described in mice (Schaffer et al., 2010).

Sphingosine-1-phosphate signaling activates YAP to set the stage for the endocrine differentiation of PP cells (Serafimidis et al., 2017) and the YAP/TEAD complex regulates the enhancer network that maintains PP cells in a proliferative state (Cebola, I. et al. Nat Cell Biol 17, 615-26, 2015). In BP cells, YAP activation mediates a mechanotransduction pathway to maintain cells in a proliferative state and its inactivation is a prerequisite for endocrine differentiation (Cebola et al., 2015; Rosado-Olivieri et al., 2019). The latter mechanism was exploited to improve the in vitro pancreatic β-cell differentiation (Hogrebe et al., 2020). As expected, gene expression of all the Hippo pathway components is strong in PP cells but there is no apparent regulation of the pathway, at least at the gene expression level, during expansion. Upon YAP inactivation and endocrine commitment, the TFs Myt1 and Neurog3 form a feed-forward loop to promote the final commitment into the endocrine lineage (Wang et al., 2008). Differentiating cells are dynamic cell populations that do not move synchronously through the successive differentiation stages and this can explain the substantial expression levels of the TFs of the endocrine programme in PP cells. The expansion procedure we propose was successful in freezing cells in a state preceding endocrine commitment as shown by the dramatic repression of all endocrine TFs.

We hypothesized that the expression kinetics of signals and signal receptors during expansion under CINI would give us indications on which signaling pathways should be either repressed or activated to promote reproducible expansion. This approach was successful and we favor the hypothesis that these are autocrine signals and receptors expressed in the PP cells themselves because this modulation remains necessary even at late passages when the percentage of PDX1^+^/SOX9^+^/NKX6-1^+^ is around 90%. However, we cannot exclude the possibility that some or even all of the inhibited signals, including the suppression or RA signaling through the withdrawal of vitamin A, may come from non-PP cells present in the culture. In this case their action would be to block expansion and/or survival of alternative cell fates. A convincing answer to this question should await single cell RNA Seq analyses but it is worth noting that a high initial PP cell density was an essential requirement for robust early expansion for all cell lines. This suggests that autocrine signals play a key role in the process.

We found that a combination of EGF, FGF2 and FGF18 promoted robust expansion of hPS cell derived PP cells. EGF promotes *NKX6-1* activation (Nostro et al., 2015) but on its own it could not support consistent PP cell expansion (CINI, Table S1). FGF2 is a widely used mitogen and whereas it enhanced PP proliferation (C1) it was less efficient than FGF18 (C5, Table S1). There was a clear synergy of the two mitogens in promoting expansion and possibly the enrichment in NKX6-1^+^ cells (C6, Table S1). NKX6-1 regulates multiple cell cycle genes (Taylor et al., 2015) and thus some of the effects of these mitogens might be indirectly reinforced through NKX6-1. The comparison of early (p3) CINI and C5 cells suggested that higher proliferation in the latter was the main driver of successful expansion. Cell death appeared also lower in C5 cells (17.2 ± 4.7% against 22.1 ± 5.0%) and thus it is likely that the combination of these effects results in reproducible expansion under C5. The high rates of cell death suggest that committed non-PP cells are primarily affected and bona fide PP cells as well as cells still competent to become bona fide PP cells compensate with higher proliferation. This question could be resolved by co-staining of the cells with PP markers as well as cell death and proliferation markers.

PP cells express the enzymes necessary to convert vitamin A into retinoic acid (RA) (Table S3) which, in turn, promotes differentiation of the pancreas progenitors (Lorberbaum et al., 2020; Martin et al., 2005; Ostrom et al., 2008; Vinckier et al., 2020). Thus, eliminating vitamin A from the expansion medium was important in order to uncouple PP expansion from differentiation. The TGFβ signaling pathway plays a complex role in the induction, maintenance and endocrine differentiation of pancreas progenitors (Guo et al., 2013; Sanvito et al., 1994; Spagnoli and Brivanlou, 2008; Tulachan et al., 2003) and this was reflected in the widely variable gene expression kinetics of receptors and ligands during CINI expansion. Accordingly, our experiments suggested that the highly specific ALK5i II inhibitor (Sanvitale et al., 2013) (C5, C6) was more efficient than the less specific A83-01 inhibitor (Tojo et al., 2005) (CINI) in promoting PP expansion. Further inhibition of the pathway with the addition of LDN193198, which targets ALK3 more efficiently than ALK5i II (Gellibert et al., 2004), dramatically reduced PP expansion (C7, Table S3). We documented that preferential expansion of progenitors, rather than cell survival, was the main mechanism of the expansion, in agreement with the idea that the expansion medium effectively decoupled proliferation from differentiation. Finally, the addition of the canonical Wnt inhibitor IWR-1 (Chen et al., 2009) significantly reduced AFP expression and promoted an efficient selection of NKX6-1^+^ cells, without affecting the proliferation rate, presumably by reducing the branching of expanding cells towards the hepatic fate.

The molecules employed in our C6 expansion medium are not, with the partial exception of FGF-18, foreign to the differentiation conditions used to generate SC-islets. RA has been used to induce the primitive gut tube conversion to PP cells and ALK5i II has been to promote the PP to endocrine progenitor conversion (Hogrebe et al., 2021; Rosado-Olivieri et al., 2019) Additionally, IWR-1 has been proposed to increase endocrine differentiation (Sharon et al., 2019). This may appear contradictory but we think that it is the combination of signals that promotes a certain cell state rather than specific signals. Accordingly, we carried out the generation of PP cells before expansion and their differentiation after expansion in exactly the same way as for PP cells that were not expanded. In these procedures we employed RA and ALK5i II as published earlier but in our hands IWR-1 did not make a difference in either the dPP or ePP to endocrine progenitor transition.

Culture conditions to expand hPS cell-derived PP cells have recently been reported (Ma et al., 2022; Nakamura et al., 2022) but suffer from a number of disadvantages as outlined in the introduction. An important feature of our expansion procedure is the reproducible enrichment in PDX1^+^/SOX9^+^/NKX6-1^+^ which can reach 90%, irrespective of the cell line used. Thus, this procedure will be particularly useful for hPS cell lines with reduced initial capacity to differentiate into PP cells. The comparison of the transcriptome profiles showed that our procedure is unique in effecting a strong upregulation of *GP2*, a unique marker of human fetal PP cells (Cogger et al., 2017; Ramond et al., 2017) associated with a high PP preference for endocrine differentiation (Aghazadeh et al., 2022; Ameri et al., 2017). Another unique feature, was the strong upregulation of *NKX6-2*, another marker of PP cells which complements *NKX6-1* function but is not retained in differentiated endocrine cells (Binot et al., 2010; Henseleit et al., 2005; Nelson et al., 2007; Pedersen et al., 2005; Schaffer et al., 2010). The FN procedure appeared to maintain higher levels of TF genes engaged in the duct program, as well as higher levels of *AFP* and *HHEX*, markers of the liver lineage. Expression of acinar *TFs* was very low in all three procedures and, as expected, the expression of *TFs* driving the endocrine program was strongly reduced in all three procedures albeit our procedure appeared more efficient in this respect, particularly with regard to *RFX3* and *RFX6* expression. It is important to point out that all three procedures resulted in noticeable upregulation of *CDX2*, a gut TF and future efforts should be directed in addressing this, using modifications of the current procedure and FC quantification of CDX2^+^ cells. Interestingly, all ePP cells clustered separately from human fetal progenitors. This could reflect genuine differences due to culture conditions or differences in developmental age, including the possibility that ePP cells may represent a transient, unstable state in vivo. Another possibility is that PP cells in vivo might be more heterogeneous as it is expected that progenitors at different positions in the developing organ would be at different developmental stage.

Importantly, ePP cells from H1, H9 and CRTD1 hPS cells all differentiated with similar efficiency to dPP cells into SC-islets containing similar numbers of β-cells of comparable functionality. This may appear surprising because ePP cells contained a much higher percentage of PDX1^+^/SOX9^+^/NKX6-1^+^ cells. It should be noted, however, that the liver marker AFP and, particularly, the gut marker CDX2 were upregulated, whereas expression of *PTF1A*, recently shown to promote endocrine differentiation of hPS cells (Miguel-Escalada et al., 2022), was essentially lost. The same pattern, and even higher upregulation of gut and liver markers, was also seen in the other expansion procedures (Ma et al., 2022; Nakamura et al., 2022).

The unlimited expansion of PP cells reported here is applicable to different hPS cell lines and presents several advantages in the efforts to scale-up the generation of islet cells, including β-cells, for the cell therapy of diabetes. It reduces the number of differentiation procedures to be carried out starting at the hPS cell stage, thus eliminating a source of variability and allows the selection of the most optimally differentiated PP cell population for subsequent expansion and storage. Since it is currently acknowledged that current differentiation procedures do not produce fully functional β-cells, these ePP cells will provide a convenient springboard to refine downstream differentiation procedures. Suitable surface markers for the selection of endocrine cells at the end of the differentiation procedure have been reported (Li et al., 2020; Veres et al., 2019) but such procedures may prove too expensive in a clinical setting. Alternatively, expanded PP cells could be directly used for transplantations as it has been shown that they can mature in vivo (Kroon et al., 2008; Shapiro et al., 2021). Whether PP cells or terminally differentiated SC-islet cells would be the best approach in future transplantations, is still discussed but, in any of these cases, the availability of a highly pure GMP-grade, hPS cell derived PP cell population has several advantages as discussed above.

The transition to full GMP conditions is expected to be relatively straightforward. Human iPS cell lines are already being generated under GMP conditions and, depending on the culture medium, the first key differentiation step (monolayer hPS cell to definitive endoderm) might need adjustment regarding seeding density, the length of time that hPS cells remain as a monolayer, confluency at the time of differentiation initiation and possibly concentrations of signals, particularly Activin A and CHIR. In our hands, the plate coating, either with Matrigel or GMP-compatible reagents such as FN or VTN-N, does not have any impact on the differentiation efficiency to definitive endoderm and subsequent stages. Otherwise, the large majority of supplements, used in hPS cell culture and differentiation into SC-islets, are small molecules and recombinant proteins which can be already purchased as GMP reagents. Some small molecules are not yet available as GMP versions but they could be manufactured upon custom request if necessary.

In summary, the expansion of PP cells will facilitate the generation of unlimited number of endocrine cells, initially for studying diabetes, drug screening for personalized medicine and eventually cell therapies. The chemically defined expansion procedure we report here will be an important step toward generating large numbers of human pancreatic endocrine cells that are of great interest for biomedical research and regenerative medicine.

## Author contributions

A.G. conceptualized the study, L.J., M.B., E.Z, V.K., R.S., D.P and I.G. performed experiments, A.G., L.J., and M.B. analyzed the data, A.G. and M.L. analyzed the transcriptome data, S.K. and K.N. generated and expanded the CRTD1 line, B.L. provided the human islets, A.G. wrote the manuscript with input and editing from L.J. and M.B., A.G. supervised and acquired funding.

## Supporting information

All sup figures with legends

All sup tables

## Acknowledgments

Research in the AG laboratory was supported by grants from the German Center for Diabetes Research (DZD)(grant 82DZD00101) and the German Research Foundation (DFG)(grants GA-2004/3-1 and IRTG 2251). Fibroblasts “Theo”, used for the generation of CRTD1 hiPSC, were a gift from Prof. Dr. med. Min Ae Lee-Kirsch, University Hospital Carl Gustav Carus, Dresden.

## MATERIALS AND METHODS

### Derivation of the CRTD1 human iPS cell line

The CRTD1 human iPS cell line (hPSCreg: CRTDi004-A) was generated from previously published foreskin fibroblasts (termed Theo) of a consenting healthy donor (Wolf et al., 2016). Isolation of cells and reprogramming to hiPS cells was approved by the ethics council of TU Dresden (EK169052010 und EK386102017). Theo fibroblasts were reprogrammed at the CRTD Stem Cell Engineering Facility at the Technical University of Dresden using the CytoTune-iPS 2.0 Sendai Reprogramming Kit (Thermo Fischer Scientific A16517) according to the supplier’s recommendations for transduction. Following transduction with the Sendai virus, cells were cultured on irradiated CF1 Mouse Embryonic Fibroblasts (Thermo Fisher Scientific 15943412) in KOSR-based medium containing 80% DMEM/F12 (Thermo Fisher Scientific 31330095), 20% KnockOut Serum Replacement (Thermo Fisher Scientific 10828028), 2 mM L-glutamine (Thermo Fisher Scientific 25030024), 1% Nonessential Amino Acids (Thermo Fisher Scientific 11140035) and 0,1 mM 2-mercaptoethanol (Thermo Fisher Scientific 21985023) supplemented with 10 ng/ml human FGF2 (Stem Cell Technologies 78003). Individual iPSC colonies were mechanically picked, expanded as clonal lines and adapted to Matrigel (Corning 354277), mTeSR1 (Stem Cell Technologies 85850) and ReLeSR (Stem Cell Technologies 05873) conditions after several passages. Master and working hiPS cell banks were established from clones with the best morphology.

To characterize the newly generated CRTD1 hiPS cell line, several tests were performed, accessible at https://hpscreg.eu/cell-line/CRTDi004-A. Pluripotency was initially analyzed by Alexa Fluor 488 conjugated anti-Oct3/4 (BD Pharmingen 560253), PE conjugated anti-Sox2 (BD Pharmingen 560291), V450 conjugated anti-SSEA4 (BD Pharmingen 561156), and Alexa Fluor 647 conjugated anti-Tra-1-60 (BD Pharmingen 560122) used according to the manufacturer’s recommendations and analyzed on a BD LSRII Flow Cytometer. Then, the three germ layer differentiation assay was performed as described previously (Cheung et al., 2011) and resulting cells were stained using the 3-germ layer Immunocytochemistry Kit (Thermo Fisher Scientific A25538) according to the instruction manual. For endoderm differentiation, a SOX17 primary antibody (Abcam ab84990) followed by Alexa Fluor 488 goat anti-mouse IgG (Thermo Fisher Scientific A11001) was used. Quantitative RT-PCR for pluripotency and trilineage spontaneous differentiation was performed according to the instruction manual of the human ES cell Primer Array (Takara Clontech).

Cells were analyzed for chromosomal abnormalities using standard G banding karyotyping. Cells were treated with 100 ng/ml KaryoMAX Colcemid solution (Thermo Fisher Scientific 15212012) for 4 h at 37°C, harvested and enlarged with 0.075 M KCl solution (Thermo Fisher Scientific 10575090) for 20 min at 37°C. After fixing with 3:1 methanol (VWR 20846.326): glacial acetic acid (VWR 20102.292), cells were spread onto glass slides and stained with Giemsa at the Institute of Human Genetics, Jena University, Germany. G-bandings of at least 20 metaphases were analyzed.

### Isolation of human pancreatic islets

Human islets were obtained through the XXX Islet Transplantation Program approved by the XXX Institutional Review Board (EK 255062022) with written informed consent obtained from each islet donor participant. Islets were isolated and purified from resected pancreas tissue according to a modified Ricordi method. Briefly, Collagenase, neutral protease (Serva Electrophoresis, Heidelberg, Germany), and Pulmozyme (Roche, Grenzach, Germany,) were infused into the main pancreatic duct. Islets were separated from exocrine tissue by centrifugation on a continuous Biocoll gradient (Biochrom AG, Berlin, Germany) in a COBE 2991 cell processor (Lakewood, CO, USA).

### Maintenance and karyotyping of human pluripotent stem cell lines

The H1 and H9 hES cell lines were purchased from WiCell (Wisconsin, USA). H1 and H9 hES cells as well as CRTD1 iPS cells were maintained on cell culture dishes coated with hES cell qualified Corning Matrigel (BD Bioscience, 354277) diluted 1:50 with DMEM/F-12 (Gibco, 21331-020) and daily changes of mTeSR1 medium (STEMCELL Technologies, 85850) supplemented with 1x penicillin/streptomycin (Gibco, 15140-122). The cells were passaged at around 70% confluency, approximately every 4 days at a ratio of 1:6 to 1:9, as small aggregates using ReLeSR (STEMCELL Technologies, 05872). Karyotyping for H1 and H9 cells was as described above for the CRTD1 iPS cell line and cells were routinely tested for mycoplasma contamination by PCR as published previously (Young et al., 2010).

### Differentiation of hPS cells to PP cells

Initially, the H1 ES cell line was differentiated to PP cells using the STEMdiff™ Pancreatic Progenitor Kit (STEMCELL Technologies, 05120) according to the manufacturer’s instructions. In short, the hES cell colonies were dissociated into single cells using TrypLE Express (Gibco, 12604-013) and seeded on Matrigel coated plates as described above at a concentration of 95,000 cells/cm^2^ in mTeSR supplemented with 20 µM ROCKi. The medium was replaced the next day by mTeSR and differentiation was initiated by replacing it with the S1d1 differentiation medium when cells were 60-70% confluent, typically two days after the initial seeding. Daily washes with DPBS (Gibco, 14190250) and media changes were done until S4d5 when the cells reached the end of PP stage. The monolayer of PP cells was then dissociated using Accumax^TM^ (STEMCELL Technologies, 07921) and cells were used for expansion under INI conditions.

Later, H1, H9 and CRTD1 iPS cells were differentiated into PP cells using an adaptation of published procedures (Mahaddalkar et al., 2020; Rezania et al., 2014; Shi et al., 2017) (Table S4). The monolayer of PP cells was then dissociated using TrypLE Express (Gibco, 12604-013) for 2 minutes and cells were used for expansion under C0 to C8 conditions.

### Expansion and cryopreservation of PP cells

The monolayer of PP cells was dissociated using TrypLE Express (Gibco, 12604-013) and cells were used for expansion. Expansion cultures were maintained on polystyrene cell culture plates (Corning, CLS3516) coated with Matrigel (Corning, 354277) or Cultrex (R&D systems, 3434-005-02), both diluted 1:50 in DMEM/F-12 or with 20 ug/mL recombinant truncated vitronectin (VTN-N) (Thermo Fischer Scientific, A31804) diluted in DMEM/F-12 (Gibco, 21331-020) for 1 hr at room temperature. PP cells were resuspended in PP expansion media CINI-C8 (Table S1) and seeded initially at a density of 3.2 × 10^5^ / cm^2^. In subsequent passages cells were seeded at a density of 2.1 × 10^5^ / cm^2^. In every passage, expansion media were supplemented, during the first day, with 10 µM ROCKi. The expansion medium was changed daily, and cells were typically passaged every 4^th^ day using TrypLE Express dissociation into single cells. Karyotyping for expanded PP cells and mycoplasma testing was as described above for the hPS cell lines.

Expanding PP cells were routinely frozen at later passages using mFreSR (Stem Cell technologies, 05854), supplemented with 20 μM ROCKi, at a density of 10 million cells/ml. To ensure proper controlled freezing (− 1 °C/min), cryotubes were placed in Mr. Frosty™ Freezing Container (Thermo Fischer Scientific, 5100-0001) at − 80 °C. After 24 hours, cryotubes were transferred to a liquid nitrogen chamber. For thawing, frozen cells were placed at 37°C and then transferred to 6ml DMEM/F-12 at room temperature for centrifugation. After spinning down the cells at 600xg, the pellet was resuspended using PP expansion media and cells were counted using the Countess II Automated Cell Counter (Thermo Fischer Scientific) and Trypan blue. Typical recovery rates were above 85%. Cells were then seeded as described above for expansion.

### Differentiation of CINI ePP cells into pancreatic endocrine cells using ALI culture

PP cells generated using the STEMdiff™ Pancreatic Progenitor Kit and expanded under CINI conditions were dissociated into single cells using Accutase^TM^ (STEMCELL Technologies, 07920) at 37 °C and then resuspended in PEP medium (Table S4) supplemented with 10 µM ROCKi at concentration of 50,000 cells/µl. The Falcon® Cell Culture Inserts (Corning, 353493) were placed in their companion plate wells (Corning, 353502) containing 1.5 ml of complete PEP media supplemented with 10 µM ROCKi. Ten droplets of 5 µl of the cell suspension were dropped on the insert to create ten 3D clusters per well, each containing 250,000 cells. Daily media changes were conducted according to Table S4 with the following changes: (a) during S5, the XX Notch inhibitor (Millipore, 565789) was added at a concentration of 10 nM (b) final glucose concentration at S6 and S7 was 20 mM and (c) no sodium pyruvate was used at S7.

### Differentiation of PP cells into SC-islets using microwells

PP cells were dissociated using TrypLE Express at 37°C for 2-3 mins and seeded in microwells of AggreWell 800 plates (STEMCELL Technologies, 34825), using the recommended procedure by the supplier. Directly differentiated and ePP cells were seeded at a density of 5000 or 2000 cells per microwell, respectively. Media were an adaptation of published procedures (Balboa et al., 2022; Mahaddalkar et al., 2020; Rezania et al., 2014; Shi et al., 2017) (Table S4). Expanded PP cells in micropatterned wells were kept in expansion media for 24 hours and, during the first day of S5, S5 medium was supplemented with one-fourth of the concentration of C6 additional factors. Media of S5-S7 were as described in Table S4 and the medium was changed daily.

### Immunofluorescence analyses

For immunofluorescence (IF), cells were cultured on 12 mm diameter Matrigel-coated coverslips (Carl Roth, P231.1) placed in 12-well wells (Corning, CLS3513) and differentiated or expanded as described above. Cells were then fixed in 4% paraformaldehyde (PFA) for 20 min at 4° C and washed with PBS. Cells were blocked and permeabilized for 1 h at RT using a 5% serum / 0.3% Triton X-100 PBS solution. Samples were then incubated at 4°C in 2.5% serum / 0.3% Triton X-100 in PBS containing the primary antibodies in the appropriate concentration (Table S6). The following day, samples were incubated for 1 h at RT in 2.5% serum / 0.3% Triton X-100 in PBS containing the appropriate secondary antibodies, conjugated with either Alexa 488, 568 or 647, at a 1:500 dilution (Table S7). The coverslip with the stained cells was then placed on a microscope slide, covered with ProLong™ Gold Antifade mounting medium with DAPI (Invitrogen, P36931) and overlayed with a rectangular coverslip.

IF images were acquired using a Zeiss Axio Observer Z1 microscope coupled with the Apotome 2.0 imaging system and with consistent exposure times for the Alexa 488, 568 and 647 channels in between passages and conditions to allow for direct comparison of the signal intensities.

### Fluorescence-activated cell sorting (FACS) of hPS cell derived PP and endocrine cells

Cells were first washed with 2% BSA in DMEM-F12 and then dissociated into single cells using TrypLE Express (Gibco, 12604-013). Following dissociation, the cells were counted using the Countess II Automated Cell Counter (Thermo Fischer Scientific) and Trypan blue. Then, cells were washed with PBS and fixed at a concentration of 4 million/ml in in 4% PFA for 10 min at 4°C. For staining with transcription factor antibodies, cells were washed with PBS and permeabilized using the Foxp3 Transcription Factor Staining Buffer Set (Invitrogen, 00-5523-00) for 1 h in dark at 4°C. Cells were then washed again using the 1x permeabilization buffer. Thereafter, 2×10^6^ cells/sample were blocked using 100 μl of 5% serum in 1x permeabilization buffer and then incubated with primary antibodies at the appropriate concentration (Table S5) in the same buffer overnight at 4°C. Then, they were washed twice with 1x permeabilization buffer and incubated with secondary antibodies at the appropriate concentration (Table S7) in the same buffer at room temperature for 1 hour in dark. For cytoplasmic factors, 2×10^6^ cells/sample were blocked for 30 mins at RT in 100 μl PBS containing 2% serum and 0.3% Triton X-100. Then cells were washed with 0.1% Triton X-100 in PBS solution and incubated with primary antibodies at the appropriate concentrations (Table S5) in the same buffer overnight. Thereafter, cells were washed twice with 0.1% Triton X-100 in PBS solution and incubated with secondary antibodies at the appropriate concentration (Table S7) at room temperature in the dark for 1 hour. For conjugated antibodies, the appropriate number of cells was used, as directed by the manufacturer, for both the isotype control and staining sample. Samples were stained overnight with conjugated antibodies and then washed with 0.1% Triton X-100 in PBS solution. FACS data were acquired using BD FACSCanto™ II and analysed using the FlowJo software.

### Proliferation and cell death assays

The EdU proliferation assay was performed with the Click-iT™ Plus EdU Alexa Flour^TM^ 488 Flow Cytometry Assay Kit (Invitrogen, C10632), that contains all the necessary reagent except PBS and BSA, and according to the kit protocol. In short, on the day of passaging the expanding cells, the expansion media was supplemented with 10 µM EdU for 2 h and cells were then dissociated and washed in 3 ml of 1% BSA in PBS. A pellet of 1 million cells was resuspended in 100 µl of Click-iT™ fixative for 15 min and washed again with 3 ml of 1% BSA in PBS. The pelleted cells were resuspended in 100 µl of permeabilization and wash reagent for 15 min. Following the incubation, 0.5 ml of the Click-iT™ Plus reaction cocktail containing 1X buffer additive, the Alexa Fluor™ 488 picolyl azide and a copper protectant in PBS, was added to the sample and incubated in the dark at RT for 30 min. The cells were then washed with 3 ml of 1X permeabilization and wash reagent. The pelleted cells were resuspended in 0.5 ml of permeabilization and wash reagent, passed through a 40 µm strainer (PluriSelect, 43-10040) and analysed on the BD FASCanto™ II.

The flow cytometry staining for necrosis and apoptosis was was performed with the FITC Annexin V Apoptosis Detection Kit with 7-AAD (Biolegend, 640922). Once the cells were incubated in 10 µM of EdU, they were detached and washed in 1% BSA in PBS and an aliquot of 250,000 cells was taken and pelleted in a 1.5 ml microcentrifugation tube. The cell pellet was resuspended in 100 µl of Annexin V Binding Buffer containing 5 µl of FITC Annexin V and 5 µl of 7-AAD Viability Staining Solution and cells were incubated for 15 min at RT in the dark. After the incubation, an additional 400 µl of Annexin V Binding Buffer was added to the sample and passed through a 40 µm strainer (PluriSelect, 43-10040) before the live cells were analysed on the BD FACSCanto™ II.

### Static glucose-stimulated insulin secretion (GSIS) assay

At the end of S7 (S7d10 – S7d14) 150 clusters were collected, washed with PBS and incubated for 1 hour in 700 μl of fresh Kreb’s buffer (2.5mM CaCl_2_, 129mM NaCl, 4.8mM KCl, 1.2mM MgSO_4_, 1.2mM KH_2_PO_4_, 1mM Na_2_HPO_4_, 5mM NaHCO_3_, 10mM HEPES, 0.1% BS, pH adjusted to 7.4 with 5M NaOH). After first incubation, clusters were incubated with Kreb’s buffer containing low glucose (2.8mM) for 1 hr, then high glucose (16.7mM) for 1 hr and finally KCl/high glucose (30 mM/16.7 mM) for 1 hr. Incubations were for exactly one hour and then supernatant was collected. Collected supernatants and pelleted cells were frozen at −80 °C until the analysis. C-peptide detection was performed using the human C-peptide ELISA kit (Mercodia, 10-1141-01), readings were taken using an ELISA plate reader and the standard curve was generated. Cell pellets were then used to isolate genomic DNA with the DNeasy Blood & Tissue kit (Qiagen, 69504) and DNA quantification was done using a Nanodrop spectrophotometer.

### RNA isolation and qRT-PCR (qPCR)

Total RNA was prepared using the RNeasy kit with on-column genomic DNA digestion (Qiagen, 74004) following the manufacturer’s instructions. First strand cDNA was prepared using the TAKARA PrimeScript RT Master Mix (TAKARA RR036A). Real-time PCR primers (Table S8) were designed using the Primer 3 software (SimGene), their specificity was ensured by *in silico* PCR and they were further evaluated by inspection of the dissociation curve. Reactions were performed with the FastStart Essential DNA Green Master mix (Roche 06924204001) using the Roche LightCycler 480 and primary results were analyzed using the on-board software. Reactions were carried out in technical triplicates from at least three independent biological samples. Relative expression values were calculated using the ΔΔCt method by normalizing to H1 undifferentiated expression levels and the *TBP* housekeeping gene.

### RNA sequencing and bioinformatics analysis

Cells were differentiated and from hPS cells as described above. Three independent samples from distinct differentiations and independent expansions were used as biological replicates. Total RNA prepared as above with an integrity number of ≥ 9 was used and subsequent steps were performed at the Biotec Sequencing Core of TU Dresden. mRNA was isolated from 1 ug of total RNA by poly-dT enrichment using the NEBNext Poly(A) mRNA Magnetic Isolation Module according to the manufacturer’s instructions. Final elution was done in 15ul 2x first strand cDNA synthesis buffer (NEBnext, NEB). After chemical fragmentation by incubating for 15 min at 94°C the sample was directly subjected to the workflow for strand specific RNA-Seq library preparation (Ultra Directional RNA Library Prep, NEB). For ligation custom adaptors were used 1: (Adaptor-Oligo 5’-ACA CTC TTT CCC TAC ACG ACG CTC TTC CGA TCT-3’, Adaptor-Oligo 2: 5’-P-GAT CGG AAG AGC ACA CGT CTG AAC TCC AGT CAC-3’). After ligation, adapters were depleted by an XP bead purification (Beckman Coulter) adding bead in a ratio of 1:1. Indexing was done during the following PCR enrichment (15 cycles) using custom amplification primers carring the index sequence indicated with ‘NNNNNN’. (Primer1: Oligo_Seq AAT GAT ACG GCG ACC ACC GAG ATC TAC ACT CTT TCC CTA CAC GAC GCT CTT CCG ATC T, primer2: GTG ACT GGA GTT CAG ACG TGT GCT CTT CCG ATC T, primer3: CAA GCA GAA GAC GGC ATA CGA GAT NNNNNN GTG ACT GGA GTT. After two more XP beads purifications (1:1) libraries were quantified using Qubit dsDNA HS Assay Kit (Invitrogen). For Illumina flowcell production, samples were equimolarly pooled and distributed on all lanes used for 75bp single read sequencing on Illumina HiSeq 2500.

After sequencing, FastQC (http://www.bioinformatics.babraham.ac.uk/) was used to perform a basic quality control of the resulting sequencing data. Fragments were aligned to the human reference genome hg38 with support of the Ensembl 104 splice sites using the aligner gsnap (v2020-12-16) (Wu and Nacu, 2010). Counts per gene and sample were obtained based on the overlap of the uniquely mapped reads with the same Ensembl annotation using featureCounts (v2.0.1) (Liao et al., 2014). Normalization of raw fragments based on library size and testing for differential expression between the different cell types/treatments was done with the DESeq2 R package (v1.30.1) (Love et al., 2014). Sample to sample Euclidean distance, Pearson’ and Spearman correlation coefficient (r) and PCA based upon the top 500 genes showing highest variance were computed to explore correlation between biological replicates and different libraries. To identify differential expressed genes, counts were fitted to the negative binomial distribution and genes were tested between conditions using the Wald test of DESeq2. Resulting p-values were corrected for multiple testing with the using Independent Hypothesis Weighting (v1.18.0) (Ignatiadis et al., 2016). Genes with a maximum of 5% false discovery rate (padj ≤ 0.05), 0.5>fold regulation>2.0 and counts above 200 were considered as significantly differentially expressed.

To directly compare the transcriptome profile of our expanded PP cells with previously published data sets, raw sequencing data of the GEO archives GSE156712 (Ma et al., 2022) were downloaded. The array express, E-MTAB-9992 archive from EBI (Nakamura et al., 2022) and EGAS00001003127 from EGA (Ramond et al., 2018) offered bam files. Here, the fastq files were extracted with picard tools (v2.25.6). All these data sets underwent the same processing procedure. Fragments/Reads were aligned to the human reference genome hg38 with support of the Ensembl 104 splice sites using the aligner gsnap (v2020-12-16). Counts per gene and sample were obtained based on the overlap of the uniquely mapped reads with the same Ensembl annotation using featureCounts (v2.0.1) (Liao et al., 2014). The various strand-specificity of the several projects was taken into account for the gene counting. Normalization of raw fragments based on library size and scaling the count on a log2 scale was done with the DESeq2 R package (v1.30.1) (Love et al., 2014) and the Variance Stabilizing Transformation (vst) function. These values were used for plotting.

Original RNA Seq data have been deposited in GEO under the GSE216266 accession number and accessible with the token sfwbsqyslfirbcj (C6 ePP transcriptome) and under the GSE216179 accession number and accessible with the token enyjwwycfjsjdub (CINI ePP transcriptome). GO and KEGG analyses have been carried out using the Enrichr suite https://maayanlab.cloud/Enrichr/ (Kuleshov et al., 2016) and GSEA analyses using the UCSD Broad Institute suite (https://www.gsea-msigdb.org/gsea/index.jsp) (Kuleshov et al., 2016). Heat maps were generated using the Morpheus application (https://software.broadinstitute.org/morpheus/)

## REFERENCES

1. Afelik, S., Pool, B., Schmerr, M., Penton, C., and Jensen, J. (2015). Wnt7b is required for epithelial progenitor growth and operates during epithelial-to-mesenchymal signaling in pancreatic development. Developmental biology 399, 204–217.

2. Afelik, S., Qu, X., Hasrouni, E., Bukys, M.A., Deering, T., Nieuwoudt, S., Rogers, W., Macdonald, R.J., and Jensen, J. (2012). Notch-mediated patterning and cell fate allocation of pancreatic progenitor cells. Development 139, 1744–1753.

3. Aghazadeh, Y., Sarangi, F., Poon, F., Nkennor, B., McGaugh, E.C., Nunes, S.S., and Nostro, M.C. (2022). GP2-enriched pancreatic progenitors give rise to functional beta cells in vivo and eliminate the risk of teratoma formation. Stem Cell Reports 17, 964–978.

4. Ameri, J., Borup, R., Prawiro, C., Ramond, C., Schachter, K.A., Scharfmann, R., and Semb, H. (2017). Efficient Generation of Glucose-Responsive Beta Cells from Isolated GP2(+) Human Pancreatic Progenitors. Cell reports 19, 36–49.

5. Amin, S., Cook, B., Zhou, T., Ghazizadeh, Z., Lis, R., Zhang, T., Khalaj, M., Crespo, M., Perera, M., Xiang, J.Z., et al. (2018). Discovery of a drug candidate for GLIS3-associated diabetes. Nature communications 9, 2681.

6. Apelqvist, A., Li, H., Sommer, L., Beatus, P., Anderson, D.J., Honjo, T., Hrabe de Angelis, M., Lendahl, U., and Edlund, H. (1999). Notch signalling controls pancreatic cell differentiation. Nature 400, 877–881.

7. Arda, H.E., Benitez, C.M., and Kim, S.K. (2013). Gene regulatory networks governing pancreas development. Developmental cell 25, 5–13.

8. Azzarelli, R., Hurley, C., Sznurkowska, M.K., Rulands, S., Hardwick, L., Gamper, I., Ali, F., McCracken, L., Hindley, C., McDuff, F., et al. (2017). Multi-site Neurogenin3 Phosphorylation Controls Pancreatic Endocrine Differentiation. Developmental cell 41, 274–286 e275.

9. Balboa, D., Barsby, T., Lithovius, V., Saarimaki-Vire, J., Omar-Hmeadi, M., Dyachok, O., Montaser, H., Lund, P.E., Yang, M., Ibrahim, H., et al. (2022). Functional, metabolic and transcriptional maturation of human pancreatic islets derived from stem cells. Nature biotechnology.

10. Bastidas-Ponce, A., Scheibner, K., Lickert, H., and Bakhti, M. (2017). Cellular and molecular mechanisms coordinating pancreas development. Development 144, 2873–2888.

11. Binot, A.C., Manfroid, I., Flasse, L., Winandy, M., Motte, P., Martial, J.A., Peers, B., and Voz, M.L. (2010). Nkx6.1 and nkx6.2 regulate alpha- and beta-cell formation in zebrafish by acting on pancreatic endocrine progenitor cells. Developmental biology 340, 397–407.

12. Brown, M.L., and Schneyer, A.L. (2010). Emerging roles for the TGFbeta family in pancreatic beta-cell homeostasis. Trends Endocrinol Metab 21, 441–448.

13. Cebola, I., Rodriguez-Segui, S.A., Cho, C.H., Bessa, J., Rovira, M., Luengo, M., Chhatriwala, M., Berry, A., Ponsa-Cobas, J., Maestro, M.A., et al. (2015). TEAD and YAP regulate the enhancer network of human embryonic pancreatic progenitors. Nature cell biology 17, 615–626.

14. Cerda-Esteban, N., Naumann, H., Ruzittu, S., Mah, N., Pongrac, I.M., Cozzitorto, C., Hommel, A., Andrade-Navarro, M.A., Bonifacio, E., and Spagnoli, F.M. (2017). Stepwise reprogramming of liver cells to a pancreas progenitor state by the transcriptional regulator Tgif2. Nature communications 8, 14127.

15. Chen, B., Dodge, M.E., Tang, W., Lu, J., Ma, Z., Fan, C.W., Wei, S., Hao, W., Kilgore, J., Williams, N.S., et al. (2009). Small molecule-mediated disruption of Wnt-dependent signaling in tissue regeneration and cancer. Nat Chem Biol 5, 100–107.

16. Chen, H., Gu, X., Liu, Y., Wang, J., Wirt, S.E., Bottino, R., Schorle, H., Sage, J., and Kim, S.K. (2011). PDGF signalling controls age-dependent proliferation in pancreatic beta-cells. Nature 478, 349–355.

17. Cogger, K.F., Sinha, A., Sarangi, F., McGaugh, E.C., Saunders, D., Dorrell, C., Mejia-Guerrero, S., Aghazadeh, Y., Rourke, J.L., Screaton, R.A., et al. (2017). Glycoprotein 2 is a specific cell surface marker of human pancreatic progenitors. Nature communications 8, 331.

18. Crisera, C.A., Rose, M.I., Connelly, P.R., Li, M., Colen, K.L., Longaker, M.T., and Gittes, G.K. (1999). The ontogeny of TGF-beta1,-beta2,-beta3, and TGF-beta receptor-II expression in the pancreas: implications for regulation of growth and differentiation. J Pediatr Surg 34, 689–693; discussion 693-684.

19. Deutsch, G., Jung, J., Zheng, M., Lora, J., and Zaret, K.S. (2001). A bipotential precursor population for pancreas and liver within the embryonic endoderm. Development 128, 871–881.

20. Dichmann, D.S., Miller, C.P., Jensen, J., Scott Heller, R., and Serup, P. (2003). Expression and misexpression of members of the FGF and TGFbeta families of growth factors in the developing mouse pancreas. Developmental dynamics: an official publication of the American Association of Anatomists 226, 663–674.

21. Du, Y., Liang, Z., Wang, S., Sun, D., Wang, X., Liew, S.Y., Lu, S., Wu, S., Jiang, Y., Wang, Y., et al. (2022). Human pluripotent stem-cell-derived islets ameliorate diabetes in non-human primates. Nature medicine 28, 272–282.

22. Duvall, E., Benitez, C.M., Tellez, K., Enge, M., Pauerstein, P.T., Li, L., Baek, S., Quake, S.R., Smith, J.P., Sheffield, N.C., et al. (2022). Single-cell transcriptome and accessible chromatin dynamics during endocrine pancreas development. Proceedings of the National Academy of Sciences of the United States of America 119, e2201267119.

23. Ertosun, M.G., Hapil, F.Z., and Osman Nidai, O. (2016). E2F1 transcription factor and its impact on growth factor and cytokine signaling. Cytokine Growth Factor Rev 31, 17–25.

24. Gao, N., Le Lay, J., Qin, W., Doliba, N., Schug, J., Fox, A.J., Smirnova, O., Matschinsky, F.M., and Kaestner, K.H. (2010). Foxa1 and Foxa2 maintain the metabolic and secretory features of the mature beta-cell. Mol Endocrinol 24, 1594–1604.

25. Gao, N., LeLay, J., Vatamaniuk, M.Z., Rieck, S., Friedman, J.R., and Kaestner, K.H. (2008). Dynamic regulation of Pdx1 enhancers by Foxa1 and Foxa2 is essential for pancreas development. Genes & development 22, 3435–3448.

26. Gellibert, F., Woolven, J., Fouchet, M.H., Mathews, N., Goodland, H., Lovegrove, V., Laroze, A., Nguyen, V.L., Sautet, S., Wang, R., et al. (2004). Identification of 1,5-naphthyridine derivatives as a novel series of potent and selective TGF-beta type I receptor inhibitors. J Med Chem 47, 4494–4506.

27. Gradwohl, G., Dierich, A., LeMeur, M., and Guillemot, F. (2000). neurogenin3 is required for the development of the four endocrine cell lineages of the pancreas. Proceedings of the National Academy of Sciences of the United States of America 97, 1607–1611.

28. Grapin-Botton, A., Majithia, A.R., and Melton, D.A. (2001). Key events of pancreas formation are triggered in gut endoderm by ectopic expression of pancreatic regulatory genes. Genes & development 15, 444–454.

29. Guo, T., Landsman, L., Li, N., and Hebrok, M. (2013). Factors expressed by murine embryonic pancreatic mesenchyme enhance generation of insulin-producing cells from hESCs. Diabetes 62, 1581–1592.

30. Hart, A., Papadopoulou, S., and Edlund, H. (2003). Fgf10 maintains notch activation, stimulates proliferation, and blocks differentiation of pancreatic epithelial cells. Developmental dynamics: an official publication of the American Association of Anatomists 228, 185–193.

31. Henseleit, K.D., Nelson, S.B., Kuhlbrodt, K., Hennings, J.C., Ericson, J., and Sander, M. (2005). NKX6 transcription factor activity is required for alpha- and beta-cell development in the pancreas. Development 132, 3139–3149.

32. Hogrebe, N.J., Augsornworawat, P., Maxwell, K.G., Velazco-Cruz, L., and Millman, J.R. (2020). Targeting the cytoskeleton to direct pancreatic differentiation of human pluripotent stem cells. Nature biotechnology 38, 460–470.

33. Hogrebe, N.J., Maxwell, K.G., Augsornworawat, P., and Millman, J.R. (2021). Generation of insulin-producing pancreatic beta cells from multiple human stem cell lines. Nature protocols 16, 4109–4143.

34. Inman, G.J., Nicolas, F.J., Callahan, J.F., Harling, J.D., Gaster, L.M., Reith, A.D., Laping, N.J., and Hill, C.S. (2002). SB-431542 is a potent and specific inhibitor of transforming growth factor-beta superfamily type I activin receptor-like kinase (ALK) receptors ALK4, ALK5, and ALK7. Mol Pharmacol 62, 65–74.

35. Jensen, J., Pedersen, E.E., Galante, P., Hald, J., Heller, R.S., Ishibashi, M., Kageyama, R., Guillemot, F., Serup, P., and Madsen, O.D. (2000). Control of endodermal endocrine development by Hes-1. Nature genetics 24, 36–44.

36. Jonsson, J., Carlsson, L., Edlund, T., and Edlund, H. (1994). Insulin-promoter-factor 1 is required for pancreas development in mice. Nature 371, 606–609.

37. Jung, J., Zheng, M., Goldfarb, M., and Zaret, K.S. (1999). Initiation of mammalian liver development from endoderm by fibroblast growth factors. Science 284, 1998–2003.

38. Kobayashi, H., Spilde, T.L., Bhatia, A.M., Buckingham, R.B., Hembree, M.J., Prasadan, K., Preuett, B.L., Imamura, M., and Gittes, G.K. (2002). Retinoid signaling controls mouse pancreatic exocrine lineage selection through epithelial-mesenchymal interactions. Gastroenterology 123, 1331–1340.

39. Konagaya, S., and Iwata, H. (2019). Chemically defined conditions for long-term maintenance of pancreatic progenitors derived from human induced pluripotent stem cells. Scientific reports 9, 640.

40. Krentz, N.A.J., van Hoof, D., Li, Z., Watanabe, A., Tang, M., Nian, C., German, M.S., and Lynn, F.C. (2017). Phosphorylation of NEUROG3 Links Endocrine Differentiation to the Cell Cycle in Pancreatic Progenitors. Developmental cell 41, 129–142 e126.

41. Kroon, E., Martinson, L.A., Kadoya, K., Bang, A.G., Kelly, O.G., Eliazer, S., Young, H., Richardson, M., Smart, N.G., Cunningham, J., et al. (2008). Pancreatic endoderm derived from human embryonic stem cells generates glucose-responsive insulin-secreting cells in vivo. Nature biotechnology 26, 443–452.

42. Lammert, E., Cleaver, O., and Melton, D. (2001). Induction of pancreatic differentiation by signals from blood vessels. Science 294, 564–567.

43. Lee, C.S., Sund, N.J., Behr, R., Herrera, P.L., and Kaestner, K.H. (2005). Foxa2 is required for the differentiation of pancreatic alpha-cells. Developmental biology 278, 484–495.

44. Lee, K., Cho, H., Rickert, R.W., Li, Q.V., Pulecio, J., Leslie, C.S., and Huangfu, D. (2019). FOXA2 Is Required for Enhancer Priming during Pancreatic Differentiation. Cell reports 28, 382–393 e387.

45. Li, X., Yang, K.Y., Chan, V.W., Leung, K.T., Zhang, X.B., Wong, A.S., Chong, C.C.N., Wang, C.C., Ku, M., and Lui, K.O. (2020). Single-Cell RNA-Seq Reveals that CD9 Is a Negative Marker of Glucose-Responsive Pancreatic beta-like Cells Derived from Human Pluripotent Stem Cells. Stem Cell Reports 15, 1111–1126.

46. Lorberbaum, D.S., Kishore, S., Rosselot, C., Sarbaugh, D., Brooks, E.P., Aragon, E., Xuan, S., Simon, O., Ghosh, D., Mendelsohn, C., et al. (2020). Retinoic acid signaling within pancreatic endocrine progenitors regulates mouse and human beta cell specification. Development 147.

47. Love, M.I., Huber, W., and Anders, S. (2014). Moderated estimation of fold change and dispersion for RNA-seq data with DESeq2. Genome biology 15, 550.

48. Ma, X., Lu, Y., Zhou, Z., Li, Q., Chen, X., Wang, W., Jin, Y., Hu, Z., Chen, G., Deng, Q., et al. (2022). Human expandable pancreatic progenitor-derived beta cells ameliorate diabetes. Sci Adv 8, eabk1826.

49. Mahaddalkar, P.U., Scheibner, K., Pfluger, S., Ansarullah, Sterr, M., Beckenbauer, J., Irmler, M., Beckers, J., Knobel, S., and Lickert, H. (2020). Generation of pancreatic beta cells from CD177(+) anterior definitive endoderm. Nature biotechnology 38, 1061–1072.

50. Mamidi, A., Prawiro, C., Seymour, P.A., de Lichtenberg, K.H., Jackson, A., Serup, P., and Semb, H. (2018). Mechanosignalling via integrins directs fate decisions of pancreatic progenitors. Nature 564, 114–118.

51. Martin, M., Gallego-Llamas, J., Ribes, V., Kedinger, M., Niederreither, K., Chambon, P., Dolle, P., and Gradwohl, G. (2005). Dorsal pancreas agenesis in retinoic acid-deficient Raldh2 mutant mice. Developmental biology 284, 399–411.

52. Masui, T., Swift, G.H., Deering, T., Shen, C., Coats, W.S., Long, Q., Elsasser, H.P., Magnuson, M.A., and MacDonald, R.J. (2010). Replacement of Rbpj with Rbpjl in the PTF1 complex controls the final maturation of pancreatic acinar cells. Gastroenterology 139, 270–280.

53. McLin, V.A., Rankin, S.A., and Zorn, A.M. (2007). Repression of Wnt/beta-catenin signaling in the anterior endoderm is essential for liver and pancreas development. Development 134, 2207–2217.

54. Meng, Z., Moroishi, T., and Guan, K.L. (2016). Mechanisms of Hippo pathway regulation. Genes & development 30, 1–17.

55. Miguel-Escalada, I., Maestro, M.A., Balboa, D., Elek, A., Bernal, A., Bernardo, E., Grau, V., Garcia-Hurtado, J., Sebe-Pedros, A., and Ferrer, J. (2022). Pancreas agenesis mutations disrupt a lead enhancer controlling a developmental enhancer cluster. Developmental cell 57, 1922–1936 e1929.

56. Millman, J.R., Xie, C., Van Dervort, A., Gurtler, M., Pagliuca, F.W., and Melton, D.A. (2016). Generation of stem cell-derived beta-cells from patients with type 1 diabetes. Nature communications 7, 11463.

57. Miralles, F., Czernichow, P., and Scharfmann, R. (1998). Follistatin regulates the relative proportions of endocrine versus exocrine tissue during pancreatic development. Development 125, 1017–1024.

58. Molotkov, A., Molotkova, N., and Duester, G. (2005). Retinoic acid generated by Raldh2 in mesoderm is required for mouse dorsal endodermal pancreas development. Developmental dynamics: an official publication of the American Association of Anatomists 232, 950–957.

59. Murtaugh, L.C., Stanger, B.Z., Kwan, K.M., and Melton, D.A. (2003). Notch signaling controls multiple steps of pancreatic differentiation. Proceedings of the National Academy of Sciences of the United States of America 100, 14920–14925.

60. Nair, G.G., Liu, J.S., Russ, H.A., Tran, S., Saxton, M.S., Chen, R., Juang, C., Li, M.L., Nguyen, V.Q., Giacometti, S., et al. (2019). Recapitulating endocrine cell clustering in culture promotes maturation of human stem-cell-derived beta cells. Nature cell biology 21, 263–274.

61. Nakamura, A., Wong, Y.F., Venturato, A., Michaut, M., Venkateswaran, S., Santra, M., Goncalves, C., Larsen, M., Leuschner, M., Kim, Y.H., et al. (2022). Long-term feeder-free culture of human pancreatic progenitors on fibronectin or matrix-free polymer potentiates beta cell differentiation. Stem Cell Reports 17, 1215–1228.

62. Nelson, S.B., Schaffer, A.E., and Sander, M. (2007). The transcription factors Nkx6.1 and Nkx6.2 possess equivalent activities in promoting beta-cell fate specification in Pdx1+ pancreatic progenitor cells. Development 134, 2491–2500.

63. Nir, T., Melton, D.A., and Dor, Y. (2007). Recovery from diabetes in mice by beta cell regeneration. The Journal of clinical investigation 117, 2553–2561.

64. Norgaard, G.A., Jensen, J.N., and Jensen, J. (2003). FGF10 signaling maintains the pancreatic progenitor cell state revealing a novel role of Notch in organ development. Developmental biology 264, 323–338.

65. Nostro, M.C., Sarangi, F., Yang, C., Holland, A., Elefanty, A.G., Stanley, E.G., Greiner, D.L., and Keller, G. (2015). Efficient generation of NKX6-1+ pancreatic progenitors from multiple human pluripotent stem cell lines. Stem Cell Reports 4, 591–604.

66. Offield, M.F., Jetton, T.L., Labosky, P.A., Ray, M., Stein, R.W., Magnuson, M.A., Hogan, B.L., and Wright, C.V. (1996). PDX-1 is required for pancreatic outgrowth and differentiation of the rostral duodenum. Development 122, 983–995.

67. Ostrom, M., Loffler, K.A., Edfalk, S., Selander, L., Dahl, U., Ricordi, C., Jeon, J., Correa-Medina, M., Diez, J., and Edlund, H. (2008). Retinoic acid promotes the generation of pancreatic endocrine progenitor cells and their further differentiation into beta-cells. PloS one 3, e2841.

68. Pagliuca, F.W., Millman, J.R., Gurtler, M., Segel, M., Van Dervort, A., Ryu, J.H., Peterson, Q.P., Greiner, D., and Melton, D.A. (2014). Generation of functional human pancreatic beta cells in vitro. Cell 159, 428–439.

69. Pedersen, J.K., Nelson, S.B., Jorgensen, M.C., Henseleit, K.D., Fujitani, Y., Wright, C.V., Sander, M., Serup, P., and Beta Cell Biology, C. (2005). Endodermal expression of Nkx6 genes depends differentially on Pdx1. Developmental biology 288, 487–501.

70. Qu, X., Afelik, S., Jensen, J.N., Bukys, M.A., Kobberup, S., Schmerr, M., Xiao, F., Nyeng, P., Veronica Albertoni, M., Grapin-Botton, A., et al. (2013). Notch-mediated post-translational control of Ngn3 protein stability regulates pancreatic patterning and cell fate commitment. Developmental biology 376, 1–12.

71. Ramond, C., Beydag-Tasoz, B.S., Azad, A., van de Bunt, M., Petersen, M.B.K., Beer, N.L., Glaser, N., Berthault, C., Gloyn, A.L., Hansson, M., et al. (2018). Understanding human fetal pancreas development using subpopulation sorting, RNA sequencing and single-cell profiling. Development 145.

72. Ramond, C., Glaser, N., Berthault, C., Ameri, J., Kirkegaard, J.S., Hansson, M., Honore, C., Semb, H., and Scharfmann, R. (2017). Reconstructing human pancreatic differentiation by mapping specific cell populations during development. Elife 6.

73. Ramzy, A., Thompson, D.M., Ward-Hartstonge, K.A., Ivison, S., Cook, L., Garcia, R.V., Loyal, J., Kim, P.T.W., Warnock, G.L., Levings, M.K., et al. (2021). Implanted pluripotent stem-cell-derived pancreatic endoderm cells secrete glucose-responsive C-peptide in patients with type 1 diabetes. Cell stem cell 28, 2047–2061 e2045.

74. Rezania, A., Bruin, J.E., Arora, P., Rubin, A., Batushansky, I., Asadi, A., O’Dwyer, S., Quiskamp, N., Mojibian, M., Albrecht, T., et al. (2014). Reversal of diabetes with insulin-producing cells derived in vitro from human pluripotent stem cells. Nature biotechnology 32, 1121–1133.

75. Roberts, W.G., Whalen, P.M., Soderstrom, E., Moraski, G., Lyssikatos, J.P., Wang, H.F., Cooper, B., Baker, D.A., Savage, D., Dalvie, D., et al. (2005). Antiangiogenic and antitumor activity of a selective PDGFR tyrosine kinase inhibitor, CP-673,451. Cancer research 65, 957–966.

76. Rosado-Olivieri, E.A., Anderson, K., Kenty, J.H., and Melton, D.A. (2019). YAP inhibition enhances the differentiation of functional stem cell-derived insulin-producing beta cells. Nature communications 10, 1464.

77. Rossi, J.M., Dunn, N.R., Hogan, B.L., and Zaret, K.S. (2001). Distinct mesodermal signals, including BMPs from the septum transversum mesenchyme, are required in combination for hepatogenesis from the endoderm. Genes & development 15, 1998–2009.

78. Rubin, S.M., Sage, J., and Skotheim, J.M. (2020). Integrating Old and New Paradigms of G1/S Control. Molecular cell 80, 183–192.

79. Russ, H.A., Parent, A.V., Ringler, J.J., Hennings, T.G., Nair, G.G., Shveygert, M., Guo, T., Puri, S., Haataja, L., Cirulli, V., et al. (2015). Controlled induction of human pancreatic progenitors produces functional beta-like cells in vitro. The EMBO journal 34, 1759–1772.

80. Sander, M., Sussel, L., Conners, J., Scheel, D., Kalamaras, J., Dela Cruz, F., Schwitzgebel, V., Hayes-Jordan, A., and German, M. (2000). Homeobox gene Nkx6.1 lies downstream of Nkx2.2 in the major pathway of beta-cell formation in the pancreas. Development 127, 5533–5540.

81. Sanvitale, C.E., Kerr, G., Chaikuad, A., Ramel, M.C., Mohedas, A.H., Reichert, S., Wang, Y., Triffitt, J.T., Cuny, G.D., Yu, P.B., et al. (2013). A new class of small molecule inhibitor of BMP signaling. PloS one 8, e62721.

82. Sanvito, F., Herrera, P.L., Huarte, J., Nichols, A., Montesano, R., Orci, L., and Vassalli, J.D. (1994). TGF-beta 1 influences the relative development of the exocrine and endocrine pancreas in vitro. Development 120, 3451–3462.

83. Scavuzzo, M.A., Hill, M.C., Chmielowiec, J., Yang, D., Teaw, J., Sheng, K., Kong, Y., Bettini, M., Zong, C., Martin, J.F., et al. (2018). Endocrine lineage biases arise in temporally distinct endocrine progenitors during pancreatic morphogenesis. Nature communications 9, 3356.

84. Schaffer, A.E., Freude, K.K., Nelson, S.B., and Sander, M. (2010). Nkx6 transcription factors and Ptf1a function as antagonistic lineage determinants in multipotent pancreatic progenitors. Developmental cell 18, 1022–1029.

85. Schaffer, A.E., Taylor, B.L., Benthuysen, J.R., Liu, J., Thorel, F., Yuan, W., Jiao, Y., Kaestner, K.H., Herrera, P.L., Magnuson, M.A., et al. (2013). Nkx6.1 controls a gene regulatory network required for establishing and maintaining pancreatic Beta cell identity. PLoS Genet 9, e1003274.

86. Seiffert, D., Bradley, J.D., Rominger, C.M., Rominger, D.H., Yang, F., Meredith, J.E., Jr., Wang, Q., Roach, A.H., Thompson, L.A., Spitz, S.M., et al. (2000). Presenilin-1 and −2 are molecular targets for gamma-secretase inhibitors. The Journal of biological chemistry 275, 34086–34091.

87. Serafimidis, I., Rodriguez-Aznar, E., Lesche, M., Yoshioka, K., Takuwa, Y., Dahl, A., Pan, D., and Gavalas, A. (2017). Pancreas lineage allocation and specification are regulated by sphingosine-1-phosphate signalling. PLoS Biol 15, e2000949.

88. Seymour, P.A., Collin, C.A., Egeskov-Madsen, A.R., Jorgensen, M.C., Shimojo, H., Imayoshi, I., de Lichtenberg, K.H., Kopan, R., Kageyama, R., and Serup, P. (2020). Jag1 Modulates an Oscillatory Dll1-Notch-Hes1 Signaling Module to Coordinate Growth and Fate of Pancreatic Progenitors. Developmental cell 52, 731–747 e738.

89. Seymour, P.A., Freude, K.K., Tran, M.N., Mayes, E.E., Jensen, J., Kist, R., Scherer, G., and Sander, M. (2007). SOX9 is required for maintenance of the pancreatic progenitor cell pool. Proceedings of the National Academy of Sciences of the United States of America 104, 1865–1870.

90. Seymour, P.A., Shih, H.P., Patel, N.A., Freude, K.K., Xie, R., Lim, C.J., and Sander, M. (2012). A Sox9/Fgf feed-forward loop maintains pancreatic organ identity. Development 139, 3363–3372.

91. Shapiro, A.M.J., Thompson, D., Donner, T.W., Bellin, M.D., Hsueh, W., Pettus, J., Wilensky, J., Daniels, M., Wang, R.M., Brandon, E.P., et al. (2021). Insulin expression and C-peptide in type 1 diabetes subjects implanted with stem cell-derived pancreatic endoderm cells in an encapsulation device. Cell Rep Med 2, 100466.

92. Sharon, N., Vanderhooft, J., Straubhaar, J., Mueller, J., Chawla, R., Zhou, Q., Engquist, E.N., Trapnell, C., Gifford, D.K., and Melton, D.A. (2019). Wnt Signaling Separates the Progenitor and Endocrine Compartments during Pancreas Development. Cell reports 27, 2281–2291 e2285.

93. Sherwood, R.I., Chen, T.Y., and Melton, D.A. (2009). Transcriptional dynamics of endodermal organ formation. Developmental dynamics: an official publication of the American Association of Anatomists 238, 29–42.

94. Shi, Z.D., Lee, K., Yang, D., Amin, S., Verma, N., Li, Q.V., Zhu, Z., Soh, C.L., Kumar, R., Evans, T., et al. (2017). Genome Editing in hPSCs Reveals GATA6 Haploinsufficiency and a Genetic Interaction with GATA4 in Human Pancreatic Development. Cell stem cell 20, 675–688 e676.

95. Shih, H.P., Kopp, J.L., Sandhu, M., Dubois, C.L., Seymour, P.A., Grapin-Botton, A., and Sander, M. (2012). A Notch-dependent molecular circuitry initiates pancreatic endocrine and ductal cell differentiation. Development 139, 2488–2499.

96. Shih, H.P., Seymour, P.A., Patel, N.A., Xie, R., Wang, A., Liu, P.P., Yeo, G.W., Magnuson, M.A., and Sander, M. (2015). A Gene Regulatory Network Cooperatively Controlled by Pdx1 and Sox9 Governs Lineage Allocation of Foregut Progenitor Cells. Cell reports 13, 326–336.

97. Spagnoli, F.M., and Brivanlou, A.H. (2008). The Gata5 target, TGIF2, defines the pancreatic region by modulating BMP signals within the endoderm. Development 135, 451–461.

98. Taylor, B.L., Benthuysen, J., and Sander, M. (2015). Postnatal beta-cell proliferation and mass expansion is dependent on the transcription factor Nkx6.1. Diabetes 64, 897–903.

99. Tojo, M., Hamashima, Y., Hanyu, A., Kajimoto, T., Saitoh, M., Miyazono, K., Node, M., and Imamura, T. (2005). The ALK-5 inhibitor A-83-01 inhibits Smad signaling and epithelial-to-mesenchymal transition by transforming growth factor-beta. Cancer Sci 96, 791–800.

100. Trott, J., Tan, E.K., Ong, S., Titmarsh, D.M., Denil, S., Giam, M., Wong, C.K., Wang, J., Shboul, M., Eio, M., et al. (2017). Long-Term Culture of Self-renewing Pancreatic Progenitors Derived from Human Pluripotent Stem Cells. Stem Cell Reports 8, 1675–1688.

101. Tulachan, S.S., Doi, R., Kawaguchi, Y., Tsuji, S., Nakajima, S., Masui, T., Koizumi, M., Toyoda, E., Mori, T., Ito, D., et al. (2003). All-trans retinoic acid induces differentiation of ducts and endocrine cells by mesenchymal/epithelial interactions in embryonic pancreas. Diabetes 52, 76–84.

102. Tulachan, S.S., Tei, E., Hembree, M., Crisera, C., Prasadan, K., Koizumi, M., Shah, S., Guo, P., Bottinger, E., and Gittes, G.K. (2007). TGF-beta isoform signaling regulates secondary transition and mesenchymal-induced endocrine development in the embryonic mouse pancreas. Developmental biology 305, 508–521.

103. Veres, A., Faust, A.L., Bushnell, H.L., Engquist, E.N., Kenty, J.H., Harb, G., Poh, Y.C., Sintov, E., Gurtler, M., Pagliuca, F.W., et al. (2019). Charting cellular identity during human in vitro beta-cell differentiation. Nature 569, 368–373.

104. Vinckier, N.K., Patel, N.A., Geusz, R.J., Wang, A., Wang, J., Matta, I., Harrington, A.R., Wortham, M., Wetton, N., Wang, J., et al. (2020). LSD1-mediated enhancer silencing attenuates retinoic acid signalling during pancreatic endocrine cell development. Nature communications 11, 2082.

105. Wang, S., Hecksher-Sorensen, J., Xu, Y., Zhao, A., Dor, Y., Rosenberg, L., Serup, P., and Gu, G. (2008). Myt1 and Ngn3 form a feed-forward expression loop to promote endocrine islet cell differentiation. Developmental biology 317, 531–540.

106. Zeng, H., Guo, M., Zhou, T., Tan, L., Chong, C.N., Zhang, T., Dong, X., Xiang, J.Z., Yu, A.S., Yue, L., et al. (2016). An Isogenic Human ESC Platform for Functional Evaluation of Genome-wide-Association-Study-Identified Diabetes Genes and Drug Discovery. Cell stem cell 19, 326–340.

107. Zhang, X., Ibrahimi, O.A., Olsen, S.K., Umemori, H., Mohammadi, M., and Ornitz, D.M. (2006). Receptor specificity of the fibroblast growth factor family. The complete mammalian FGF family. The Journal of biological chemistry 281, 15694–15700.

108. Zhou, Q., Law, A.C., Rajagopal, J., Anderson, W.J., Gray, P.A., and Melton, D.A. (2007). A multipotent progenitor domain guides pancreatic organogenesis. Developmental cell 13, 103–114.

109. Zhu, S., Russ, H.A., Wang, X., Zhang, M., Ma, T., Xu, T., Tang, S., Hebrok, M., and Ding, S. (2016a). Human pancreatic beta-like cells converted from fibroblasts. Nature communications 7, 10080.

110. Zhu, Z., Li, Q.V., Lee, K., Rosen, B.P., Gonzalez, F., Soh, C.L., and Huangfu, D. (2016b). Genome Editing of Lineage Determinants in Human Pluripotent Stem Cells Reveals Mechanisms of Pancreatic Development and Diabetes. Cell stem cell 18, 755–768.

## REFERENCES

111. Balboa, D., Barsby, T., Lithovius, V., Saarimaki-Vire, J., Omar-Hmeadi, M., Dyachok, O., Montaser, H., Lund, P.E., Yang, M., Ibrahim, H., et al. (2022). Functional, metabolic and transcriptional maturation of human pancreatic islets derived from stem cells. Nature biotechnology.

112. Cheung, A.Y., Horvath, L.M., Grafodatskaya, D., Pasceri, P., Weksberg, R., Hotta, A., Carrel, L., and Ellis, J. (2011). Isolation of MECP2-null Rett Syndrome patient hiPS cells and isogenic controls through X-chromosome inactivation. Hum Mol Genet 20, 2103–2115.

113. Ignatiadis, N., Klaus, B., Zaugg, J.B., and Huber, W. (2016). Data-driven hypothesis weighting increases detection power in genome-scale multiple testing. Nature methods 13, 577–580.

114. Kuleshov, M.V., Jones, M.R., Rouillard, A.D., Fernandez, N.F., Duan, Q., Wang, Z., Koplev, S., Jenkins, S.L., Jagodnik, K.M., Lachmann, A., et al. (2016). Enrichr: a comprehensive gene set enrichment analysis web server 2016 update. Nucleic acids research 44, W90–97.

115. Liao, Y., Smyth, G.K., and Shi, W. (2014). featureCounts: an efficient general purpose program for assigning sequence reads to genomic features. Bioinformatics 30, 923–930.

116. Love, M.I., Huber, W., and Anders, S. (2014). Moderated estimation of fold change and dispersion for RNA-seq data with DESeq2. Genome biology 15, 550.

117. Ma, X., Lu, Y., Zhou, Z., Li, Q., Chen, X., Wang, W., Jin, Y., Hu, Z., Chen, G., Deng, Q., et al. (2022). Human expandable pancreatic progenitor-derived beta cells ameliorate diabetes. Sci Adv 8, eabk1826.

118. Mahaddalkar, P.U., Scheibner, K., Pfluger, S., Ansarullah, Sterr, M., Beckenbauer, J., Irmler, M., Beckers, J., Knobel, S., and Lickert, H. (2020). Generation of pancreatic beta cells from CD177(+) anterior definitive endoderm. Nature biotechnology 38, 1061–1072.

119. Nakamura, A., Wong, Y.F., Venturato, A., Michaut, M., Venkateswaran, S., Santra, M., Goncalves, C., Larsen, M., Leuschner, M., Kim, Y.H., et al. (2022). Long-term feeder-free culture of human pancreatic progenitors on fibronectin or matrix-free polymer potentiates beta cell differentiation. Stem Cell Reports.

120. Ramond, C., Beydag-Tasoz, B.S., Azad, A., van de Bunt, M., Petersen, M.B.K., Beer, N.L., Glaser, N., Berthault, C., Gloyn, A.L., Hansson, M., et al. (2018). Understanding human fetal pancreas development using subpopulation sorting, RNA sequencing and single-cell profiling. Development 145.

121. Rezania, A., Bruin, J.E., Arora, P., Rubin, A., Batushansky, I., Asadi, A., O’Dwyer, S., Quiskamp, N., Mojibian, M., Albrecht, T., et al. (2014). Reversal of diabetes with insulin-producing cells derived in vitro from human pluripotent stem cells. Nature biotechnology 32, 1121–1133.

122. Shi, Z.D., Lee, K., Yang, D., Amin, S., Verma, N., Li, Q.V., Zhu, Z., Soh, C.L., Kumar, R., Evans, T., et al. (2017). Genome Editing in hPSCs Reveals GATA6 Haploinsufficiency and a Genetic Interaction with GATA4 in Human Pancreatic Development. Cell stem cell 20, 675–688 e676.

123. Wolf, C., Rapp, A., Berndt, N., Staroske, W., Schuster, M., Dobrick-Mattheuer, M., Kretschmer, S., Konig, N., Kurth, T., Wieczorek, D., et al. (2016). RPA and Rad51 constitute a cell intrinsic mechanism to protect the cytosol from self DNA. Nature communications 7, 11752.

124. Wu, T.D., and Nacu, S. (2010). Fast and SNP-tolerant detection of complex variants and splicing in short reads. Bioinformatics 26, 873–881.

125. Young, L., Sung, J., Stacey, G., and Masters, J.R. (2010). Detection of Mycoplasma in cell cultures. Nature protocols 5, 929–934.

