## Supplementary material for "Regulation of multiple signaling pathways promotes the consistent expansion of human pancreatic progenitors in defined conditions": All sup figures with legends

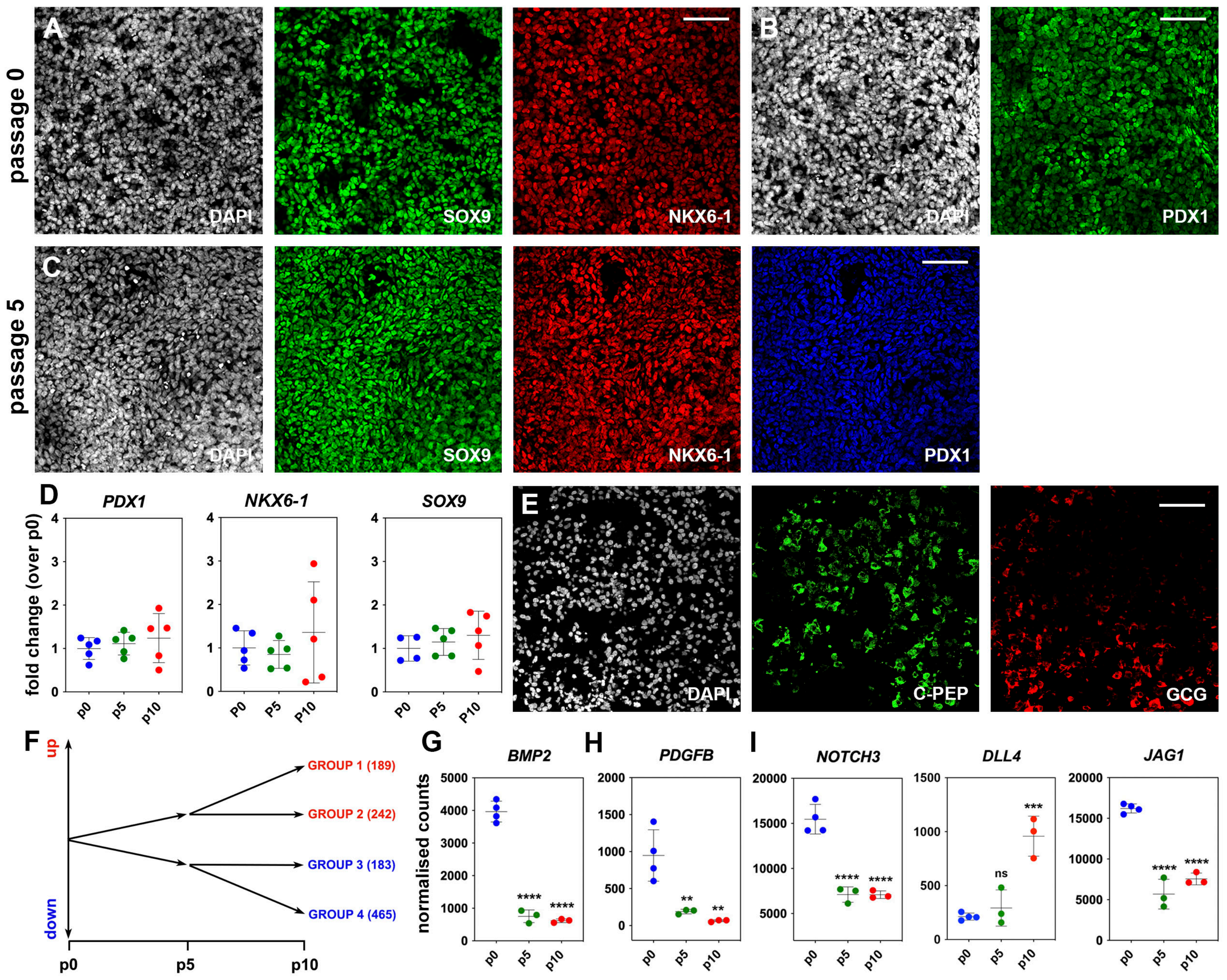

A
B
C
D
E
F
G
H
I
passage 0
passage 5
DAPI
SOX9
NKX6-1
PDX1
C-PEP
GCG
PDX1
NKX6-1
SOX9
fold change (over p0)
p0
p5
p10
P0
P5
P10
up
down
GROUP 1 (189)
GROUP 2 (242)
GROUP 3 (183)
GROUP 4 (465)
BMP2
PDGFB
NOTCH3
DLL4
JAG1
normalised counts
****
**
***
ns


Figure S1

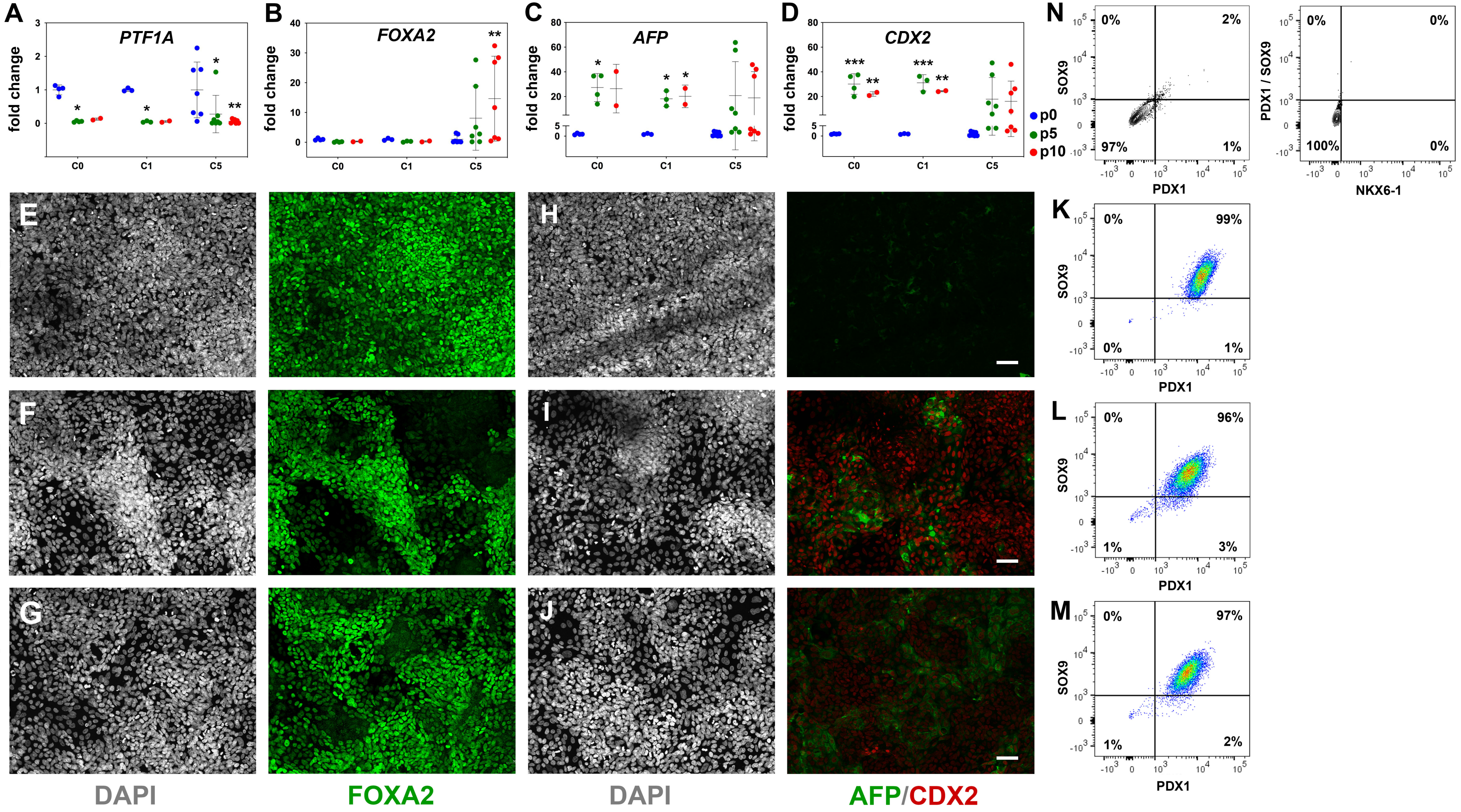

A
PTF1A
fold change
B
FOXA2
C
AFP
D
CDX2
C0
C1
C5
p0
p5
p10
N
SOX9
PDX1
PDX1 / SOX9
NKX6-1
K
L
M
E
F
G
H
I
J
passage 0
C5 passage 5
C5 passage 10
DAPI
FOXA2
DAPI
AFP/CDX2


Figure S2

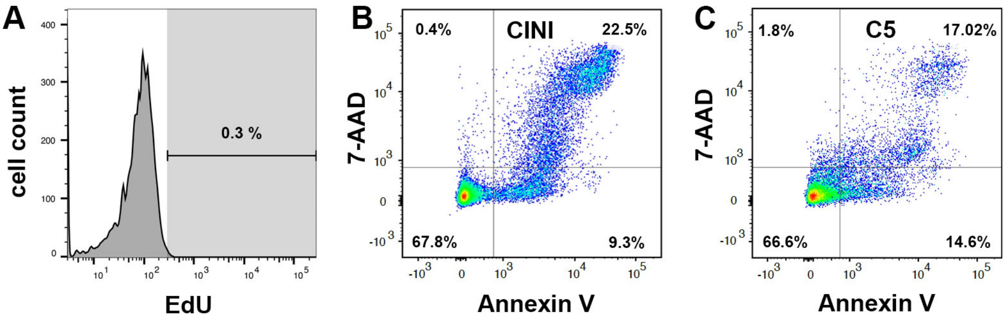
A
cell count
400
300
200
100
0
0.3 %
EdU
B
7-AAD
0.4%
CINI
22.5%
67.8%
9.3%
Annexin V
C
7-AAD
1.8%
C5
17.02%
66.6%
14.6%
Annexin V

Figure S3

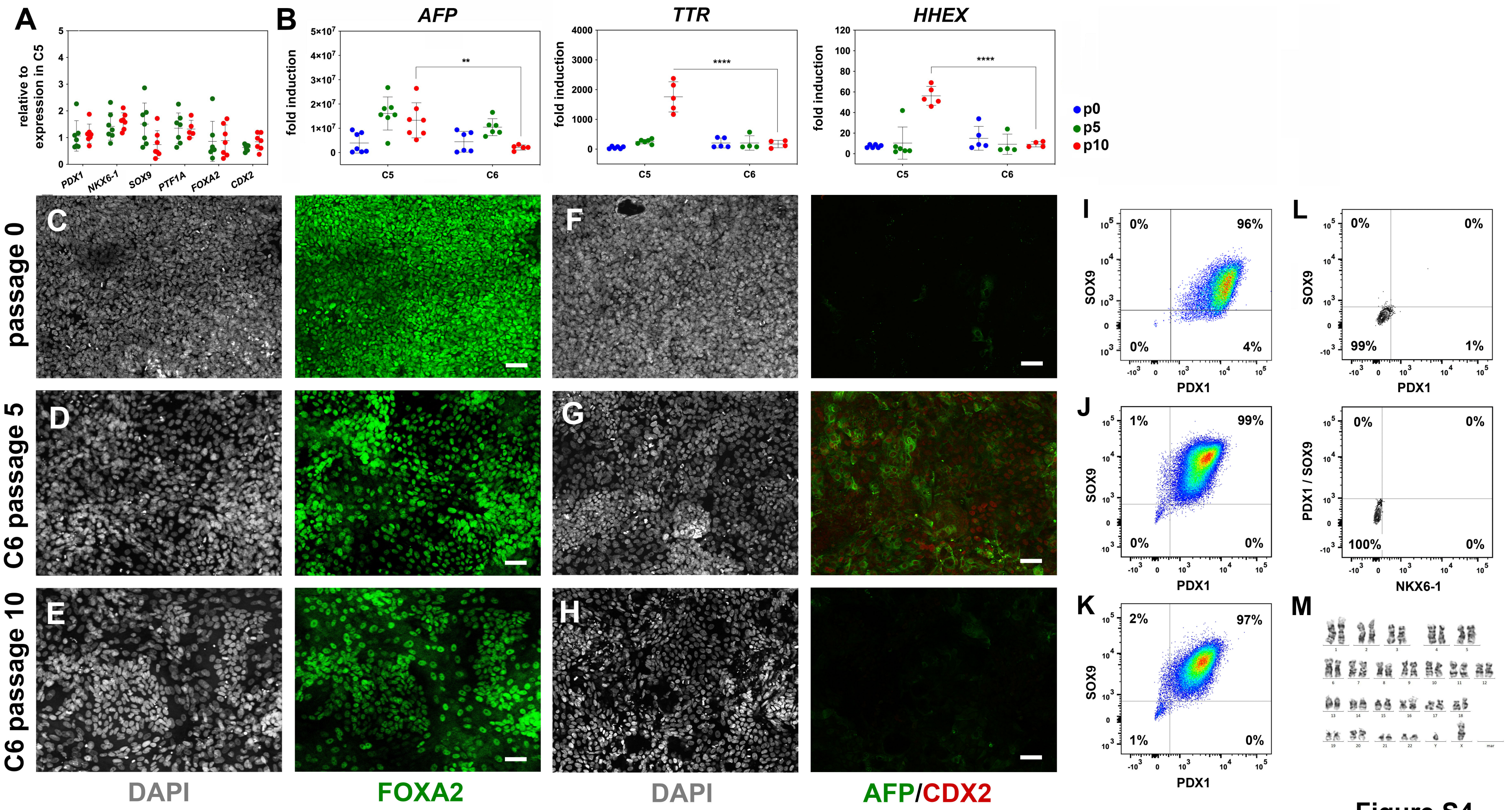
A
relative to expression in C5
PDX1
NKX6-1
SOX9
PTF1A
FOXA2
CDX2
B
AFP
TTR
HHEX
fold induction
C5
C6
p0
p5
p10
passage 0
C6 passage 5
C6 passage 10
DAPI
FOXA2
DAPI
AFP/CDX2
SOX9
PDX1
PDX1 / SOX9
NKX6-1

Figure S4

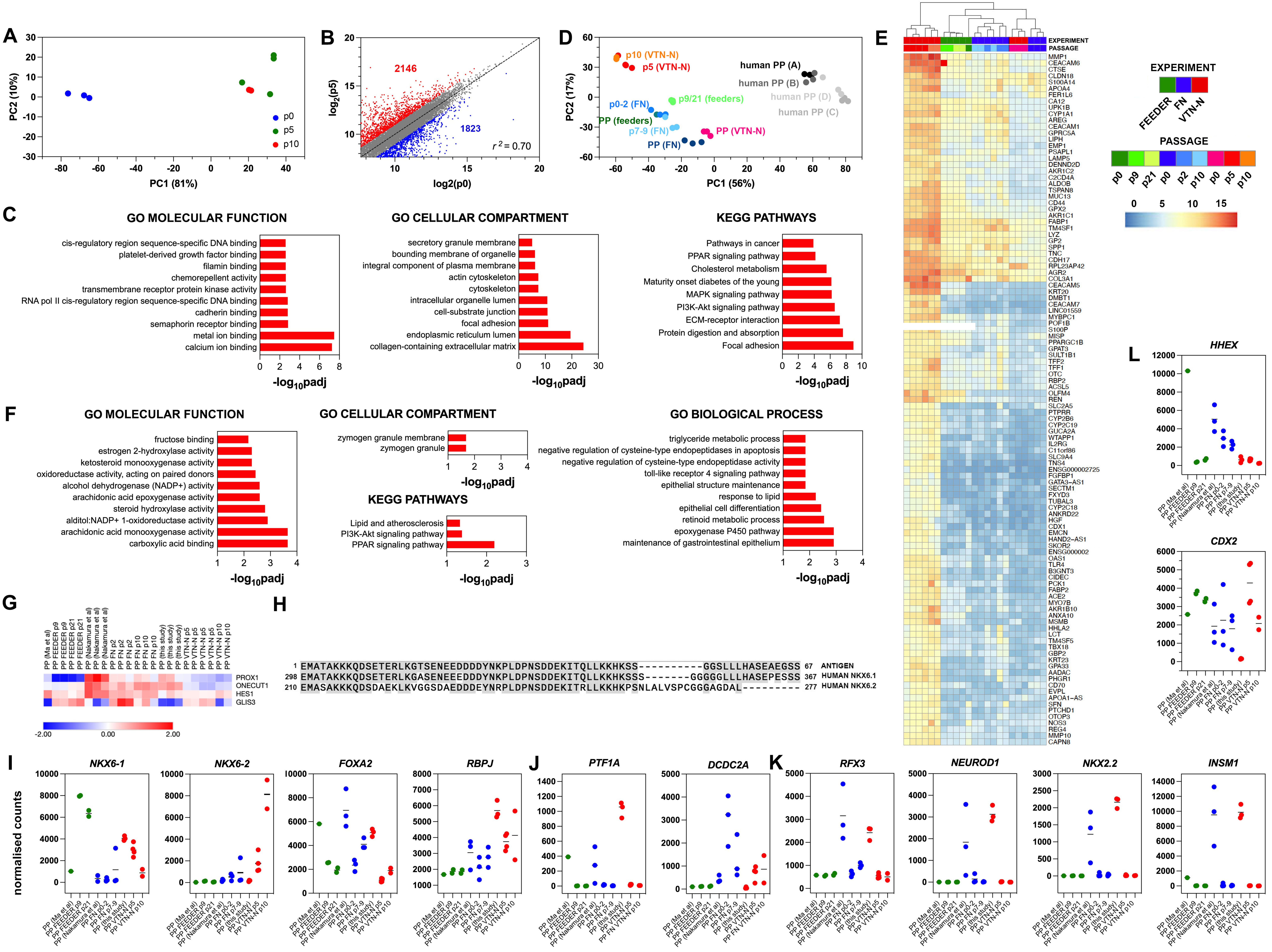
A
PC2 (10%)
PC1 (81%)
p0
p5
p10
B
log2(p5)
log2(p0)
2146
1823
$r^2$ = 0.70
D
p10 (VTN-N)
p5 (VTN-N)
human PP (A)
human PP (B)
human PP (D)
human PP (C)
p0-2 (FN)
p9/21 (feeders)
PP (feeders)
p7-9 (FN)
PP (VTN-N)
PP (FN)
PC2 (17%)
PC1 (56%)
E
EXPERIMENT
PASSAGE
FEEDER
FN
VTN-N
p0
p9
p21
p0
p2
p10
p0
p5
p10
0
5
10
15
MMP1
CEACAM6
CTSE
CLDN18
S100A14
APOA4
FER1L6
CA12
UPK1B
CYP1A1
AREG
CEACAM1
GPRC5A
LIPH
EMP1
PSAPL1
LAMP5
DENND2D
AKR1C2
C2CD4A
ALDOB
TSPAN8
MUC13
CD44
GPX2
AKR1C1
FABP1
TM4SF1
LYZ
GP2
SPP1
TNC
CDH17
RPL23AP42
AGR2
COL3A1
CEACAM5
KRT20
DMBT1
CEACAM7
LINC01559
MYBPC1
POF1B
S100P
MISP
PPARGC1B
GPAT3
SULT1B1
TFF2
TFF1
OTC
RBP2
ACSL5
OLFM4
REN
SLC2A5
PTPRR
CYP2B6
CYP2C19
GUCA2A
WTAPP1
IL2RG
C11orf86
SLC9A4
TNS4
ENSG000002725
FGFBP1
GATA3-AS1
SECTM1
FXYD3
TUBAL3
CYP2C18
ANKRD22
HGF
CDX1
EMCN
HAND2-AS1
SKOR2
ENSG000002
OAS1
TLR4
B3GNT3
CIDEC
PCK1
FABP2
ACE2
MYO7B
AKR1B10
ANXA10
MSMB
HHLA2
LCT
TM4SF5
TBX18
GBP2
KRT23
GPA33
AADAC
PHGR1
CD70
EVPL
APOA1-AS
SFN
PTCHD1
OTOP3
NOS3
REG4
MMP10
CAPN8
C
GO MOLECULAR FUNCTION
cis-regulatory region sequence-specific DNA binding
platelet-derived growth factor binding
filamin binding
chemorepellent activity
transmembrane receptor protein kinase activity
RNA pol II cis-regulatory region sequence-specific DNA binding
cadherin binding
semaphorin receptor binding
metal ion binding
calcium ion binding
-log10padj
GO CELLULAR COMPARTMENT
secretory granule membrane
bounding membrane of organelle
integral component of plasma membrane
actin cytoskeleton
cytoskeleton
intracellular organelle lumen
cell-substrate junction
focal adhesion
endoplasmic reticulum lumen
collagen-containing extracellular matrix
-log10padj
KEGG PATHWAYS
Pathways in cancer
PPAR signaling pathway
Cholesterol metabolism
Maturity onset diabetes of the young
MAPK signaling pathway
PI3K-Akt signaling pathway
ECM-receptor interaction
Protein digestion and absorption
Focal adhesion
-log10padj
F
GO MOLECULAR FUNCTION
fructose binding
estrogen 2-hydroxylase activity
ketosteroid monooxygenase activity
oxidoreductase activity, acting on paired donors
alcohol dehydrogenase (NADP+) activity
arachidonic acid epoxygenase activity
steroid hydroxylase activity
alditol:NADP+ 1-oxidoreductase activity
arachidonic acid monooxygenase activity
carboxylic acid binding
-log10padj
GO CELLULAR COMPARTMENT
zymogen granule membrane
zymogen granule
-log10padj
KEGG PATHWAYS
Lipid and atherosclerosis
PI3K-Akt signaling pathway
PPAR signaling pathway
-log10padj
GO BIOLOGICAL PROCESS
triglyceride metabolic process
negative regulation of cysteine-type endopeptidases in apoptosis
negative regulation of cysteine-type endopeptidase activity
toll-like receptor 4 signaling pathway
epithelial structure maintenance
response to lipid
epithelial cell differentiation
retinoid metabolic process
epoxygenase P450 pathway
maintenance of gastrointestinal epithelium
-log10padj
G
PP (Ma et al)
PP FEEDER p9
PP FEEDER p9
PP FEEDER p21
PP FEEDER p21
PP (Nakamura et al)
PP (Nakamura et al)
PP (Nakamura et al)
PP FN p2
PP FN p2
PP FN p2
PP FN p10
PP FN p10
PP FN p10
PP (this study)
PP (this study)
PP (this study)
PP VTN-N p5
PP VTN-N p5
PP VTN-N p5
PP VTN-N p10
PP VTN-N p10
PP VTN-N p10
PROX1
ONECUT1
HES1
GLIS3
-2.00
0.00
2.00
H
1 EMATAKKKQDSETERLKGTSENEEDDDYNKPLDPNSDDEKITQLLKKHKSS----------GGSLLLHASEAEGSS 67 ANTIGEN
298 EMATAKKKQDSETERLKGASENEEDDDYNKPLDPNSDDEKITQLLKKHKSSS-------GGGGGLLLHASEPESSS 367 HUMAN NKX6.1
210 EMASAKKKQDSDAEKLKVGGSDAEDDDEYNRPLDPNSDDEKITRLLKKHKPSNLALVSPCGGGAGDAL------- 277 HUMAN NKX6.2
I
normalised counts
NKX6-1
NKX6-2
FOXA2
RBPJ
J
PTF1A
DCDC2A
K
RFX3
NEUROD1
NKX2.2
INSM1
L
HHEX
CDX2
PP (Ma et al)
PP FEEDER p9
PP FEEDER p21
PP (Nakamura et al)
PP FN p0-2
PP FN p7-9
PP (this study)
PP VTN-N p5
PP VTN-N p10

Figure S5

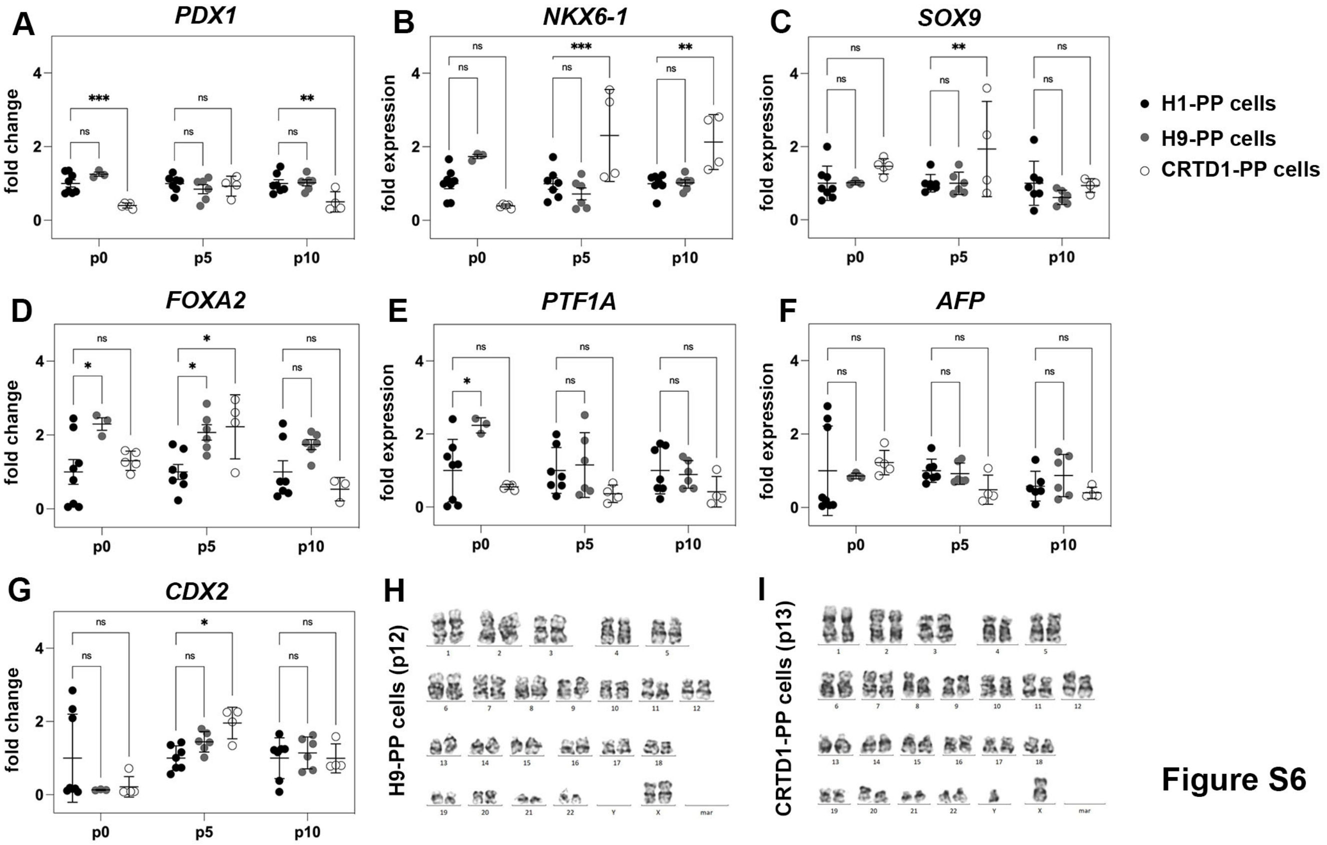

A
PDX1
fold change
B
NKX6-1
fold expression
C
SOX9
fold expression
D
FOXA2
fold change
E
PTF1A
fold expression
F
AFP
fold expression
G
CDX2
fold change
p0
p5
p10
ns
*
**
***
H1-PP cells
H9-PP cells
CRTD1-PP cells
H
H9-PP cells (p12)
I
CRTD1-PP cells (p13)


Figure S6

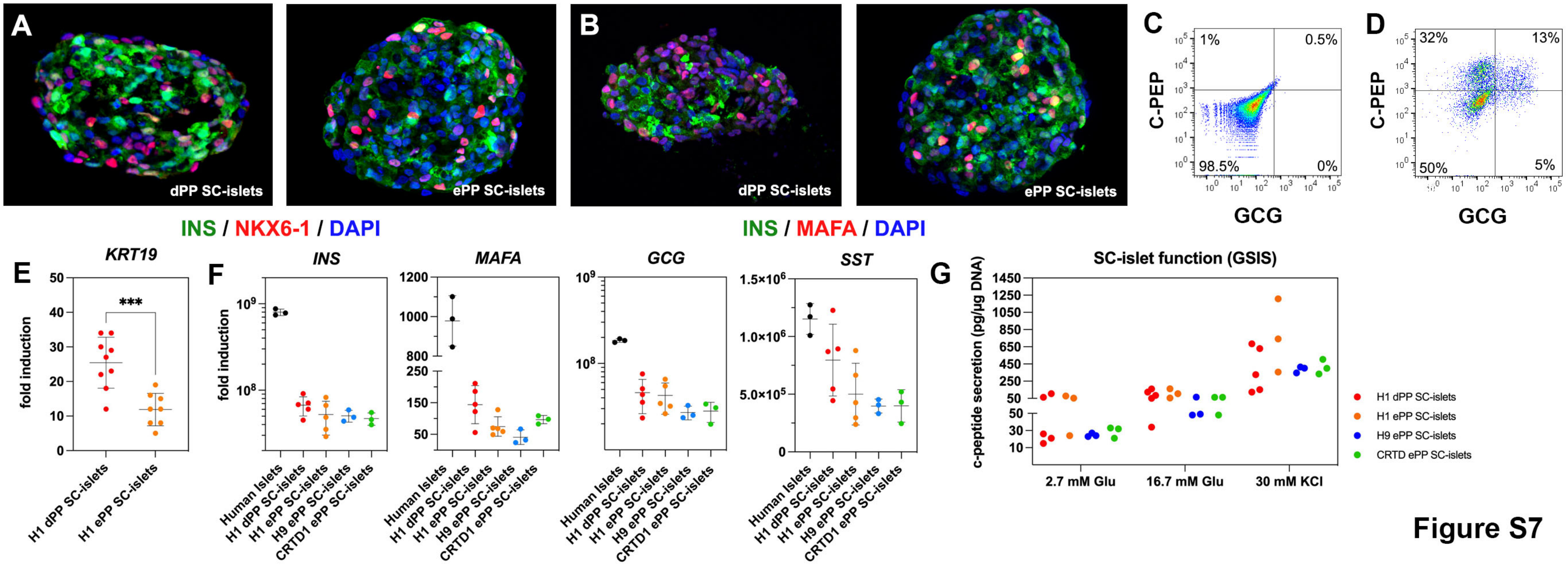
A
dPP SC-islets
ePP SC-islets
INS / NKX6-1 / DAPI
B
dPP SC-islets
ePP SC-islets
INS / MAFA / DAPI
C
C-PEP
GCG
1%
0.5%
98.5%
0%
D
C-PEP
GCG
32%
13%
50%
5%
E
KRT19
fold induction
***
H1 dPP SC-islets
H1 ePP SC-islets
F
INS
MAFA
GCG
SST
fold induction
Human Islets
H1 dPP SC-islets
H1 ePP SC-islets
H9 ePP SC-islets
CRTD1 ePP SC-islets
G
SC-islet function (GSIS)
c-peptide secretion (pg/µg DNA)
2.7 mM Glu
16.7 mM Glu
30 mM KCl
H1 dPP SC-islets
H1 ePP SC-islets
H9 ePP SC-islets
CRTD ePP SC-islets

Figure S7

**Figure S1 Regulated genes and signaling pathways during PP expansion under initial condition (CINI)**
**(A-C)** Immunofluorescence staining of H1-derived PPs showed that expression of PP markers at p0 (A, B) was retained for at least five passages (p5) of expansion under the initial condition (CINI) (C).
**(D)** qPCR analyses indicated that transcript levels of pancreatic progenitor markers were retained at p5 and p10. Values at p5 and p10 were normalized against the value of the corresponding expansion at p0 to give the calculated fold change. Values at p0 were normalized against the average p0 value. Horizontal lines represent the mean ± standard deviation (SD).
**(E)** Stained cryosections of CINI ePP-derived endocrine cell clusters showing expression of endocrine markers.
**(F)** Schematic showing the number of genes significantly up- or down-regulated after filtering the RNA Seq data for 1.6 ≤ fold change ≤ 0.6, normalized counts ≥ 100 and *padj* ≤ 0.05.
**(G-I)** Average transcript levels in normalized RNA Seq counts of p0 (n=4), p5 (n=3) and p10 (n=3) PP cells for genes encoding regulated ligands or receptors of the TGFβ (G), PDGF (H), and NOTCH signaling pathways (I).

Scale bar corresponds to 50 µm (A, B, C).

**Figure S2 Reproducible expansion of PP cells under condition 5 (C5)**
**(A-D)** Gene expression profile of C0-, C1- and C5-expanded cells as shown by qPCR for the PP markers *PTF1A* (A) and *FOXA2* (B) as well as the liver marker *AFP* (C) and the gut marker *CDX2* (D). Expression is normalized against the expression of each marker at p0.
**(E-J)** Representative images of immunofluorescent staining of p0 PP cells and C5-expanded cells at p5 and p10 for FOXA2 (E-G) and AFP/CDX2 expression (H-J).
**(K-M)** Flow cytometry analysis of p0 PP cells and C5-expanded cells at p5 and p10 for $PDX1^+/SOX9^+$ cells.
**(N)** Flow cytometry analysis of PP cells incubated with only secondary antibodies.

Horizontal lines represent the mean ± SD. Statistical tests were two-way ANOVA with Tukey's test, using p0 as the control condition for the comparison with $p \leq 0.033$ (*), $p \leq 0.002$ (**), ≤ 0.0002 (***) and ≤ 0.0001 (****). Scale bar corresponds to 50 µm.

**Figure S3 Expansion of PP cells promotes primarily their proliferation rather than their survival.**
**(A)** Cells without EdU incorporation but processed in the same manner as the sample were used to set the threshold for the quantification.
**(B, C)** Dot plot diagrams of the flow cytometry assay for cell death. Dead cells present in CINI- (B) and C5- (C) ePP cells appear in the upper right quadrant as cells positive for both Annexin V-FITC and 7-AAD

**Figure S4 Reproducible expansion in condition 6 promotes PP identity**

**(A)** Gene expression profile of C6-expanded PP cells p5 (green) and p10 (red) as determined by qPCR. Values are expressed as fold change relative to the expression values in C5-expanded cells at the corresponding passage number.
**(B)** The expression of the hepatic markers *AFP*, *TTR* and *HHEX*, as determined by qPCR, were significantly lower in C6- than in C5- expanded cells by p10. Fold induction was calculated with reference to expression levels at the hPS cell stage.
**(C-H)** Representative images of immunofluorescent staining of p0 PP cells and C6-expanded cells at p5 and p10 for FOXA2 (C-E) and AFP/CDX2 expression (F-H).
**(I-K)** Flow cytometry analysis of p0 PP cells (I) and C6 ePP cells at p5 (J) and p10 (K) for $PDX1^+/SOX9^+$ cells.
**(L)** Flow cytometry analysis of PP cells incubated with only secondary antibodies.
**(M)** Karyotyping of C6-expanded PP cells after sixteen passages showed no chromosomal abnormalities.

Horizontal lines represent the mean ± SD. Statistical tests were two-way ANOVA with Tukey's test, using p0 as the control condition for the comparison with $p \leq 0.033$ (*), $p \leq 0.002$ (**), ≤ 0.0002 (***) and ≤ 0.0001 (****). Scale bar corresponds to 50 μm.

**Figure S5 Expansion under C6 stabilizes PP cell identity by repressing endocrine differentiation.**
**(A)** PCA analysis of our p0 as well as p5 and p10 expanded PP cells.
**(B)** Correlation analyses of the transcriptome profile of p0 PP cells and p5 ePP cells. The numbers of upregulated and downregulated genes (normalized counts ≥ 200 and 0.5 ≥ FC ≥ 2) are shown in red and blue, respectively and r is the correlation coefficient.
**(C)** Most affected molecular functions, cellular compartments and KEGG pathways between p0 and p10.
**(D)** PCA of feeder ePP cells and their corresponding p0 PP cells (shades of green), fibronectin (FN) ePP cells and their corresponding p0 cells (shades of blue), vitronectin-N (VTN-N) ePP cells and corresponding p0 cells (shades of red) as well as different populations of human fetal pancreas progenitor cells (shades of grey). Darker shades correspond to earlier cells.
**(E)** Heat map of the genes contributing to the difference between our ePP cells (VTN-N expanded) and those of others (feeder or FN expanded), derived using the variance stabilizing transformation (vst). Darker shades of red correspond to stronger upregulation.
**(F)** Most affected molecular functions, cellular compartments, biological processes and KEGG pathways in genes that separate our ePP cells (VTN-N expanded) from those of others (feeder or FN expanded).
**(G)** Comparative heat map of the expression levels of duct TFs in feeder or FN or VTN-N ePP and their corresponding p0 cells.
**(H)** Alignment of the mouse Nkx6-1 antigen, used to generate the NKX6-1 antibody, with the human NKX6-1 and NKX6-2 sequences.
**(I-L)** Comparative expression levels of progenitor (I), MPC and BP (J), endocrine (L) as well as liver and gut (J) markers in normalized RNA-Seq counts.

**Figure S6 Expansion of H9-derived and CRTD1-derived PP cells**

**(A-G)** Gene expression profile of PP cells derived from H1, H9 and the CRTD1 hPS cells at p0, p5 and p10 as shown by qPCR for the progenitor markers *PDX1* (A)*, NKX6.1* (B), *SOX9* (C) *FOXA2* (D) and *PTF1A* (E) as well as for the liver marker *AFP* (F) and the gut marker *CDX2* (G). Expression levels are normalized against the expression levels of H1-PP cells at the corresponding passages (p0, p5 and p10).
**(H, I)** Karyotyping of C6-expanded H9-PP cells after twelve passages (H) and C6-expanded CRTD1-PP cells after thirteen passages (I) showed no chromosomal abnormalities.

**Figure S7 Differentiation of ePP cells into SC-islets containing functional β-cells.**
**(A, B)** Immunofluorescence analysis of SC-islets for INS / NKX6-1 expression (A) or INS / MAFA expression (B) derived from p0 PP cells (dPP) or expanded PP cells for at least ten passages (ePP).
**(C, D)** Representative flow cytometry analysis of SC-islet cells incubated with only secondary antibodies (C) or with both C-PEP and GCG primary and corresponding secondary antibodies (D).
**(E)** *KRT19* expression in relation to expression levels in hPS cells (fold induction) as determined by qPCR in SC-islets derived from H1-derived dPP and ePP cells.
**(F)** Expression levels of *INS*, *GCG*, *MAFA*, and *SST* in relation to expression levels in hPS cells (fold induction) as determined by qPCR.
**(G)** Normalized secreted C-peptide levels after exposure to basal conditions with 2.8 mM glucose and following sequential stimulation by 16.7 mM glucose and 16.7 mM glucose / 30 mM KCl (30 mM KCl). The absolute secreted C-peptide levels were normalized against cell numbers (DNA content).

Horizontal lines represent the mean ± SD. Statistical tests were two-way ANOVA with Tukey's test, using p0 as the control condition for the comparison with $p \leq 0.033$ (*), $p \leq 0.002$ (**), ≤ 0.0002 (***) and ≤ 0.0001 (****). Scale bar corresponds to 100 µm (J, K).
