## Supplementary material for "Regulation of multiple signaling pathways promotes the consistent expansion of human pancreatic progenitors in defined conditions": All sup tables

| | | Final concentration | CINI | C0 | C1 | C2 | C3 | C4 | C5 | C6 | C7 | C8 |
|---|---|---|---|---|---|---|---|---|---|---|---|---|
| Basal Components | B-27 Supplement, serum-free with Vitamin A (Gibco, 17504044) | 1X | • | | | | | | | | | |
| | B-27 Supplement, serum-free without Vitamin A (Gibco, 12587010) | 1X | | • | • | • | • | • | • | • | • | • |
| | GlutaMAX (100X) (Gibco, 35050038) | 1X | • | • | • | • | • | • | • | • | • | • |
| | BSA Fatty acid-free, low-endotoxin (20,000X) (LSP, BSA-68700) | 5 ug/ml | • | • | • | • | • | • | • | • | • | • |
| | PEN/STREP (100X) (Gibco, 15140-122) | 1X | • | • | • | • | • | • | • | • | • | • |
| | DMEM High Glucose (Gibco, 21969035) | 1X | • | • | • | • | • | • | • | • | • | • |
| | | Final concentration | CINI | C0 | C1 | C2 | C3 | C4 | C5 | C6 | C7 | C8 |
| Supplements | A83-01, ALK5/4/7 inhibition (R&D systems, 2939/10) | 0.5 uM | • | | | | | | | | | |
| | Human thermostable K128N FGF2 (Biomol ABE-32-8372-10) | 10 ng/ml | • | • | | | | | • | • | • | • |
| | Human EGF, premium grade (Miltenyi Biotec, 130-097-750) | 50 ng/ml | • | • | • | • | • | • | • | • | • | • |
| | StemMACS RepSox, ALK5 inhibition (Miltenyi Biotec, 130-117-340) | 10 uM | | • | • | • | • | • | • | • | • | • |
| | FGF18, Human (GenScript, Z03011) | 20 ng/ml | | | • | • | • | • | • | • | • | • |
| | y-Secretase Inhibitor XXI, Notch inhibition (Millipore, 565790) | 0.1 uM | | | | • | | • | | | | |
| | CP 673451, PDGF inhibition (MedChemExpress, HY-12050) | 0.1 uM | | | | | • | • | | | | |
| | Endo-IWR-1, Canonical Wnt inhibition (PeproTech, 1128234) | 10 uM | | | | | | | | • | | • |
| | LDN 193189, BMP Type I Receptor inhibition (Tocris, 6053) | 0.1 uM | | | | | | | | | • | • |
| | Doubling Time (Td) (days) in H1-PP cells | | ND | 3.9 | 3.6 | no expansion | no expansion | no expansion | 2.3 | 2.2 | 17.3 | 3.5 |

**Table S1** Expansion conditions tested and corresponding doubling times. ND; not determined

| Ensembl ID | Gene Symbol | PP P0 | PP P5 | PP P10 | log2(fold change) P0 to P5 | log2(fold change) P5 to P10 | Description |
|---|---|---|---|---|---|---|---|
| | | | | | **GROUP 1** | | |
| ENSG00000120437 | ACAT2 | 836 | 2212 | 5434 | 1,4 | 1,3 | acetyl-CoA acetyltransferase 2 [Source:HGNC Symbol;Acc:HGNC:94] |
| ENSG00000114948 | ADAM23 | 270 | 583 | 966 | 1,11 | 0,73 | ADAM metallopeptidase domain 23 [Source:HGNC Symbol;Acc:HGNC:202] |
| ENSG00000116771 | AGMAT | 68 | 177 | 313 | 1,39 | 0,82 | agmatinase [Source:HGNC Symbol;Acc:HGNC:18407] |
| ENSG00000106305 | AIMP2 | 294 | 550 | 1011 | 0,91 | 0,88 | aminoacyl tRNA synthetase complex interacting multifunctional protein 2 [Source:HGNC Symbol;Acc:HGNC:20609] |
| ENSG00000100478 | AP4S1 | 96 | 184 | 301 | 0,94 | 0,71 | adaptor related protein complex 4 sigma 1 subunit [Source:HGNC Symbol;Acc:HGNC:575] |
| ENSG00000084674 | APOB | 578 | 1693 | 4647 | 1,55 | 1,46 | apolipoprotein B [Source:HGNC Symbol;Acc:HGNC:603] |
| ENSG00000137135 | ARHGEF39 | 310 | 587 | 1007 | 0,92 | 0,78 | Rho guanine nucleotide exchange factor 39 [Source:HGNC Symbol;Acc:HGNC:25909] |
| ENSG00000105011 | ASF1B | 564 | 1229 | 2080 | 1,12 | 0,76 | anti-silencing function 1B histone chaperone [Source:HGNC Symbol;Acc:HGNC:20996] |
| ENSG00000169136 | ATF5 | 139 | 242 | 805 | 0,8 | 1,74 | activating transcription factor 5 [Source:HGNC Symbol;Acc:HGNC:790] |
| ENSG00000198513 | ATL1 | 99 | 174 | 375 | 0,82 | 1,11 | atlastin GTPase 1 [Source:HGNC Symbol;Acc:HGNC:11231] |
| ENSG00000157087 | ATP2B2 | 13 | 250 | 684 | 4,24 | 1,46 | ATPase plasma membrane Ca2+ transporting 2 [Source:HGNC Symbol;Acc:HGNC:815] |
| ENSG00000172318 | B3GALT1 | 658 | 2191 | 3931 | 1,74 | 0,84 | beta-1,3-galactosyltransferase 1 [Source:HGNC Symbol;Acc:HGNC:916] |
| ENSG00000162630 | B3GALT2 | 37 | 207 | 390 | 2,5 | 0,92 | beta-1,3-galactosyltransferase 2 [Source:HGNC Symbol;Acc:HGNC:917] |
| ENSG00000197299 | BLM | 310 | 629 | 1101 | 1,02 | 0,81 | Bloom syndrome RecQ like helicase [Source:HGNC Symbol;Acc:HGNC:1058] |
| ENSG00000261236 | BOP1 | 331 | 749 | 1447 | 1,18 | 0,95 | block of proliferation 1 [Source:HGNC Symbol;Acc:HGNC:15519] |
| ENSG00000139618 | BRCA2 | 227 | 492 | 952 | 1,12 | 0,95 | BRCA2, DNA repair associated [Source:HGNC Symbol;Acc:HGNC:1101] |
| ENSG00000164076 | CAMKV | 14 | 151 | 310 | 3,41 | 1,04 | CaM kinase like vesicle associated [Source:HGNC Symbol;Acc:HGNC:28788] |
| ENSG00000122483 | CCDC18 | 145 | 304 | 595 | 1,07 | 0,97 | coiled-coil domain containing 18 [Source:HGNC Symbol;Acc:HGNC:30370] |
| ENSG00000175305 | CCNE2 | 25 | 83 | 363 | 1,74 | 2,13 | cyclin E2 [Source:HGNC Symbol;Acc:HGNC:1590] |
| ENSG00000167775 | CD320 | 944 | 1590 | 2549 | 0,75 | 0,68 | CD320 molecule [Source:HGNC Symbol;Acc:HGNC:16692] |
| ENSG00000117877 | CD3EAP | 87 | 197 | 436 | 1,18 | 1,15 | CD3e molecule associated protein [Source:HGNC Symbol;Acc:HGNC:24219] |
| ENSG00000164045 | CDC25A | 384 | 1029 | 2293 | 1,42 | 1,16 | cell division cycle 25A [Source:HGNC Symbol;Acc:HGNC:1725] |
| ENSG00000093009 | CDC45 | 181 | 431 | 863 | 1,26 | 1 | cell division cycle 45 [Source:HGNC Symbol;Acc:HGNC:1739] |
| ENSG00000094804 | CDC6 | 530 | 1099 | 2467 | 1,06 | 1,17 | cell division cycle 6 [Source:HGNC Symbol;Acc:HGNC:1744] |
| ENSG00000176749 | CDK5R1 | 116 | 218 | 362 | 0,92 | 0,73 | cyclin dependent kinase 5 regulatory subunit 1 [Source:HGNC Symbol;Acc:HGNC:1775] |
| ENSG00000245848 | CEBPA | 982 | 1847 | 3442 | 0,91 | 0,9 | CCAAT/enhancer binding protein alpha [Source:HGNC Symbol;Acc:HGNC:1833] |
| ENSG00000117724 | CENPF | 3054 | 5910 | 11883 | 0,95 | 1,01 | centromere protein F [Source:HGNC Symbol;Acc:HGNC:1857] |
| ENSG00000102384 | CENPI | 188 | 416 | 709 | 1,15 | 0,77 | centromere protein I [Source:HGNC Symbol;Acc:HGNC:3968] |
| ENSG00000100162 | CENPM | 164 | 278 | 496 | 0,76 | 0,83 | centromere protein M [Source:HGNC Symbol;Acc:HGNC:18352] |
| ENSG00000166451 | CENPN | 289 | 594 | 999 | 1,04 | 0,75 | centromere protein N [Source:HGNC Symbol;Acc:HGNC:30873] |
| ENSG00000151725 | CENPU | 617 | 1302 | 2800 | 1,08 | 1,11 | centromere protein U [Source:HGNC Symbol;Acc:HGNC:21348] |
| ENSG00000167670 | CHAF1A | 732 | 1699 | 3019 | 1,22 | 0,83 | chromatin assembly factor 1 subunit A [Source:HGNC Symbol;Acc:HGNC:1910] |
| ENSG00000159259 | CHAF1B | 403 | 681 | 1175 | 0,76 | 0,79 | chromatin assembly factor 1 subunit B [Source:HGNC Symbol;Acc:HGNC:1911] |
| ENSG00000128973 | CLN6 | 292 | 489 | 1152 | 0,74 | 1,24 | ceroid-lipofuscinosis, neuronal 6, late infantile, variant [Source:HGNC Symbol;Acc:HGNC:2077] |
| ENSG00000092853 | CLSPN | 300 | 663 | 1633 | 1,15 | 1,3 | claspin [Source:HGNC Symbol;Acc:HGNC:19715] |
| ENSG00000102879 | CORO1A | 50 | 138 | 299 | 1,45 | 1,12 | coronin 1A [Source:HGNC Symbol;Acc:HGNC:2252] |
| ENSG00000080618 | CPB2 | 37 | 73 | 271 | 0,96 | 1,9 | carboxypeptidase B2 [Source:HGNC Symbol;Acc:HGNC:2300] |
| ENSG00000060566 | CREB3L3 | 106 | 236 | 462 | 1,15 | 0,97 | cAMP responsive element binding protein 3 like 3 [Source:HGNC Symbol;Acc:HGNC:18855] |
| ENSG00000171793 | CTPS1 | 556 | 1463 | 2721 | 1,4 | 0,89 | CTP synthase 1 [Source:HGNC Symbol;Acc:HGNC:2519] |
| ENSG00000103018 | CYB5B | 1693 | 2786 | 4638 | 0,72 | 0,74 | cytochrome b5 type B [Source:HGNC Symbol;Acc:HGNC:24374] |
| ENSG00000241095 | CYP51A1P1 | 158 | 263 | 449 | 0,73 | 0,77 | cytochrome P450 family 51 subfamily A member 1 pseudogene 1 [Source:HGNC Symbol;Acc:HGNC:20245] |
| ENSG00000198924 | DCLRE1A | 364 | 588 | 945 | 0,69 | 0,68 | DNA cross-link repair 1A [Source:HGNC Symbol;Acc:HGNC:17660] |
| ENSG00000182810 | DDX28 | 147 | 286 | 526 | 0,96 | 0,88 | DEAD-box helicase 28 [Source:HGNC Symbol;Acc:HGNC:17330] |
| ENSG00000035499 | DEPDC1B | 612 | 1161 | 1970 | 0,92 | 0,76 | DEP domain containing 1B [Source:HGNC Symbol;Acc:HGNC:24902] |
| ENSG00000116133 | DHCR24 | 7416 | 13137 | 27533 | 0,82 | 1,07 | 24-dehydrocholesterol reductase [Source:HGNC Symbol;Acc:HGNC:2859] |
| ENSG00000228716 | DHFR | 1189 | 3012 | 5116 | 1,34 | 0,76 | dihydrofolate reductase [Source:HGNC Symbol;Acc:HGNC:2861] |
| ENSG00000231607 | DLEU2 | 94 | 170 | 373 | 0,86 | 1,13 | deleted in lymphocytic leukemia 2 (non-protein coding) [Source:HGNC Symbol;Acc:HGNC:13748] |
| ENSG00000011332 | DPF1 | 28 | 114 | 421 | 2,02 | 1,89 | double PHD fingers 1 [Source:HGNC Symbol;Acc:HGNC:20225] |
| ENSG00000245750 | DRAIC | 13 | 181 | 561 | 3,87 | 1,63 | downregulated RNA in cancer, inhibitor of cell invasion and migration [Source:HGNC Symbol;Acc:HGNC:27082] |
| ENSG00000136982 | DSCC1 | 122 | 364 | 602 | 1,58 | 0,72 | DNA replication and sister chromatid cohesion 1 [Source:HGNC Symbol;Acc:HGNC:24453] |
| ENSG00000143476 | DTL | 346 | 1239 | 2167 | 1,84 | 0,81 | denticleless E3 ubiquitin protein ligase homolog [Source:HGNC Symbol;Acc:HGNC:30288] |
| ENSG00000101412 | E2F1 | 256 | 901 | 1521 | 1,82 | 0,76 | E2F transcription factor 1 [Source:HGNC Symbol;Acc:HGNC:3113] |
| ENSG00000007968 | E2F2 | 108 | 261 | 692 | 1,28 | 1,41 | E2F transcription factor 2 [Source:HGNC Symbol;Acc:HGNC:3114] |
| ENSG00000165891 | E2F7 | 398 | 895 | 1917 | 1,17 | 1,1 | E2F transcription factor 7 [Source:HGNC Symbol;Acc:HGNC:23820] |
| ENSG00000118894 | EEF2KMT | 110 | 216 | 386 | 0,98 | 0,84 | eukaryotic elongation factor 2 lysine methyltransferase [Source:HGNC Symbol;Acc:HGNC:32221] |
| ENSG00000141642 | ELAC1 | 80 | 148 | 324 | 0,88 | 1,13 | elaC ribonuclease Z 1 [Source:HGNC Symbol;Acc:HGNC:14197] |
| ENSG00000186871 | ERCC6L | 168 | 408 | 666 | 1,28 | 0,71 | ERCC excision repair 6 like, spindle assembly checkpoint helicase [Source:HGNC Symbol;Acc:HGNC:20794] |
| ENSG00000171320 | ESCO2 | 129 | 424 | 717 | 1,72 | 0,76 | establishment of sister chromatid cohesion N-acetyltransferase 2 [Source:HGNC Symbol;Acc:HGNC:27230] |
| ENSG00000174371 | EXO1 | 194 | 498 | 857 | 1,36 | 0,78 | exonuclease 1 [Source:HGNC Symbol;Acc:HGNC:3511] |
| ENSG00000026103 | FAS | 90 | 156 | 314 | 0,78 | 1,01 | Fas cell surface death receptor [Source:HGNC Symbol;Acc:HGNC:11920] |
| ENSG00000112029 | FBXO5 | 443 | 1046 | 1748 | 1,24 | 0,74 | F-box protein 5 [Source:HGNC Symbol;Acc:HGNC:13584] |
| ENSG00000168496 | FEN1 | 617 | 1326 | 2970 | 1,1 | 1,16 | flap structure-specific endonuclease 1 [Source:HGNC Symbol;Acc:HGNC:3650] |
| ENSG00000156427 | FGF18 | 332 | 8917 | 15769 | 4,75 | 0,82 | fibroblast growth factor 18 [Source:HGNC Symbol;Acc:HGNC:3674] |
| ENSG00000100350 | FOXRED2 | 466 | 920 | 1799 | 0,98 | 0,97 | FAD dependent oxidoreductase domain containing 2 [Source:HGNC Symbol;Acc:HGNC:26264] |
| ENSG00000089280 | FUS | 5614 | 10051 | 24940 | 0,84 | 1,31 | FUS RNA binding protein [Source:HGNC Symbol;Acc:HGNC:4010] |
| ENSG00000100116 | GCAT | 111 | 523 | 880 | 2,23 | 0,75 | glycine C-acetyltransferase [Source:HGNC Symbol;Acc:HGNC:4188] |
| ENSG00000145990 | GFOD1 | 44 | 299 | 547 | 2,76 | 0,87 | glucose-fructose oxidoreductase domain containing 1 [Source:HGNC Symbol;Acc:HGNC:21096] |
| ENSG00000101003 | GINS1 | 647 | 1070 | 1726 | 0,73 | 0,69 | GINS complex subunit 1 [Source:HGNC Symbol;Acc:HGNC:28980] |
| ENSG00000131153 | GINS2 | 275 | 682 | 1329 | 1,31 | 0,96 | GINS complex subunit 2 [Source:HGNC Symbol;Acc:HGNC:24575] |
| ENSG00000181938 | GINS3 | 164 | 270 | 733 | 0,72 | 1,44 | GINS complex subunit 3 [Source:HGNC Symbol;Acc:HGNC:25851] |
| ENSG00000147536 | GINS4 | 512 | 1085 | 1753 | 1,08 | 0,69 | GINS complex subunit 4 [Source:HGNC Symbol;Acc:HGNC:28226] |
| ENSG00000160818 | GPATCH4 | 339 | 684 | 1327 | 1,02 | 0,96 | G-patch domain containing 4 [Source:HGNC Symbol;Acc:HGNC:25982] |
| ENSG00000198822 | GRM3 | 51 | 266 | 524 | 2,37 | 0,98 | glutamate metabotropic receptor 3 [Source:HGNC Symbol;Acc:HGNC:4595] |
| ENSG00000188486 | H2AFX | 1755 | 2965 | 4868 | 0,76 | 0,72 | H2A histone family member X [Source:HGNC Symbol;Acc:HGNC:4739] |
| ENSG00000164032 | H2AFZ | 5656 | 10058 | 17776 | 0,83 | 0,82 | H2A histone family member Z [Source:HGNC Symbol;Acc:HGNC:4741] |
| ENSG00000119969 | HELLS | 674 | 1302 | 2103 | 0,95 | 0,69 | helicase, lymphoid-specific [Source:HGNC Symbol;Acc:HGNC:4861] |
| ENSG00000144485 | HES6 | 552 | 1474 | 2936 | 1,42 | 0,99 | hes family bHLH transcription factor 6 [Source:HGNC Symbol;Acc:HGNC:18254] |
| ENSG00000135547 | HEY2 | 30 | 238 | 439 | 2,99 | 0,88 | hes related family bHLH transcription factor with YRPW motif 2 [Source:HGNC Symbol;Acc:HGNC:4881] |
| ENSG00000214821 | HMGB1P4 | 118 | 193 | 350 | 0,72 | 0,86 | high mobility group box 1 pseudogene 4 [Source:HGNC Symbol;Acc:HGNC:4996] |
| ENSG00000112972 | HMGCS1 | 3875 | 8379 | 15710 | 1,11 | 0,91 | 3-hydroxy-3-methylglutaryl-CoA synthase 1 [Source:HGNC Symbol;Acc:HGNC:5007] |
| ENSG00000198830 | HMGN2 | 3467 | 7851 | 12849 | 1,18 | 0,71 | high mobility group nucleosomal binding domain 2 [Source:HGNC Symbol;Acc:HGNC:4986] |
| ENSG00000249014 | HMGN2P4 | 209 | 443 | 772 | 1,09 | 0,8 | high mobility group nucleosomal binding domain 2 pseudogene 4 [Source:HGNC Symbol;Acc:HGNC:33567] |
| ENSG00000234664 | HMGN2P5 | 178 | 451 | 867 | 1,34 | 0,94 | high mobility group nucleosomal binding domain 2 pseudogene 5 [Source:HGNC Symbol;Acc:HGNC:33568] |
| ENSG00000099783 | HNRNPM | 3111 | 5951 | 17956 | 0,94 | 1,59 | heterogeneous nuclear ribonucleoprotein M [Source:HGNC Symbol;Acc:HGNC:5046] |
| ENSG00000067064 | IDI1 | 986 | 1971 | 3208 | 1 | 0,7 | isopentenyl-diphosphate delta isomerase 1 [Source:HGNC Symbol;Acc:HGNC:5387] |
| ENSG00000099869 | IGF2-AS | 843 | 1559 | 3104 | 0,89 | 0,99 | IGF2 antisense RNA [Source:HGNC Symbol;Acc:HGNC:14062] |
| ENSG00000166130 | IKBIP | 446 | 889 | 1486 | 1 | 0,74 | IKBKB interacting protein [Source:HGNC Symbol;Acc:HGNC:26430] |
| ENSG00000156113 | KCNMA1 | 12 | 100 | 435 | 3,02 | 2,12 | potassium calcium-activated channel subfamily M alpha 1 [Source:HGNC Symbol;Acc:HGNC:6284] |
| ENSG00000155666 | KDM8 | 86 | 149 | 251 | 0,8 | 0,75 | lysine demethylase 8 [Source:HGNC Symbol;Acc:HGNC:25840] |
| ENSG00000259417 | LINC01314 | 276 | 531 | 1020 | 0,94 | 0,94 | long intergenic non-protein coding RNA 1314 [Source:HGNC Symbol;Acc:HGNC:50507] |
| ENSG00000166035 | LIPC | 751 | 2085 | 3671 | 1,47 | 0,82 | lipase C, hepatic type [Source:HGNC Symbol;Acc:HGNC:6619] |
| ENSG00000113368 | LMNB1 | 2851 | 6416 | 10920 | 1,17 | 0,77 | lamin B1 [Source:HGNC Symbol;Acc:HGNC:6637] |
| ENSG00000157193 | LRP8 | 66 | 331 | 611 | 2,33 | 0,88 | LDL receptor related protein 8 [Source:HGNC Symbol;Acc:HGNC:6700] |
| ENSG00000128606 | LRRC17 | 123 | 668 | 1079 | 2,44 | 0,69 | leucine rich repeat containing 17 [Source:HGNC Symbol;Acc:HGNC:16895] |
| ENSG00000065328 | MCM10 | 260 | 751 | 1667 | 1,54 | 1,15 | minichromosome maintenance 10 replication initiation factor [Source:HGNC Symbol;Acc:HGNC:18043] |
| ENSG00000073111 | MCM2 | 1510 | 4386 | 7362 | 1,54 | 0,75 | minichromosome maintenance complex component 2 [Source:HGNC Symbol;Acc:HGNC:6944] |
| ENSG00000104738 | MCM4 | 2119 | 3921 | 7116 | 0,89 | 0,86 | minichromosome maintenance complex component 4 [Source:HGNC Symbol;Acc:HGNC:6947] |
| ENSG00000100297 | MCM5 | 1015 | 2463 | 3985 | 1,28 | 0,69 | minichromosome maintenance complex component 5 [Source:HGNC Symbol;Acc:HGNC:6948] |
| ENSG00000076003 | MCM6 | 1279 | 2465 | 4031 | 0,95 | 0,71 | minichromosome maintenance complex component 6 [Source:HGNC Symbol;Acc:HGNC:6949] |
| ENSG00000166508 | MCM7 | 3791 | 8110 | 13970 | 1,1 | 0,78 | minichromosome maintenance complex component 7 [Source:HGNC Symbol;Acc:HGNC:6950] |
| ENSG00000172878 | METAP1D | 64 | 105 | 183 | 0,72 | 0,8 | methionyl aminopeptidase type 1D, mitochondrial [Source:HGNC Symbol;Acc:HGNC:32583] |
| ENSG00000037897 | METTL1 | 87 | 190 | 361 | 1,13 | 0,92 | methyltransferase like 1 [Source:HGNC Symbol;Acc:HGNC:7030] |
| ENSG00000170854 | MINA | 135 | 348 | 612 | 1,37 | 0,82 | MYC induced nuclear antigen [Source:HGNC Symbol;Acc:HGNC:19441] |
| ENSG00000215417 | MIR17HG | 48 | 84 | 212 | 0,82 | 1,34 | miR-17-92a-1 cluster host gene [Source:HGNC Symbol;Acc:HGNC:23564] |
| ENSG00000148773 | MKI67 | 3675 | 5927 | 9508 | 0,69 | 0,68 | marker of proliferation Ki-67 [Source:HGNC Symbol;Acc:HGNC:7107] |
| ENSG00000164077 | MON1A | 93 | 169 | 285 | 0,87 | 0,75 | MON1 homolog A, secretory trafficking associated [Source:HGNC Symbol;Acc:HGNC:28207] |
| ENSG00000175806 | MSRA | 115 | 233 | 393 | 1,03 | 0,75 | methionine sulfoxide reductase A [Source:HGNC Symbol;Acc:HGNC:7377] |
| ENSG00000100714 | MTHFD1 | 757 | 2143 | 3584 | 1,5 | 0,74 | methylenetetrahydrofolate dehydrogenase, cyclohydrolase and formyltetrahydrofolate synthetase 1 [Source:HGNC Symbol;Acc:HGNC:7432] |
| ENSG00000110921 | MVK | 342 | 564 | 1076 | 0,72 | 0,93 | mevalonate kinase [Source:HGNC Symbol;Acc:HGNC:7530] |
| ENSG00000118513 | MYB | 16 | 84 | 250 | 2,39 | 1,57 | MYB proto-oncogene, transcription factor [Source:HGNC Symbol;Acc:HGNC:7545] |
| ENSG00000101057 | MYBL2 | 1214 | 2369 | 3812 | 0,97 | 0,69 | MYB proto-oncogene like 2 [Source:HGNC Symbol;Acc:HGNC:7548] |
| ENSG00000109805 | NCAPG | 821 | 2083 | 3474 | 1,34 | 0,74 | non-SMC condensin I complex subunit G [Source:HGNC Symbol;Acc:HGNC:24304] |
| ENSG00000025770 | NCAPH2 | 831 | 1361 | 2245 | 0,71 | 0,72 | non-SMC condensin II complex subunit H2 [Source:HGNC Symbol;Acc:HGNC:25071] |
| ENSG00000177707 | NECTIN3 | 610 | 1676 | 2828 | 1,46 | 0,75 | nectin cell adhesion molecule 3 [Source:HGNC Symbol;Acc:HGNC:17664] |
| ENSG00000171208 | NETO2 | 868 | 1504 | 3314 | 0,79 | 1,14 | neuropilin and tolloid like 2 [Source:HGNC Symbol;Acc:HGNC:14644] |
| ENSG00000239672 | NME1 | 222 | 513 | 1141 | 1,21 | 1,15 | NME/NM23 nucleoside diphosphate kinase 1 [Source:HGNC Symbol;Acc:HGNC:7849] |
| ENSG00000055044 | NOP58 | 1007 | 1690 | 3232 | 0,75 | 0,94 | NOP58 ribonucleoprotein [Source:HGNC Symbol;Acc:HGNC:29926] |
| ENSG00000181019 | NQO1 | 359 | 1864 | 6258 | 2,38 | 1,75 | NAD(P)H quinone dehydrogenase 1 [Source:HGNC Symbol;Acc:HGNC:2874] |
| ENSG00000131910 | NR0B2 | 216 | 562 | 1383 | 1,38 | 1,3 | nuclear receptor subfamily 0 group B member 2 [Source:HGNC Symbol;Acc:HGNC:7961] |
| ENSG00000119900 | OGFRL1 | 1101 | 1792 | 2896 | 0,7 | 0,69 | opioid growth factor receptor like 1 [Source:HGNC Symbol;Acc:HGNC:21378] |
| ENSG00000162073 | PAQR4 | 222 | 366 | 900 | 0,72 | 1,3 | progestin and adipoQ receptor family member 4 [Source:HGNC Symbol;Acc:HGNC:26386] |
| ENSG00000132646 | PCNA | 1272 | 2392 | 6003 | 0,91 | 1,33 | proliferating cell nuclear antigen [Source:HGNC Symbol;Acc:HGNC:8729] |
| ENSG00000163710 | PCOLCE2 | 44 | 186 | 309 | 2,06 | 0,73 | procollagen C-endopeptidase enhancer 2 [Source:HGNC Symbol;Acc:HGNC:8739] |
| ENSG00000178921 | PFAS | 923 | 2088 | 3424 | 1,18 | 0,71 | phosphoribosylformylglycinamidine synthase [Source:HGNC Symbol;Acc:HGNC:8863] |
| ENSG00000277161 | PIGW | 103 | 227 | 514 | 1,13 | 1,18 | phosphatidylinositol glycan anchor biosynthesis class W [Source:HGNC Symbol;Acc:HGNC:23213] |
| ENSG00000117461 | PIK3R3 | 591 | 1541 | 2986 | 1,38 | 0,95 | phosphoinositide-3-kinase regulatory subunit 3 [Source:HGNC Symbol;Acc:HGNC:8981] |
| ENSG00000100078 | PLA2G3 | 33 | 130 | 336 | 1,98 | 1,37 | phospholipase A2 group III [Source:HGNC Symbol;Acc:HGNC:17934] |
| ENSG00000052126 | PLEKHA5 | 1527 | 2877 | 4674 | 0,91 | 0,7 | pleckstrin homology domain containing A5 [Source:HGNC Symbol;Acc:HGNC:30036] |
| ENSG00000014138 | POLA2 | 561 | 938 | 1708 | 0,74 | 0,86 | DNA polymerase alpha 2, accessory subunit [Source:HGNC Symbol;Acc:HGNC:30073] |
| ENSG00000100479 | POLE2 | 101 | 327 | 565 | 1,7 | 0,79 | DNA polymerase epsilon 2, accessory subunit [Source:HGNC Symbol;Acc:HGNC:9178] |
| ENSG00000177700 | POLR2L | 766 | 1238 | 2140 | 0,69 | 0,79 | RNA polymerase II subunit L [Source:HGNC Symbol;Acc:HGNC:9199] |
| ENSG00000221988 | PPT2 | 409 | 697 | 1229 | 0,77 | 0,82 | palmitoyl-protein thioesterase 2 [Source:HGNC Symbol;Acc:HGNC:9326] |
| ENSG00000057657 | PRDM1 | 42 | 124 | 387 | 1,58 | 1,64 | PR/SET domain 1 [Source:HGNC Symbol;Acc:HGNC:9346] |
| ENSG00000198890 | PRMT6 | 413 | 734 | 1245 | 0,83 | 0,76 | protein arginine methyltransferase 6 [Source:HGNC Symbol;Acc:HGNC:18241] |
| ENSG00000250799 | PRODH2 | 29 | 168 | 283 | 2,55 | 0,75 | proline dehydrogenase 2 [Source:HGNC Symbol;Acc:HGNC:17325] |
| ENSG00000135069 | PSAT1 | 599 | 2526 | 5716 | 2,08 | 1,18 | phosphoserine aminotransferase 1 [Source:HGNC Symbol;Acc:HGNC:19129] |
| ENSG00000165916 | PSMC3 | 2063 | 3440 | 5577 | 0,74 | 0,7 | proteasome 26S subunit, ATPase 3 [Source:HGNC Symbol;Acc:HGNC:9549] |
| ENSG00000104524 | PYCRL | 201 | 386 | 776 | 0,95 | 1,01 | pyrroline-5-carboxylate reductase-like [Source:HGNC Symbol;Acc:HGNC:25846] |
| ENSG00000152932 | RAB3C | 231 | 717 | 1543 | 1,63 | 1,11 | RAB3C, member RAS oncogene family [Source:HGNC Symbol;Acc:HGNC:30269] |
| ENSG00000051180 | RAD51 | 243 | 399 | 677 | 0,72 | 0,76 | RAD51 recombinase [Source:HGNC Symbol;Acc:HGNC:9817] |
| ENSG00000099901 | RANBP1 | 2008 | 3401 | 5681 | 0,76 | 0,74 | RAN binding protein 1 [Source:HGNC Symbol;Acc:HGNC:9847] |
| ENSG00000080839 | RBL1 | 200 | 437 | 988 | 1,13 | 1,18 | RB transcriptional corepressor like 1 [Source:HGNC Symbol;Acc:HGNC:9893] |
| ENSG00000160957 | RECQL4 | 715 | 1145 | 2001 | 0,68 | 0,81 | RecQ like helicase 4 [Source:HGNC Symbol;Acc:HGNC:9949] |
| ENSG00000133119 | RFC3 | 343 | 907 | 1452 | 1,41 | 0,68 | replication factor C subunit 3 [Source:HGNC Symbol;Acc:HGNC:9971] |
| ENSG00000234171 | RNASEH1-AS1 | 65 | 118 | 206 | 0,87 | 0,81 | RNASEH1 antisense RNA 1 [Source:HGNC Symbol;Acc:HGNC:49289] |
| ENSG00000171848 | RRM2 | 1233 | 2698 | 6885 | 1,13 | 1,35 | ribonucleotide reductase regulatory subunit M2 [Source:HGNC Symbol;Acc:HGNC:10452] |
| ENSG00000179041 | RRS1 | 166 | 356 | 575 | 1,11 | 0,69 | ribosome biogenesis regulator homolog [Source:HGNC Symbol;Acc:HGNC:17083] |
| ENSG00000144285 | SCN1A | 74 | 347 | 1095 | 2,22 | 1,66 | sodium voltage-gated channel alpha subunit 1 [Source:HGNC Symbol;Acc:HGNC:10585] |
| ENSG00000170099 | SERPINA6 | 418 | 886 | 1554 | 1,08 | 0,81 | serpin family A member 6 [Source:HGNC Symbol;Acc:HGNC:1540] |
| ENSG00000156398 | SFXN2 | 166 | 323 | 983 | 0,96 | 1,6 | sideroflexin 2 [Source:HGNC Symbol;Acc:HGNC:16086] |
| ENSG00000165480 | SKA3 | 216 | 634 | 1073 | 1,55 | 0,76 | spindle and kinetochore associated complex subunit 3 [Source:HGNC Symbol;Acc:HGNC:20262] |
| ENSG00000115902 | SLC1A4 | 64 | 159 | 583 | 1,34 | 1,87 | solute carrier family 1 member 4 [Source:HGNC Symbol;Acc:HGNC:10942] |
| ENSG00000145217 | SLC26A1 | 37 | 111 | 321 | 1,61 | 1,53 | solute carrier family 26 member 1 [Source:HGNC Symbol;Acc:HGNC:10993] |
| ENSG00000142494 | SLC47A1 | 59 | 351 | 851 | 2,57 | 1,28 | solute carrier family 47 member 1 [Source:HGNC Symbol;Acc:HGNC:25588] |
| ENSG00000139514 | SLC7A1 | 954 | 1553 | 2543 | 0,7 | 0,71 | solute carrier family 7 member 1 [Source:HGNC Symbol;Acc:HGNC:11057] |
| ENSG00000151012 | SLC7A11 | 75 | 294 | 1122 | 1,97 | 1,93 | solute carrier family 7 member 11 [Source:HGNC Symbol;Acc:HGNC:11059] |
| ENSG00000157734 | SNX22 | 151 | 249 | 595 | 0,72 | 1,26 | sorting nexin 22 [Source:HGNC Symbol;Acc:HGNC:16315] |
| ENSG00000140263 | SORD | 1202 | 2013 | 3544 | 0,74 | 0,82 | sorbitol dehydrogenase [Source:HGNC Symbol;Acc:HGNC:11184] |
| ENSG00000259479 | SORD2P | 1657 | 2768 | 4863 | 0,74 | 0,81 | sorbitol dehydrogenase 2, pseudogene [Source:HGNC Symbol;Acc:HGNC:49919] |
| ENSG00000104549 | SQLE | 2193 | 3660 | 6358 | 0,74 | 0,8 | squalene epoxidase [Source:HGNC Symbol;Acc:HGNC:11279] |
| ENSG00000161547 | SRSF2 | 3041 | 5636 | 10125 | 0,89 | 0,85 | serine and arginine rich splicing factor 2 [Source:HGNC Symbol;Acc:HGNC:10783] |
| ENSG00000112081 | SRSF3 | 3736 | 6942 | 13388 | 0,89 | 0,95 | serine and arginine rich splicing factor 3 [Source:HGNC Symbol;Acc:HGNC:10785] |
| ENSG00000147465 | STAR | 38 | 191 | 721 | 2,36 | 1,92 | steroidogenic acute regulatory protein [Source:HGNC Symbol;Acc:HGNC:11359] |
| ENSG00000197457 | STMN3 | 660 | 1686 | 2872 | 1,35 | 0,77 | stathmin 3 [Source:HGNC Symbol;Acc:HGNC:15926] |
| ENSG00000182253 | SYNM | 54 | 218 | 510 | 2,02 | 1,23 | synemin [Source:HGNC Symbol;Acc:HGNC:24466] |
| ENSG00000187325 | TAF9B | 240 | 584 | 961 | 1,28 | 0,72 | TATA-box binding protein associated factor 9b [Source:HGNC Symbol;Acc:HGNC:17306] |

| | | | | | | | |
|---|---|---|---|---|---|---|---|
| ENSG00000136111 | TBC1D4 | 766 | 1334 | 2186 | 0,8 | 0,71 | TBC1 domain family member 4 [Source:HGNC Symbol;Acc:HGNC:19165] |
| ENSG00000075131 | TIPIN | 92 | 164 | 302 | 0,84 | 0,88 | TIMELESS interacting protein [Source:HGNC Symbol;Acc:HGNC:30750] |
| ENSG00000158164 | TMSB15A | 309 | 720 | 1291 | 1,22 | 0,84 | thymosin beta 15a [Source:HGNC Symbol;Acc:HGNC:30744] |
| ENSG00000132773 | TOE1 | 122 | 197 | 626 | 0,69 | 1,67 | target of EGR1, member 1 (nuclear) [Source:HGNC Symbol;Acc:HGNC:15954] |
| ENSG00000186854 | TRABD2A | 64 | 203 | 717 | 1,67 | 1,82 | TraB domain containing 2A [Source:HGNC Symbol;Acc:HGNC:27013] |
| ENSG00000183763 | TRAIP | 247 | 402 | 671 | 0,7 | 0,74 | TRAF interacting protein [Source:HGNC Symbol;Acc:HGNC:30764] |
| ENSG00000104691 | UBXN8 | 137 | 260 | 435 | 0,92 | 0,75 | UBX domain protein 8 [Source:HGNC Symbol;Acc:HGNC:30307] |
| ENSG00000154277 | UCHL1 | 1616 | 2978 | 5276 | 0,88 | 0,83 | ubiquitin C-terminal hydrolase L1 [Source:HGNC Symbol;Acc:HGNC:12513] |
| ENSG00000276043 | UHRF1 | 426 | 1134 | 2694 | 1,41 | 1,25 | ubiquitin like with PHD and ring finger domains 1 [Source:HGNC Symbol;Acc:HGNC:12556] |
| ENSG00000135763 | URB2 | 234 | 429 | 885 | 0,87 | 1,04 | URB2 ribosome biogenesis 2 homolog (S. cerevisiae) [Source:HGNC Symbol;Acc:HGNC:28967] |
| ENSG00000162607 | USP1 | 1414 | 2785 | 4486 | 0,98 | 0,69 | ubiquitin specific peptidase 1 [Source:HGNC Symbol;Acc:HGNC:12607] |
| ENSG00000120800 | UTP20 | 407 | 711 | 1450 | 0,8 | 1,03 | UTP20, small subunit processome component [Source:HGNC Symbol;Acc:HGNC:17897] |
| ENSG00000038427 | VCAN | 27717 | 68892 | 111842 | 1,31 | 0,7 | versican [Source:HGNC Symbol;Acc:HGNC:2464] |
| ENSG00000160193 | WDR4 | 125 | 238 | 485 | 0,93 | 1,03 | WD repeat domain 4 [Source:HGNC Symbol;Acc:HGNC:12756] |
| ENSG00000075702 | WDR62 | 438 | 736 | 1440 | 0,75 | 0,97 | WD repeat domain 62 [Source:HGNC Symbol;Acc:HGNC:24502] |
| ENSG00000126215 | XRCC3 | 350 | 582 | 1112 | 0,73 | 0,93 | X-ray repair cross complementing 3 [Source:HGNC Symbol;Acc:HGNC:12830] |
| ENSG00000152422 | XRCC4 | 751 | 1245 | 2066 | 0,73 | 0,73 | X-ray repair cross complementing 4 [Source:HGNC Symbol;Acc:HGNC:12831] |
| ENSG00000132846 | ZBED3 | 1022 | 1709 | 2814 | 0,74 | 0,72 | zinc finger BED-type containing 3 [Source:HGNC Symbol;Acc:HGNC:20711] |
| ENSG00000187801 | ZFP69B | 68 | 126 | 219 | 0,9 | 0,8 | ZFP69 zinc finger protein B [Source:HGNC Symbol;Acc:HGNC:28053] |
| ENSG00000165244 | ZNF367 | 152 | 288 | 487 | 0,92 | 0,76 | zinc finger protein 367 [Source:HGNC Symbol;Acc:HGNC:18320] |
| ENSG00000144792 | ZNF660 | 165 | 273 | 492 | 0,73 | 0,85 | zinc finger protein 660 [Source:HGNC Symbol;Acc:HGNC:26720] |
| ENSG00000179965 | ZNF771 | 137 | 267 | 477 | 0,96 | 0,84 | zinc finger protein 771 [Source:HGNC Symbol;Acc:HGNC:29653] |
| ENSG00000122952 | ZWINT | 537 | 1757 | 3442 | 1,71 | 0,97 | ZW10 interacting kinetochore protein [Source:HGNC Symbol;Acc:HGNC:13195] |

**GROUP 2**

| | | | | | | | |
|---|---|---|---|---|---|---|---|
| ENSG00000183044 | ABAT | 400 | 917 | 1334 | 1,2 | 0,54 | 4-aminobutyrate aminotransferase [Source:HGNC Symbol;Acc:HGNC:23] |
| ENSG00000144827 | ABHD10 | 746 | 1320 | 1046 | 0,82 | -0,34 | abhydrolase domain containing 10 [Source:HGNC Symbol;Acc:HGNC:25656] |
| ENSG00000254893 | AC113404.1 | 1669 | 4678 | 3554 | 1,49 | -0,4 | Ras-related protein Rap-1b-like protein [Source:UniProtKB/Swiss-Prot;Acc:A6NIZ1] |
| ENSG00000117054 | ACADM | 809 | 1843 | 2288 | 1,19 | 0,31 | acyl-CoA dehydrogenase, C-4 to C-12 straight chain [Source:HGNC Symbol;Acc:HGNC:89] |
| ENSG00000136518 | ACTL6A | 1098 | 1855 | 2574 | 0,76 | 0,47 | actin like 6A [Source:HGNC Symbol;Acc:HGNC:24124] |
| ENSG00000008277 | ADAM22 | 261 | 553 | 807 | 1,09 | 0,55 | ADAM metallopeptidase domain 22 [Source:HGNC Symbol;Acc:HGNC:201] |
| ENSG00000239900 | ADSL | 1461 | 2472 | 3052 | 0,76 | 0,3 | adenylosuccinate lyase [Source:HGNC Symbol;Acc:HGNC:291] |
| ENSG00000155966 | AFF2 | 226 | 2218 | 1485 | 3,3 | -0,58 | AF4/FMR2 family member 2 [Source:HGNC Symbol;Acc:HGNC:3776] |
| ENSG00000038002 | AGA | 343 | 668 | 1027 | 0,96 | 0,62 | aspartylglucosaminidase [Source:HGNC Symbol;Acc:HGNC:318] |
| ENSG00000123505 | AMD1 | 1996 | 3360 | 4252 | 0,75 | 0,34 | adenosylmethionine decarboxylase 1 [Source:HGNC Symbol;Acc:HGNC:457] |
| ENSG00000135409 | AMHR2 | 554 | 1359 | 1895 | 1,3 | 0,48 | anti-Mullerian hormone receptor type 2 [Source:HGNC Symbol;Acc:HGNC:465] |
| ENSG00000101935 | AMMECR1 | 684 | 1110 | 1570 | 0,7 | 0,5 | Alport syndrome, mental retardation, midface hypoplasia and elliptocytosis chromosomal region gene 1 [Source:HGNC Symbol;Acc:HGNC:467] |
| ENSG00000196510 | ANAPC7 | 717 | 1160 | 1598 | 0,69 | 0,46 | anaphase promoting complex subunit 7 [Source:HGNC Symbol;Acc:HGNC:17380] |
| ENSG00000011426 | ANLN | 1889 | 3862 | 5240 | 1,03 | 0,44 | anillin actin binding protein [Source:HGNC Symbol;Acc:HGNC:14082] |
| ENSG00000197043 | ANXA6 | 2944 | 6789 | 4803 | 1,21 | -0,5 | annexin A6 [Source:HGNC Symbol;Acc:HGNC:544] |
| ENSG00000112379 | ARFGEF3 | 4147 | 6802 | 4325 | 0,71 | -0,65 | ARFGEF family member 3 [Source:HGNC Symbol;Acc:HGNC:21213] |
| ENSG00000187951 | ARHGAP11B | 631 | 1119 | 1525 | 0,83 | 0,45 | Rho GTPase activating protein 11B [Source:HGNC Symbol;Acc:HGNC:15782] |
| ENSG00000105676 | ARMC6 | 585 | 980 | 1510 | 0,75 | 0,62 | armadillo repeat containing 6 [Source:HGNC Symbol;Acc:HGNC:25049] |
| ENSG00000172379 | ARNT2 | 1962 | 3207 | 4197 | 0,71 | 0,39 | aryl hydrocarbon receptor nuclear translocator 2 [Source:HGNC Symbol;Acc:HGNC:16876] |
| ENSG00000111875 | ASF1A | 555 | 1031 | 1274 | 0,89 | 0,31 | anti-silencing function 1A histone chaperone [Source:HGNC Symbol;Acc:HGNC:20995] |
| ENSG00000115966 | ATF2 | 1673 | 3117 | 2606 | 0,9 | -0,26 | activating transcription factor 2 [Source:HGNC Symbol;Acc:HGNC:784] |
| ENSG00000169857 | AVEN | 163 | 307 | 433 | 0,92 | 0,5 | apoptosis and caspase activation inhibitor [Source:HGNC Symbol;Acc:HGNC:13509] |
| ENSG00000118276 | B4GALT6 | 1112 | 1944 | 1360 | 0,81 | -0,52 | beta-1,4-galactosyltransferase 6 [Source:HGNC Symbol;Acc:HGNC:929] |
| ENSG00000174684 | B4GAT1 | 1432 | 2419 | 1788 | 0,76 | -0,44 | beta-1,4-glucuronyltransferase 1 [Source:HGNC Symbol;Acc:HGNC:15685] |
| ENSG00000137936 | BCAR3 | 215 | 1300 | 2031 | 2,6 | 0,64 | breast cancer anti-estrogen resistance 3 [Source:HGNC Symbol;Acc:HGNC:973] |
| ENSG00000107949 | BCCIP | 1187 | 2048 | 2798 | 0,79 | 0,45 | BRCA2 and CDKN1A interacting protein [Source:HGNC Symbol;Acc:HGNC:978] |
| ENSG00000144857 | BOC | 284 | 574 | 351 | 1,02 | -0,71 | BOC cell adhesion associated, oncogene regulated [Source:HGNC Symbol;Acc:HGNC:17173] |
| ENSG00000012048 | BRCA1 | 551 | 993 | 1548 | 0,85 | 0,64 | BRCA1, DNA repair associated [Source:HGNC Symbol;Acc:HGNC:1100] |
| ENSG00000108561 | C1QBP | 1395 | 2567 | 3533 | 0,88 | 0,46 | complement C1q binding protein [Source:HGNC Symbol;Acc:HGNC:1243] |
| ENSG00000253250 | C8orf88 | 189 | 312 | 218 | 0,72 | -0,52 | chromosome 8 open reading frame 88 [Source:HGNC Symbol;Acc:HGNC:44672] |
| ENSG00000163006 | CCDC138 | 272 | 612 | 809 | 1,17 | 0,4 | coiled-coil domain containing 138 [Source:HGNC Symbol;Acc:HGNC:26531] |
| ENSG00000149548 | CCDC15 | 83 | 139 | 213 | 0,74 | 0,62 | coiled-coil domain containing 15 [Source:HGNC Symbol;Acc:HGNC:25798] |
| ENSG00000198624 | CCDC69 | 326 | 547 | 388 | 0,75 | -0,5 | coiled-coil domain containing 69 [Source:HGNC Symbol;Acc:HGNC:24487] |
| ENSG00000091986 | CCDC80 | 1857 | 3052 | 1927 | 0,72 | -0,66 | coiled-coil domain containing 80 [Source:HGNC Symbol;Acc:HGNC:30649] |
| ENSG00000150753 | CCT5 | 4300 | 7569 | 10114 | 0,82 | 0,42 | chaperonin containing TCP1 subunit 5 [Source:HGNC Symbol;Acc:HGNC:1618] |
| ENSG00000241399 | CD302 | 329 | 1115 | 805 | 1,76 | -0,47 | CD302 molecule [Source:HGNC Symbol;Acc:HGNC:30843] |
| ENSG00000167775 | CD320 | 944 | 1590 | 2549 | 0,75 | 0,68 | CD320 molecule [Source:HGNC Symbol;Acc:HGNC:16692] |
| ENSG00000097046 | CDC7 | 457 | 844 | 1256 | 0,88 | 0,57 | cell division cycle 7 [Source:HGNC Symbol;Acc:HGNC:1745] |
| ENSG00000134371 | CDC73 | 1368 | 2192 | 2645 | 0,68 | 0,27 | cell division cycle 73 [Source:HGNC Symbol;Acc:HGNC:16783] |
| ENSG00000184661 | CDCA2 | 446 | 955 | 1347 | 1,1 | 0,49 | cell division cycle associated 2 [Source:HGNC Symbol;Acc:HGNC:14623] |
| ENSG00000111665 | CDCA3 | 514 | 843 | 1196 | 0,71 | 0,5 | cell division cycle associated 3 [Source:HGNC Symbol;Acc:HGNC:14624] |
| ENSG00000170779 | CDCA4 | 579 | 1225 | 1889 | 1,08 | 0,62 | cell division cycle associated 4 [Source:HGNC Symbol;Acc:HGNC:14625] |
| ENSG00000146670 | CDCA5 | 864 | 1823 | 2666 | 1,08 | 0,55 | cell division cycle associated 5 [Source:HGNC Symbol;Acc:HGNC:14626] |
| ENSG00000144354 | CDCA7 | 251 | 1958 | 2984 | 2,96 | 0,61 | cell division cycle associated 7 [Source:HGNC Symbol;Acc:HGNC:14628] |
| ENSG00000135446 | CDK4 | 2958 | 5500 | 7588 | 0,9 | 0,46 | cyclin dependent kinase 4 [Source:HGNC Symbol;Acc:HGNC:1773] |
| ENSG00000147889 | CDKN2A | 10 | 800 | 1199 | 6,35 | 0,58 | cyclin dependent kinase inhibitor 2A [Source:HGNC Symbol;Acc:HGNC:1787] |
| ENSG00000138778 | CENPE | 1061 | 1952 | 3065 | 0,88 | 0,65 | centromere protein E [Source:HGNC Symbol;Acc:HGNC:1856] |
| ENSG00000138092 | CENPO | 877 | 1735 | 2329 | 0,98 | 0,43 | centromere protein O [Source:HGNC Symbol;Acc:HGNC:28152] |
| ENSG00000164323 | CFAP97 | 1596 | 3086 | 2214 | 0,95 | -0,48 | cilia and flagella associated protein 97 [Source:HGNC Symbol;Acc:HGNC:29276] |
| ENSG00000149554 | CHEK1 | 740 | 1394 | 2049 | 0,91 | 0,56 | checkpoint kinase 1 [Source:HGNC Symbol;Acc:HGNC:1925] |
| ENSG00000169684 | CHRNA5 | 130 | 324 | 468 | 1,32 | 0,53 | cholinergic receptor nicotinic alpha 5 subunit [Source:HGNC Symbol;Acc:HGNC:1959] |
| ENSG00000136108 | CKAP2 | 1694 | 2741 | 3661 | 0,69 | 0,42 | cytoskeleton associated protein 2 [Source:HGNC Symbol;Acc:HGNC:1990] |
| ENSG00000164237 | CMBL | 787 | 1311 | 871 | 0,74 | -0,59 | carboxymethylenebutenolidase homolog [Source:HGNC Symbol;Acc:HGNC:25090] |
| ENSG00000100528 | CNIH1 | 2615 | 4505 | 3397 | 0,78 | -0,41 | cornichon family AMPA receptor auxiliary protein 1 [Source:HGNC Symbol;Acc:HGNC:19431] |
| ENSG00000162377 | COA7 | 324 | 731 | 1126 | 1,17 | 0,62 | cytochrome c oxidase assembly factor 7 (putative) [Source:HGNC Symbol;Acc:HGNC:25716] |
| ENSG00000141030 | COPS3 | 1029 | 1759 | 2458 | 0,77 | 0,48 | COP9 signalosome subunit 3 [Source:HGNC Symbol;Acc:HGNC:2239] |
| ENSG00000088882 | CPXM1 | 5997 | 18069 | 12556 | 1,59 | -0,53 | carboxypeptidase X, M14 family member 1 [Source:HGNC Symbol;Acc:HGNC:15771] |
| ENSG00000170275 | CRTAP | 2976 | 5518 | 3606 | 0,89 | -0,61 | cartilage associated protein [Source:HGNC Symbol;Acc:HGNC:2379] |
| ENSG00000124207 | CSE1L | 3176 | 5901 | 8477 | 0,89 | 0,52 | chromosome segregation 1 like [Source:HGNC Symbol;Acc:HGNC:2431] |
| ENSG00000172115 | CYCS | 1026 | 1731 | 2420 | 0,76 | 0,48 | cytochrome c, somatic [Source:HGNC Symbol;Acc:HGNC:19986] |
| ENSG00000006634 | DBF4 | 436 | 913 | 1204 | 1,07 | 0,4 | DBF4 zinc finger [Source:HGNC Symbol;Acc:HGNC:17364] |
| ENSG00000198924 | DCLRE1A | 364 | 588 | 945 | 0,69 | 0,68 | DNA cross-link repair 1A [Source:HGNC Symbol;Acc:HGNC:17660] |
| ENSG00000118655 | DCLRE1B | 259 | 475 | 710 | 0,88 | 0,58 | DNA cross-link repair 1B [Source:HGNC Symbol;Acc:HGNC:17641] |
| ENSG00000165490 | DDIAS | 150 | 353 | 554 | 1,24 | 0,65 | DNA damage induced apoptosis suppressor [Source:HGNC Symbol;Acc:HGNC:26351] |
| ENSG00000138346 | DNA2 | 536 | 1341 | 2055 | 1,32 | 0,62 | DNA replication helicase/nuclease 2 [Source:HGNC Symbol;Acc:HGNC:2939] |
| ENSG00000176410 | DNAJC30 | 111 | 186 | 292 | 0,75 | 0,65 | DnaJ heat shock protein family (Hsp40) member C30 [Source:HGNC Symbol;Acc:HGNC:16410] |
| ENSG00000136048 | DRAM1 | 592 | 1473 | 1859 | 1,31 | 0,34 | DNA damage regulated autophagy modulator 1 [Source:HGNC Symbol;Acc:HGNC:25645] |
| ENSG00000139318 | DUSP6 | 3155 | 18501 | 13796 | 2,55 | -0,42 | dual specificity phosphatase 6 [Source:HGNC Symbol;Acc:HGNC:3072] |
| ENSG00000128951 | DUT | 573 | 924 | 1278 | 0,69 | 0,47 | deoxyuridine triphosphatase [Source:HGNC Symbol;Acc:HGNC:3078] |
| ENSG00000134874 | DZIP1 | 1216 | 2361 | 1806 | 0,96 | -0,39 | DAZ interacting zinc finger protein 1 [Source:HGNC Symbol;Acc:HGNC:20908] |
| ENSG00000233057 | EEF1A1P14 | 102 | 170 | 114 | 0,74 | -0,58 | eukaryotic translation elongation factor 1 alpha 1 pseudogene 14 [Source:HGNC Symbol;Acc:HGNC:3197] |
| ENSG00000163577 | EIF5A2 | 78 | 160 | 104 | 1,04 | -0,63 | eukaryotic translation initiation factor 5A2 [Source:HGNC Symbol;Acc:HGNC:3301] |
| ENSG00000170522 | ELOVL6 | 2316 | 4018 | 5989 | 0,79 | 0,58 | ELOVL fatty acid elongase 6 [Source:HGNC Symbol;Acc:HGNC:15829] |
| ENSG00000074800 | ENO1 | 51071 | 88820 | 59484 | 0,8 | -0,58 | enolase 1 [Source:HGNC Symbol;Acc:HGNC:3350] |
| ENSG00000135476 | ESPL1 | 988 | 1752 | 2580 | 0,83 | 0,56 | extra spindle pole bodies like 1, separase [Source:HGNC Symbol;Acc:HGNC:16856] |
| ENSG00000123737 | EXOSC9 | 661 | 1104 | 1396 | 0,74 | 0,34 | exosome component 9 [Source:HGNC Symbol;Acc:HGNC:9137] |
| ENSG00000149485 | FADS1 | 5755 | 9980 | 13746 | 0,79 | 0,46 | fatty acid desaturase 1 [Source:HGNC Symbol;Acc:HGNC:3574] |
| ENSG00000144554 | FANCD2 | 1062 | 1745 | 2371 | 0,72 | 0,44 | Fanconi anemia complementation group D2 [Source:HGNC Symbol;Acc:HGNC:3585] |
| ENSG00000145743 | FBXL17 | 654 | 1044 | 818 | 0,68 | -0,35 | F-box and leucine rich repeat protein 17 [Source:HGNC Symbol;Acc:HGNC:13615] |
| ENSG00000164946 | FREM1 | 1724 | 21215 | 13035 | 3,62 | -0,7 | FRAS1 related extracellular matrix 1 [Source:HGNC Symbol;Acc:HGNC:23399] |
| ENSG00000255883 | FUNDC2P1 | 194 | 330 | 217 | 0,77 | -0,61 | FUN14 domain containing 2 pseudogene 1 [Source:HGNC Symbol;Acc:HGNC:17253] |
| ENSG00000165060 | FXN | 303 | 515 | 647 | 0,76 | 0,33 | frataxin [Source:HGNC Symbol;Acc:HGNC:3951] |
| ENSG00000228232 | GAPDHP1 | 127 | 230 | 151 | 0,86 | -0,6 | glyceraldehyde-3-phosphate dehydrogenase pseudogene 1 [Source:HGNC Symbol;Acc:HGNC:4159] |
| ENSG00000006007 | GDE1 | 4395 | 8852 | 5526 | 1,01 | -0,68 | glycerophosphodiester phosphodiesterase 1 [Source:HGNC Symbol;Acc:HGNC:29644] |
| ENSG00000179409 | GEMIN4 | 812 | 1322 | 1938 | 0,7 | 0,55 | gem nuclear organelle associated protein 4 [Source:HGNC Symbol;Acc:HGNC:15717] |
| ENSG00000023572 | GLRX2 | 83 | 162 | 256 | 0,97 | 0,65 | glutaredoxin 2 [Source:HGNC Symbol;Acc:HGNC:16065] |
| ENSG00000172380 | GNG12 | 1305 | 2232 | 1423 | 0,77 | -0,65 | G protein subunit gamma 12 [Source:HGNC Symbol;Acc:HGNC:19663] |
| ENSG00000158292 | GPR153 | 605 | 1266 | 778 | 1,07 | -0,7 | G protein-coupled receptor 153 [Source:HGNC Symbol;Acc:HGNC:23618] |
| ENSG00000122034 | GTF3A | 1229 | 1999 | 2682 | 0,7 | 0,42 | general transcription factor IIIA [Source:HGNC Symbol;Acc:HGNC:4662] |
| ENSG00000155115 | GTF3C6 | 598 | 1104 | 1494 | 0,89 | 0,44 | general transcription factor IIIC subunit 6 [Source:HGNC Symbol;Acc:HGNC:20872] |
| ENSG00000075218 | GTSE1 | 776 | 1240 | 1737 | 0,68 | 0,49 | G2 and S-phase expressed 1 [Source:HGNC Symbol;Acc:HGNC:13698] |
| ENSG00000105968 | H2AFV | 2907 | 5316 | 7426 | 0,87 | 0,48 | H2A histone family member V [Source:HGNC Symbol;Acc:HGNC:20664] |
| ENSG00000128708 | HAT1 | 588 | 1030 | 1407 | 0,81 | 0,45 | histone acetyltransferase 1 [Source:HGNC Symbol;Acc:HGNC:4821] |
| ENSG00000167220 | HDHD2 | 948 | 1612 | 1286 | 0,77 | -0,33 | haloacid dehalogenase like hydrolase domain containing 2 [Source:HGNC Symbol;Acc:HGNC:25364] |
| ENSG00000110422 | HIPK3 | 1656 | 3130 | 3785 | 0,92 | 0,27 | homeodomain interacting protein kinase 3 [Source:HGNC Symbol;Acc:HGNC:4915] |
| ENSG00000149929 | HIRIP3 | 303 | 671 | 974 | 1,15 | 0,54 | HIRA interacting protein 3 [Source:HGNC Symbol;Acc:HGNC:4917] |
| ENSG00000071794 | HLTF | 2695 | 4332 | 2838 | 0,68 | -0,61 | helicase like transcription factor [Source:HGNC Symbol;Acc:HGNC:11099] |
| ENSG00000189403 | HMGB1 | 4063 | 8236 | 11783 | 1,02 | 0,52 | high mobility group box 1 [Source:HGNC Symbol;Acc:HGNC:4983] |
| ENSG00000132967 | HMGB1P5 | 3780 | 6797 | 9857 | 0,85 | 0,54 | high mobility group box 1 pseudogene 5 [Source:HGNC Symbol;Acc:HGNC:4997] |
| ENSG00000113161 | HMGCR | 5552 | 10124 | 15674 | 0,87 | 0,63 | 3-hydroxy-3-methylglutaryl-CoA reductase [Source:HGNC Symbol;Acc:HGNC:5006] |
| ENSG00000138668 | HNRNPD | 7763 | 13117 | 18588 | 0,76 | 0,5 | heterogeneous nuclear ribonucleoprotein D [Source:HGNC Symbol;Acc:HGNC:5036] |
| ENSG00000153976 | HS3ST3A1 | 173 | 1545 | 952 | 3,15 | -0,7 | heparan sulfate-glucosamine 3-sulfotransferase 3A1 [Source:HGNC Symbol;Acc:HGNC:5196] |
| ENSG00000155304 | HSPA13 | 2300 | 4308 | 3414 | 0,91 | -0,34 | heat shock protein family A (Hsp70) member 13 [Source:HGNC Symbol;Acc:HGNC:11375] |
| ENSG00000113013 | HSPA9 | 5068 | 8853 | 11751 | 0,8 | 0,41 | heat shock protein family A (Hsp70) member 9 [Source:HGNC Symbol;Acc:HGNC:5244] |
| ENSG00000226666 | HSPA9P1 | 117 | 206 | 293 | 0,82 | 0,51 | heat shock protein family A (Hsp70) member 9 pseudogene 1 [Source:HGNC Symbol;Acc:HGNC:24915] |
| ENSG00000076662 | ICAM3 | 51 | 108 | 165 | 1,08 | 0,61 | intercellular adhesion molecule 3 [Source:HGNC Symbol;Acc:HGNC:5346] |
| ENSG00000185201 | IFITM2 | 2433 | 4074 | 2480 | 0,74 | -0,72 | interferon induced transmembrane protein 2 [Source:HGNC Symbol;Acc:HGNC:5413] |
| ENSG00000103742 | IGDCC4 | 1231 | 2062 | 1510 | 0,74 | -0,45 | immunoglobulin superfamily DCC subclass member 4 [Source:HGNC Symbol;Acc:HGNC:13770] |
| ENSG00000144730 | IL17RD | 1401 | 2324 | 1825 | 0,73 | -0,35 | interleukin 17 receptor D [Source:HGNC Symbol;Acc:HGNC:17616] |
| ENSG00000186480 | INSIG1 | 563 | 932 | 1443 | 0,73 | 0,63 | insulin induced gene 1 [Source:HGNC Symbol;Acc:HGNC:6083] |
| ENSG00000151151 | IPMK | 885 | 1473 | 948 | 0,74 | -0,64 | inositol polyphosphate multikinase [Source:HGNC Symbol;Acc:HGNC:20739] |
| ENSG00000178202 | KDELC2 | 987 | 2255 | 2780 | 1,19 | 0,3 | KDEL motif containing 2 [Source:HGNC Symbol;Acc:HGNC:28496] |
| ENSG00000138160 | KIF11 | 1587 | 3078 | 4367 | 0,96 | 0,5 | kinesin family member 11 [Source:HGNC Symbol;Acc:HGNC:6388] |
| ENSG00000186185 | KIF18B | 713 | 1189 | 1883 | 0,74 | 0,66 | kinesin family member 18B [Source:HGNC Symbol;Acc:HGNC:27102] |
| ENSG00000182481 | KPNA2 | 4403 | 7912 | 10266 | 0,85 | 0,38 | karyopherin subunit alpha 2 [Source:HGNC Symbol;Acc:HGNC:6395] |
| ENSG00000106003 | LFNG | 672 | 1162 | 1818 | 0,79 | 0,65 | LFNG O-fucosylpeptide 3-beta-N-acetylglucosaminyltransferase [Source:HGNC Symbol;Acc:HGNC:6560] |
| ENSG00000105486 | LIG1 | 1444 | 2348 | 3179 | 0,7 | 0,44 | DNA ligase 1 [Source:HGNC Symbol;Acc:HGNC:6598] |
| ENSG00000169756 | LIMS1 | 2332 | 4261 | 3413 | 0,87 | -0,32 | LIM zinc finger domain containing 1 [Source:HGNC Symbol;Acc:HGNC:6616] |
| ENSG00000165501 | LRR1 | 250 | 534 | 669 | 1,09 | 0,33 | leucine rich repeat protein 1 [Source:HGNC Symbol;Acc:HGNC:19742] |
| ENSG00000175061 | LRRC75A-AS1 | 12644 | 20467 | 14733 | 0,69 | -0,47 | LRRC75A antisense RNA 1 [Source:HGNC Symbol;Acc:HGNC:28619] |
| ENSG00000133739 | LRRCC1 | 241 | 402 | 571 | 0,74 | 0,5 | leucine rich repeat and coiled-coil centrosomal protein 1 [Source:HGNC Symbol;Acc:HGNC:29373] |
| ENSG00000101367 | MAPRE1 | 2697 | 4407 | 5681 | 0,71 | 0,37 | microtubule associated protein RP/EB family member 1 [Source:HGNC Symbol;Acc:HGNC:6890] |
| ENSG00000247626 | MARS2 | 218 | 355 | 517 | 0,7 | 0,54 | methionyl-tRNA synthetase 2, mitochondrial [Source:HGNC Symbol;Acc:HGNC:25133] |
| ENSG00000131844 | MCCC2 | 1424 | 2301 | 2942 | 0,69 | 0,35 | methylcrotonoyl-CoA carboxylase 2 [Source:HGNC Symbol;Acc:HGNC:6937] |
| ENSG00000112118 | MCM3 | 3714 | 6396 | 9840 | 0,78 | 0,62 | minichromosome maintenance complex component 3 [Source:HGNC Symbol;Acc:HGNC:6945] |
| ENSG00000134138 | MEIS2 | 2776 | 7447 | 9490 | 1,42 | 0,35 | Meis homeobox 2 [Source:HGNC Symbol;Acc:HGNC:7001] |
| ENSG00000168282 | MGAT2 | 763 | 1302 | 1720 | 0,77 | 0,4 | mannosyl (alpha-1,6-)-glycoprotein beta-1,2-N-acetylglucosaminyltransferase [Source:HGNC Symbol;Acc:HGNC:7045] |
| ENSG00000148773 | MKI67 | 3675 | 5927 | 9508 | 0,69 | 0,68 | marker of proliferation Ki-67 [Source:HGNC Symbol;Acc:HGNC:7107] |
| ENSG00000132763 | MMACHC | 196 | 343 | 534 | 0,81 | 0,64 | methylmalonic aciduria (cobalamin deficiency) cblC type, with homocystinuria [Source:HGNC Symbol;Acc:HGNC:24525] |
| ENSG00000156103 | MMP16 | 834 | 12123 | 7877 | 3,86 | -0,62 | matrix metallopeptidase 16 [Source:HGNC Symbol;Acc:HGNC:7162] |
| ENSG00000051825 | MPHOSPH9 | 660 | 1059 | 1485 | 0,68 | 0,49 | M-phase phosphoprotein 9 [Source:HGNC Symbol;Acc:HGNC:7215] |
| ENSG00000066382 | MPPED2 | 73 | 310 | 212 | 2,09 | -0,54 | metallophosphoesterase domain containing 2 [Source:HGNC Symbol;Acc:HGNC:1180] |
| ENSG00000137547 | MRPL15 | 491 | 815 | 1127 | 0,73 | 0,47 | mitochondrial ribosomal protein L15 [Source:HGNC Symbol;Acc:HGNC:14054] |
| ENSG00000053372 | MRTO4 | 649 | 1103 | 1665 | 0,77 | 0,59 | MRT4 homolog, ribosome maturation factor [Source:HGNC Symbol;Acc:HGNC:18477] |
| ENSG00000095002 | MSH2 | 1400 | 2604 | 3686 | 0,9 | 0,5 | mutS homolog 2 [Source:HGNC Symbol;Acc:HGNC:7325] |
| ENSG00000132780 | NASP | 6263 | 10858 | 17115 | 0,79 | 0,66 | nuclear autoantigenic sperm protein [Source:HGNC Symbol;Acc:HGNC:7644] |
| ENSG00000121152 | NCAPH | 486 | 1184 | 1871 | 1,29 | 0,66 | non-SMC condensin I complex subunit H [Source:HGNC Symbol;Acc:HGNC:1112] |
| ENSG00000266412 | NCOA4 | 4952 | 8178 | 6812 | 0,72 | -0,26 | nuclear receptor coactivator 4 [Source:HGNC Symbol;Acc:HGNC:7671] |
| ENSG00000123545 | NDUFAF4 | 179 | 286 | 374 | 0,68 | 0,38 | NADH:ubiquinone oxidoreductase complex assembly factor 4 [Source:HGNC Symbol;Acc:HGNC:21034] |
| ENSG00000189362 | NEMP2 | 223 | 460 | 605 | 1,05 | 0,4 | nuclear envelope integral membrane protein 2 [Source:HGNC Symbol;Acc:HGNC:33700] |
| ENSG00000072736 | NFATC3 | 981 | 1586 | 2114 | 0,69 | 0,41 | nuclear factor of activated T-cells 3 [Source:HGNC Symbol;Acc:HGNC:7777] |
| ENSG00000130935 | NOL11 | 1091 | 1767 | 2259 | 0,7 | 0,35 | nucleolar protein 11 [Source:HGNC Symbol;Acc:HGNC:24557] |
| ENSG00000148200 | NR6A1 | 3180 | 5992 | 7711 | 0,91 | 0,36 | nuclear receptor subfamily 6 group A member 1 [Source:HGNC Symbol;Acc:HGNC:7985] |
| ENSG00000113569 | NUP155 | 1190 | 2040 | 2753 | 0,78 | 0,43 | nucleoporin 155 [Source:HGNC Symbol;Acc:HGNC:8063] |
| ENSG00000091651 | ORC6 | 615 | 1084 | 1608 | 0,82 | 0,57 | origin recognition complex subunit 6 [Source:HGNC Symbol;Acc:HGNC:17151] |

| | | | | | | | |
|---|---|---|---|---|---|---|---|
| ENSG00000083720 | OXCT1 | 251 | 1082 | 1498 | 2,11 | 0,47 | 3-oxoacid CoA-transferase 1 [Source:HGNC Symbol;Acc:HGNC:8527] |
| ENSG00000111845 | PAK1IP1 | 210 | 443 | 650 | 1,08 | 0,55 | PAK1 interacting protein 1 [Source:HGNC Symbol;Acc:HGNC:20882] |
| ENSG00000173599 | PC | 511 | 1177 | 1773 | 1,2 | 0,59 | pyruvate carboxylase [Source:HGNC Symbol;Acc:HGNC:8636] |
| ENSG00000142657 | PGD | 4224 | 6869 | 9258 | 0,7 | 0,43 | phosphogluconate dehydrogenase [Source:HGNC Symbol;Acc:HGNC:8891] |
| ENSG00000087842 | PIR | 58 | 229 | 358 | 1,99 | 0,64 | pirin [Source:HGNC Symbol;Acc:HGNC:30048] |
| ENSG00000171033 | PKIA | 364 | 847 | 1323 | 1,22 | 0,64 | protein kinase (cAMP-dependent, catalytic) inhibitor alpha [Source:HGNC Symbol;Acc:HGNC:9017] |
| ENSG00000114805 | PLCH1 | 451 | 929 | 1245 | 1,04 | 0,42 | phospholipase C eta 1 [Source:HGNC Symbol;Acc:HGNC:29185] |
| ENSG00000143850 | PLEKHA6 | 2702 | 4956 | 7148 | 0,88 | 0,53 | pleckstrin homology domain containing A6 [Source:HGNC Symbol;Acc:HGNC:17053] |
| ENSG00000100979 | PLTP | 1154 | 2019 | 1446 | 0,81 | -0,48 | phospholipid transfer protein [Source:HGNC Symbol;Acc:HGNC:9093] |
| ENSG00000004399 | PLXND1 | 4656 | 11516 | 7960 | 1,31 | -0,53 | plexin D1 [Source:HGNC Symbol;Acc:HGNC:9107] |
| ENSG00000115946 | PNO1 | 261 | 429 | 635 | 0,72 | 0,57 | partner of NOB1 homolog [Source:HGNC Symbol;Acc:HGNC:32790] |
| ENSG00000148229 | POLE3 | 1040 | 1991 | 2805 | 0,94 | 0,49 | DNA polymerase epsilon 3, accessory subunit [Source:HGNC Symbol;Acc:HGNC:13546] |
| ENSG00000051341 | POLQ | 410 | 685 | 946 | 0,74 | 0,46 | DNA polymerase theta [Source:HGNC Symbol;Acc:HGNC:9186] |
| ENSG00000196262 | PPIA | 8273 | 13638 | 18554 | 0,72 | 0,44 | peptidylprolyl isomerase A [Source:HGNC Symbol;Acc:HGNC:9253] |
| ENSG00000236334 | PPIAL4G | 583 | 964 | 1346 | 0,73 | 0,48 | peptidylprolyl isomerase A like 4G [Source:HGNC Symbol;Acc:HGNC:33996] |
| ENSG00000251495 | PPIAP11 | 913 | 1461 | 2056 | 0,68 | 0,49 | peptidylprolyl isomerase A pseudogene 11 [Source:HGNC Symbol;Acc:HGNC:9263] |
| ENSG00000137168 | PPIL1 | 619 | 1076 | 1689 | 0,8 | 0,65 | peptidylprolyl isomerase like 1 [Source:HGNC Symbol;Acc:HGNC:9260] |
| ENSG00000186298 | PPP1CC | 6001 | 10662 | 7577 | 0,83 | -0,49 | protein phosphatase 1 catalytic subunit gamma [Source:HGNC Symbol;Acc:HGNC:9283] |
| ENSG00000131238 | PPT1 | 2188 | 3754 | 4992 | 0,78 | 0,41 | palmitoyl-protein thioesterase 1 [Source:HGNC Symbol;Acc:HGNC:9325] |
| ENSG00000198056 | PRIM1 | 239 | 531 | 824 | 1,15 | 0,63 | primase (DNA) subunit 1 [Source:HGNC Symbol;Acc:HGNC:9369] |
| ENSG00000100033 | PRODH | 282 | 556 | 399 | 0,98 | -0,48 | proline dehydrogenase 1 [Source:HGNC Symbol;Acc:HGNC:9453] |
| ENSG00000099256 | PRTFDC1 | 222 | 366 | 557 | 0,72 | 0,61 | phosphoribosyl transferase domain containing 1 [Source:HGNC Symbol;Acc:HGNC:23333] |
| ENSG00000131470 | PSMC3IP | 218 | 418 | 632 | 0,94 | 0,6 | PSMC3 interacting protein [Source:HGNC Symbol;Acc:HGNC:17928] |
| ENSG00000183527 | PSMG1 | 378 | 686 | 988 | 0,86 | 0,53 | proteasome assembly chaperone 1 [Source:HGNC Symbol;Acc:HGNC:3043] |
| ENSG00000184489 | PTP4A3 | 1594 | 2849 | 1806 | 0,84 | -0,66 | protein tyrosine phosphatase type IVA, member 3 [Source:HGNC Symbol;Acc:HGNC:9636] |
| ENSG00000171016 | PYGO1 | 337 | 663 | 528 | 0,98 | -0,33 | pygopus family PHD finger 1 [Source:HGNC Symbol;Acc:HGNC:30256] |
| ENSG00000213339 | QTRT1 | 485 | 932 | 1289 | 0,94 | 0,47 | queuine tRNA-ribosyltransferase catalytic subunit 1 [Source:HGNC Symbol;Acc:HGNC:23797] |
| ENSG00000137502 | RAB30 | 369 | 664 | 498 | 0,85 | -0,41 | RAB30, member RAS oncogene family [Source:HGNC Symbol;Acc:HGNC:9770] |
| ENSG00000127314 | RAP1B | 629 | 1643 | 1193 | 1,39 | -0,46 | RAP1B, member of RAS oncogene family [Source:HGNC Symbol;Acc:HGNC:9857] |
| ENSG00000244462 | RBM12 | 1882 | 3515 | 5555 | 0,9 | 0,66 | RNA binding motif protein 12 [Source:HGNC Symbol;Acc:HGNC:9898] |
| ENSG00000102317 | RBM3 | 2422 | 4084 | 6333 | 0,75 | 0,63 | RNA binding motif (RNP1, RRM) protein 3 [Source:HGNC Symbol;Acc:HGNC:9900] |
| ENSG00000133119 | RFC3 | 343 | 907 | 1452 | 1,41 | 0,68 | replication factor C subunit 3 [Source:HGNC Symbol;Acc:HGNC:9971] |
| ENSG00000163918 | RFC4 | 650 | 1097 | 1565 | 0,76 | 0,51 | replication factor C subunit 4 [Source:HGNC Symbol;Acc:HGNC:9972] |
| ENSG00000133111 | RFXAP | 150 | 318 | 506 | 1,09 | 0,67 | regulatory factor X associated protein [Source:HGNC Symbol;Acc:HGNC:9988] |
| ENSG00000187994 | RINL | 146 | 285 | 175 | 0,96 | -0,71 | Ras and Rab interactor like [Source:HGNC Symbol;Acc:HGNC:24795] |
| ENSG00000178966 | RMI1 | 368 | 733 | 1119 | 0,99 | 0,61 | RecQ mediated genome instability 1 [Source:HGNC Symbol;Acc:HGNC:25764] |
| ENSG00000104889 | RNASEH2A | 575 | 1217 | 1778 | 1,08 | 0,55 | ribonuclease H2 subunit A [Source:HGNC Symbol;Acc:HGNC:18518] |
| ENSG00000101695 | RNF125 | 170 | 371 | 233 | 1,13 | -0,67 | ring finger protein 125 [Source:HGNC Symbol;Acc:HGNC:21150] |
| ENSG00000117748 | RPA2 | 607 | 1239 | 1652 | 1,03 | 0,41 | replication protein A2 [Source:HGNC Symbol;Acc:HGNC:10290] |
| ENSG00000233913 | RPL10P9 | 7214 | 12609 | 8529 | 0,81 | -0,56 | ribosomal protein L10 pseudogene 9 [Source:HGNC Symbol;Acc:HGNC:35579] |
| ENSG00000167325 | RRM1 | 2136 | 4283 | 5748 | 1 | 0,42 | ribonucleotide reductase catalytic subunit M1 [Source:HGNC Symbol;Acc:HGNC:10451] |
| ENSG00000067533 | RRP15 | 422 | 1008 | 1349 | 1,25 | 0,42 | ribosomal RNA processing 15 homolog [Source:HGNC Symbol;Acc:HGNC:24255] |
| ENSG00000182010 | RTKN2 | 400 | 1152 | 1605 | 1,53 | 0,48 | rhotekin 2 [Source:HGNC Symbol;Acc:HGNC:19364] |
| ENSG00000168061 | SAC3D1 | 304 | 553 | 830 | 0,87 | 0,59 | SAC3 domain containing 1 [Source:HGNC Symbol;Acc:HGNC:30179] |
| ENSG00000164105 | SAP30 | 241 | 431 | 613 | 0,84 | 0,51 | Sin3A associated protein 30 [Source:HGNC Symbol;Acc:HGNC:10532] |
| ENSG00000100934 | SEC23A | 4708 | 7973 | 6071 | 0,76 | -0,39 | Sec23 homolog A, coat complex II component [Source:HGNC Symbol;Acc:HGNC:10701] |
| ENSG00000086475 | SEPHS1 | 2519 | 4125 | 5497 | 0,71 | 0,41 | selenophosphate synthetase 1 [Source:HGNC Symbol;Acc:HGNC:19685] |
| ENSG00000171241 | SHCBP1 | 237 | 647 | 878 | 1,45 | 0,44 | SHC binding and spindle associated 1 [Source:HGNC Symbol;Acc:HGNC:29547] |
| ENSG00000180730 | SHISA2 | 482 | 10645 | 15014 | 4,46 | 0,5 | shisa family member 2 [Source:HGNC Symbol;Acc:HGNC:20366] |
| ENSG00000154839 | SKA1 | 274 | 738 | 1133 | 1,43 | 0,62 | spindle and kinetochore associated complex subunit 1 [Source:HGNC Symbol;Acc:HGNC:28109] |
| ENSG00000182628 | SKA2 | 828 | 2031 | 2730 | 1,3 | 0,43 | spindle and kinetochore associated complex subunit 2 [Source:HGNC Symbol;Acc:HGNC:28006] |
| ENSG00000145604 | SKP2 | 970 | 1821 | 2797 | 0,91 | 0,62 | S-phase kinase associated protein 2 [Source:HGNC Symbol;Acc:HGNC:10901] |
| ENSG00000125520 | SLC2A4RG | 973 | 1715 | 2229 | 0,82 | 0,38 | SLC2A4 regulator [Source:HGNC Symbol;Acc:HGNC:15930] |
| ENSG00000108055 | SMC3 | 9874 | 17600 | 27340 | 0,83 | 0,64 | structural maintenance of chromosomes 3 [Source:HGNC Symbol;Acc:HGNC:2468] |
| ENSG00000184602 | SNN | 2163 | 3590 | 5399 | 0,73 | 0,59 | stannin [Source:HGNC Symbol;Acc:HGNC:11149] |
| ENSG00000167088 | SNRPD1 | 959 | 1908 | 2925 | 0,99 | 0,62 | small nuclear ribonucleoprotein D1 polypeptide [Source:HGNC Symbol;Acc:HGNC:11158] |
| ENSG00000110693 | SOX6 | 1270 | 2314 | 3398 | 0,87 | 0,55 | SRY-box 6 [Source:HGNC Symbol;Acc:HGNC:16421] |
| ENSG00000136158 | SPRY2 | 6294 | 15737 | 20053 | 1,32 | 0,35 | sprouty RTK signaling antagonist 2 [Source:HGNC Symbol;Acc:HGNC:11270] |
| ENSG00000145545 | SRD5A1 | 582 | 933 | 1208 | 0,68 | 0,37 | steroid 5 alpha-reductase 1 [Source:HGNC Symbol;Acc:HGNC:11284] |
| ENSG00000116649 | SRM | 601 | 1136 | 1788 | 0,92 | 0,66 | spermidine synthase [Source:HGNC Symbol;Acc:HGNC:11296] |
| ENSG00000101955 | SRPX | 198 | 505 | 338 | 1,35 | -0,58 | sushi repeat containing protein, X-linked [Source:HGNC Symbol;Acc:HGNC:11309] |
| ENSG00000004866 | ST7 | 238 | 412 | 585 | 0,79 | 0,51 | suppression of tumorigenicity 7 [Source:HGNC Symbol;Acc:HGNC:11351] |
| ENSG00000117632 | STMN1 | 7986 | 18433 | 27180 | 1,21 | 0,56 | stathmin 1 [Source:HGNC Symbol;Acc:HGNC:6510] |
| ENSG00000152455 | SUV39H2 | 427 | 928 | 1259 | 1,12 | 0,44 | suppressor of variegation 3-9 homolog 2 [Source:HGNC Symbol;Acc:HGNC:17287] |
| ENSG00000198168 | SVIP | 921 | 2654 | 4199 | 1,53 | 0,66 | small VCP interacting protein [Source:HGNC Symbol;Acc:HGNC:25238] |
| ENSG00000013810 | TACC3 | 1705 | 2934 | 4255 | 0,78 | 0,54 | transforming acidic coiled-coil containing protein 3 [Source:HGNC Symbol;Acc:HGNC:11524] |
| ENSG00000137492 | THAP12 | 883 | 1629 | 1984 | 0,88 | 0,28 | THAP domain containing 12 [Source:HGNC Symbol;Acc:HGNC:9440] |
| ENSG00000159445 | THEM4 | 231 | 542 | 709 | 1,23 | 0,39 | thioesterase superfamily member 4 [Source:HGNC Symbol;Acc:HGNC:17947] |
| ENSG00000111602 | TIMELESS | 2524 | 4410 | 6688 | 0,81 | 0,6 | timeless circadian clock [Source:HGNC Symbol;Acc:HGNC:11813] |
| ENSG00000100575 | TIMM9 | 478 | 773 | 564 | 0,69 | -0,46 | translocase of inner mitochondrial membrane 9 [Source:HGNC Symbol;Acc:HGNC:11819] |
| ENSG00000140332 | TLE3 | 1559 | 3756 | 5336 | 1,27 | 0,51 | transducin like enhancer of split 3 [Source:HGNC Symbol;Acc:HGNC:11839] |
| ENSG00000137210 | TMEM14B | 397 | 636 | 894 | 0,68 | 0,49 | transmembrane protein 14B [Source:HGNC Symbol;Acc:HGNC:21384] |
| ENSG00000174695 | TMEM167A | 6411 | 11531 | 14392 | 0,85 | 0,32 | transmembrane protein 167A [Source:HGNC Symbol;Acc:HGNC:28330] |
| ENSG00000146802 | TMEM168 | 485 | 778 | 491 | 0,68 | -0,66 | transmembrane protein 168 [Source:HGNC Symbol;Acc:HGNC:25826] |
| ENSG00000155099 | TMEM55A | 826 | 1363 | 943 | 0,72 | -0,53 | transmembrane protein 55A [Source:HGNC Symbol;Acc:HGNC:25452] |
| ENSG00000109084 | TMEM97 | 1463 | 3500 | 5159 | 1,26 | 0,56 | transmembrane protein 97 [Source:HGNC Symbol;Acc:HGNC:28106] |
| ENSG00000071539 | TRIP13 | 232 | 793 | 1072 | 1,78 | 0,44 | thyroid hormone receptor interactor 13 [Source:HGNC Symbol;Acc:HGNC:12307] |
| ENSG00000174173 | TRMT10C | 429 | 696 | 896 | 0,7 | 0,37 | tRNA methyltransferase 10C, mitochondrial RNase P subunit [Source:HGNC Symbol;Acc:HGNC:26022] |
| ENSG00000198860 | TSEN15 | 283 | 576 | 807 | 1,03 | 0,49 | tRNA splicing endonuclease subunit 15 [Source:HGNC Symbol;Acc:HGNC:16791] |
| ENSG00000113638 | TTC33 | 823 | 1325 | 994 | 0,69 | -0,42 | tetratricopeptide repeat domain 33 [Source:HGNC Symbol;Acc:HGNC:29959] |
| ENSG00000198431 | TXNRD1 | 2656 | 4626 | 7171 | 0,8 | 0,63 | thioredoxin reductase 1 [Source:HGNC Symbol;Acc:HGNC:12437] |
| ENSG00000176890 | TYMS | 1578 | 4131 | 6073 | 1,39 | 0,56 | thymidylate synthetase [Source:HGNC Symbol;Acc:HGNC:12441] |
| ENSG00000177889 | UBE2N | 1625 | 2698 | 3699 | 0,73 | 0,46 | ubiquitin conjugating enzyme E2 N [Source:HGNC Symbol;Acc:HGNC:12492] |
| ENSG00000012963 | UBR7 | 566 | 952 | 1367 | 0,75 | 0,52 | ubiquitin protein ligase E3 component n-recognin 7 (putative) [Source:HGNC Symbol;Acc:HGNC:20344] |
| ENSG00000058056 | USP13 | 449 | 937 | 1280 | 1,06 | 0,45 | ubiquitin specific peptidase 13 (isopeptidase T-3) [Source:HGNC Symbol;Acc:HGNC:12611] |
| ENSG00000148429 | USP6NL | 1598 | 2738 | 3593 | 0,78 | 0,39 | USP6 N-terminal like [Source:HGNC Symbol;Acc:HGNC:16858] |
| ENSG00000141076 | UTP4 | 897 | 1564 | 2067 | 0,8 | 0,4 | UTP4, small subunit processome component [Source:HGNC Symbol;Acc:HGNC:1983] |
| ENSG00000143494 | VASH2 | 189 | 433 | 557 | 1,2 | 0,36 | vasohibin 2 [Source:HGNC Symbol;Acc:HGNC:25723] |
| ENSG00000026025 | VIM | 27569 | 60612 | 41608 | 1,14 | -0,54 | vimentin [Source:HGNC Symbol;Acc:HGNC:12692] |
| ENSG00000092470 | WDR76 | 164 | 432 | 672 | 1,4 | 0,64 | WD repeat domain 76 [Source:HGNC Symbol;Acc:HGNC:25773] |
| ENSG00000126562 | WNK4 | 604 | 1057 | 672 | 0,81 | -0,65 | WNK lysine deficient protein kinase 4 [Source:HGNC Symbol;Acc:HGNC:14544] |
| ENSG00000128245 | YWHAH | 1878 | 3054 | 4348 | 0,7 | 0,51 | tyrosine 3-monooxygenase/tryptophan 5-monooxygenase activation protein eta [Source:HGNC Symbol;Acc:HGNC:12853] |
| ENSG00000105939 | ZC3HAV1 | 1049 | 2057 | 2877 | 0,97 | 0,48 | zinc finger CCCH-type containing, antiviral 1 [Source:HGNC Symbol;Acc:HGNC:23721] |
| ENSG00000196247 | ZNF107 | 443 | 853 | 1091 | 0,95 | 0,36 | zinc finger protein 107 [Source:HGNC Symbol;Acc:HGNC:12887] |
| ENSG00000156853 | ZNF689 | 1063 | 1748 | 2099 | 0,72 | 0,26 | zinc finger protein 689 [Source:HGNC Symbol;Acc:HGNC:25173] |
| ENSG00000196081 | ZNF724 | 93 | 182 | 265 | 0,97 | 0,54 | zinc finger protein 724 [Source:HGNC Symbol;Acc:HGNC:32460] |
| ENSG00000185869 | ZNF829 | 75 | 135 | 201 | 0,84 | 0,58 | zinc finger protein 829 [Source:HGNC Symbol;Acc:HGNC:34032] |

**GROUP 3**

| | | | | | | | |
|---|---|---|---|---|---|---|---|
| ENSG00000167107 | ACSF2 | 2858 | 1579 | 1108 | -0,86 | -0,51 | acyl-CoA synthetase family member 2 [Source:HGNC Symbol;Acc:HGNC:26101] |
| ENSG00000151651 | ADAM8 | 1942 | 260 | 159 | -2,9 | -0,71 | ADAM metallopeptidase domain 8 [Source:HGNC Symbol;Acc:HGNC:215] |
| ENSG00000121281 | ADCY7 | 1394 | 667 | 405 | -1,06 | -0,72 | adenylate cyclase 7 [Source:HGNC Symbol;Acc:HGNC:238] |
| ENSG00000072071 | ADGRL1 | 5369 | 2650 | 1989 | -1,02 | -0,41 | adhesion G protein-coupled receptor L1 [Source:HGNC Symbol;Acc:HGNC:20973] |
| ENSG00000136383 | ALPK3 | 11723 | 4914 | 5832 | -1,25 | 0,25 | alpha kinase 3 [Source:HGNC Symbol;Acc:HGNC:17574] |
| ENSG00000139211 | AMIGO2 | 482 | 133 | 85 | -1,86 | -0,65 | adhesion molecule with Ig like domain 2 [Source:HGNC Symbol;Acc:HGNC:24073] |
| ENSG00000164331 | ANKRA2 | 1086 | 601 | 387 | -0,85 | -0,64 | ankyrin repeat family A member 2 [Source:HGNC Symbol;Acc:HGNC:13208] |
| ENSG00000164236 | ANKRD33B | 773 | 412 | 281 | -0,91 | -0,55 | ankyrin repeat domain 33B [Source:HGNC Symbol;Acc:HGNC:35240] |
| ENSG00000156381 | ANKRD9 | 2626 | 1426 | 1076 | -0,88 | -0,41 | ankyrin repeat domain 9 [Source:HGNC Symbol;Acc:HGNC:20096] |
| ENSG00000163516 | ANKZF1 | 3012 | 1766 | 1163 | -0,77 | -0,6 | ankyrin repeat and zinc finger domain containing 1 [Source:HGNC Symbol;Acc:HGNC:25527] |
| ENSG00000122359 | ANXA11 | 5393 | 3224 | 2427 | -0,74 | -0,41 | annexin A11 [Source:HGNC Symbol;Acc:HGNC:535] |
| ENSG00000143412 | ANXA9 | 534 | 49 | 30 | -3,44 | -0,72 | annexin A9 [Source:HGNC Symbol;Acc:HGNC:547] |
| ENSG00000089820 | ARHGAP4 | 467 | 237 | 144 | -0,98 | -0,72 | Rho GTPase activating protein 4 [Source:HGNC Symbol;Acc:HGNC:674] |
| ENSG00000130762 | ARHGEF16 | 4162 | 874 | 660 | -2,25 | -0,4 | Rho guanine nucleotide exchange factor 16 [Source:HGNC Symbol;Acc:HGNC:15515] |
| ENSG00000099889 | ARVCF | 2083 | 883 | 1260 | -1,24 | 0,51 | armadillo repeat gene deleted in velocardiofacial syndrome [Source:HGNC Symbol;Acc:HGNC:728] |
| ENSG00000104763 | ASAH1 | 2950 | 1777 | 1295 | -0,73 | -0,46 | N-acylsphingosine amidohydrolase 1 [Source:HGNC Symbol;Acc:HGNC:735] |
| ENSG00000162650 | ATXN7L2 | 575 | 327 | 522 | -0,81 | 0,68 | ataxin 7 like 2 [Source:HGNC Symbol;Acc:HGNC:28713] |
| ENSG00000153094 | BCL2L11 | 1164 | 662 | 971 | -0,82 | 0,55 | BCL2 like 11 [Source:HGNC Symbol;Acc:HGNC:994] |
| ENSG00000104081 | BMF | 8004 | 1695 | 1188 | -2,24 | -0,51 | Bcl2 modifying factor [Source:HGNC Symbol;Acc:HGNC:24132] |
| ENSG00000167535 | CACNB3 | 2867 | 1541 | 1186 | -0,9 | -0,38 | calcium voltage-gated channel auxiliary subunit beta 3 [Source:HGNC Symbol;Acc:HGNC:1403] |
| ENSG00000012822 | CALCOCO1 | 3871 | 1894 | 1254 | -1,03 | -0,59 | calcium binding and coiled-coil domain 1 [Source:HGNC Symbol;Acc:HGNC:29306] |
| ENSG00000079691 | CARMIL1 | 2109 | 1126 | 1586 | -0,91 | 0,49 | capping protein regulator and myosin 1 linker 1 [Source:HGNC Symbol;Acc:HGNC:21581] |
| ENSG00000130940 | CASZ1 | 1010 | 141 | 85 | -2,85 | -0,73 | castor zinc finger 1 [Source:HGNC Symbol;Acc:HGNC:26002] |
| ENSG00000105971 | CAV2 | 403 | 228 | 137 | -0,82 | -0,73 | caveolin 2 [Source:HGNC Symbol;Acc:HGNC:1528] |
| ENSG00000171219 | CDC42BPG | 2295 | 563 | 405 | -2,03 | -0,48 | CDC42 binding protein kinase gamma [Source:HGNC Symbol;Acc:HGNC:29829] |
| ENSG00000163075 | CFAP221 | 698 | 380 | 599 | -0,88 | 0,65 | cilia and flagella associated protein 221 [Source:HGNC Symbol;Acc:HGNC:33720] |
| ENSG00000184697 | CLDN6 | 16534 | 4373 | 6550 | -1,92 | 0,58 | claudin 6 [Source:HGNC Symbol;Acc:HGNC:2048] |
| ENSG00000038532 | CLEC16A | 1592 | 796 | 1113 | -1 | 0,48 | C-type lectin domain family 16 member A [Source:HGNC Symbol;Acc:HGNC:29013] |
| ENSG00000182372 | CLN8 | 1583 | 661 | 506 | -1,26 | -0,38 | ceroid-lipofuscinosis, neuronal 8 [Source:HGNC Symbol;Acc:HGNC:2079] |
| ENSG00000183723 | CMTM4 | 6233 | 3770 | 2934 | -0,73 | -0,36 | CKLF like MARVEL transmembrane domain containing 4 [Source:HGNC Symbol;Acc:HGNC:19175] |
| ENSG00000204650 | CRHR1-IT1 | 1400 | 771 | 518 | -0,86 | -0,57 | CRHR1 intronic transcript 1 [Source:EntrezGene;Acc:147081] |
| ENSG00000103316 | CRYM | 412 | 213 | 134 | -0,95 | -0,67 | crystallin mu [Source:HGNC Symbol;Acc:HGNC:2418] |
| ENSG00000104218 | CSPP1 | 2765 | 1345 | 2107 | -1,04 | 0,65 | centrosome and spindle pole associated protein 1 [Source:HGNC Symbol;Acc:HGNC:26193] |
| ENSG00000168772 | CXXC4 | 1517 | 460 | 327 | -1,72 | -0,5 | CXXC finger protein 4 [Source:HGNC Symbol;Acc:HGNC:24593] |
| ENSG00000166347 | CYB5A | 2468 | 1264 | 948 | -0,97 | -0,41 | cytochrome b5 type A [Source:HGNC Symbol;Acc:HGNC:2570] |
| ENSG00000051523 | CYBA | 3292 | 1756 | 1156 | -0,91 | -0,6 | cytochrome b-245 alpha chain [Source:HGNC Symbol;Acc:HGNC:2577] |
| ENSG00000276644 | DACH1 | 2785 | 1124 | 1657 | -1,31 | 0,56 | dachshund family transcription factor 1 [Source:HGNC Symbol;Acc:HGNC:2663] |
| ENSG00000179532 | DNHD1 | 1048 | 454 | 282 | -1,21 | -0,69 | dynein heavy chain domain 1 [Source:HGNC Symbol;Acc:HGNC:26532] |
| ENSG00000107554 | DNMBP | 2776 | 1600 | 1302 | -0,8 | -0,3 | dynamin binding protein [Source:HGNC Symbol;Acc:HGNC:30373] |
| ENSG00000110042 | DTX4 | 4015 | 1004 | 613 | -2 | -0,71 | deltex E3 ubiquitin ligase 4 [Source:HGNC Symbol;Acc:HGNC:29151] |
| ENSG00000163412 | EIF4E3 | 332 | 166 | 228 | -1 | 0,46 | eukaryotic translation initiation factor 4E family member 3 [Source:HGNC Symbol;Acc:HGNC:31837] |
| ENSG00000158711 | ELK4 | 1737 | 1001 | 661 | -0,8 | -0,6 | ELK4, ETS transcription factor [Source:HGNC Symbol;Acc:HGNC:3326] |
| ENSG00000132205 | EMILIN2 | 436 | 101 | 62 | -2,11 | -0,71 | elastin microfibril interfacer 2 [Source:HGNC Symbol;Acc:HGNC:19881] |
| ENSG00000167280 | ENGASE | 2193 | 926 | 627 | -1,24 | -0,56 | endo-beta-N-acetylglucosaminidase [Source:HGNC Symbol;Acc:HGNC:24622] |
| ENSG00000099219 | ERMP1 | 2933 | 1742 | 1059 | -0,75 | -0,72 | endoplasmic reticulum metallopeptidase 1 [Source:HGNC Symbol;Acc:HGNC:23703] |
| ENSG00000072840 | EVC | 1531 | 914 | 1240 | -0,74 | 0,44 | EvC ciliary complex subunit 1 [Source:HGNC Symbol;Acc:HGNC:3497] |
| ENSG00000173040 | EVC2 | 491 | 147 | 91 | -1,75 | -0,69 | EvC ciliary complex subunit 2 [Source:HGNC Symbol;Acc:HGNC:19747] |
| ENSG00000142449 | FBN3 | 4826 | 2315 | 1595 | -1,06 | -0,54 | fibrillin 3 [Source:HGNC Symbol;Acc:HGNC:18794] |
| ENSG00000127084 | FGD3 | 994 | 261 | 377 | -1,93 | 0,53 | FYVE, RhoGEF and PH domain containing 3 [Source:HGNC Symbol;Acc:HGNC:16027] |
| ENSG00000070404 | FSTL3 | 1276 | 487 | 365 | -1,39 | -0,42 | follistatin like 3 [Source:HGNC Symbol;Acc:HGNC:3973] |
| ENSG00000140564 | FURIN | 8479 | 2540 | 1705 | -1,74 | -0,57 | furin, paired basic amino acid cleaving enzyme [Source:HGNC Symbol;Acc:HGNC:8568] |
| ENSG00000141012 | GALNS | 592 | 326 | 486 | -0,86 | 0,58 | galactosamine (N-acetyl)-6-sulfatase [Source:HGNC Symbol;Acc:HGNC:4122] |
| ENSG00000164574 | GALNT10 | 1765 | 869 | 1109 | -1,02 | 0,35 | polypeptide N-acetylgalactosaminyltransferase 10 [Source:HGNC Symbol;Acc:HGNC:19873] |
| ENSG00000185340 | GAS2L1 | 376 | 195 | 275 | -0,95 | 0,5 | growth arrest specific 2 like 1 [Source:HGNC Symbol;Acc:HGNC:16955] |
| ENSG00000130513 | GDF15 | 3347 | 1772 | 1094 | -0,92 | -0,7 | growth differentiation factor 15 [Source:HGNC Symbol;Acc:HGNC:30142] |
| ENSG00000166123 | GPT2 | 3263 | 1559 | 2158 | -1,07 | 0,47 | glutamic--pyruvic transaminase 2 [Source:HGNC Symbol;Acc:HGNC:18062] |
| ENSG00000100429 | HDAC10 | 157 | 91 | 61 | -0,79 | -0,57 | histone deacetylase 10 [Source:HGNC Symbol;Acc:HGNC:18128] |
| ENSG00000010818 | HIVEP2 | 1300 | 754 | 511 | -0,79 | -0,56 | human immunodeficiency virus type I enhancer binding protein 2 [Source:HGNC Symbol;Acc:HGNC:4921] |
| ENSG00000003147 | ICA1 | 1395 | 258 | 164 | -2,44 | -0,65 | islet cell autoantigen 1 [Source:HGNC Symbol;Acc:HGNC:5343] |
| ENSG00000185436 | IFNLR1 | 555 | 335 | 412 | -0,73 | 0,3 | interferon lambda receptor 1 [Source:HGNC Symbol;Acc:HGNC:18584] |
| ENSG00000138002 | IFT172 | 836 | 460 | 727 | -0,86 | 0,66 | intraflagellar transport 172 [Source:HGNC Symbol;Acc:HGNC:30391] |
| ENSG00000115457 | IGFBP2 | 19496 | 10135 | 6560 | -0,94 | -0,63 | insulin like growth factor binding protein 2 [Source:HGNC Symbol;Acc:HGNC:5471] |
| ENSG00000269335 | IKBKG | 812 | 477 | 627 | -0,77 | 0,4 | inhibitor of kappa light polypeptide gene enhancer in B-cells, kinase gamma [Source:HGNC Symbol;Acc:HGNC:5961] |
| ENSG00000144711 | IQSEC1 | 1976 | 1117 | 808 | -0,82 | -0,47 | IQ motif and Sec7 domain 1 [Source:HGNC Symbol;Acc:HGNC:29112] |
| ENSG00000135424 | ITGA7 | 831 | 163 | 110 | -2,35 | -0,58 | integrin subunit alpha 7 [Source:HGNC Symbol;Acc:HGNC:6143] |
| ENSG00000137171 | KLC4 | 2030 | 848 | 604 | -1,26 | -0,49 | kinesin light chain 4 [Source:HGNC Symbol;Acc:HGNC:21624] |
| ENSG00000119138 | KLF9 | 570 | 147 | 91 | -1,96 | -0,68 | Kruppel like factor 9 [Source:HGNC Symbol;Acc:HGNC:1123] |
| ENSG00000109790 | KLHL5 | 4071 | 2023 | 1683 | -1,01 | -0,27 | kelch like family member 5 [Source:HGNC Symbol;Acc:HGNC:6356] |
| ENSG00000135338 | LCA5 | 636 | 355 | 221 | -0,84 | -0,69 | LCA5, lebercilin [Source:HGNC Symbol;Acc:HGNC:31923] |
| ENSG00000224081 | LINC01057 | 862 | 312 | 214 | -1,47 | -0,54 | long intergenic non-protein coding RNA 1057 [Source:HGNC Symbol;Acc:HGNC:49057] |
| ENSG00000228794 | LINC01128 | 458 | 253 | 331 | -0,85 | 0,39 | long intergenic non-protein coding RNA 1128 [Source:HGNC Symbol;Acc:HGNC:49377] |

| | | | | | | | |
|---|---|---|---|---|---|---|---|
| ENSG00000073350 | LLGL2 | 7787 | 3139 | 2510 | -1,31 | -0,32 | LLGL2, scribble cell polarity complex component [Source:HGNC Symbol;Acc:HGNC:6629] |
| ENSG00000176454 | LPCAT4 | 3360 | 1881 | 1410 | -0,84 | -0,42 | lysophosphatidylcholine acyltransferase 4 [Source:HGNC Symbol;Acc:HGNC:30059] |
| ENSG00000127603 | MACF1 | 11310 | 6455 | 9492 | -0,81 | 0,56 | microtubule-actin crosslinking factor 1 [Source:HGNC Symbol;Acc:HGNC:13664] |
| ENSG00000198517 | MAFK | 2529 | 645 | 438 | -1,97 | -0,55 | MAF bZIP transcription factor K [Source:HGNC Symbol;Acc:HGNC:6782] |
| ENSG00000198934 | MAGEE1 | 509 | 268 | 177 | -0,92 | -0,6 | MAGE family member E1 [Source:HGNC Symbol;Acc:HGNC:24934] |
| ENSG00000139793 | MBNL2 | 2756 | 1295 | 860 | -1,09 | -0,59 | muscleblind like splicing regulator 2 [Source:HGNC Symbol;Acc:HGNC:16746] |
| ENSG00000067365 | METTL22 | 1961 | 784 | 545 | -1,32 | -0,52 | methyltransferase like 22 [Source:HGNC Symbol;Acc:HGNC:28368] |
| ENSG00000157227 | MMP14 | 23004 | 13733 | 9832 | -0,74 | -0,48 | matrix metallopeptidase 14 [Source:HGNC Symbol;Acc:HGNC:7160] |
| ENSG00000133131 | MORC4 | 908 | 491 | 316 | -0,89 | -0,63 | MORC family CW-type zinc finger 4 [Source:HGNC Symbol;Acc:HGNC:23485] |
| ENSG00000170873 | MTSS1 | 1563 | 672 | 427 | -1,22 | -0,65 | MTSS1, I-BAR domain containing [Source:HGNC Symbol;Acc:HGNC:20443] |
| ENSG00000059728 | MXD1 | 888 | 452 | 291 | -0,97 | -0,63 | MAX dimerization protein 1 [Source:HGNC Symbol;Acc:HGNC:6761] |
| ENSG00000172936 | MYD88 | 941 | 428 | 260 | -1,14 | -0,72 | myeloid differentiation primary response 88 [Source:HGNC Symbol;Acc:HGNC:7562] |
| ENSG00000167306 | MYO5B | 2211 | 352 | 215 | -2,65 | -0,71 | myosin VB [Source:HGNC Symbol;Acc:HGNC:7603] |
| ENSG00000274180 | NATD1 | 1238 | 559 | 349 | -1,15 | -0,68 | N-acetyltransferase domain containing 1 [Source:HGNC Symbol;Acc:HGNC:30770] |
| ENSG00000134369 | NAV1 | 2311 | 927 | 1361 | -1,32 | 0,55 | neuron navigator 1 [Source:HGNC Symbol;Acc:HGNC:15989] |
| ENSG00000160796 | NBEAL2 | 10013 | 2350 | 3158 | -2,09 | 0,43 | neurobeachin like 2 [Source:HGNC Symbol;Acc:HGNC:31928] |
| ENSG00000273136 | NBPF26 | 999 | 546 | 426 | -0,87 | -0,36 | neuroblastoma breakpoint family member 26 [Source:HGNC Symbol;Acc:HGNC:49571] |
| ENSG00000165795 | NDRG2 | 2005 | 1065 | 699 | -0,91 | -0,61 | NDRG family member 2 [Source:HGNC Symbol;Acc:HGNC:14460] |
| ENSG00000140398 | NEIL1 | 975 | 260 | 163 | -1,91 | -0,68 | nei like DNA glycosylase 1 [Source:HGNC Symbol;Acc:HGNC:18448] |
| ENSG00000170322 | NFRKB | 2923 | 1549 | 2242 | -0,92 | 0,53 | nuclear factor related to kappaB binding protein [Source:HGNC Symbol;Acc:HGNC:7802] |
| ENSG00000151014 | NOCT | 412 | 211 | 292 | -0,96 | 0,47 | nocturnin [Source:HGNC Symbol;Acc:HGNC:14254] |
| ENSG00000134250 | NOTCH2 | 15770 | 7135 | 5053 | -1,14 | -0,5 | notch 2 [Source:HGNC Symbol;Acc:HGNC:7882] |
| ENSG00000215440 | NPEPL1 | 1283 | 485 | 291 | -1,4 | -0,73 | aminopeptidase-like 1 [Source:HGNC Symbol;Acc:HGNC:16244] |
| ENSG00000135124 | P2RX4 | 1741 | 932 | 676 | -0,9 | -0,46 | purinergic receptor P2X 4 [Source:HGNC Symbol;Acc:HGNC:8535] |
| ENSG00000132849 | PATJ | 3641 | 1606 | 994 | -1,18 | -0,69 | PATJ, crumbs cell polarity complex component [Source:HGNC Symbol;Acc:HGNC:28881] |
| ENSG00000273344 | PAXIP1-AS1 | 329 | 176 | 111 | -0,9 | -0,66 | PAXIP1 antisense RNA 1 (head to head) [Source:HGNC Symbol;Acc:HGNC:27328] |
| ENSG00000177839 | PCDHB9 | 248 | 142 | 95 | -0,81 | -0,59 | protocadherin beta 9 [Source:HGNC Symbol;Acc:HGNC:8694] |
| ENSG00000160613 | PCSK7 | 1364 | 801 | 1054 | -0,77 | 0,4 | proprotein convertase subtilisin/kexin type 7 [Source:HGNC Symbol;Acc:HGNC:8748] |
| ENSG00000172572 | PDE3A | 4645 | 1647 | 2370 | -1,5 | 0,52 | phosphodiesterase 3A [Source:HGNC Symbol;Acc:HGNC:8778] |
| ENSG00000142102 | PGGHG | 11024 | 6231 | 4393 | -0,82 | -0,5 | protein-glucosylgalactosylhydroxylysine glucosidase [Source:HGNC Symbol;Acc:HGNC:26210] |
| ENSG00000171608 | PIK3CD | 988 | 441 | 284 | -1,16 | -0,64 | phosphatidylinositol-4,5-bisphosphate 3-kinase catalytic subunit delta [Source:HGNC Symbol;Acc:HGNC:8977] |
| ENSG00000164050 | PLXNB1 | 15053 | 6940 | 10824 | -1,12 | 0,64 | plexin B1 [Source:HGNC Symbol;Acc:HGNC:9103] |
| ENSG00000186866 | POFUT2 | 4794 | 2714 | 2089 | -0,82 | -0,38 | protein O-fucosyltransferase 2 [Source:HGNC Symbol;Acc:HGNC:14683] |
| ENSG00000118898 | PPL | 5046 | 990 | 612 | -2,35 | -0,69 | periplakin [Source:HGNC Symbol;Acc:HGNC:9273] |
| ENSG00000104881 | PPP1R13L | 1533 | 394 | 251 | -1,96 | -0,65 | protein phosphatase 1 regulatory subunit 13 like [Source:HGNC Symbol;Acc:HGNC:18838] |
| ENSG00000106617 | PRKAG2 | 351 | 208 | 268 | -0,75 | 0,37 | protein kinase AMP-activated non-catalytic subunit gamma 2 [Source:HGNC Symbol;Acc:HGNC:9386] |
| ENSG00000188191 | PRKAR1B | 1033 | 503 | 330 | -1,04 | -0,61 | protein kinase cAMP-dependent type I regulatory subunit beta [Source:HGNC Symbol;Acc:HGNC:9390] |
| ENSG00000067606 | PRKCZ | 1866 | 1067 | 709 | -0,81 | -0,59 | protein kinase C zeta [Source:HGNC Symbol;Acc:HGNC:9412] |
| ENSG00000173482 | PTPRM | 2140 | 785 | 536 | -1,45 | -0,55 | protein tyrosine phosphatase, receptor type M [Source:HGNC Symbol;Acc:HGNC:9675] |
| ENSG00000156675 | RAB11FIP1 | 5817 | 2647 | 1780 | -1,14 | -0,57 | RAB11 family interacting protein 1 [Source:HGNC Symbol;Acc:HGNC:30265] |
| ENSG00000139998 | RAB15 | 934 | 531 | 825 | -0,81 | 0,64 | RAB15, member RAS oncogene family [Source:HGNC Symbol;Acc:HGNC:20150] |
| ENSG00000105514 | RAB3D | 1268 | 743 | 592 | -0,77 | -0,33 | RAB3D, member RAS oncogene family [Source:HGNC Symbol;Acc:HGNC:9779] |
| ENSG00000177548 | RABEP2 | 732 | 385 | 591 | -0,93 | 0,62 | rabaptin, RAB GTPase binding effector protein 2 [Source:HGNC Symbol;Acc:HGNC:24817] |
| ENSG00000136828 | RALGPS1 | 1703 | 439 | 303 | -1,95 | -0,54 | Ral GEF with PH domain and SH3 binding motif 1 [Source:HGNC Symbol;Acc:HGNC:16851] |
| ENSG00000076864 | RAP1GAP | 2893 | 1331 | 999 | -1,12 | -0,41 | RAP1 GTPase activating protein [Source:HGNC Symbol;Acc:HGNC:9858] |
| ENSG00000076067 | RBMS2 | 3896 | 1609 | 1166 | -1,28 | -0,46 | RNA binding motif single stranded interacting protein 2 [Source:HGNC Symbol;Acc:HGNC:9909] |
| ENSG00000117602 | RCAN3 | 1906 | 1140 | 757 | -0,74 | -0,59 | RCAN family member 3 [Source:HGNC Symbol;Acc:HGNC:3042] |
| ENSG00000205517 | RGL3 | 4506 | 917 | 645 | -2,3 | -0,51 | ral guanine nucleotide dissociation stimulator like 3 [Source:HGNC Symbol;Acc:HGNC:30282] |
| ENSG00000183421 | RIPK4 | 2503 | 1368 | 1015 | -0,87 | -0,43 | receptor interacting serine/threonine kinase 4 [Source:HGNC Symbol;Acc:HGNC:496] |
| ENSG00000034677 | RNF19A | 1776 | 1057 | 779 | -0,75 | -0,44 | ring finger protein 19A, RBR E3 ubiquitin protein ligase [Source:HGNC Symbol;Acc:HGNC:13432] |
| ENSG00000108375 | RNF43 | 763 | 439 | 342 | -0,8 | -0,36 | ring finger protein 43 [Source:HGNC Symbol;Acc:HGNC:18505] |
| ENSG00000069667 | RORA | 519 | 269 | 167 | -0,95 | -0,69 | RAR related orphan receptor A [Source:HGNC Symbol;Acc:HGNC:10258] |
| ENSG00000269609 | RPARP-AS1 | 464 | 183 | 125 | -1,34 | -0,56 | RPARP antisense RNA 1 [Source:HGNC Symbol;Acc:HGNC:45238] |
| ENSG00000117616 | RSRP1 | 2202 | 1159 | 715 | -0,93 | -0,7 | arginine and serine rich protein 1 [Source:HGNC Symbol;Acc:HGNC:25234] |
| ENSG00000197956 | S100A6 | 5288 | 2053 | 1495 | -1,36 | -0,46 | S100 calcium binding protein A6 [Source:HGNC Symbol;Acc:HGNC:10496] |
| ENSG00000198794 | SCAMP5 | 2211 | 679 | 874 | -1,7 | 0,36 | secretory carrier membrane protein 5 [Source:HGNC Symbol;Acc:HGNC:30386] |
| ENSG00000124145 | SDC4 | 12198 | 6145 | 4026 | -0,99 | -0,61 | syndecan 4 [Source:HGNC Symbol;Acc:HGNC:10661] |
| ENSG00000001617 | SEMA3F | 1419 | 458 | 328 | -1,63 | -0,48 | semaphorin 3F [Source:HGNC Symbol;Acc:HGNC:10728] |
| ENSG00000187764 | SEMA4D | 1820 | 1094 | 828 | -0,73 | -0,4 | semaphorin 4D [Source:HGNC Symbol;Acc:HGNC:10732] |
| ENSG00000120057 | SFRP5 | 8725 | 2109 | 1315 | -2,05 | -0,68 | secreted frizzled related protein 5 [Source:HGNC Symbol;Acc:HGNC:10779] |
| ENSG00000187531 | SIRT7 | 585 | 300 | 427 | -0,96 | 0,51 | sirtuin 7 [Source:HGNC Symbol;Acc:HGNC:14935] |
| ENSG00000140199 | SLC12A6 | 1895 | 1117 | 868 | -0,76 | -0,37 | solute carrier family 12 member 6 [Source:HGNC Symbol;Acc:HGNC:10914] |
| ENSG00000174327 | SLC16A13 | 305 | 159 | 255 | -0,94 | 0,68 | solute carrier family 16 member 13 [Source:HGNC Symbol;Acc:HGNC:31037] |
| ENSG00000197375 | SLC22A5 | 1265 | 430 | 280 | -1,56 | -0,62 | solute carrier family 22 member 5 [Source:HGNC Symbol;Acc:HGNC:10969] |
| ENSG00000197119 | SLC25A29 | 6600 | 2491 | 3331 | -1,41 | 0,42 | solute carrier family 25 member 29 [Source:HGNC Symbol;Acc:HGNC:20116] |
| ENSG00000080493 | SLC4A4 | 13648 | 3764 | 5529 | -1,86 | 0,55 | solute carrier family 4 member 4 [Source:HGNC Symbol;Acc:HGNC:11030] |
| ENSG00000130821 | SLC6A8 | 2442 | 1415 | 951 | -0,79 | -0,57 | solute carrier family 6 member 8 [Source:HGNC Symbol;Acc:HGNC:11055] |
| ENSG00000089060 | SLC8B1 | 707 | 281 | 199 | -1,33 | -0,49 | solute carrier family 8 member B1 [Source:HGNC Symbol;Acc:HGNC:26175] |
| ENSG00000065923 | SLC9A7 | 1141 | 475 | 314 | -1,26 | -0,6 | solute carrier family 9 member A7 [Source:HGNC Symbol;Acc:HGNC:17123] |
| ENSG00000166949 | SMAD3 | 4660 | 2016 | 1559 | -1,21 | -0,37 | SMAD family member 3 [Source:HGNC Symbol;Acc:HGNC:6769] |
| ENSG00000120693 | SMAD9 | 2125 | 918 | 560 | -1,21 | -0,71 | SMAD family member 9 [Source:HGNC Symbol;Acc:HGNC:6774] |
| ENSG00000105771 | SMG9 | 1470 | 882 | 1137 | -0,74 | 0,37 | SMG9, nonsense mediated mRNA decay factor [Source:HGNC Symbol;Acc:HGNC:25763] |
| ENSG00000144451 | SPAG16 | 1047 | 569 | 441 | -0,88 | -0,37 | sperm associated antigen 16 [Source:HGNC Symbol;Acc:HGNC:23225] |
| ENSG00000144228 | SPOPL | 2196 | 1265 | 917 | -0,8 | -0,47 | speckle type BTB/POZ protein like [Source:HGNC Symbol;Acc:HGNC:27934] |
| ENSG00000173898 | SPTBN2 | 1787 | 813 | 1233 | -1,14 | 0,6 | spectrin beta, non-erythrocytic 2 [Source:HGNC Symbol;Acc:HGNC:11276] |
| ENSG00000242294 | STAG3L5P | 405 | 228 | 352 | -0,83 | 0,63 | stromal antigen 3-like 5 pseudogene [Source:HGNC Symbol;Acc:HGNC:48896] |
| ENSG00000214530 | STARD10 | 6433 | 3793 | 2421 | -0,76 | -0,65 | StAR related lipid transfer domain containing 10 [Source:HGNC Symbol;Acc:HGNC:10666] |
| ENSG00000163482 | STK36 | 2391 | 1008 | 1542 | -1,25 | 0,61 | serine/threonine kinase 36 [Source:HGNC Symbol;Acc:HGNC:17209] |
| ENSG00000076944 | STXBP2 | 1394 | 691 | 550 | -1,01 | -0,33 | syntaxin binding protein 2 [Source:HGNC Symbol;Acc:HGNC:11445] |
| ENSG00000097096 | SYDE2 | 626 | 193 | 284 | -1,7 | 0,56 | synapse defective Rho GTPase homolog 2 [Source:HGNC Symbol;Acc:HGNC:25841] |
| ENSG00000100321 | SYNGR1 | 4196 | 2483 | 1529 | -0,76 | -0,7 | synaptogyrin 1 [Source:HGNC Symbol;Acc:HGNC:11498] |
| ENSG00000158710 | TAGLN2 | 7973 | 3835 | 2355 | -1,06 | -0,7 | transgelin 2 [Source:HGNC Symbol;Acc:HGNC:11554] |
| ENSG00000183597 | TANGO2 | 368 | 188 | 256 | -0,97 | 0,44 | transport and golgi organization 2 homolog [Source:HGNC Symbol;Acc:HGNC:25439] |
| ENSG00000110719 | TCIRG1 | 3890 | 1478 | 942 | -1,4 | -0,65 | T-cell immune regulator 1, ATPase H+ transporting V0 subunit a3 [Source:HGNC Symbol;Acc:HGNC:11647] |
| ENSG00000167302 | TEPSIN | 787 | 455 | 654 | -0,79 | 0,52 | TEPSIN, adaptor related protein complex 4 accessory protein [Source:HGNC Symbol;Acc:HGNC:26458] |
| ENSG00000072274 | TFRC | 13600 | 6821 | 4806 | -1 | -0,51 | transferrin receptor [Source:HGNC Symbol;Acc:HGNC:11763] |
| ENSG00000105329 | TGFB1 | 4610 | 1949 | 1192 | -1,24 | -0,71 | transforming growth factor beta 1 [Source:HGNC Symbol;Acc:HGNC:11766] |
| ENSG00000127666 | TICAM1 | 386 | 159 | 111 | -1,28 | -0,52 | toll like receptor adaptor molecule 1 [Source:HGNC Symbol;Acc:HGNC:18348] |
| ENSG00000163659 | TIPARP | 1698 | 971 | 678 | -0,81 | -0,52 | TCDD inducible poly(ADP-ribose) polymerase [Source:HGNC Symbol;Acc:HGNC:23696] |
| ENSG00000162542 | TMCO4 | 387 | 141 | 88 | -1,46 | -0,68 | transmembrane and coiled-coil domains 4 [Source:HGNC Symbol;Acc:HGNC:27393] |
| ENSG00000181284 | TMEM102 | 447 | 257 | 184 | -0,8 | -0,48 | transmembrane protein 102 [Source:HGNC Symbol;Acc:HGNC:26722] |
| ENSG00000152078 | TMEM56 | 1886 | 974 | 646 | -0,95 | -0,59 | transmembrane protein 56 [Source:HGNC Symbol;Acc:HGNC:26477] |
| ENSG00000116857 | TMEM9 | 5641 | 3218 | 2489 | -0,81 | -0,37 | transmembrane protein 9 [Source:HGNC Symbol;Acc:HGNC:18823] |
| ENSG00000078804 | TP53INP2 | 1152 | 479 | 732 | -1,27 | 0,61 | tumor protein p53 inducible nuclear protein 2 [Source:HGNC Symbol;Acc:HGNC:16104] |
| ENSG00000112343 | TRIM38 | 1374 | 669 | 457 | -1,04 | -0,55 | tripartite motif containing 38 [Source:HGNC Symbol;Acc:HGNC:10059] |
| ENSG00000166436 | TRIM66 | 457 | 223 | 348 | -1,04 | 0,64 | tripartite motif containing 66 [Source:HGNC Symbol;Acc:HGNC:29005] |
| ENSG00000167333 | TRIM68 | 806 | 302 | 188 | -1,42 | -0,68 | tripartite motif containing 68 [Source:HGNC Symbol;Acc:HGNC:21161] |
| ENSG00000087077 | TRIP6 | 2743 | 1500 | 1139 | -0,87 | -0,4 | thyroid hormone receptor interactor 6 [Source:HGNC Symbol;Acc:HGNC:12311] |
| ENSG00000121297 | TSHZ3 | 515 | 311 | 431 | -0,73 | 0,47 | teashirt zinc finger homeobox 3 [Source:HGNC Symbol;Acc:HGNC:30700] |
| ENSG00000099282 | TSPAN15 | 968 | 478 | 342 | -1,02 | -0,48 | tetraspanin 15 [Source:HGNC Symbol;Acc:HGNC:23298] |
| ENSG00000005379 | TSPOAP1 | 1553 | 546 | 372 | -1,51 | -0,56 | TSPO associated protein 1 [Source:HGNC Symbol;Acc:HGNC:16831] |
| ENSG00000133985 | TTC9 | 575 | 304 | 417 | -0,92 | 0,46 | tetratricopeptide repeat domain 9 [Source:HGNC Symbol;Acc:HGNC:20267] |
| ENSG00000173218 | VANGL1 | 1104 | 651 | 1008 | -0,76 | 0,63 | VANGL planar cell polarity protein 1 [Source:HGNC Symbol;Acc:HGNC:15512] |
| ENSG00000171044 | XKR6 | 197 | 106 | 70 | -0,9 | -0,6 | XK related 6 [Source:HGNC Symbol;Acc:HGNC:27806] |
| ENSG00000130733 | YIPF2 | 3347 | 1945 | 1307 | -0,78 | -0,57 | Yip1 domain family member 2 [Source:HGNC Symbol;Acc:HGNC:28476] |
| ENSG00000175155 | YPEL2 | 2850 | 955 | 631 | -1,58 | -0,6 | yippee like 2 [Source:HGNC Symbol;Acc:HGNC:18326] |
| ENSG00000090238 | YPEL3 | 1352 | 496 | 316 | -1,45 | -0,65 | yippee like 3 [Source:HGNC Symbol;Acc:HGNC:18327] |
| ENSG00000175048 | ZDHHC14 | 349 | 171 | 261 | -1,03 | 0,61 | zinc finger DHHC-type containing 14 [Source:HGNC Symbol;Acc:HGNC:20341] |
| ENSG00000174586 | ZNF497 | 207 | 115 | 183 | -0,85 | 0,67 | zinc finger protein 497 [Source:HGNC Symbol;Acc:HGNC:23714] |
| ENSG00000188171 | ZNF626 | 597 | 301 | 443 | -0,99 | 0,56 | zinc finger protein 626 [Source:HGNC Symbol;Acc:HGNC:30461] |
| ENSG00000151612 | ZNF827 | 1459 | 848 | 623 | -0,78 | -0,45 | zinc finger protein 827 [Source:HGNC Symbol;Acc:HGNC:27193] |
| ENSG00000106479 | ZNF862 | 1094 | 645 | 460 | -0,76 | -0,49 | zinc finger protein 862 [Source:HGNC Symbol;Acc:HGNC:34519] |

**GROUP 4**

| | | | | | | | |
|---|---|---|---|---|---|---|---|
| ENSG00000107331 | ABCA2 | 6642 | 3838 | 1571 | -0,79 | -1,29 | ATP binding cassette subfamily A member 2 [Source:HGNC Symbol;Acc:HGNC:32] |
| ENSG00000108846 | ABCC3 | 207 | 47 | 13 | -2,15 | -1,87 | ATP binding cassette subfamily C member 3 [Source:HGNC Symbol;Acc:HGNC:54] |
| ENSG00000114626 | ABTB1 | 710 | 271 | 150 | -1,39 | -0,86 | ankyrin repeat and BTB domain containing 1 [Source:HGNC Symbol;Acc:HGNC:18275] |
| ENSG00000181513 | ACBD4 | 703 | 370 | 212 | -0,93 | -0,8 | acyl-CoA binding domain containing 4 [Source:HGNC Symbol;Acc:HGNC:23337] |
| ENSG00000110455 | ACCS | 505 | 146 | 54 | -1,79 | -1,44 | 1-aminocyclopropane-1-carboxylate synthase homolog (inactive) [Source:HGNC Symbol;Acc:HGNC:23989] |
| ENSG00000102575 | ACP5 | 250 | 42 | 11 | -2,57 | -1,94 | acid phosphatase 5, tartrate resistant [Source:HGNC Symbol;Acc:HGNC:124] |
| ENSG00000049192 | ADAMTS6 | 350 | 55 | 18 | -2,68 | -1,57 | ADAM metallopeptidase with thrombospondin type 1 motif 6 [Source:HGNC Symbol;Acc:HGNC:222] |
| ENSG00000121753 | ADGRB2 | 970 | 570 | 296 | -0,77 | -0,95 | adhesion G protein-coupled receptor B2 [Source:HGNC Symbol;Acc:HGNC:944] |
| ENSG00000164393 | ADGRF2 | 1127 | 52 | 18 | -4,43 | -1,6 | adhesion G protein-coupled receptor F2 [Source:HGNC Symbol;Acc:HGNC:18991] |
| ENSG00000153294 | ADGRF4 | 736 | 37 | 8 | -4,32 | -2,21 | adhesion G protein-coupled receptor F4 [Source:HGNC Symbol;Acc:HGNC:19011] |
| ENSG00000205336 | ADGRG1 | 1200 | 264 | 151 | -2,18 | -0,81 | adhesion G protein-coupled receptor G1 [Source:HGNC Symbol;Acc:HGNC:4512] |
| ENSG00000150594 | ADRA2A | 397 | 15 | 5 | -4,75 | -1,45 | adrenoceptor alpha 2A [Source:HGNC Symbol;Acc:HGNC:281] |
| ENSG00000184160 | ADRA2C | 455 | 42 | 22 | -3,44 | -0,93 | adrenoceptor alpha 2C [Source:HGNC Symbol;Acc:HGNC:283] |
| ENSG00000185100 | ADSSL1 | 801 | 355 | 149 | -1,18 | -1,26 | adenylosuccinate synthase like 1 [Source:HGNC Symbol;Acc:HGNC:20093] |
| ENSG00000169129 | AFAP1L2 | 313 | 80 | 21 | -1,97 | -1,94 | actin filament associated protein 1 like 2 [Source:HGNC Symbol;Acc:HGNC:25901] |
| ENSG00000176092 | AIM1L | 1217 | 237 | 95 | -2,36 | -1,32 | absent in melanoma 1-like [Source:HGNC Symbol;Acc:HGNC:17295] |
| ENSG00000117020 | AKT3 | 1621 | 632 | 119 | -1,36 | -2,41 | AKT serine/threonine kinase 3 [Source:HGNC Symbol;Acc:HGNC:393] |
| ENSG00000170017 | ALCAM | 2455 | 763 | 216 | -1,69 | -1,82 | activated leukocyte cell adhesion molecule [Source:HGNC Symbol;Acc:HGNC:400] |
| ENSG00000165092 | ALDH1A1 | 4374 | 1131 | 172 | -1,95 | -2,72 | aldehyde dehydrogenase 1 family member A1 [Source:HGNC Symbol;Acc:HGNC:402] |
| ENSG00000012779 | ALOX5 | 293 | 26 | 5 | -3,51 | -2,48 | arachidonate 5-lipoxygenase [Source:HGNC Symbol;Acc:HGNC:435] |
| ENSG00000178038 | ALS2CL | 1696 | 175 | 42 | -3,28 | -2,05 | ALS2 C-terminal like [Source:HGNC Symbol;Acc:HGNC:20605] |
| ENSG00000166126 | AMN | 318 | 125 | 24 | -1,34 | -2,38 | amnion associated transmembrane protein [Source:HGNC Symbol;Acc:HGNC:14604] |
| ENSG00000151150 | ANK3 | 2078 | 1094 | 483 | -0,93 | -1,18 | ankyrin 3 [Source:HGNC Symbol;Acc:HGNC:494] |
| ENSG00000185101 | ANO9 | 3949 | 1488 | 518 | -1,41 | -1,52 | anoctamin 9 [Source:HGNC Symbol;Acc:HGNC:20679] |
| ENSG00000011201 | ANOS1 | 272 | 34 | 12 | -3,01 | -1,46 | anosmin 1 [Source:HGNC Symbol;Acc:HGNC:6211] |
| ENSG00000131480 | AOC2 | 322 | 90 | 50 | -1,84 | -0,86 | amine oxidase, copper containing 2 [Source:HGNC Symbol;Acc:HGNC:549] |
| ENSG00000243811 | APOBEC3D | 260 | 121 | 47 | -1,11 | -1,37 | apolipoprotein B mRNA editing enzyme catalytic subunit 3D [Source:HGNC Symbol;Acc:HGNC:17354] |
| ENSG00000159314 | ARHGAP27 | 791 | 210 | 89 | -1,91 | -1,24 | Rho GTPase activating protein 27 [Source:HGNC Symbol;Acc:HGNC:31813] |
| ENSG00000180448 | ARHGAP45 | 583 | 204 | 59 | -1,51 | -1,79 | Rho GTPase activating protein 45 [Source:HGNC Symbol;Acc:HGNC:17102] |
| ENSG00000241484 | ARHGAP8 | 640 | 14 | 0 | -5,49 | -5,33 | Rho GTPase activating protein 8 [Source:HGNC Symbol;Acc:HGNC:677] |
| ENSG00000111348 | ARHGDIB | 189 | 95 | 27 | -0,99 | -1,82 | Rho GDP dissociation inhibitor beta [Source:HGNC Symbol;Acc:HGNC:679] |
| ENSG00000204959 | ARHGEF34P | 1799 | 384 | 72 | -2,23 | -2,41 | Rho guanine nucleotide exchange factor 34, pseudogene [Source:HGNC Symbol;Acc:HGNC:38086] |
| ENSG00000213214 | ARHGEF35 | 862 | 216 | 44 | -1,99 | -2,31 | Rho guanine nucleotide exchange factor 35 [Source:HGNC Symbol;Acc:HGNC:33846] |
| ENSG00000050327 | ARHGEF5 | 443 | 118 | 27 | -1,91 | -2,1 | Rho guanine nucleotide exchange factor 5 [Source:HGNC Symbol;Acc:HGNC:13209] |
| ENSG00000126947 | ARMCX1 | 978 | 572 | 120 | -0,77 | -2,25 | armadillo repeat containing, X-linked 1 [Source:HGNC Symbol;Acc:HGNC:18073] |
| ENSG00000184867 | ARMCX2 | 2143 | 1224 | 424 | -0,81 | -1,53 | armadillo repeat containing, X-linked 2 [Source:HGNC Symbol;Acc:HGNC:16869] |
| ENSG00000196440 | ARMCX4 | 932 | 290 | 76 | -1,68 | -1,93 | armadillo repeat containing, X-linked 4 [Source:HGNC Symbol;Acc:HGNC:28615] |
| ENSG00000174939 | ASPHD1 | 343 | 61 | 15 | -2,5 | -2,03 | aspartate beta-hydroxylase domain containing 1 [Source:HGNC Symbol;Acc:HGNC:27380] |
| ENSG00000130707 | ASS1 | 240 | 132 | 33 | -0,87 | -1,99 | argininosuccinate synthase 1 [Source:HGNC Symbol;Acc:HGNC:758] |
| ENSG00000064270 | ATP2C2 | 852 | 221 | 33 | -1,95 | -2,74 | ATPase secretory pathway Ca2+ transporting 2 [Source:HGNC Symbol;Acc:HGNC:29103] |
| ENSG00000081923 | ATP8B1 | 1593 | 225 | 37 | -2,82 | -2,59 | ATPase phospholipid transporting 8B1 [Source:HGNC Symbol;Acc:HGNC:3706] |
| ENSG00000172232 | AZU1 | 255 | 62 | 17 | -2,04 | -1,87 | azurocidin 1 [Source:HGNC Symbol;Acc:HGNC:913] |
| ENSG00000109956 | B3GAT1 | 664 | 108 | 17 | -2,62 | -2,7 | beta-1,3-glucuronyltransferase 1 [Source:HGNC Symbol;Acc:HGNC:921] |
| ENSG00000156966 | B3GNT7 | 1071 | 295 | 106 | -1,86 | -1,47 | UDP-GlcNAc:betaGal beta-1,3-N-acetylglucosaminyltransferase 7 [Source:HGNC Symbol;Acc:HGNC:18811] |
| ENSG00000237172 | B3GNT9 | 308 | 170 | 71 | -0,86 | -1,25 | UDP-GlcNAc:betaGal beta-1,3-N-acetylglucosaminyltransferase 9 [Source:HGNC Symbol;Acc:HGNC:28714] |
| ENSG00000007516 | BAIAP3 | 831 | 497 | 260 | -0,74 | -0,93 | BAI1 associated protein 3 [Source:HGNC Symbol;Acc:HGNC:948] |
| ENSG00000119866 | BCL11A | 268 | 10 | 1 | -4,79 | -3,31 | B-cell CLL/lymphoma 11A [Source:HGNC Symbol;Acc:HGNC:13221] |
| ENSG00000113916 | BCL6 | 1484 | 535 | 212 | -1,47 | -1,33 | B-cell CLL/lymphoma 6 [Source:HGNC Symbol;Acc:HGNC:1001] |
| ENSG00000136717 | BIN1 | 2808 | 1469 | 830 | -0,93 | -0,82 | bridging integrator 1 [Source:HGNC Symbol;Acc:HGNC:1052] |
| ENSG00000281406 | BLACAT1 | 731 | 159 | 26 | -2,2 | -2,61 | bladder cancer associated transcript 1 (non-protein coding) [Source:HGNC Symbol;Acc:HGNC:48597] |
| ENSG00000176720 | BOK | 917 | 192 | 70 | -2,26 | -1,45 | BOK, BCL2 family apoptosis regulator [Source:HGNC Symbol;Acc:HGNC:1087] |
| ENSG00000119411 | BSPRY | 626 | 58 | 6 | -3,42 | -3,19 | B-box and SPRY domain containing [Source:HGNC Symbol;Acc:HGNC:18232] |
| ENSG00000107738 | C10orf54 | 588 | 225 | 103 | -1,39 | -1,12 | chromosome 10 open reading frame 54 [Source:HGNC Symbol;Acc:HGNC:30085] |
| ENSG00000165863 | C10orf82 | 916 | 204 | 23 | -2,17 | -3,12 | chromosome 10 open reading frame 82 [Source:HGNC Symbol;Acc:HGNC:28500] |
| ENSG00000109944 | C11orf63 | 210 | 92 | 52 | -1,19 | -0,83 | chromosome 11 open reading frame 63 [Source:HGNC Symbol;Acc:HGNC:26288] |
| ENSG00000253313 | C1orf210 | 887 | 156 | 50 | -2,5 | -1,66 | chromosome 1 open reading frame 210 [Source:HGNC Symbol;Acc:HGNC:28755] |

| | | | | | | | |
|---|---|---|---|---|---|---|---|
| ENSG00000159403 | C1R | 403 | 244 | 105 | -0,73 | -1,21 | complement C1r [Source:HGNC Symbol;Acc:HGNC:1246] |
| ENSG00000174403 | C20orf166-AS1 | 339 | 28 | 3 | -3,58 | -3,45 | C20orf166 antisense RNA 1 [Source:HGNC Symbol;Acc:HGNC:26393] |
| ENSG00000273045 | C2orf15 | 273 | 91 | 49 | -1,58 | -0,9 | chromosome 2 open reading frame 15 [Source:HGNC Symbol;Acc:HGNC:28436] |
| ENSG00000100346 | CACNA1I | 331 | 44 | 2 | -2,91 | -4,26 | calcium voltage-gated channel subunit alpha1 I [Source:HGNC Symbol;Acc:HGNC:1396] |
| ENSG00000130433 | CACNG6 | 958 | 285 | 54 | -1,75 | -2,39 | calcium voltage-gated channel auxiliary subunit gamma 6 [Source:HGNC Symbol;Acc:HGNC:13625] |
| ENSG00000142408 | CACNG8 | 270 | 32 | 7 | -3,07 | -2,2 | calcium voltage-gated channel auxiliary subunit gamma 8 [Source:HGNC Symbol;Acc:HGNC:13628] |
| ENSG00000183049 | CAMK1D | 516 | 107 | 40 | -2,26 | -1,43 | calcium/calmodulin dependent protein kinase ID [Source:HGNC Symbol;Acc:HGNC:19341] |
| ENSG00000042493 | CAPG | 2006 | 312 | 74 | -2,68 | -2,07 | capping actin protein, gelsolin like [Source:HGNC Symbol;Acc:HGNC:1474] |
| ENSG00000182472 | CAPN12 | 767 | 313 | 98 | -1,29 | -1,67 | calpain 12 [Source:HGNC Symbol;Acc:HGNC:13249] |
| ENSG00000162909 | CAPN2 | 1805 | 646 | 218 | -1,48 | -1,56 | calpain 2 [Source:HGNC Symbol;Acc:HGNC:1479] |
| ENSG00000130940 | CASZ1 | 1010 | 141 | 85 | -2,85 | -0,73 | castor zinc finger 1 [Source:HGNC Symbol;Acc:HGNC:26002] |
| ENSG00000105971 | CAV2 | 403 | 228 | 137 | -0,82 | -0,73 | caveolin 2 [Source:HGNC Symbol;Acc:HGNC:1528] |
| ENSG00000142273 | CBLC | 1076 | 156 | 57 | -2,79 | -1,45 | Cbl proto-oncogene C [Source:HGNC Symbol;Acc:HGNC:15961] |
| ENSG00000128596 | CCDC136 | 877 | 424 | 195 | -1,05 | -1,12 | coiled-coil domain containing 136 [Source:HGNC Symbol;Acc:HGNC:22225] |
| ENSG00000213213 | CCDC183 | 697 | 342 | 91 | -1,03 | -1,9 | coiled-coil domain containing 183 [Source:HGNC Symbol;Acc:HGNC:28236] |
| ENSG00000223745 | CCDC18-AS1 | 675 | 331 | 193 | -1,03 | -0,78 | CCDC18 antisense RNA 1 [Source:HGNC Symbol;Acc:HGNC:52262] |
| ENSG00000215039 | CD27-AS1 | 526 | 128 | 42 | -2,04 | -1,6 | CD27 antisense RNA 1 [Source:HGNC Symbol;Acc:HGNC:43896] |
| ENSG00000196352 | CD55 | 295 | 145 | 26 | -1,02 | -2,49 | CD55 molecule (Cromer blood group) [Source:HGNC Symbol;Acc:HGNC:2665] |
| ENSG00000137101 | CD72 | 263 | 108 | 40 | -1,28 | -1,43 | CD72 molecule [Source:HGNC Symbol;Acc:HGNC:1696] |
| ENSG00000110651 | CD81 | 6418 | 3702 | 1438 | -0,79 | -1,36 | CD81 molecule [Source:HGNC Symbol;Acc:HGNC:1701] |
| ENSG00000085117 | CD82 | 301 | 114 | 29 | -1,41 | -1,97 | CD82 molecule [Source:HGNC Symbol;Acc:HGNC:6210] |
| ENSG00000010278 | CD9 | 421 | 165 | 12 | -1,35 | -3,79 | CD9 molecule [Source:HGNC Symbol;Acc:HGNC:1709] |
| ENSG00000163814 | CDCP1 | 401 | 48 | 4 | -3,05 | -3,61 | CUB domain containing protein 1 [Source:HGNC Symbol;Acc:HGNC:24357] |
| ENSG00000062038 | CDH3 | 5884 | 152 | 24 | -5,28 | -2,66 | cadherin 3 [Source:HGNC Symbol;Acc:HGNC:1762] |
| ENSG00000117266 | CDK18 | 1557 | 613 | 123 | -1,34 | -2,32 | cyclin dependent kinase 18 [Source:HGNC Symbol;Acc:HGNC:8751] |
| ENSG00000105810 | CDK6 | 934 | 144 | 23 | -2,7 | -2,61 | cyclin dependent kinase 6 [Source:HGNC Symbol;Acc:HGNC:1777] |
| ENSG00000110318 | CEP126 | 326 | 101 | 51 | -1,69 | -0,98 | centrosomal protein 126 [Source:HGNC Symbol;Acc:HGNC:29264] |
| ENSG00000167123 | CERCAM | 1348 | 729 | 412 | -0,89 | -0,82 | cerebral endothelial cell adhesion molecule [Source:HGNC Symbol;Acc:HGNC:23723] |
| ENSG00000105792 | CFAP69 | 218 | 58 | 30 | -1,91 | -0,95 | cilia and flagella associated protein 69 [Source:HGNC Symbol;Acc:HGNC:26107] |
| ENSG00000090539 | CHRD | 1400 | 660 | 359 | -1,08 | -0,88 | chordin [Source:HGNC Symbol;Acc:HGNC:1949] |
| ENSG00000182022 | CHST15 | 289 | 25 | 1 | -3,55 | -5,31 | carbohydrate sulfotransferase 15 [Source:HGNC Symbol;Acc:HGNC:18137] |
| ENSG00000186510 | CLCNKA | 336 | 62 | 18 | -2,45 | -1,79 | chloride voltage-gated channel Ka [Source:HGNC Symbol;Acc:HGNC:2026] |
| ENSG00000184908 | CLCNKB | 291 | 77 | 17 | -1,93 | -2,18 | chloride voltage-gated channel Kb [Source:HGNC Symbol;Acc:HGNC:2027] |
| ENSG00000164007 | CLDN19 | 644 | 149 | 40 | -2,11 | -1,9 | claudin 19 [Source:HGNC Symbol;Acc:HGNC:2040] |
| ENSG00000165215 | CLDN3 | 2088 | 185 | 82 | -3,5 | -1,17 | claudin 3 [Source:HGNC Symbol;Acc:HGNC:2045] |
| ENSG00000189143 | CLDN4 | 9844 | 372 | 86 | -4,73 | -2,11 | claudin 4 [Source:HGNC Symbol;Acc:HGNC:2046] |
| ENSG00000181885 | CLDN7 | 3859 | 259 | 12 | -3,9 | -4,37 | claudin 7 [Source:HGNC Symbol;Acc:HGNC:2049] |
| ENSG00000112782 | CLIC5 | 404 | 61 | 17 | -2,73 | -1,79 | chloride intracellular channel 5 [Source:HGNC Symbol;Acc:HGNC:13517] |
| ENSG00000198515 | CNGA1 | 189 | 80 | 38 | -1,24 | -1,09 | cyclic nucleotide gated channel alpha 1 [Source:HGNC Symbol;Acc:HGNC:2148] |
| ENSG00000142675 | CNKSR1 | 1079 | 110 | 23 | -3,29 | -2,28 | connector enhancer of kinase suppressor of Ras 1 [Source:HGNC Symbol;Acc:HGNC:19700] |
| ENSG00000130176 | CNN1 | 600 | 124 | 61 | -2,28 | -1,01 | calponin 1 [Source:HGNC Symbol;Acc:HGNC:2155] |
| ENSG00000060718 | COL11A1 | 3876 | 437 | 59 | -3,15 | -2,87 | collagen type XI alpha 1 chain [Source:HGNC Symbol;Acc:HGNC:2186] |
| ENSG00000164692 | COL1A2 | 25057 | 672 | 49 | -5,22 | -3,79 | collagen type I alpha 2 chain [Source:HGNC Symbol;Acc:HGNC:2198] |
| ENSG00000187498 | COL4A1 | 12714 | 3753 | 931 | -1,76 | -2,01 | collagen type IV alpha 1 chain [Source:HGNC Symbol;Acc:HGNC:2202] |
| ENSG00000134871 | COL4A2 | 23169 | 6423 | 1399 | -1,85 | -2,2 | collagen type IV alpha 2 chain [Source:HGNC Symbol;Acc:HGNC:2203] |
| ENSG00000130635 | COL5A1 | 5659 | 1100 | 160 | -2,36 | -2,78 | collagen type V alpha 1 chain [Source:HGNC Symbol;Acc:HGNC:2209] |
| ENSG00000142173 | COL6A2 | 8253 | 3835 | 952 | -1,11 | -2,01 | collagen type VI alpha 2 chain [Source:HGNC Symbol;Acc:HGNC:2212] |
| ENSG00000139117 | CPNE8 | 196 | 96 | 4 | -1,03 | -4,43 | copine 8 [Source:HGNC Symbol;Acc:HGNC:23498] |
| ENSG00000169169 | CPT1C | 2498 | 764 | 436 | -1,71 | -0,81 | carnitine palmitoyltransferase 1C [Source:HGNC Symbol;Acc:HGNC:18540] |
| ENSG00000163703 | CRELD1 | 3700 | 2021 | 1056 | -0,87 | -0,94 | cysteine rich with EGF like domains 1 [Source:HGNC Symbol;Acc:HGNC:14630] |
| ENSG00000103196 | CRISPLD2 | 332 | 25 | 11 | -3,73 | -1,17 | cysteine rich secretory protein LCCL domain containing 2 [Source:HGNC Symbol;Acc:HGNC:25248] |
| ENSG00000072832 | CRMP1 | 445 | 55 | 14 | -3,03 | -1,93 | collapsin response mediator protein 1 [Source:HGNC Symbol;Acc:HGNC:2365] |
| ENSG00000172346 | CSDC2 | 249 | 33 | 13 | -2,94 | -1,35 | cold shock domain containing C2 [Source:HGNC Symbol;Acc:HGNC:30359] |
| ENSG00000161921 | CXCL16 | 241 | 51 | 7 | -2,24 | -2,96 | C-X-C motif chemokine ligand 16 [Source:HGNC Symbol;Acc:HGNC:16642] |
| ENSG00000166394 | CYB5R2 | 442 | 207 | 44 | -1,09 | -2,22 | cytochrome b5 reductase 2 [Source:HGNC Symbol;Acc:HGNC:24376] |
| ENSG00000135929 | CYP27A1 | 1365 | 530 | 249 | -1,36 | -1,09 | cytochrome P450 family 27 subfamily A member 1 [Source:HGNC Symbol;Acc:HGNC:2605] |
| ENSG00000205795 | CYS1 | 709 | 71 | 13 | -3,31 | -2,5 | cystin 1 [Source:HGNC Symbol;Acc:HGNC:18525] |
| ENSG0000003249 | DBNDD1 | 1162 | 151 | 40 | -2,94 | -1,91 | dysbindin domain containing 1 [Source:HGNC Symbol;Acc:HGNC:28455] |
| ENSG00000163354 | DCST2 | 199 | 70 | 34 | -1,5 | -1,04 | DC-STAMP domain containing 2 [Source:HGNC Symbol;Acc:HGNC:26562] |
| ENSG00000204580 | DDR1 | 9346 | 3699 | 2102 | -1,34 | -0,82 | discoidin domain receptor tyrosine kinase 1 [Source:HGNC Symbol;Acc:HGNC:2730] |
| ENSG00000107984 | DKK1 | 626 | 52 | 9 | -3,59 | -2,53 | dickkopf WNT signaling pathway inhibitor 1 [Source:HGNC Symbol;Acc:HGNC:2891] |
| ENSG00000161249 | DMKN | 3253 | 1178 | 616 | -1,47 | -0,93 | dermokine [Source:HGNC Symbol;Acc:HGNC:25063] |
| ENSG00000184911 | DMRTC1B | 247 | 83 | 23 | -1,58 | -1,84 | DMRT like family C1B [Source:HGNC Symbol;Acc:HGNC:31686] |
| ENSG00000116675 | DNAJC6 | 551 | 298 | 150 | -0,89 | -0,99 | DnaJ heat shock protein family (Hsp40) member C6 [Source:HGNC Symbol;Acc:HGNC:15469] |
| ENSG00000105612 | DNASE2 | 1854 | 994 | 528 | -0,9 | -0,91 | deoxyribonuclease 2, lysosomal [Source:HGNC Symbol;Acc:HGNC:2960] |
| ENSG00000205683 | DPF3 | 248 | 79 | 24 | -1,66 | -1,74 | double PHD fingers 3 [Source:HGNC Symbol;Acc:HGNC:17427] |
| ENSG00000176978 | DPP7 | 2219 | 763 | 208 | -1,54 | -1,87 | dipeptidyl peptidase 7 [Source:HGNC Symbol;Acc:HGNC:14892] |
| ENSG00000177103 | DSCAML1 | 292 | 54 | 24 | -2,44 | -1,19 | DS cell adhesion molecule like 1 [Source:HGNC Symbol;Acc:HGNC:14656] |
| ENSG00000134769 | DTNA | 1120 | 57 | 10 | -4,3 | -2,46 | dystrobrevin alpha [Source:HGNC Symbol;Acc:HGNC:3057] |
| ENSG00000140279 | DUOX2 | 375 | 28 | 3 | -3,74 | -3,24 | dual oxidase 2 [Source:HGNC Symbol;Acc:HGNC:13273] |
| ENSG00000121310 | ECHDC2 | 1578 | 752 | 424 | -1,07 | -0,83 | enoyl-CoA hydratase domain containing 2 [Source:HGNC Symbol;Acc:HGNC:23408] |
| ENSG00000101210 | EEF1A2 | 4794 | 148 | 28 | -5,01 | -2,43 | eukaryotic translation elongation factor 1 alpha 2 [Source:HGNC Symbol;Acc:HGNC:3192] |
| ENSG00000169242 | EFNA1 | 1538 | 585 | 252 | -1,39 | -1,22 | ephrin A1 [Source:HGNC Symbol;Acc:HGNC:3221] |
| ENSG00000143590 | EFNA3 | 221 | 111 | 53 | -1 | -1,07 | ephrin A3 [Source:HGNC Symbol;Acc:HGNC:3223] |
| ENSG00000163435 | ELF3 | 3211 | 796 | 190 | -2,01 | -2,07 | E74 like ETS transcription factor 3 [Source:HGNC Symbol;Acc:HGNC:3318] |
| ENSG00000102034 | ELF4 | 921 | 460 | 206 | -1 | -1,16 | E74 like ETS transcription factor 4 [Source:HGNC Symbol;Acc:HGNC:3319] |
| ENSG00000166897 | ELFN2 | 266 | 19 | 2 | -3,85 | -3,01 | extracellular leucine rich repeat and fibronectin type III domain containing 2 [Source:HGNC Symbol;Acc:HGNC:29396] |
| ENSG00000125746 | EML2 | 1826 | 629 | 334 | -1,54 | -0,91 | echinoderm microtubule associated protein like 2 [Source:HGNC Symbol;Acc:HGNC:18035] |
| ENSG00000112796 | ENPP5 | 466 | 67 | 7 | -2,8 | -3,28 | ectonucleotide pyrophosphatase/phosphodiesterase 5 (putative) [Source:HGNC Symbol;Acc:HGNC:13717] |
| ENSG00000088367 | EPB41L1 | 2668 | 1195 | 553 | -1,16 | -1,11 | erythrocyte membrane protein band 4.1 like 1 [Source:HGNC Symbol;Acc:HGNC:3378] |
| ENSG00000082397 | EPB41L3 | 1425 | 665 | 329 | -1,1 | -1,02 | erythrocyte membrane protein band 4.1 like 3 [Source:HGNC Symbol;Acc:HGNC:3380] |
| ENSG00000129595 | EPB41L4A | 581 | 159 | 50 | -1,87 | -1,67 | erythrocyte membrane protein band 4.1 like 4A [Source:HGNC Symbol;Acc:HGNC:13278] |
| ENSG00000146904 | EPHA1 | 2298 | 321 | 126 | -2,84 | -1,35 | EPH receptor A1 [Source:HGNC Symbol;Acc:HGNC:3385] |
| ENSG00000080224 | EPHA6 | 420 | 86 | 45 | -2,29 | -0,93 | EPH receptor A6 [Source:HGNC Symbol;Acc:HGNC:19296] |
| ENSG00000106123 | EPHB6 | 687 | 43 | 3 | -3,99 | -4 | EPH receptor B6 [Source:HGNC Symbol;Acc:HGNC:3396] |
| ENSG00000105131 | EPHX3 | 260 | 18 | 2 | -3,84 | -3,35 | epoxide hydrolase 3 [Source:HGNC Symbol;Acc:HGNC:23760] |
| ENSG00000261150 | EPPK1 | 2823 | 372 | 129 | -2,92 | -1,53 | epiplakin 1 [Source:HGNC Symbol;Acc:HGNC:15577] |
| ENSG00000131037 | EPS8L1 | 1574 | 79 | 22 | -4,32 | -1,84 | EPS8 like 1 [Source:HGNC Symbol;Acc:HGNC:21295] |
| ENSG00000177106 | EPS8L2 | 6961 | 1826 | 975 | -1,93 | -0,9 | EPS8 like 2 [Source:HGNC Symbol;Acc:HGNC:21296] |
| ENSG00000086619 | ERO1B | 244 | 73 | 18 | -1,74 | -1,99 | endoplasmic reticulum oxidoreductase 1 beta [Source:HGNC Symbol;Acc:HGNC:14355] |
| ENSG00000164283 | ESM1 | 638 | 42 | 1 | -3,94 | -5,94 | endothelial cell specific molecule 1 [Source:HGNC Symbol;Acc:HGNC:3466] |
| ENSG00000187609 | EXD3 | 2011 | 898 | 395 | -1,16 | -1,18 | exonuclease 3'-5' domain containing 3 [Source:HGNC Symbol;Acc:HGNC:26023] |
| ENSG00000158769 | F11R | 6578 | 2155 | 1162 | -1,61 | -0,89 | F11 receptor [Source:HGNC Symbol;Acc:HGNC:14685] |
| ENSG00000164251 | F2RL1 | 1938 | 349 | 45 | -2,47 | -2,95 | F2R like trypsin receptor 1 [Source:HGNC Symbol;Acc:HGNC:3538] |
| ENSG00000177706 | FAM20C | 879 | 177 | 96 | -2,32 | -0,88 | FAM20C, golgi associated secretory pathway kinase [Source:HGNC Symbol;Acc:HGNC:22140] |
| ENSG00000116661 | FBXO2 | 3134 | 260 | 61 | -3,59 | -2,09 | F-box protein 2 [Source:HGNC Symbol;Acc:HGNC:13581] |
| ENSG00000163013 | FBXO41 | 916 | 57 | 18 | -4,02 | -1,66 | F-box protein 41 [Source:HGNC Symbol;Acc:HGNC:29409] |
| ENSG00000132879 | FBXO44 | 1667 | 415 | 219 | -2 | -0,92 | F-box protein 44 [Source:HGNC Symbol;Acc:HGNC:24847] |
| ENSG00000130475 | FCHO1 | 230 | 44 | 13 | -2,38 | -1,78 | FCH domain only 1 [Source:HGNC Symbol;Acc:HGNC:29002] |
| ENSG00000088340 | FER1L4 | 1336 | 629 | 142 | -1,09 | -2,15 | fer-1 like family member 4, pseudogene [Source:HGNC Symbol;Acc:HGNC:15801] |
| ENSG00000101311 | FERMT1 | 310 | 18 | 3 | -4,13 | -2,31 | fermitin family member 1 [Source:HGNC Symbol;Acc:HGNC:15889] |
| ENSG00000182511 | FES | 460 | 212 | 123 | -1,11 | -0,79 | FES proto-oncogene, tyrosine kinase [Source:HGNC Symbol;Acc:HGNC:3657] |
| ENSG00000128591 | FLNC | 5849 | 1538 | 771 | -1,93 | -1 | filamin C [Source:HGNC Symbol;Acc:HGNC:3756] |
| ENSG00000102755 | FLT1 | 896 | 407 | 40 | -1,14 | -3,33 | fms related tyrosine kinase 1 [Source:HGNC Symbol;Acc:HGNC:3763] |
| ENSG00000075426 | FOSL2 | 1720 | 438 | 167 | -1,97 | -1,39 | FOS like 2, AP-1 transcription factor subunit [Source:HGNC Symbol;Acc:HGNC:3798] |
| ENSG00000151474 | FRMD4A | 3835 | 978 | 426 | -1,97 | -1,2 | FERM domain containing 4A [Source:HGNC Symbol;Acc:HGNC:25491] |
| ENSG00000172461 | FUT9 | 666 | 33 | 4 | -4,32 | -3,23 | fucosyltransferase 9 [Source:HGNC Symbol;Acc:HGNC:4020] |
| ENSG00000139112 | GABARAPL1 | 5533 | 2860 | 1513 | -0,95 | -0,92 | GABA type A receptor associated protein like 1 [Source:HGNC Symbol;Acc:HGNC:4068] |
| ENSG00000115339 | GALNT3 | 2330 | 222 | 42 | -3,39 | -2,41 | polypeptide N-acetylgalactosaminyltransferase 3 [Source:HGNC Symbol;Acc:HGNC:4125] |
| ENSG00000139629 | GALNT6 | 386 | 37 | 2 | -3,39 | -3,98 | polypeptide N-acetylgalactosaminyltransferase 6 [Source:HGNC Symbol;Acc:HGNC:4128] |
| ENSG00000141013 | GAS8 | 337 | 19 | 2 | -4,12 | -3,56 | growth arrest specific 8 [Source:HGNC Symbol;Acc:HGNC:4166] |
| ENSG00000130700 | GATA5 | 374 | 130 | 26 | -1,52 | -2,33 | GATA binding protein 5 [Source:HGNC Symbol;Acc:HGNC:15802] |
| ENSG00000266010 | GATA6-AS1 | 1696 | 731 | 244 | -1,21 | -1,58 | GATA6 antisense RNA 1 (head to head) [Source:HGNC Symbol;Acc:HGNC:48840] |
| ENSG00000171766 | GATM | 2257 | 897 | 429 | -1,33 | -1,06 | glycine amidinotransferase [Source:HGNC Symbol;Acc:HGNC:4175] |
| ENSG00000148288 | GBGT1 | 243 | 58 | 15 | -2,07 | -2,02 | globoside alpha-1,3-N-acetylgalactosaminyltransferase 1 [Source:HGNC Symbol;Acc:HGNC:20460] |
| ENSG00000119125 | GDA | 483 | 103 | 15 | -2,23 | -2,8 | guanine deaminase [Source:HGNC Symbol;Acc:HGNC:4212] |
| ENSG00000137960 | GIPC2 | 280 | 52 | 14 | -2,44 | -1,85 | GIPC PDZ domain containing family member 2 [Source:HGNC Symbol;Acc:HGNC:18177] |
| ENSG00000149328 | GLB1L2 | 558 | 69 | 5 | -3,02 | -3,8 | galactosidase beta 1 like 2 [Source:HGNC Symbol;Acc:HGNC:25129] |
| ENSG00000076716 | GPC4 | 1375 | 173 | 74 | -2,99 | -1,23 | glypican 4 [Source:HGNC Symbol;Acc:HGNC:4452] |
| ENSG00000077585 | GPR137B | 238 | 40 | 3 | -2,57 | -3,63 | G protein-coupled receptor 137B [Source:HGNC Symbol;Acc:HGNC:11862] |
| ENSG00000180758 | GPR157 | 468 | 275 | 165 | -0,77 | -0,74 | G protein-coupled receptor 157 [Source:HGNC Symbol;Acc:HGNC:23687] |
| ENSG00000013588 | GPRC5A | 1483 | 265 | 24 | -2,49 | -3,46 | G protein-coupled receptor class C group 5 member A [Source:HGNC Symbol;Acc:HGNC:9836] |
| ENSG00000178075 | GRAMD1C | 411 | 246 | 122 | -0,74 | -1,01 | GRAM domain containing 1C [Source:HGNC Symbol;Acc:HGNC:25252] |
| ENSG00000141738 | GRB7 | 2867 | 405 | 52 | -2,82 | -2,95 | growth factor receptor bound protein 7 [Source:HGNC Symbol;Acc:HGNC:4567] |
| ENSG00000105737 | GRIK5 | 922 | 325 | 25 | -1,51 | -3,69 | glutamate ionotropic receptor kainate type subunit 5 [Source:HGNC Symbol;Acc:HGNC:4583] |
| ENSG00000116032 | GRIN3B | 394 | 145 | 28 | -1,44 | -2,38 | glutamate ionotropic receptor NMDA type subunit 3B [Source:HGNC Symbol;Acc:HGNC:16768] |
| ENSG00000073605 | GSDMB | 2770 | 993 | 489 | -1,48 | -1,02 | gasdermin B [Source:HGNC Symbol;Acc:HGNC:23690] |
| ENSG00000134184 | GSTM1 | 676 | 329 | 47 | -1,04 | -2,81 | glutathione S-transferase mu 1 [Source:HGNC Symbol;Acc:HGNC:4632] |
| ENSG00000213366 | GSTM2 | 2312 | 1211 | 651 | -0,93 | -0,9 | glutathione S-transferase mu 2 [Source:HGNC Symbol;Acc:HGNC:4634] |
| ENSG00000134202 | GSTM3 | 3533 | 1848 | 1052 | -0,93 | -0,81 | glutathione S-transferase mu 3 [Source:HGNC Symbol;Acc:HGNC:4635] |
| ENSG00000065621 | GSTO2 | 249 | 76 | 43 | -1,71 | -0,82 | glutathione S-transferase omega 2 [Source:HGNC Symbol;Acc:HGNC:23064] |
| ENSG00000112599 | GUCA1B | 187 | 61 | 34 | -1,62 | -0,87 | guanylate cyclase activator 1B [Source:HGNC Symbol;Acc:HGNC:4679] |
| ENSG00000061918 | GUCY1B3 | 311 | 102 | 44 | -1,61 | -1,22 | guanylate cyclase 1 soluble subunit beta [Source:HGNC Symbol;Acc:HGNC:4687] |
| ENSG00000228223 | HCG11 | 447 | 245 | 69 | -0,87 | -1,82 | HLA complex group 11 (non-protein coding) [Source:HGNC Symbol;Acc:HGNC:17707] |
| ENSG00000127311 | HELB | 389 | 223 | 129 | -0,8 | -0,79 | DNA helicase B [Source:HGNC Symbol;Acc:HGNC:17196] |
| ENSG00000114315 | HES1 | 3883 | 2173 | 1207 | -0,84 | -0,85 | hes family bHLH transcription factor 1 [Source:HGNC Symbol;Acc:HGNC:5192] |
| ENSG00000164683 | HEY1 | 1771 | 726 | 258 | -1,29 | -1,49 | hes related family bHLH transcription factor with YRPW motif 1 [Source:HGNC Symbol;Acc:HGNC:4880] |
| ENSG00000206503 | HLA-A | 5998 | 3183 | 1426 | -0,91 | -1,16 | major histocompatibility complex, class I, A [Source:HGNC Symbol;Acc:HGNC:4931] |
| ENSG00000234745 | HLA-B | 2976 | 175 | 19 | -4,09 | -3,16 | major histocompatibility complex, class I, B [Source:HGNC Symbol;Acc:HGNC:4932] |
| ENSG00000204525 | HLA-C | 8688 | 2422 | 1428 | -1,84 | -0,76 | major histocompatibility complex, class I, C [Source:HGNC Symbol;Acc:HGNC:4933] |
| ENSG00000223865 | HLA-DPB1 | 333 | 5 | 0 | -6 | -4,83 | major histocompatibility complex, class II, DP beta 1 [Source:HGNC Symbol;Acc:HGNC:4940] |
| ENSG00000179344 | HLA-DQB1 | 539 | 9 | 0 | -5,94 | -4,63 | major histocompatibility complex, class II, DQ beta 1 [Source:HGNC Symbol;Acc:HGNC:4944] |
| ENSG00000204592 | HLA-E | 3886 | 2230 | 949 | -0,8 | -1,23 | major histocompatibility complex, class I, E [Source:HGNC Symbol;Acc:HGNC:4962] |
| ENSG00000206341 | HLA-H | 1531 | 727 | 343 | -1,07 | -1,08 | major histocompatibility complex, class I, H (pseudogene) [Source:HGNC Symbol;Acc:HGNC:4965] |
| ENSG00000105707 | HPN | 9510 | 4807 | 1521 | -0,98 | -1,66 | hepsin [Source:HGNC Symbol;Acc:HGNC:5155] |
| ENSG00000171004 | HS6ST2 | 1321 | 297 | 151 | -2,15 | -0,97 | heparan sulfate 6-O-sulfotransferase 2 [Source:HGNC Symbol;Acc:HGNC:19133] |
| ENSG00000087076 | HSD17B14 | 462 | 250 | 145 | -0,88 | -0,79 | hydroxysteroid 17-beta dehydrogenase 14 [Source:HGNC Symbol;Acc:HGNC:23238] |
| ENSG00000126803 | HSPA2 | 196 | 117 | 60 | -0,74 | -0,96 | heat shock protein family A (Hsp70) member 2 [Source:HGNC Symbol;Acc:HGNC:5235] |
| ENSG00000174498 | IGDCC3 | 6784 | 2772 | 1128 | -1,29 | -1,3 | immunoglobulin superfamily DCC subclass member 3 [Source:HGNC Symbol;Acc:HGNC:9700] |
| ENSG00000115461 | IGFBP5 | 18662 | 3652 | 727 | -2,35 | -2,33 | insulin like growth factor binding protein 5 [Source:HGNC Symbol;Acc:HGNC:5474] |
| ENSG00000147255 | IGSF1 | 1083 | 594 | 343 | -0,87 | -0,79 | immunoglobulin superfamily member 1 [Source:HGNC Symbol;Acc:HGNC:5948] |
| ENSG00000198825 | INPP5F | 4521 | 2310 | 1322 | -0,97 | -0,8 | inositol polyphosphate-5-phosphatase F [Source:HGNC Symbol;Acc:HGNC:17054] |
| ENSG00000117595 | IRF6 | 1460 | 107 | 4 | -3,77 | -4,81 | interferon regulatory factor 6 [Source:HGNC Symbol;Acc:HGNC:6121] |
| ENSG00000005884 | ITGA3 | 4100 | 920 | 303 | -2,16 | -1,6 | integrin subunit alpha 3 [Source:HGNC Symbol;Acc:HGNC:6139] |
| ENSG00000132470 | ITGB4 | 966 | 250 | 49 | -1,95 | -2,37 | integrin subunit beta 4 [Source:HGNC Symbol;Acc:HGNC:6158] |
| ENSG00000115221 | ITGB6 | 1838 | 73 | 9 | -4,66 | -3,01 | integrin subunit beta 6 [Source:HGNC Symbol;Acc:HGNC:6161] |
| ENSG00000105639 | JAK3 | 344 | 65 | 4 | -2,41 | -3,87 | Janus kinase 3 [Source:HGNC Symbol;Acc:HGNC:6193] |
| ENSG00000154118 | JPH3 | 1047 | 59 | 5 | -4,15 | -3,57 | junctophilin 3 [Source:HGNC Symbol;Acc:HGNC:14203] |
| ENSG00000183960 | KCNH8 | 490 | 27 | 7 | -4,2 | -1,97 | potassium voltage-gated channel subfamily H member 8 [Source:HGNC Symbol;Acc:HGNC:18864] |
| ENSG00000135750 | KCNK1 | 447 | 6 | 0 | -6,26 | -4,04 | potassium two pore domain channel subfamily K member 1 [Source:HGNC Symbol;Acc:HGNC:6272] |
| ENSG00000099337 | KCNK6 | 629 | 116 | 20 | -2,44 | -2,57 | potassium two pore domain channel subfamily K member 6 [Source:HGNC Symbol;Acc:HGNC:6281] |
| ENSG00000178695 | KCTD12 | 678 | 244 | 81 | -1,48 | -1,59 | potassium channel tetramerization domain containing 12 [Source:HGNC Symbol;Acc:HGNC:14678] |
| ENSG00000175707 | KDF1 | 660 | 111 | 16 | -2,57 | -2,83 | keratinocyte differentiation factor 1 [Source:HGNC Symbol;Acc:HGNC:26624] |
| ENSG00000112232 | KHDRBS2 | 308 | 124 | 59 | -1,31 | -1,07 | KH RNA binding domain containing, signal transduction associated 2 [Source:HGNC Symbol;Acc:HGNC:18114] |
| ENSG00000157404 | KIT | 452 | 153 | 48 | -1,57 | -1,65 | KIT proto-oncogene receptor tyrosine kinase [Source:HGNC Symbol;Acc:HGNC:6342] |
| ENSG00000102349 | KLF8 | 243 | 39 | 4 | -2,64 | -3,31 | Kruppel like factor 8 [Source:HGNC Symbol;Acc:HGNC:6351] |
| ENSG00000171798 | KNDC1 | 545 | 41 | 3 | -3,74 | -3,64 | kinase non-catalytic C-lobe domain containing 1 [Source:HGNC Symbol;Acc:HGNC:29374] |
| ENSG00000159166 | LAD1 | 1905 | 408 | 156 | -2,22 | -1,39 | ladinin 1 [Source:HGNC Symbol;Acc:HGNC:6472] |
| ENSG00000172037 | LAMB2 | 15830 | 7238 | 4106 | -1,13 | -0,82 | laminin subunit beta 2 [Source:HGNC Symbol;Acc:HGNC:6487] |
| ENSG00000204381 | LAYN | 575 | 112 | 67 | -2,37 | -0,74 | layilin [Source:HGNC Symbol;Acc:HGNC:29471] |
| ENSG00000186007 | LEMD1 | 596 | 80 | 9 | -2,9 | -3,21 | LEM domain containing 1 [Source:HGNC Symbol;Acc:HGNC:18725] |

| | | | | | | | |
|---|---|---|---|---|---|---|---|
| ENSG00000116977 | LGALS8 | 1889 | 800 | 309 | -1,24 | -1,37 | galectin 8 [Source:HGNC Symbol;Acc:HGNC:6569] |
| ENSG00000153012 | LGI2 | 321 | 20 | 7 | -3,99 | -1,51 | leucine rich repeat LGI family member 2 [Source:HGNC Symbol;Acc:HGNC:18710] |
| ENSG00000064042 | LIMCH1 | 1765 | 812 | 433 | -1,12 | -0,91 | LIM and calponin homology domains 1 [Source:HGNC Symbol;Acc:HGNC:29191] |
| ENSG00000072163 | LIMS2 | 238 | 112 | 59 | -1,09 | -0,92 | LIM zinc finger domain containing 2 [Source:HGNC Symbol;Acc:HGNC:16084] |
| ENSG00000251533 | LINC00605 | 311 | 70 | 8 | -2,16 | -3,21 | long intergenic non-protein coding RNA 605 [Source:HGNC Symbol;Acc:HGNC:43928] |
| ENSG00000245060 | LINC00847 | 248 | 126 | 53 | -0,98 | -1,27 | long intergenic non-protein coding RNA 847 [Source:HGNC Symbol;Acc:HGNC:45050] |
| ENSG00000281205 | LINC00950 | 356 | 155 | 82 | -1,2 | -0,92 | long intergenic non-protein coding RNA 950 [Source:HGNC Symbol;Acc:HGNC:28098] |
| ENSG00000204054 | LINC00963 | 1175 | 666 | 377 | -0,82 | -0,82 | long intergenic non-protein coding RNA 963 [Source:HGNC Symbol;Acc:HGNC:48716] |
| ENSG00000163898 | LIPH | 395 | 155 | 24 | -1,35 | -2,69 | lipase H [Source:HGNC Symbol;Acc:HGNC:18483] |
| ENSG00000142235 | LMTK3 | 978 | 343 | 140 | -1,51 | -1,29 | lemur tyrosine kinase 3 [Source:HGNC Symbol;Acc:HGNC:19295] |
| ENSG00000072201 | LNX1 | 1006 | 349 | 43 | -1,53 | -3 | ligand of numb-protein X 1 [Source:HGNC Symbol;Acc:HGNC:6657] |
| ENSG00000129038 | LOXL1 | 720 | 345 | 100 | -1,06 | -1,78 | lysyl oxidase like 1 [Source:HGNC Symbol;Acc:HGNC:6665] |
| ENSG00000261801 | LOXL1-AS1 | 235 | 90 | 18 | -1,38 | -2,32 | LOXL1 antisense RNA 1 [Source:HGNC Symbol;Acc:HGNC:44169] |
| ENSG00000138131 | LOXL4 | 3161 | 169 | 61 | -4,23 | -1,45 | lysyl oxidase like 4 [Source:HGNC Symbol;Acc:HGNC:17171] |
| ENSG00000087253 | LPCAT2 | 264 | 71 | 22 | -1,89 | -1,65 | lysophosphatidylcholine acyltransferase 2 [Source:HGNC Symbol;Acc:HGNC:26032] |
| ENSG00000010626 | LRRC23 | 302 | 177 | 101 | -0,77 | -0,81 | leucine rich repeat containing 23 [Source:HGNC Symbol;Acc:HGNC:19138] |
| ENSG00000150551 | LYPD1 | 591 | 21 | 5 | -4,78 | -2,02 | LY6/PLAUR domain containing 1 [Source:HGNC Symbol;Acc:HGNC:28431] |
| ENSG00000183742 | MACC1 | 987 | 108 | 43 | -3,18 | -1,35 | MACC1, MET transcriptional regulator [Source:HGNC Symbol;Acc:HGNC:30215] |
| ENSG00000147676 | MAL2 | 3936 | 1179 | 240 | -1,74 | -2,3 | mal, T-cell differentiation protein 2 (gene/pseudogene) [Source:HGNC Symbol;Acc:HGNC:13634] |
| ENSG00000101460 | MAP1LC3A | 401 | 180 | 92 | -1,16 | -0,96 | microtubule associated protein 1 light chain 3 alpha [Source:HGNC Symbol;Acc:HGNC:6838] |
| ENSG00000267278 | MAP3K14-AS1 | 244 | 42 | 16 | -2,53 | -1,37 | MAP3K14 antisense RNA 1 [Source:HGNC Symbol;Acc:HGNC:44359] |
| ENSG00000107968 | MAP3K8 | 614 | 134 | 55 | -2,2 | -1,29 | mitogen-activated protein kinase kinase kinase 8 [Source:HGNC Symbol;Acc:HGNC:6860] |
| ENSG00000156711 | MAPK13 | 3188 | 730 | 122 | -2,13 | -2,58 | mitogen-activated protein kinase 13 [Source:HGNC Symbol;Acc:HGNC:6875] |
| ENSG00000181085 | MAPK15 | 941 | 138 | 47 | -2,77 | -1,54 | mitogen-activated protein kinase 15 [Source:HGNC Symbol;Acc:HGNC:24667] |
| ENSG00000152939 | MARVELD2 | 1809 | 750 | 413 | -1,27 | -0,86 | MARVEL domain containing 2 [Source:HGNC Symbol;Acc:HGNC:26401] |
| ENSG00000055732 | MCOLN3 | 1329 | 166 | 9 | -3 | -4,27 | mucolipin 3 [Source:HGNC Symbol;Acc:HGNC:13358] |
| ENSG00000105419 | MEIS3 | 1594 | 77 | 18 | -4,38 | -2,12 | Meis homeobox 3 [Source:HGNC Symbol;Acc:HGNC:29537] |
| ENSG00000163975 | MELTF | 1384 | 722 | 264 | -0,94 | -1,45 | melanotransferrin [Source:HGNC Symbol;Acc:HGNC:7037] |
| ENSG00000170430 | MGMT | 479 | 83 | 5 | -2,53 | -3,91 | O-6-methylguanine-DNA methyltransferase [Source:HGNC Symbol;Acc:HGNC:7059] |
| ENSG00000141854 | MISP3 | 252 | 105 | 51 | -1,27 | -1,07 | MISP family member 3 [Source:HGNC Symbol;Acc:HGNC:26963] |
| ENSG00000196611 | MMP1 | 933 | 135 | 9 | -2,79 | -3,95 | matrix metallopeptidase 1 [Source:HGNC Symbol;Acc:HGNC:7155] |
| ENSG00000126005 | MMP24-AS1 | 1340 | 725 | 358 | -0,89 | -1,01 | MMP24 antisense RNA 1 [Source:HGNC Symbol;Acc:HGNC:44421] |
| ENSG00000079931 | MOXD1 | 1240 | 519 | 237 | -1,26 | -1,13 | monooxygenase DBH like 1 [Source:HGNC Symbol;Acc:HGNC:21063] |
| ENSG00000011028 | MRC2 | 8939 | 4350 | 1301 | -1,04 | -1,74 | mannose receptor C type 2 [Source:HGNC Symbol;Acc:HGNC:16875] |
| ENSG00000101825 | MXRA5 | 212 | 76 | 12 | -1,47 | -2,65 | matrix remodeling associated 5 [Source:HGNC Symbol;Acc:HGNC:7539] |
| ENSG00000162576 | MXRA8 | 788 | 193 | 34 | -2,03 | -2,52 | matrix remodeling associated 8 [Source:HGNC Symbol;Acc:HGNC:7542] |
| ENSG00000101335 | MYL9 | 2272 | 584 | 337 | -1,96 | -0,79 | myosin light chain 9 [Source:HGNC Symbol;Acc:HGNC:15754] |
| ENSG00000266714 | MYO15B | 3254 | 647 | 243 | -2,33 | -1,41 | myosin XVB [Source:HGNC Symbol;Acc:HGNC:14083] |
| ENSG00000136274 | NACAD | 583 | 282 | 138 | -1,05 | -1,04 | NAC alpha domain containing [Source:HGNC Symbol;Acc:HGNC:22196] |
| ENSG00000237886 | NALT1 | 225 | 121 | 61 | -0,89 | -1 | NOTCH1 associated lncRNA in T-cell acute lymphoblastic leukemia 1 [Source:HGNC Symbol;Acc:HGNC:51192] |
| ENSG00000184454 | NCMAP | 276 | 49 | 13 | -2,49 | -1,95 | non-compact myelin associated protein [Source:HGNC Symbol;Acc:HGNC:29332] |
| ENSG00000078114 | NEBL | 668 | 373 | 121 | -0,84 | -1,63 | nebulette [Source:HGNC Symbol;Acc:HGNC:16932] |
| ENSG00000100968 | NFATC4 | 3707 | 1519 | 700 | -1,29 | -1,12 | nuclear factor of activated T-cells 4 [Source:HGNC Symbol;Acc:HGNC:7778] |
| ENSG00000103024 | NME3 | 773 | 400 | 136 | -0,95 | -1,55 | NME/NM23 nucleoside diphosphate kinase 3 [Source:HGNC Symbol;Acc:HGNC:7851] |
| ENSG00000104967 | NOVA2 | 203 | 85 | 5 | -1,25 | -3,95 | NOVA alternative splicing regulator 2 [Source:HGNC Symbol;Acc:HGNC:7887] |
| ENSG00000188747 | NOXA1 | 658 | 167 | 62 | -1,98 | -1,42 | NADPH oxidase activator 1 [Source:HGNC Symbol;Acc:HGNC:10668] |
| ENSG00000107281 | NPDC1 | 2877 | 847 | 287 | -1,76 | -1,56 | neural proliferation, differentiation and control 1 [Source:HGNC Symbol;Acc:HGNC:7899] |
| ENSG00000215440 | NPEPL1 | 1283 | 485 | 291 | -1,4 | -0,73 | aminopeptidase-like 1 [Source:HGNC Symbol;Acc:HGNC:16244] |
| ENSG00000168743 | NPNT | 906 | 21 | 8 | -5,45 | -1,28 | nephronectin [Source:HGNC Symbol;Acc:HGNC:27405] |
| ENSG00000169418 | NPR1 | 222 | 94 | 6 | -1,25 | -3,87 | natriuretic peptide receptor 1 [Source:HGNC Symbol;Acc:HGNC:7943] |
| ENSG00000118257 | NRP2 | 1244 | 327 | 64 | -1,93 | -2,36 | neuropilin 2 [Source:HGNC Symbol;Acc:HGNC:8005] |
| ENSG00000246082 | NUDT16P1 | 248 | 86 | 17 | -1,52 | -2,33 | nudix hydrolase 16 pseudogene 1 [Source:HGNC Symbol;Acc:HGNC:27189] |
| ENSG00000166924 | NYAP1 | 423 | 242 | 71 | -0,81 | -1,79 | neuronal tyrosine phosphorylated phosphoinositide-3-kinase adaptor 1 [Source:HGNC Symbol;Acc:HGNC:22009] |
| ENSG00000205978 | NYNRIN | 4514 | 2426 | 1345 | -0,9 | -0,85 | NYN domain and retroviral integrase containing [Source:HGNC Symbol;Acc:HGNC:20165] |
| ENSG00000177989 | ODF3B | 225 | 81 | 8 | -1,48 | -3,28 | outer dense fiber of sperm tails 3B [Source:HGNC Symbol;Acc:HGNC:34388] |
| ENSG00000197444 | OGDHL | 1107 | 209 | 71 | -2,41 | -1,56 | oxoglutarate dehydrogenase-like [Source:HGNC Symbol;Acc:HGNC:25590] |
| ENSG00000105088 | OLFM2 | 2643 | 828 | 261 | -1,67 | -1,66 | olfactomedin 2 [Source:HGNC Symbol;Acc:HGNC:17189] |
| ENSG00000185585 | OLFML2A | 692 | 115 | 67 | -2,59 | -0,8 | olfactomedin like 2A [Source:HGNC Symbol;Acc:HGNC:27270] |
| ENSG00000125510 | OPRL1 | 252 | 88 | 46 | -1,51 | -0,94 | opioid related nociceptin receptor 1 [Source:HGNC Symbol;Acc:HGNC:8155] |
| ENSG00000085465 | OVGP1 | 602 | 217 | 118 | -1,47 | -0,88 | oviductal glycoprotein 1 [Source:HGNC Symbol;Acc:HGNC:8524] |
| ENSG00000172818 | OVOL1 | 557 | 80 | 17 | -2,8 | -2,23 | ovo like transcriptional repressor 1 [Source:HGNC Symbol;Acc:HGNC:8525] |
| ENSG00000072682 | P4HA2 | 3054 | 1698 | 759 | -0,85 | -1,16 | prolyl 4-hydroxylase subunit alpha 2 [Source:HGNC Symbol;Acc:HGNC:8547] |
| ENSG00000178467 | P4HTM | 1696 | 467 | 55 | -1,86 | -3,08 | prolyl 4-hydroxylase, transmembrane [Source:HGNC Symbol;Acc:HGNC:28858] |
| ENSG00000171759 | PAH | 1554 | 80 | 21 | -4,27 | -1,91 | phenylalanine hydroxylase [Source:HGNC Symbol;Acc:HGNC:8582] |
| ENSG00000149269 | PAK1 | 4809 | 2361 | 1336 | -1,03 | -0,82 | p21 (RAC1) activated kinase 1 [Source:HGNC Symbol;Acc:HGNC:8590] |
| ENSG00000116117 | PARD3B | 1029 | 527 | 287 | -0,96 | -0,88 | par-3 family cell polarity regulator beta [Source:HGNC Symbol;Acc:HGNC:14446] |
| ENSG00000169851 | PCDH7 | 990 | 381 | 106 | -1,38 | -1,84 | protocadherin 7 [Source:HGNC Symbol;Acc:HGNC:8659] |
| ENSG00000250120 | PCDHA10 | 310 | 187 | 59 | -0,73 | -1,66 | protocadherin alpha 10 [Source:HGNC Symbol;Acc:HGNC:8664] |
| ENSG00000102109 | PCSK1N | 937 | 190 | 76 | -2,3 | -1,33 | proprotein convertase subtilisin/kexin type 1 inhibitor [Source:HGNC Symbol;Acc:HGNC:17301] |
| ENSG00000065989 | PDE4A | 208 | 121 | 62 | -0,78 | -0,96 | phosphodiesterase 4A [Source:HGNC Symbol;Acc:HGNC:8780] |
| ENSG00000197461 | PDGFA | 3143 | 543 | 106 | -2,53 | -2,35 | platelet derived growth factor subunit A [Source:HGNC Symbol;Acc:HGNC:8799] |
| ENSG00000100311 | PDGFB | 948 | 189 | 64 | -2,33 | -1,55 | platelet derived growth factor subunit B [Source:HGNC Symbol;Acc:HGNC:8800] |
| ENSG00000120913 | PDLIM2 | 378 | 161 | 38 | -1,24 | -2,09 | PDZ and LIM domain 2 [Source:HGNC Symbol;Acc:HGNC:13992] |
| ENSG00000067057 | PFKP | 2195 | 823 | 174 | -1,41 | -2,24 | phosphofructokinase, platelet [Source:HGNC Symbol;Acc:HGNC:8878] |
| ENSG00000103335 | PIEZO1 | 6749 | 1200 | 591 | -2,49 | -1,02 | piezo type mechanosensitive ion channel component 1 [Source:HGNC Symbol;Acc:HGNC:28993] |
| ENSG00000100100 | PIK3IP1 | 595 | 291 | 127 | -1,03 | -1,19 | phosphoinositide-3-kinase interacting protein 1 [Source:HGNC Symbol;Acc:HGNC:24942] |
| ENSG00000176485 | PLA2G16 | 1090 | 483 | 277 | -1,17 | -0,8 | phospholipase A2 group XVI [Source:HGNC Symbol;Acc:HGNC:17825] |
| ENSG00000153246 | PLA2R1 | 486 | 290 | 58 | -0,75 | -2,32 | phospholipase A2 receptor 1 [Source:HGNC Symbol;Acc:HGNC:9042] |
| ENSG00000145287 | PLAC8 | 265 | 56 | 17 | -2,23 | -1,77 | placenta specific 8 [Source:HGNC Symbol;Acc:HGNC:19254] |
| ENSG00000122861 | PLAU | 1263 | 604 | 253 | -1,06 | -1,26 | plasminogen activator, urokinase [Source:HGNC Symbol;Acc:HGNC:9052] |
| ENSG00000115896 | PLCL1 | 1424 | 80 | 16 | -4,15 | -2,27 | phospholipase C like 1 [Source:HGNC Symbol;Acc:HGNC:9063] |
| ENSG00000008323 | PLEKHG6 | 1631 | 500 | 178 | -1,7 | -1,49 | pleckstrin homology and RhoGEF domain containing G6 [Source:HGNC Symbol;Acc:HGNC:25562] |
| ENSG00000141934 | PLPP2 | 1158 | 47 | 5 | -4,64 | -3,27 | phospholipid phosphatase 2 [Source:HGNC Symbol;Acc:HGNC:9230] |
| ENSG00000129951 | PLPPR3 | 3497 | 1828 | 1001 | -0,94 | -0,87 | phospholipid phosphatase related 3 [Source:HGNC Symbol;Acc:HGNC:23497] |
| ENSG00000114698 | PLSCR4 | 432 | 150 | 79 | -1,52 | -0,93 | phospholipid scramblase 4 [Source:HGNC Symbol;Acc:HGNC:16497] |
| ENSG00000185664 | PMEL | 334 | 88 | 52 | -1,93 | -0,76 | premelanosome protein [Source:HGNC Symbol;Acc:HGNC:10880] |
| ENSG00000183837 | PNMA3 | 665 | 223 | 72 | -1,57 | -1,65 | paraneoplastic Ma antigen 3 [Source:HGNC Symbol;Acc:HGNC:18742] |
| ENSG00000100344 | PNPLA3 | 290 | 86 | 48 | -1,75 | -0,84 | patatin like phospholipase domain containing 3 [Source:HGNC Symbol;Acc:HGNC:18590] |
| ENSG00000143847 | PPFIA4 | 1222 | 710 | 277 | -0,78 | -1,36 | PTPRF interacting protein alpha 4 [Source:HGNC Symbol;Acc:HGNC:9248] |
| ENSG00000135447 | PPP1R1A | 637 | 69 | 30 | -3,2 | -1,23 | protein phosphatase 1 regulatory inhibitor subunit 1A [Source:HGNC Symbol;Acc:HGNC:9286] |
| ENSG00000131771 | PPP1R1B | 252 | 105 | 21 | -1,27 | -2,34 | protein phosphatase 1 regulatory inhibitor subunit 1B [Source:HGNC Symbol;Acc:HGNC:9287] |
| ENSG00000162148 | PPP1R32 | 505 | 132 | 51 | -1,94 | -1,38 | protein phosphatase 1 regulatory subunit 32 [Source:HGNC Symbol;Acc:HGNC:28869] |
| ENSG00000173281 | PPP1R3B | 2188 | 896 | 401 | -1,29 | -1,16 | protein phosphatase 1 regulatory subunit 3B [Source:HGNC Symbol;Acc:HGNC:14942] |
| ENSG00000119938 | PPP1R3C | 1200 | 626 | 300 | -0,94 | -1,06 | protein phosphatase 1 regulatory subunit 3C [Source:HGNC Symbol;Acc:HGNC:9293] |
| ENSG00000027075 | PRKCH | 230 | 40 | 18 | -2,51 | -1,22 | protein kinase C eta [Source:HGNC Symbol;Acc:HGNC:9403] |
| ENSG00000171867 | PRNP | 1214 | 139 | 11 | -3,13 | -3,72 | prion protein [Source:HGNC Symbol;Acc:HGNC:9449] |
| ENSG00000228672 | PROB1 | 416 | 177 | 53 | -1,23 | -1,77 | proline rich basic protein 1 [Source:HGNC Symbol;Acc:HGNC:41906] |
| ENSG00000230461 | PROX1-AS1 | 724 | 406 | 130 | -0,84 | -1,64 | PROX1 antisense RNA 1 [Source:HGNC Symbol;Acc:HGNC:43656] |
| ENSG00000164099 | PRSS12 | 620 | 167 | 30 | -1,89 | -2,47 | protease, serine 12 [Source:HGNC Symbol;Acc:HGNC:9477] |
| ENSG00000112812 | PRSS16 | 798 | 249 | 136 | -1,68 | -0,87 | protease, serine 16 [Source:HGNC Symbol;Acc:HGNC:9480] |
| ENSG00000005001 | PRSS22 | 2586 | 42 | 5 | -5,93 | -3,11 | protease, serine 22 [Source:HGNC Symbol;Acc:HGNC:14368] |
| ENSG00000151006 | PRSS53 | 234 | 125 | 68 | -0,91 | -0,88 | protease, serine 53 [Source:HGNC Symbol;Acc:HGNC:34407] |
| ENSG00000052344 | PRSS8 | 5851 | 401 | 53 | -3,87 | -2,93 | protease, serine 8 [Source:HGNC Symbol;Acc:HGNC:9491] |
| ENSG00000059915 | PSD | 224 | 90 | 29 | -1,32 | -1,63 | pleckstrin and Sec7 domain containing [Source:HGNC Symbol;Acc:HGNC:9507] |
| ENSG00000146005 | PSD2 | 517 | 114 | 14 | -2,18 | -3,07 | pleckstrin and Sec7 domain containing 2 [Source:HGNC Symbol;Acc:HGNC:19092] |
| ENSG00000124212 | PTGIS | 2186 | 501 | 241 | -2,13 | -1,05 | prostaglandin I2 synthase [Source:HGNC Symbol;Acc:HGNC:9603] |
| ENSG00000110786 | PTPN5 | 248 | 7 | 1 | -5,25 | -3,3 | protein tyrosine phosphatase, non-receptor type 5 [Source:HGNC Symbol;Acc:HGNC:9657] |
| ENSG00000080031 | PTPRH | 241 | 70 | 24 | -1,79 | -1,54 | protein tyrosine phosphatase, receptor type H [Source:HGNC Symbol;Acc:HGNC:9672] |
| ENSG00000168994 | PXDC1 | 560 | 197 | 91 | -1,51 | -1,13 | PX domain containing 1 [Source:HGNC Symbol;Acc:HGNC:21361] |
| ENSG00000115828 | QPCT | 724 | 268 | 121 | -1,44 | -1,15 | glutaminyl-peptide cyclotransferase [Source:HGNC Symbol;Acc:HGNC:9753] |
| ENSG00000139832 | RAB20 | 262 | 114 | 43 | -1,2 | -1,42 | RAB20, member RAS oncogene family [Source:HGNC Symbol;Acc:HGNC:18260] |
| ENSG00000132698 | RAB25 | 3480 | 103 | 11 | -5,08 | -3,3 | RAB25, member RAS oncogene family [Source:HGNC Symbol;Acc:HGNC:18238] |
| ENSG00000154917 | RAB6B | 1290 | 277 | 126 | -2,22 | -1,15 | RAB6B, member RAS oncogene family [Source:HGNC Symbol;Acc:HGNC:14902] |
| ENSG00000128340 | RAC2 | 362 | 147 | 59 | -1,3 | -1,33 | ras-related C3 botulinum toxin substrate 2 (rho family, small GTP binding protein Rac2) [Source:HGNC Symbol;Acc:HGNC:9802] |
| ENSG00000131831 | RAI2 | 309 | 59 | 9 | -2,4 | -2,73 | retinoic acid induced 2 [Source:HGNC Symbol;Acc:HGNC:9835] |
| ENSG00000079337 | RAPGEF3 | 223 | 110 | 29 | -1,03 | -1,94 | Rap guanine nucleotide exchange factor 3 [Source:HGNC Symbol;Acc:HGNC:16629] |
| ENSG00000106538 | RARRES2 | 965 | 460 | 134 | -1,07 | -1,78 | retinoic acid receptor responder 2 [Source:HGNC Symbol;Acc:HGNC:9868] |
| ENSG00000198915 | RASGEF1A | 255 | 44 | 5 | -2,53 | -3,21 | RasGEF domain family member 1A [Source:HGNC Symbol;Acc:HGNC:24246] |
| ENSG00000138670 | RASGEF1B | 326 | 42 | 8 | -2,96 | -2,41 | RasGEF domain family member 1B [Source:HGNC Symbol;Acc:HGNC:24881] |
| ENSG00000203867 | RBM20 | 334 | 105 | 21 | -1,67 | -2,33 | RNA binding motif protein 20 [Source:HGNC Symbol;Acc:HGNC:27424] |
| ENSG00000139890 | REM2 | 505 | 171 | 91 | -1,56 | -0,91 | RRAD and GEM like GTPase 2 [Source:HGNC Symbol;Acc:HGNC:20248] |
| ENSG00000117152 | RGS4 | 275 | 148 | 48 | -0,89 | -1,64 | regulator of G-protein signaling 4 [Source:HGNC Symbol;Acc:HGNC:10000] |
| ENSG00000135828 | RNASEL | 261 | 101 | 38 | -1,37 | -1,4 | ribonuclease L [Source:HGNC Symbol;Acc:HGNC:10050] |
| ENSG00000146373 | RNF217 | 866 | 524 | 303 | -0,73 | -0,79 | ring finger protein 217 [Source:HGNC Symbol;Acc:HGNC:21487] |
| ENSG00000185008 | ROBO2 | 7406 | 1613 | 505 | -2,2 | -1,68 | roundabout guidance receptor 2 [Source:HGNC Symbol;Acc:HGNC:10250] |
| ENSG00000154134 | ROBO3 | 246 | 26 | 8 | -3,25 | -1,68 | roundabout guidance receptor 3 [Source:HGNC Symbol;Acc:HGNC:13433] |
| ENSG00000072133 | RPS6KA6 | 1203 | 698 | 347 | -0,79 | -1,01 | ribosomal protein S6 kinase A6 [Source:HGNC Symbol;Acc:HGNC:10435] |
| ENSG00000189334 | S100A14 | 1209 | 119 | 14 | -3,35 | -3,06 | S100 calcium binding protein A14 [Source:HGNC Symbol;Acc:HGNC:18901] |
| ENSG00000167100 | SAMD14 | 719 | 234 | 125 | -1,62 | -0,92 | sterile alpha motif domain containing 14 [Source:HGNC Symbol;Acc:HGNC:27312] |
| ENSG00000130066 | SAT1 | 7886 | 3922 | 1274 | -1,01 | -1,62 | spermidine/spermine N1-acetyltransferase 1 [Source:HGNC Symbol;Acc:HGNC:10540] |
| ENSG00000183873 | SCN5A | 250 | 19 | 10 | -3,71 | -1,03 | sodium voltage-gated channel alpha subunit 5 [Source:HGNC Symbol;Acc:HGNC:10593] |
| ENSG00000111319 | SCNN1A | 4713 | 76 | 26 | -5,96 | -1,53 | sodium channel epithelial 1 alpha subunit [Source:HGNC Symbol;Acc:HGNC:10599] |
| ENSG00000125775 | SDCBP2 | 212 | 108 | 20 | -0,97 | -2,43 | syndecan binding protein 2 [Source:HGNC Symbol;Acc:HGNC:15756] |
| ENSG00000012171 | SEMA3B | 2185 | 644 | 302 | -1,76 | -1,09 | semaphorin 3B [Source:HGNC Symbol;Acc:HGNC:10724] |
| ENSG00000137872 | SEMA6D | 209 | 121 | 15 | -0,79 | -3 | semaphorin 6D [Source:HGNC Symbol;Acc:HGNC:16770] |
| ENSG00000179918 | SEPHS2 | 2956 | 651 | 283 | -2,18 | -1,2 | selenophosphate synthetase 2 [Source:HGNC Symbol;Acc:HGNC:19686] |
| ENSG00000135919 | SERPINE2 | 1394 | 493 | 139 | -1,5 | -1,83 | serpin family E member 2 [Source:HGNC Symbol;Acc:HGNC:8951] |
| ENSG00000174938 | SEZ6L2 | 3117 | 1003 | 489 | -1,63 | -1,04 | seizure related 6 homolog like 2 [Source:HGNC Symbol;Acc:HGNC:30844] |
| ENSG00000107819 | SFXN3 | 1640 | 246 | 73 | -2,74 | -1,75 | sideroflexin 3 [Source:HGNC Symbol;Acc:HGNC:16087] |
| ENSG00000125731 | SH2D3A | 1368 | 229 | 53 | -2,58 | -2,1 | SH2 domain containing 3A [Source:HGNC Symbol;Acc:HGNC:16885] |
| ENSG00000214193 | SH3D21 | 1130 | 395 | 175 | -1,52 | -1,17 | SH3 domain containing 21 [Source:HGNC Symbol;Acc:HGNC:26236] |
| ENSG00000035115 | SH3YL1 | 3520 | 821 | 435 | -2,1 | -0,92 | SH3 and SYLF domain containing 1 [Source:HGNC Symbol;Acc:HGNC:29546] |
| ENSG00000162105 | SHANK2 | 2905 | 679 | 384 | -2,1 | -0,82 | SH3 and multiple ankyrin repeat domains 2 [Source:HGNC Symbol;Acc:HGNC:14295] |
| ENSG00000198892 | SHISA4 | 343 | 105 | 39 | -1,71 | -1,45 | shisa family member 4 [Source:HGNC Symbol;Acc:HGNC:27139] |
| ENSG00000158352 | SHROOM4 | 837 | 500 | 290 | -0,74 | -0,79 | shroom family member 4 [Source:HGNC Symbol;Acc:HGNC:29215] |
| ENSG00000187164 | SHTN1 | 847 | 102 | 25 | -3,05 | -2,02 | shootin 1 [Source:HGNC Symbol;Acc:HGNC:29319] |
| ENSG00000185187 | SIGIRR | 1678 | 359 | 183 | -2,23 | -0,97 | single Ig and TIR domain containing [Source:HGNC Symbol;Acc:HGNC:30575] |
| ENSG00000112246 | SIM1 | 953 | 154 | 28 | -2,63 | -2,47 | single-minded family bHLH transcription factor 1 [Source:HGNC Symbol;Acc:HGNC:10882] |
| ENSG00000158296 | SLC13A3 | 224 | 56 | 32 | -2,01 | -0,81 | solute carrier family 13 member 3 [Source:HGNC Symbol;Acc:HGNC:14430] |
| ENSG00000146039 | SLC17A4 | 907 | 76 | 15 | -3,58 | -2,37 | solute carrier family 17 member 4 [Source:HGNC Symbol;Acc:HGNC:10932] |
| ENSG00000101194 | SLC17A9 | 2948 | 201 | 10 | -3,88 | -4,36 | solute carrier family 17 member 9 [Source:HGNC Symbol;Acc:HGNC:16192] |
| ENSG00000106688 | SLC1A1 | 405 | 185 | 108 | -1,13 | -0,78 | solute carrier family 1 member 1 [Source:HGNC Symbol;Acc:HGNC:10939] |
| ENSG00000092096 | SLC22A17 | 800 | 299 | 163 | -1,42 | -0,88 | solute carrier family 22 member 17 [Source:HGNC Symbol;Acc:HGNC:23095] |
| ENSG00000110628 | SLC22A18 | 836 | 65 | 8 | -3,68 | -2,97 | solute carrier family 22 member 18 [Source:HGNC Symbol;Acc:HGNC:10964] |
| ENSG00000153291 | SLC25A27 | 539 | 238 | 104 | -1,18 | -1,19 | solute carrier family 25 member 27 [Source:HGNC Symbol;Acc:HGNC:21065] |
| ENSG00000146411 | SLC2A12 | 542 | 268 | 92 | -1,02 | -1,54 | solute carrier family 2 member 12 [Source:HGNC Symbol;Acc:HGNC:18067] |
| ENSG00000173262 | SLC2A14 | 1353 | 189 | 57 | -2,84 | -1,72 | solute carrier family 2 member 14 [Source:HGNC Symbol;Acc:HGNC:18301] |
| ENSG00000059804 | SLC2A3 | 6787 | 696 | 56 | -3,29 | -3,63 | solute carrier family 2 member 3 [Source:HGNC Symbol;Acc:HGNC:11007] |
| ENSG00000110660 | SLC35F2 | 578 | 161 | 35 | -1,84 | -2,18 | solute carrier family 35 member F2 [Source:HGNC Symbol;Acc:HGNC:23615] |
| ENSG00000139540 | SLC39A5 | 2628 | 929 | 289 | -1,5 | -1,68 | solute carrier family 39 member 5 [Source:HGNC Symbol;Acc:HGNC:20502] |
| ENSG00000204385 | SLC44A4 | 1791 | 142 | 15 | -3,66 | -3,25 | solute carrier family 44 member 4 [Source:HGNC Symbol;Acc:HGNC:13941] |
| ENSG00000155465 | SLC7A7 | 895 | 217 | 50 | -2,04 | -2,11 | solute carrier family 7 member 7 [Source:HGNC Symbol;Acc:HGNC:11065] |
| ENSG00000092068 | SLC7A8 | 1437 | 733 | 128 | -0,97 | -2,52 | solute carrier family 7 member 8 [Source:HGNC Symbol;Acc:HGNC:11066] |
| ENSG00000137491 | SLCO2B1 | 352 | 124 | 35 | -1,51 | -1,81 | solute carrier organic anion transporter family member 2B1 [Source:HGNC Symbol;Acc:HGNC:10962] |
| ENSG00000166750 | SLFN5 | 833 | 262 | 123 | -1,67 | -1,1 | schlafen family member 5 [Source:HGNC Symbol;Acc:HGNC:28286] |
| ENSG00000184347 | SLIT3 | 1692 | 412 | 56 | -2,04 | -2,86 | slit guidance ligand 3 [Source:HGNC Symbol;Acc:HGNC:11087] |
| ENSG00000170545 | SMAGP | 227 | 99 | 54 | -1,19 | -0,87 | small cell adhesion glycoprotein [Source:HGNC Symbol;Acc:HGNC:26918] |
| ENSG00000178947 | SMIM10L2A | 885 | 360 | 164 | -1,3 | -1,13 | small integral membrane protein 10 like 2A [Source:HGNC Symbol;Acc:HGNC:34499] |
| ENSG00000108018 | SORCS1 | 1334 | 621 | 100 | -1,1 | -2,62 | sortilin related VPS10 domain containing receptor 1 [Source:HGNC Symbol;Acc:HGNC:16697] |
| ENSG00000184985 | SORCS2 | 251 | 120 | 62 | -1,07 | -0,96 | sortilin related VPS10 domain containing receptor 2 [Source:HGNC Symbol;Acc:HGNC:16698] |
| ENSG00000186212 | SOWAHB | 456 | 120 | 28 | -1,93 | -2,08 | sosondowah ankyrin repeat domain family member B [Source:HGNC Symbol;Acc:HGNC:32958] |

| | | | | | | | |
|---|---|---|---|---|---|---|---|
| ENSG00000067066 | SP100 | 265 | 89 | 44 | -1,58 | -1,01 | SP100 nuclear antigen [Source:HGNC Symbol;Acc:HGNC:11206] |
| ENSG00000166145 | SPINT1 | 13756 | 1667 | 253 | -3,04 | -2,72 | serine peptidase inhibitor, Kunitz type 1 [Source:HGNC Symbol;Acc:HGNC:11246] |
| ENSG00000167642 | SPINT2 | 14613 | 4920 | 1604 | -1,57 | -1,62 | serine peptidase inhibitor, Kunitz type 2 [Source:HGNC Symbol;Acc:HGNC:11247] |
| ENSG00000152377 | SPOCK1 | 1276 | 53 | 14 | -4,58 | -1,96 | SPARC/osteonectin, cwcv and kazal like domains proteoglycan 1 [Source:HGNC Symbol;Acc:HGNC:11251] |
| ENSG00000107742 | SPOCK2 | 9186 | 2100 | 199 | -2,13 | -3,4 | SPARC/osteonectin, cwcv and kazal like domains proteoglycan 2 [Source:HGNC Symbol;Acc:HGNC:13564] |
| ENSG00000118785 | SPP1 | 18387 | 359 | 123 | -5,68 | -1,55 | secreted phosphoprotein 1 [Source:HGNC Symbol;Acc:HGNC:11255] |
| ENSG00000277363 | SRCIN1 | 372 | 86 | 39 | -2,11 | -1,15 | SRC kinase signaling inhibitor 1 [Source:HGNC Symbol;Acc:HGNC:29506] |
| ENSG00000146700 | SSC4D | 2432 | 601 | 347 | -2,02 | -0,79 | scavenger receptor cysteine rich family member with 4 domains [Source:HGNC Symbol;Acc:HGNC:14461] |
| ENSG00000172830 | SSH3 | 1757 | 721 | 299 | -1,28 | -1,27 | slingshot protein phosphatase 3 [Source:HGNC Symbol;Acc:HGNC:30581] |
| ENSG00000149418 | ST14 | 6558 | 208 | 18 | -4,98 | -3,51 | suppression of tumorigenicity 14 [Source:HGNC Symbol;Acc:HGNC:11344] |
| ENSG00000178078 | STAP2 | 1458 | 571 | 228 | -1,35 | -1,32 | signal transducing adaptor family member 2 [Source:HGNC Symbol;Acc:HGNC:30430] |
| ENSG00000113739 | STC2 | 944 | 157 | 33 | -2,59 | -2,27 | stanniocalcin 2 [Source:HGNC Symbol;Acc:HGNC:11374] |
| ENSG00000157214 | STEAP2 | 351 | 196 | 74 | -0,84 | -1,4 | STEAP2 metalloreductase [Source:HGNC Symbol;Acc:HGNC:17885] |
| ENSG00000127561 | SYNGR3 | 448 | 153 | 60 | -1,55 | -1,37 | synaptogyrin 3 [Source:HGNC Symbol;Acc:HGNC:11501] |
| ENSG00000019505 | SYT13 | 1314 | 484 | 188 | -1,44 | -1,36 | synaptotagmin 13 [Source:HGNC Symbol;Acc:HGNC:14962] |
| ENSG00000184292 | TACSTD2 | 945 | 10 | 1 | -6,55 | -3,33 | tumor-associated calcium signal transducer 2 [Source:HGNC Symbol;Acc:HGNC:11530] |
| ENSG00000168394 | TAP1 | 226 | 43 | 10 | -2,39 | -2,08 | transporter 1, ATP binding cassette subfamily B member [Source:HGNC Symbol;Acc:HGNC:43] |
| ENSG00000143178 | TBX19 | 210 | 119 | 47 | -0,82 | -1,33 | T-box 19 [Source:HGNC Symbol;Acc:HGNC:11596] |
| ENSG00000165929 | TC2N | 1483 | 709 | 322 | -1,06 | -1,14 | tandem C2 domains, nuclear [Source:HGNC Symbol;Acc:HGNC:19859] |
| ENSG00000204219 | TCEA3 | 573 | 248 | 96 | -1,21 | -1,37 | transcription elongation factor A3 [Source:HGNC Symbol;Acc:HGNC:11615] |
| ENSG00000166046 | TCP11L2 | 364 | 127 | 56 | -1,52 | -1,19 | t-complex 11 like 2 [Source:HGNC Symbol;Acc:HGNC:28627] |
| ENSG00000132749 | TESMIN | 774 | 147 | 86 | -2,4 | -0,78 | testis expressed metallothionein like protein [Source:HGNC Symbol;Acc:HGNC:7446] |
| ENSG00000105825 | TFPI2 | 653 | 53 | 12 | -3,62 | -2,12 | tissue factor pathway inhibitor 2 [Source:HGNC Symbol;Acc:HGNC:11761] |
| ENSG00000137801 | THBS1 | 1804 | 860 | 442 | -1,07 | -0,96 | thrombospondin 1 [Source:HGNC Symbol;Acc:HGNC:11785] |
| ENSG00000144229 | THSD7B | 321 | 74 | 22 | -2,11 | -1,76 | thrombospondin type 1 domain containing 7B [Source:HGNC Symbol;Acc:HGNC:29348] |
| ENSG00000223573 | TINCR | 292 | 30 | 7 | -3,29 | -2,15 | tissue differentiation-inducing non-protein coding RNA [Source:HGNC Symbol;Acc:HGNC:14607] |
| ENSG00000105289 | TJP3 | 1781 | 217 | 63 | -3,04 | -1,79 | tight junction protein 3 [Source:HGNC Symbol;Acc:HGNC:11829] |
| ENSG00000167608 | TMC4 | 4423 | 183 | 28 | -4,59 | -2,71 | transmembrane channel like 4 [Source:HGNC Symbol;Acc:HGNC:22998] |
| ENSG00000179178 | TMEM125 | 1151 | 35 | 8 | -5,03 | -2,23 | transmembrane protein 125 [Source:HGNC Symbol;Acc:HGNC:28275] |
| ENSG00000178826 | TMEM139 | 238 | 76 | 13 | -1,64 | -2,53 | transmembrane protein 139 [Source:HGNC Symbol;Acc:HGNC:22058] |
| ENSG00000011638 | TMEM159 | 1092 | 391 | 165 | -1,48 | -1,25 | transmembrane protein 159 [Source:HGNC Symbol;Acc:HGNC:30136] |
| ENSG00000002933 | TMEM176A | 323 | 80 | 17 | -2,01 | -2,25 | transmembrane protein 176A [Source:HGNC Symbol;Acc:HGNC:24930] |
| ENSG00000198133 | TMEM229B | 813 | 260 | 64 | -1,65 | -2,03 | transmembrane protein 229B [Source:HGNC Symbol;Acc:HGNC:20130] |
| ENSG00000182107 | TMEM30B | 1799 | 71 | 14 | -4,66 | -2,31 | transmembrane protein 30B [Source:HGNC Symbol;Acc:HGNC:27254] |
| ENSG00000121900 | TMEM54 | 539 | 70 | 27 | -2,94 | -1,4 | transmembrane protein 54 [Source:HGNC Symbol;Acc:HGNC:24143] |
| ENSG00000125895 | TMEM74B | 991 | 337 | 144 | -1,56 | -1,22 | transmembrane protein 74B [Source:HGNC Symbol;Acc:HGNC:15893] |
| ENSG00000167105 | TMEM92 | 421 | 95 | 17 | -2,15 | -2,46 | transmembrane protein 92 [Source:HGNC Symbol;Acc:HGNC:26579] |
| ENSG00000184012 | TMPRSS2 | 1574 | 243 | 24 | -2,69 | -3,34 | transmembrane protease, serine 2 [Source:HGNC Symbol;Acc:HGNC:11876] |
| ENSG00000137648 | TMPRSS4 | 241 | 25 | 2 | -3,26 | -3,56 | transmembrane protease, serine 4 [Source:HGNC Symbol;Acc:HGNC:11878] |
| ENSG00000173535 | TNFRSF10C | 840 | 385 | 141 | -1,12 | -1,45 | TNF receptor superfamily member 10c [Source:HGNC Symbol;Acc:HGNC:11906] |
| ENSG00000127863 | TNFRSF19 | 699 | 127 | 44 | -2,46 | -1,54 | TNF receptor superfamily member 19 [Source:HGNC Symbol;Acc:HGNC:11915] |
| ENSG00000174292 | TNK1 | 311 | 73 | 39 | -2,09 | -0,89 | tyrosine kinase non receptor 1 [Source:HGNC Symbol;Acc:HGNC:11940] |
| ENSG00000248290 | TNXA | 240 | 44 | 10 | -2,44 | -2,23 | tenascin XA (pseudogene) [Source:HGNC Symbol;Acc:HGNC:11975] |
| ENSG00000168477 | TNXB | 1125 | 214 | 46 | -2,4 | -2,22 | tenascin XB [Source:HGNC Symbol;Acc:HGNC:11976] |
| ENSG00000146242 | TPBG | 1758 | 430 | 254 | -2,03 | -0,76 | trophoblast glycoprotein [Source:HGNC Symbol;Acc:HGNC:12004] |
| ENSG00000168016 | TRANK1 | 1548 | 899 | 338 | -0,78 | -1,41 | tetratricopeptide repeat and ankyrin repeat containing 1 [Source:HGNC Symbol;Acc:HGNC:29011] |
| ENSG00000132109 | TRIM21 | 511 | 295 | 70 | -0,79 | -2,07 | tripartite motif containing 21 [Source:HGNC Symbol;Acc:HGNC:11312] |
| ENSG00000121236 | TRIM6 | 458 | 67 | 6 | -2,77 | -3,37 | tripartite motif containing 6 [Source:HGNC Symbol;Acc:HGNC:16277] |
| ENSG00000182463 | TSHZ2 | 1329 | 686 | 395 | -0,95 | -0,8 | teashirt zinc finger homeobox 2 [Source:HGNC Symbol;Acc:HGNC:13010] |
| ENSG00000215845 | TSTD1 | 749 | 147 | 11 | -2,35 | -3,77 | thiosulfate sulfurtransferase like domain containing 1 [Source:HGNC Symbol;Acc:HGNC:35410] |
| ENSG00000085831 | TTC39A | 611 | 255 | 153 | -1,26 | -0,74 | tetratricopeptide repeat domain 39A [Source:HGNC Symbol;Acc:HGNC:18657] |
| ENSG00000166402 | TUB | 1403 | 505 | 63 | -1,48 | -3 | tubby bipartite transcription factor [Source:HGNC Symbol;Acc:HGNC:12406] |
| ENSG00000137766 | UNC13C | 501 | 39 | 10 | -3,69 | -2,03 | unc-13 homolog C [Source:HGNC Symbol;Acc:HGNC:23149] |
| ENSG00000156687 | UNC5D | 275 | 37 | 7 | -2,89 | -2,46 | unc-5 netrin receptor D [Source:HGNC Symbol;Acc:HGNC:18634] |
| ENSG00000168140 | VASN | 1161 | 653 | 301 | -0,83 | -1,12 | vasorin [Source:HGNC Symbol;Acc:HGNC:18517] |
| ENSG00000112715 | VEGFA | 15553 | 5905 | 3217 | -1,4 | -0,88 | vascular endothelial growth factor A [Source:HGNC Symbol;Acc:HGNC:12680] |
| ENSG00000136059 | VILL | 329 | 65 | 32 | -2,35 | -1,01 | villin like [Source:HGNC Symbol;Acc:HGNC:30906] |
| ENSG00000110002 | VWA5A | 1030 | 212 | 101 | -2,28 | -1,06 | von Willebrand factor A domain containing 5A [Source:HGNC Symbol;Acc:HGNC:6658] |
| ENSG00000187260 | WDR86 | 1506 | 734 | 169 | -1,04 | -2,12 | WD repeat domain 86 [Source:HGNC Symbol;Acc:HGNC:28020] |
| ENSG00000101443 | WFDC2 | 4923 | 663 | 75 | -2,89 | -3,14 | WAP four-disulfide core domain 2 [Source:HGNC Symbol;Acc:HGNC:15939] |
| ENSG00000018408 | WWTR1 | 2382 | 1365 | 802 | -0,8 | -0,77 | WW domain containing transcription regulator 1 [Source:HGNC Symbol;Acc:HGNC:24042] |
| ENSG00000103489 | XYLT1 | 665 | 47 | 6 | -3,83 | -2,99 | xylosyltransferase 1 [Source:HGNC Symbol;Acc:HGNC:15516] |
| ENSG00000006047 | YBX2 | 471 | 165 | 47 | -1,51 | -1,83 | Y-box binding protein 2 [Source:HGNC Symbol;Acc:HGNC:17948] |
| ENSG00000159714 | ZDHHC1 | 437 | 246 | 143 | -0,83 | -0,78 | zinc finger DHHC-type containing 1 [Source:HGNC Symbol;Acc:HGNC:17916] |
| ENSG00000147394 | ZNF185 | 682 | 226 | 97 | -1,59 | -1,22 | zinc finger protein 185 (LIM domain) [Source:HGNC Symbol;Acc:HGNC:12976] |
| ENSG00000148143 | ZNF462 | 1674 | 820 | 351 | -1,03 | -1,22 | zinc finger protein 462 [Source:HGNC Symbol;Acc:HGNC:21684] |
| ENSG00000197162 | ZNF785 | 1436 | 9 | 1 | -7,25 | -2,84 | zinc finger protein 785 [Source:HGNC Symbol;Acc:HGNC:26496] |

**Table S2** Regulated genes in CINI expansion. In group 1 there are genes continuously upregulated in p5 and p10, in group 2 genes upregulated at p5 but stable at p10, in group 3 genes downregulated at p5 but stable in p10 and in group 4 genes continuously downregulated.

| Ensembl ID | Gene Symbol | PP P0 1 | PP P0 2 | PP P0 3 | PP P0 4 | PP P5 1 | PP P5 2 | PP P5 2 | PP P10 1 | PP P10 2 | PP P10 3 | Description |
|---|---|---|---|---|---|---|---|---|---|---|---|---|
| **TGFB LIGANDS AND RECEPTORS** | | | | | | | | | | | | |
| ENSG00000123999 | INHA | 843 | 1019 | 1843 | 562 | 55 | 111 | 84 | 53 | 44 | 160 | inhibin alpha subunit [Source:HGNC Symbol;Acc:HGNC:6065] |
| ENSG00000135414 | GDF11 | 2653 | 2097 | 2375 | 2185 | 3805 | 4514 | 3764 | 3822 | 3625 | 3475 | growth differentiation factor 11 [Source:HGNC Symbol;Acc:HGNC:4216] |
| ENSG00000125845 | BMP2 | 3613 | 4345 | 4085 | 3823 | 795 | 926 | 541 | 557 | 626 | 668 | bone morphogenetic protein 2 [Source:HGNC Symbol;Acc:HGNC:1069] |
| ENSG00000101144 | BMP7 | 462 | 366 | 691 | 282 | 881 | 700 | 647 | 1017 | 1127 | 1070 | bone morphogenetic protein 7 [Source:HGNC Symbol;Acc:HGNC:1074] |
| ENSG00000163235 | TGFA | 874 | 714 | 677 | 830 | 49 | 75 | 108 | 81 | 50 | 298 | transforming growth factor alpha [Source:HGNC Symbol;Acc:HGNC:11765] |
| ENSG00000105329 | TGFB1 | 4806 | 5648 | 4606 | 3379 | 1719 | 2386 | 1741 | 1316 | 998 | 1262 | transforming growth factor beta 1 [Source:HGNC Symbol;Acc:HGNC:11766] |
| ENSG00000140682 | TGFB1I1 | 324 | 510 | 436 | 340 | 257 | 470 | 390 | 271 | 199 | 263 | transforming growth factor beta 1 induced transcript 1 [Source:HGNC Symbol;Acc:HGNC:11767] |
| ENSG00000092969 | TGFB2 | 680 | 986 | 1285 | 759 | 248 | 409 | 450 | 405 | 502 | 546 | transforming growth factor beta 2 [Source:HGNC Symbol;Acc:HGNC:11768] |
| ENSG00000119699 | TGFB3 | 123 | 165 | 244 | 281 | 140 | 266 | 295 | 178 | 150 | 338 | transforming growth factor beta 3 [Source:HGNC Symbol;Acc:HGNC:11769] |
| ENSG00000115170 | ACVR1 (ALK2) | 1097 | 1140 | 828 | 998 | 745 | 844 | 785 | 816 | 891 | 818 | activin A receptor type 1 [Source:HGNC Symbol;Acc:HGNC:171] |
| ENSG00000135503 | ACVR1B (ALK4) | 4989 | 3394 | 4119 | 3676 | 4235 | 4002 | 4226 | 4790 | 5412 | 4636 | activin A receptor type 1B [Source:HGNC Symbol;Acc:HGNC:172] |
| ENSG00000139567 | ACVRL1 (ALK1) | 73 | 63 | 41 | 46 | 103 | 178 | 122 | 126 | 93 | 129 | activin A receptor like type 1 [Source:HGNC Symbol;Acc:HGNC:175] |
| ENSG00000107779 | BMPR1A (ALK3) | 2728 | 2159 | 1892 | 2266 | 2062 | 2106 | 2044 | 2216 | 2333 | 2176 | bone morphogenetic protein receptor type 1A [Source:HGNC Symbol;Acc:HGNC:1076] |
| ENSG00000106799 | TGFBR1 (ALK5) | 3435 | 2536 | 3338 | 2969 | 3712 | 4021 | 4520 | 4852 | 4861 | 4616 | transforming growth factor beta receptor 1 [Source:HGNC Symbol;Acc:HGNC:11772] |
| ENSG00000138696 | BMPR1B (ALK6) | 263 | 217 | 211 | 317 | 387 | 189 | 278 | 362 | 361 | 338 | bone morphogenetic protein receptor type 1B [Source:HGNC Symbol;Acc:HGNC:1077] |
| ENSG00000121989 | ACVR2A | 1130 | 1087 | 616 | 1038 | 1319 | 1577 | 1461 | 1896 | 2211 | 1607 | activin A receptor type 2A [Source:HGNC Symbol;Acc:HGNC:173] |
| ENSG00000114739 | ACVR2B | 6065 | 4850 | 5996 | 5126 | 6700 | 6027 | 6887 | 7592 | 7369 | 5696 | activin A receptor type 2B [Source:HGNC Symbol;Acc:HGNC:174] |
| ENSG00000163513 | TGFBR2 | 1845 | 1912 | 1568 | 1556 | 2177 | 2443 | 2676 | 2859 | 3086 | 2989 | transforming growth factor beta receptor 2 [Source:HGNC Symbol;Acc:HGNC:11773] |
| ENSG00000204217 | BMPR2 | 2521 | 2240 | 1967 | 2323 | 1593 | 1742 | 1431 | 1182 | 887 | 796 | bone morphogenetic protein receptor type 2 [Source:HGNC Symbol;Acc:HGNC:1078] |
| **FGF LIGANDS AND RECEPTORS** | | | | | | | | | | | | |
| ENSG00000161958 | FGF11 | 144 | 187 | 207 | 167 | 83 | 172 | 74 | 25 | 28 | 13 | fibroblast growth factor 11 [Source:HGNC Symbol;Acc:HGNC:3667] |
| ENSG00000114279 | FGF12 | 121 | 155 | 194 | 91 | 5 | 2 | 16 | 5 | 0 | 4 | fibroblast growth factor 12 [Source:HGNC Symbol;Acc:HGNC:3668] |
| ENSG00000129682 | FGF13 | 327 | 286 | 294 | 321 | 165 | 18 | 4 | 0 | 0 | 3 | fibroblast growth factor 13 [Source:HGNC Symbol;Acc:HGNC:3670] |
| ENSG00000102466 | FGF14 | 41 | 9 | 191 | 159 | 5 | 35 | 14 | 1 | 3 | 0 | fibroblast growth factor 14 [Source:HGNC Symbol;Acc:HGNC:3671] |
| ENSG00000156427 | FGF18 | 524 | 202 | 333 | 270 | 7103 | 9338 | 10310 | 15389 | 17133 | 14785 | fibroblast growth factor 18 [Source:HGNC Symbol;Acc:HGNC:3674] |
| ENSG00000138685 | FGF2 | 183 | 180 | 199 | 191 | 338 | 397 | 430 | 527 | 540 | 281 | fibroblast growth factor 2 [Source:HGNC Symbol;Acc:HGNC:3676] |
| ENSG00000102678 | FGF9 | 243 | 259 | 116 | 194 | 209 | 267 | 357 | 282 | 352 | 459 | fibroblast growth factor 9 [Source:HGNC Symbol;Acc:HGNC:3687] |
| ENSG00000077782 | FGFR1 | 5194 | 5720 | 5752 | 5709 | 4498 | 4971 | 5531 | 5200 | 5426 | 4851 | fibroblast growth factor receptor 1 [Source:HGNC Symbol;Acc:HGNC:3688] |
| ENSG00000066468 | FGFR2 | 6615 | 3566 | 4031 | 3975 | 2584 | 2415 | 2948 | 3188 | 3048 | 2618 | fibroblast growth factor receptor 2 [Source:HGNC Symbol;Acc:HGNC:3689] |
| ENSG00000068078 | FGFR3 | 19069 | 19177 | 14087 | 16752 | 8440 | 7643 | 7451 | 7469 | 7232 | 7756 | fibroblast growth factor receptor 3 [Source:HGNC Symbol;Acc:HGNC:3690] |
| ENSG00000160867 | FGFR4 | 17275 | 13539 | 11906 | 10908 | 12364 | 11956 | 13309 | 15865 | 17372 | 16523 | fibroblast growth factor receptor 4 [Source:HGNC Symbol;Acc:HGNC:3691] |
| **RETINOIC ACID ENZYMES AND RECEPTORS** | | | | | | | | | | | | |
| ENSG00000165092 | ALDH1A1 | 4940 | 3619 | 5576 | 3361 | 2939 | 302 | 151 | 134 | 100 | 280 | aldehyde dehydrogenase 1 family member A1 [Source:HGNC Symbol;Acc:HGNC:402] |
| ENSG00000072042 | RDH11 | 5764 | 4635 | 4016 | 4821 | 4245 | 3881 | 3866 | 4657 | 5113 | 4684 | retinol dehydrogenase 11 (all-trans/9-cis/11-cis) [Source:HGNC Symbol;Acc:HGNC:17964] |
| ENSG00000131759 | RARA | 1727 | 1834 | 1546 | 1454 | 1560 | 1666 | 1421 | 1048 | 1428 | 1416 | retinoic acid receptor alpha [Source:HGNC Symbol;Acc:HGNC:9864] |
| ENSG00000172819 | RARG | 757 | 952 | 566 | 748 | 429 | 819 | 977 | 417 | 367 | 510 | retinoic acid receptor gamma [Source:HGNC Symbol;Acc:HGNC:9866] |
| ENSG00000186350 | RXRA | 4378 | 3838 | 3748 | 2973 | 2864 | 2930 | 3132 | 2952 | 3233 | 3222 | retinoid X receptor alpha [Source:HGNC Symbol;Acc:HGNC:10477] |
| ENSG00000204231 | RXRB | 2076 | 2789 | 2298 | 2654 | 1358 | 1688 | 1760 | 1352 | 1445 | 1550 | retinoid X receptor beta [Source:HGNC Symbol;Acc:HGNC:10478] |
| **NOTCH LIGANDS, RECEPTORS AND EFFECTORS** | | | | | | | | | | | | |
| ENSG00000148400 | NOTCH1 | 7662 | 5343 | 6394 | 6129 | 2648 | 3592 | 5785 | 5212 | 4597 | 5075 | notch 1 [Source:HGNC Symbol;Acc:HGNC:7881] |
| ENSG00000134250 | NOTCH2 | 16764 | 15711 | 12307 | 18299 | 8621 | 7134 | 5649 | 5851 | 4526 | 4783 | notch 2 [Source:HGNC Symbol;Acc:HGNC:7882] |
| ENSG00000074181 | NOTCH3 | 17705 | 15681 | 14224 | 14246 | 6120 | 7521 | 7686 | 7568 | 6920 | 6784 | notch 3 [Source:HGNC Symbol;Acc:HGNC:7883] |
| ENSG00000204301 | NOTCH4 | 173 | 62 | 305 | 254 | 68 | 120 | 117 | 64 | 62 | 70 | notch 4 [Source:HGNC Symbol;Acc:HGNC:7884] |
| ENSG00000101384 | JAG1 | 16084 | 16783 | 15493 | 16498 | 5197 | 4153 | 7704 | 8394 | 7152 | 7110 | jagged 1 [Source:HGNC Symbol;Acc:HGNC:6188] |
| ENSG00000184916 | JAG2 | 500 | 778 | 614 | 674 | 515 | 606 | 482 | 188 | 175 | 128 | jagged 2 [Source:HGNC Symbol;Acc:HGNC:6189] |
| ENSG00000198719 | DLL1 | 1310 | 824 | 3056 | 1230 | 298 | 657 | 1275 | 1228 | 1069 | 1485 | delta like canonical Notch ligand 1 [Source:HGNC Symbol;Acc:HGNC:2908] |
| ENSG00000090932 | DLL3 | 51 | 18 | 108 | 37 | 142 | 714 | 2147 | 1833 | 1438 | 2251 | delta like canonical Notch ligand 3 [Source:HGNC Symbol;Acc:HGNC:2909] |
| ENSG00000128917 | DLL4 | 211 | 177 | 207 | 257 | 159 | 239 | 481 | 755 | 1117 | 1004 | delta like canonical Notch ligand 4 [Source:HGNC Symbol;Acc:HGNC:2910] |
| ENSG00000135547 | HEY2 | 43 | 23 | 34 | 21 | 204 | 230 | 280 | 388 | 277 | 652 | hes related family bHLH transcription factor with YRPW motif 2 [Source:HGNC Symbol;Acc:HGNC:4881] |
| ENSG00000163909 | HEYL | 13 | 61 | 230 | 27 | 31 | 44 | 71 | 22 | 4 | 52 | hes related family bHLH transcription factor with YRPW motif-like [Source:HGNC Symbol;Acc:HGNC:4882] |
| ENSG00000114315 | HES1 | 3890 | 3196 | 4427 | 4020 | 2250 | 2115 | 2153 | 1438 | 1172 | 1011 | hes family bHLH transcription factor 1 [Source:HGNC Symbol;Acc:HGNC:5192] |
| **PDGF LIGANDS AND RECEPTORS** | | | | | | | | | | | | |
| ENSG00000197461 | PDGFA | 2917 | 2861 | 3700 | 3094 | 674 | 544 | 411 | 100 | 85 | 134 | platelet derived growth factor subunit A [Source:HGNC Symbol;Acc:HGNC:8799] |
| ENSG00000100311 | PDGFB | 602 | 774 | 1406 | 1009 | 210 | 153 | 204 | 72 | 49 | 72 | platelet derived growth factor subunit B [Source:HGNC Symbol;Acc:HGNC:8800] |
| ENSG00000145431 | PDGFC | 3190 | 2903 | 1939 | 2983 | 1257 | 1614 | 1302 | 1301 | 1556 | 1520 | platelet derived growth factor C [Source:HGNC Symbol;Acc:HGNC:8801] |
| ENSG00000170962 | PDGFD | 752 | 1006 | 724 | 791 | 1437 | 1996 | 920 | 1124 | 1463 | 1545 | platelet derived growth factor D [Source:HGNC Symbol;Acc:HGNC:30620] |
| ENSG00000134853 | PDGFRA | 1655 | 1737 | 1190 | 1190 | 1676 | 1842 | 2532 | 2929 | 1885 | 1650 | platelet derived growth factor receptor alpha [Source:HGNC Symbol;Acc:HGNC:8803] |
| ENSG00000113721 | PDGFRB | 186 | 730 | 225 | 183 | 947 | 1877 | 2766 | 244 | 151 | 571 | platelet derived growth factor receptor beta [Source:HGNC Symbol;Acc:HGNC:8804] |
| **WNT LIGANDS AND RECEPTORS** | | | | | | | | | | | | |
| ENSG00000108379 | WNT3 | 262 | 254 | 286 | 224 | 215 | 159 | 215 | 263 | 230 | 227 | Wnt family member 3 [Source:HGNC Symbol;Acc:HGNC:12782] |
| ENSG00000162552 | WNT4 | 67 | 67 | 149 | 104 | 21 | 60 | 97 | 48 | 39 | 108 | Wnt family member 4 [Source:HGNC Symbol;Acc:HGNC:12783] |
| ENSG00000114251 | WNT5A | 69 | 194 | 119 | 59 | 179 | 337 | 517 | 243 | 131 | 131 | Wnt family member 5A [Source:HGNC Symbol;Acc:HGNC:12784] |
| ENSG00000188064 | WNT7B | 270 | 330 | 583 | 207 | 60 | 181 | 263 | 131 | 176 | 320 | Wnt family member 7B [Source:HGNC Symbol;Acc:HGNC:12787] |
| ENSG00000075290 | WNT8B | 95 | 86 | 111 | 48 | 179 | 57 | 53 | 61 | 37 | 42 | Wnt family member 8B [Source:HGNC Symbol;Acc:HGNC:12789] |
| ENSG00000157240 | FZD1 | 673 | 792 | 465 | 464 | 917 | 1053 | 1350 | 1446 | 1557 | 1479 | frizzled class receptor 1 [Source:HGNC Symbol;Acc:HGNC:4038] |
| ENSG00000180340 | FZD2 | 2049 | 1814 | 2638 | 1230 | 1496 | 1499 | 2276 | 1973 | 1857 | 1698 | frizzled class receptor 2 [Source:HGNC Symbol;Acc:HGNC:4040] |
| ENSG00000104290 | FZD3 | 926 | 663 | 808 | 794 | 741 | 526 | 629 | 942 | 643 | 706 | frizzled class receptor 3 [Source:HGNC Symbol;Acc:HGNC:4041] |
| ENSG00000174804 | FZD4 | 1766 | 1667 | 1276 | 1444 | 1193 | 904 | 1128 | 1576 | 1446 | 1305 | frizzled class receptor 4 [Source:HGNC Symbol;Acc:HGNC:4042] |
| ENSG00000163251 | FZD5 | 15559 | 9700 | 14256 | 12793 | 9898 | 6820 | 10908 | 21429 | 24156 | 15162 | frizzled class receptor 5 [Source:HGNC Symbol;Acc:HGNC:4043] |
| ENSG00000164930 | FZD6 | 1026 | 893 | 890 | 1121 | 852 | 786 | 838 | 756 | 774 | 739 | frizzled class receptor 6 [Source:HGNC Symbol;Acc:HGNC:4044] |
| ENSG00000155760 | FZD7 | 6381 | 3752 | 5850 | 3896 | 6301 | 3686 | 3318 | 4926 | 5291 | 5034 | frizzled class receptor 7 [Source:HGNC Symbol;Acc:HGNC:4045] |
| ENSG00000177283 | FZD8 | 3356 | 2529 | 2734 | 2241 | 1578 | 2367 | 1883 | 1766 | 1430 | 1172 | frizzled class receptor 8 [Source:HGNC Symbol;Acc:HGNC:4046] |
| ENSG00000188763 | FZD9 | 119 | 113 | 214 | 73 | 322 | 673 | 712 | 402 | 248 | 411 | frizzled class receptor 9 [Source:HGNC Symbol;Acc:HGNC:4047] |
| ENSG00000162337 | LRP5 | 8074 | 6695 | 6381 | 5916 | 8033 | 8158 | 7563 | 8418 | 7951 | 8161 | LDL receptor related protein 6 [Source:HGNC Symbol;Acc:HGNC:6698] |
| ENSG00000070018 | LRP6 | 7345 | 5932 | 6140 | 6304 | 5924 | 5546 | 6336 | 7229 | 5613 | 6483 | LDL receptor related protein 6 [Source:HGNC Symbol;Acc:HGNC:6698] |
| ENSG00000185483 | ROR1 | 1914 | 1313 | 1987 | 1268 | 915 | 811 | 1437 | 1465 | 1686 | 1521 | receptor tyrosine kinase like orphan receptor 1 [Source:HGNC Symbol;Acc:HGNC:10256] |
| ENSG00000169071 | ROR2 | 2476 | 1772 | 1803 | 1653 | 3574 | 3735 | 3802 | 3481 | 3970 | 3371 | receptor-like tyrosine kinase [Source:HGNC Symbol;Acc:HGNC:10481] |
| ENSG00000163785 | RYK | 2337 | 2179 | 2096 | 2431 | 1962 | 1821 | 1907 | 2388 | 2157 | 2054 | receptor-like tyrosine kinase [Source:HGNC Symbol;Acc:HGNC:10481] |
| **HIPPO SIGNALLING PATHWAY COMPONENTS** | | | | | | | | | | | | |
| ENSG00000018408 | WWTR1 | 2270 | 2405 | 2451 | 2404 | 1350 | 1606 | 1140 | 791 | 671 | 945 | WW domain containing transcription regulator 1 [Source:HGNC Symbol;Acc:HGNC:24042] |
| ENSG00000162105 | SHANK2 | 2731 | 2719 | 2881 | 3291 | 767 | 918 | 353 | 322 | 423 | 405 | SH3 and multiple ankyrin repeat domains 2 [Source:HGNC Symbol;Acc:HGNC:14295] |

**Table S3** Gene regulation of components of selected signaling pathways during expansion in CINI

| Stage | Duration (days) | Basal media | Supplements |
| --- | --- | --- | --- |
| **S1 (DE)** | **3** | MCDB131 (Life Technologies, 10372-019)<br>2.5mM Glucose (Sigma, G8769)<br>1x Glutmax (Life Technologies, 35050038)<br>1.5 g/l NaHCO3 (Sigma-Aldrich, S6297)<br>0.5 % BSA (Serva, 11945)<br>1x Pen/Strep (Life technologies, 15140-122) | 100 ng/ml Activin A (Peprotech, 120-14P) (all three days)<br>3.0 µM of CHIR (Tocris, SML1046) (1st day)<br>0.3 µM of CHIR (Tocris, SML1046) (2nd day) |
| **S2 (PGT)** | **2** | | 50 ng/ml FGF7 (Peprotech, 100-19)<br>0.25 mM Vitamin C (Tocris, 4055)<br>IWP-2 (Tocris, 3533) |
| **S3 (PF)** | **2** | MCDB131 (Life Technologies, 10372-019)<br>4.5 mM Glucose (Sigma, G8769)<br>1x Glutmax (Life Technologies (35050038)<br>1.5 g/l NaHCO3 (Sigma-Aldrich S6297)<br>2% BSA (Serva, 11945)<br>1x Pen/Strep (Life technologies, 15140-122)<br>0.5% IST-X (Life technologies, 51500056) | 50 ng/ml FGF7 (Peprotech, 100-19)<br>0.25 mM Vitamin C (Tocris, 4055)<br>0.25 µM SANT1 (Sigma, S4572)<br>100 nM LDN (Sigma, SML0559)<br>1 µM Retinoic Acid (Sigma, R2625)<br>200 nM TPB (Merck, 565740) |
| **S4 (PP)** | **3** | | 0.25 mM Vitamin C (Tocris, 4055)<br>2 ng/ml FGF7 (Peprotech, 100-19)<br>0.25 µM SANT1 (Sigma, S4572)<br>0.1 µM Retinoic Acid (Sigma, R2625)<br>200 nM LDN (Sigma, SML0559)<br>100 nM TPB (Merck, 565740) |
| **S5 (PEP)** | **4** | MCDB131 (Life Technologies, 10372-019)<br>14.5 mM Glucose (Sigma, G8769<br>1x Glutmax (Life Technologies, 35050038)<br>1.5 g/l NaHCO3 (Sigma-Aldrich, S6297)<br>2% BSA (Serva, 11945)<br>1x Pen/Strep (Life Technologies, 15140-122)<br>0.5 x IST-X (Life Technologies, 51500056)<br>1.8u/ml Heparin (Sigma, 2106-10VL) | 0.25 µM SANT1 (Sigma, S4572)<br>0.05 µM Retinoic Acid (Sigma, R2625)<br>100 nM LDN (Sigma, SML0559)<br>10 µM ALK5i (Milltenybiotech, 130117340)<br>1 µM T3 (Sigma, T6397)<br>10 µM ZnSO4 (Sigma, Z0251) |
| **S6** | **7** | | 100 nM LDN (Sigma, SML0559)<br>10 nM XX (Millipore, 565789)<br>10 µM ALK5i (Millteny Biotech, 130117340)<br>1 µM T3 (Sigma, T6397)<br>10 µM ZnSO4 (Sigma, Z0251) |
| **S7** | **10-14** | MCDB131 (Life Technologies, 10372-019)<br>1x Glutmax (Life Technologies, 35050038)<br>1.5 g/l NaHCO3 (Sigma-Aldrich, S6297)<br>0.5 mM Sodium Pyruvate (Lonza BE13-115E)<br>2% BSA (Serva, 11945)<br>1x Pen/Strep (Life Technologies, 15140-122)<br>0.5 x IST-X (Life Technologies, 51500056)<br>1.8u/ml Heparin (Sigma, 2106-10VL) | 1mM N-Cyst (Sigma, A9165)<br>2 µM R428 (Selleckchem, S2841)<br>10 µM ALK5i (Millteny Biotech, 130117340)<br>1 µM T3 (Sigma, T6397)<br>10 µM ZnSO4 (Sigma, Z0251)<br>10 µM Trolox (Millipore, 648471) |

**Table S4** Basal media and supplements for S1-S7

| GENE SYMBOL | PP (Ma et al) | PP FEEDER p9 | PP FEEDER p9 | PP FEEDER p21 | PP FEEDER p21 | PP (Nakamura et al) | PP (Nakamura et al) | PP (Nakamura et al) | PP FN p2 | PP FN p2 | PP FN p2 | PP FN p10 | PP FN p10 | PP FN p10 | PP (this study) | PP (this study) | PP (this study) | PP VTN-N p5 | PP VTN-N p5 | PP VTN-N p5 | PP VTN-N p5 | PP FN VTN-N p10 | PP FN VTN-N p10 |
|---|---|---|---|---|---|---|---|---|---|---|---|---|---|---|---|---|---|---|---|---|---|---|---|
| **DUCT MARKERS** | | | | | | | | | | | | | | | | | | | | | | | |
| *AQP1* | 193 | 2 | 8 | 20 | 18 | 12 | 24 | 21 | 94 | 8 | 13 | 18 | 14 | 13 | 20 | 20 | 19 | 131 | 28 | 35 | 637 | 446 | 110 |
| *AREG* | 83 | 39 | 45 | 137 | 139 | 12 | 2 | 17 | 1149 | 2283 | 5099 | 118 | 853 | 65 | 23 | 2 | 8 | 1023 | 1745 | 1523 | 1203 | 763 | 1245 |
| *C3* | 33 | 6 | 14 | 12 | 11 | 91 | 87 | 27 | 14809 | 14360 | 2610 | 5545 | 72876 | 3744 | 5 | 3 | 9 | 1753 | 2853 | 2623 | 2743 | 7477 | 4792 |
| *CA2* | 403 | 1264 | 1210 | 1154 | 1317 | 896 | 1110 | 504 | 4408 | 881 | 779 | 735 | 400 | 751 | 197 | 138 | 165 | 1518 | 2133 | 1896 | 3742 | 6486 | 8136 |
| *CA8* | 264 | 34 | 36 | 16 | 25 | 19 | 79 | 125 | 29 | 17 | 33 | 5 | 72 | 17 | 176 | 146 | 200 | 189 | 146 | 133 | 130 | 96 | 34 |
| *CFTR* | 185 | 1605 | 1554 | 2635 | 2485 | 51 | 79 | 27 | 2048 | 256 | 195 | 6 | 18 | 4 | 68 | 38 | 86 | 13267 | 5428 | 5922 | 3744 | 1123 | 1114 |
| *GLIS3* | 339 | 1271 | 1214 | 1451 | 1671 | 909 | 406 | 594 | 1397 | 2480 | 2158 | 1016 | 1407 | 794 | 413 | 501 | 390 | 1091 | 1290 | 1211 | 1055 | 430 | 780 |
| *HES1* | 3714 | 1656 | 1635 | 1241 | 1611 | 6717 | 4262 | 4248 | 2416 | 2573 | 2197 | 1848 | 2769 | 2623 | 1043 | 1486 | 1963 | 1389 | 1304 | 1326 | 1375 | 1963 | 1541 |
| *KRT18* | 8873 | 14574 | 15178 | 18441 | 18095 | 11144 | 7325 | 8854 | 10092 | 14752 | 21729 | 5003 | 11180 | 6217 | 9107 | 7361 | 6304 | 27754 | 30001 | 28695 | 25614 | 26148 | 25318 |
| *KRT19* | 17083 | 7688 | 7521 | 5829 | 5645 | 29425 | 20221 | 18348 | 13215 | 14536 | 28208 | 7394 | 7675 | 7379 | 11320 | 10262 | 9854 | 18458 | 25466 | 24719 | 24271 | 35269 | 32969 |
| *KRT23* | 2 | 0 | 0 | 0 | 0 | 1 | 3 | 8 | 28 | 18 | 44 | 2 | 8 | 0 | 1 | 9 | 3 | 312 | 1361 | 1347 | 568 | 216 | 311 |
| *MUC1* | 62 | 7 | 1 | 11 | 14 | 75 | 122 | 145 | 28 | 43 | 106 | 50 | 47 | 36 | 99 | 74 | 82 | 84 | 87 | 67 | 168 | 560 | 155 |
| *ONECUT1* | 720 | 640 | 663 | 1044 | 1123 | 8662 | 5990 | 2706 | 1622 | 2810 | 1600 | 3610 | 3879 | 2834 | 2628 | 2675 | 3602 | 605 | 315 | 314 | 563 | 479 | 521 |
| *P2RY1* | 233 | 319 | 292 | 269 | 330 | 121 | 107 | 78 | 267 | 140 | 201 | 85 | 65 | 55 | 86 | 189 | 159 | 2089 | 1214 | 1179 | 2686 | 800 | 1924 |
| *PROX1* | 1489 | 441 | 436 | 473 | 592 | 6543 | 9069 | 5728 | 2403 | 2421 | 1825 | 2678 | 2413 | 1754 | 3269 | 1815 | 3033 | 1796 | 1420 | 1348 | 1469 | 1131 | 1283 |
| *SERPINE1* | 118 | 15 | 31 | 121 | 68 | 11 | 8 | 9 | 913 | 1082 | 3032 | 107 | 1550 | 76 | 4 | 5 | 6 | 429 | 330 | 325 | 565 | 1668 | 623 |
| *SLC4A4* | 2134 | 6380 | 6621 | 7657 | 7638 | 2407 | 2667 | 3733 | 1386 | 4273 | 5669 | 1736 | 5272 | 1626 | 7167 | 8939 | 8766 | 7639 | 3891 | 4046 | 5796 | 4139 | 5073 |
| *SPP1* | 2234 | 146 | 172 | 481 | 232 | 20 | 18 | 66 | 1088 | 5635 | 4727 | 93 | 1070 | 69 | 5 | 12 | 14 | 5279 | 966 | 927 | 11302 | 4056 | 7743 |
| *TM4SF1* | 531 | 346 | 335 | 442 | 389 | 19 | 8 | 30 | 2432 | 5426 | 6328 | 391 | 11074 | 399 | 13 | 79 | 72 | 18017 | 18004 | 16945 | 21119 | 44632 | 42232 |
| **ACINAR MARKERS** | | | | | | | | | | | | | | | | | | | | | | | |
| *BHLHA15* | 69 | 19 | 4 | 5 | 10 | 82 | 56 | 54 | 97 | 65 | 62 | 74 | 206 | 113 | 49 | 46 | 37 | 38 | 35 | 39 | 26 | 76 | 71 |
| *CPA1* | 32 | 4 | 4 | 1 | 0 | 50 | 172 | 363 | 24 | 42 | 15 | 57 | 258 | 74 | 371 | 407 | 333 | 108 | 76 | 68 | 100 | 184 | 46 |
| *CPA2* | 138 | 55 | 55 | 72 | 61 | 53 | 209 | 1168 | 1005 | 182 | 592 | 13 | 24 | 12 | 1210 | 686 | 466 | 174 | 67 | 58 | 75 | 95 | 34 |
| *CPE* | 8624 | 5380 | 5679 | 7232 | 6127 | 5353 | 8237 | 14202 | 4435 | 3617 | 9884 | 2123 | 1741 | 1980 | 11898 | 12162 | 8479 | 1085 | 2117 | 1978 | 1738 | 866 | 646 |
| *PRSS1* | 564 | 0 | 0 | 0 | 0 | 2461 | 874 | 690 | 59 | 54 | 82 | 11 | 0 | 13 | 777 | 327 | 479 | 78 | 60 | 76 | 63 | 45 | 31 |
| *PTF1A* | 391 | 1 | 4 | 0 | 3 | 36 | 278 | 526 | 9 | 17 | 22 | 4 | 7 | 2 | 911 | 1059 | 1112 | 23 | 9 | 15 | 25 | 7 | 7 |
| *RBPJL* | 8 | 0 | 0 | 1 | 0 | 3 | 5 | 15 | 0 | 2 | 0 | 0 | 0 | 0 | 17 | 14 | 7 | 3 | 1 | 1 | 1 | 2 | 0 |
| *RHOV* | 42 | 77 | 70 | 53 | 34 | 15 | 19 | 50 | 20 | 58 | 69 | 22 | 15 | 18 | 64 | 31 | 26 | 253 | 211 | 190 | 182 | 329 | 213 |
| *RNASE1* | 463 | 24 | 5 | 2 | 9 | 226 | 218 | 212 | 345 | 141 | 295 | 105 | 11 | 71 | 70 | 85 | 72 | 423 | 702 | 722 | 2277 | 3440 | 6507 |
| **ENDOCRINE MARKERS** | | | | | | | | | | | | | | | | | | | | | | | |
| *CHGA* | 695 | 86 | 77 | 120 | 107 | 4802 | 69260 | 186320 | 2315 | 608 | 38794 | 2913 | 122 | 4327 | 200359 | 129425 | 130414 | 735 | 2446 | 2614 | 425 | 35 | 59 |
| *FOXA2* | 5814 | 2578 | 2547 | 1740 | 2128 | 8755 | 6474 | 5620 | 1818 | 2776 | 2446 | 3828 | 3827 | 4645 | 4742 | 5137 | 5374 | 1250 | 948 | 973 | 1215 | 2105 | 1690 |
| *GCG* | 2 | 0 | 0 | 0 | 0 | 1 | 17 | 22 | 204 | 1 | 177 | 0 | 0 | 0 | 3499 | 266 | 647 | 31 | 8 | 15 | 22 | 0 | 0 |
| *GP2* | 122 | 107 | 144 | 121 | 113 | 3 | 11 | 24 | 219 | 913 | 767 | 662 | 962 | 374 | 26 | 335 | 251 | 4325 | 2275 | 2664 | 11226 | 19630 | 42497 |
| *INS* | 0 | 0 | 0 | 0 | 0 | 0 | 6 | 38 | 14 | 0 | 123 | 0 | 0 | 0 | 1446 | 152 | 207 | 55 | 14 | 10 | 1 | 0 | 0 |
| *INSM1* | 1110 | 14 | 31 | 15 | 15 | 5339 | 9952 | 13278 | 25 | 2 | 383 | 101 | 3 | 59 | 9523 | 9134 | 10945 | 30 | 29 | 33 | 35 | 2 | 14 |
| *NEUROD1* | 4 | 4 | 1 | 2 | 8 | 313 | 1631 | 3583 | 34 | 7 | 390 | 16 | 2 | 29 | 3543 | 2817 | 3016 | 9 | 23 | 22 | 8 | 1 | 1 |
| *NEUROG3* | 77 | 7 | 5 | 3 | 6 | 1584 | 4181 | 6949 | 8 | 2 | 28 | 104 | 0 | 88 | 5729 | 3341 | 4558 | 3 | 5 | 9 | 3 | 1 | 2 |
| *NKX2-2* | 6 | 7 | 3 | 2 | 3 | 386 | 1412 | 1877 | 7 | 2 | 104 | 60 | 1 | 39 | 2264 | 1978 | 2250 | 39 | 9 | 11 | 11 | 14 | 7 |
| *NKX6-1* | 1024 | 8015 | 7936 | 6082 | 6631 | 26 | 86 | 631 | 113 | 436 | 200 | 199 | 3153 | 141 | 4035 | 4300 | 3863 | 3742 | 2798 | 3072 | 2325 | 1203 | 558 |
| *NKX6-1/2* | 1030 | 8022 | 7939 | 6084 | 6633 | 412 | 1497 | 2507 | 120 | 437 | 304 | 258 | 3154 | 181 | 6299 | 6278 | 6112 | 3781 | 2807 | 3083 | 2335 | 1218 | 565 |
| *NKX6-2* | 30 | 151 | 111 | 69 | 39 | 228 | 95 | 305 | 159 | 610 | 791 | 243 | 2281 | 204 | 85 | 215 | 192 | 3024 | 1110 | 1153 | 1774 | 9459 | 6797 |
| *PAX4* | 0 | 0 | 1 | 3 | 4 | 457 | 1886 | 2937 | 5 | 2 | 62 | 120 | 5 | 76 | 1649 | 1462 | 1776 | 1 | 1 | 2 | 0 | 0 | 1 |
| *PDX1* | 8993 | 4344 | 4164 | 3639 | 3660 | 5416 | 4586 | 4899 | 2254 | 3436 | 2748 | 3503 | 5169 | 3435 | 6426 | 6606 | 7739 | 4484 | 3508 | 3628 | 3707 | 3384 | 4136 |
| *RBPJ* | 1679 | 1989 | 1754 | 1969 | 1748 | 3739 | 3439 | 1966 | 2164 | 2760 | 1349 | 3393 | 2776 | 2124 | 5470 | 5293 | 6340 | 4240 | 4122 | 3440 | 3164 | 2601 | 5665 |
| *RFX3* | 579 | 578 | 534 | 575 | 655 | 2178 | 4541 | 2753 | 506 | 758 | 628 | 998 | 903 | 1119 | 2584 | 2081 | 2600 | 468 | 688 | 597 | 444 | 651 | 356 |
| *RFX6* | 499 | 770 | 843 | 1090 | 1135 | 4795 | 4672 | 2618 | 1621 | 123 | 398 | 1600 | 111 | 1506 | 2897 | 1990 | 2557 | 6 | 21 | 24 | 6 | 4 | 12 |
| *SOX9* | 5366 | 6139 | 6062 | 4409 | 6196 | 6647 | 6361 | 8838 | 6010 | 12499 | 5133 | 8767 | 10194 | 8092 | 5849 | 6377 | 5710 | 3517 | 3217 | 3148 | 3605 | 5708 | 3450 |
| **MPC AND BP MARKERS** | | | | | | | | | | | | | | | | | | | | | | | |
| *PTF1A* | 391 | 1 | 4 | 0 | 3 | 36 | 278 | 526 | 9 | 17 | 22 | 4 | 7 | 2 | 911 | 1059 | 1112 | 23 | 9 | 15 | 25 | 7 | 7 |
| *CPA1* | 32 | 4 | 4 | 1 | 0 | 50 | 172 | 363 | 24 | 42 | 15 | 57 | 258 | 74 | 371 | 407 | 333 | 108 | 76 | 68 | 100 | 184 | 46 |
| *DCDC2A* | 100 | 112 | 121 | 135 | 108 | 359 | 322 | 606 | 1855 | 4051 | 3233 | 615 | 2373 | 881 | 97 | 36 | 38 | 259 | 787 | 758 | 973 | 266 | 1458 |
| **LIVER AND GUT MARKERS** | | | | | | | | | | | | | | | | | | | | | | | |
| *AFP* | 43055 | 2378 | 2605 | 2636 | 2654 | 9238 | 32423 | 14262 | 1268706 | 234011 | 120526 | 183745 | 131218 | 148309 | 12315 | 2596 | 11978 | 5298 | 9358 | 9315 | 35124 | 26676 | 18943 |
| *CDX2* | 2575 | 3850 | 3690 | 3443 | 3258 | 3133 | 1605 | 1051 | 4202 | 905 | 1650 | 2233 | 645 | 2497 | 156 | 130 | 172 | 3190 | 5360 | 5288 | 3295 | 1737 | 2419 |
| *HHEX* | 10315 | 337 | 402 | 572 | 758 | 6612 | 4818 | 3701 | 2069 | 3768 | 2962 | 2279 | 2650 | 1793 | 961 | 299 | 613 | 743 | 488 | 540 | 724 | 249 | 229 |
| *TTR* | 5295 | 78 | 94 | 142 | 111 | 12132 | 8680 | 4552 | 183515 | 11224 | 34773 | 1890 | 2589 | 1638 | 2984 | 1323 | 2315 | 621 | 1147 | 1068 | 1934 | 848 | 646 |

**TABLE S5** Normalised RNA-Seq counts of selected genes from feeder expanded cells, fibronectin expanded cells and vitronectin-N expanded cells

| Antigen | Species | Supplier, cat no | Application | Dilution |
|---|---|---|---|---|
| AFP | Mouse | Abcam, ab3980 | IF | 1:200 |
| CDX2 | Rabbit | Abcam, ab76541 | IF | 1:100 |
| C-PEP-Alexa 647 | Mouse | BD pharmigen, 565831 | FC | 1:200 |
| FOXA2 | Rabbit | Merck, 07-633 | IF | 1:200 |
| GCG | Mouse | Sigma, G2654 | IF | 1:500 |
| GCG | Rabbit | Invitrogen, PA5-13442 | FC | 1:500 |
| GP-2 | Mouse | MBL D277-3 | IF | 1:300 |
| HNF1b | Goat | Santa Cruz, sc-7411 | IF | 1:50 |
| INS | Guinea pig | Abam, ab7842 | IF | 1:1000 |
| NEUROD1 | Mouse | Abcam, ab60704 | IF/FC | 1:500/1:200 |
| NGN3 | Sheep | R&D, AF3444 | IF/FC | 1:100 |
| NKX6.1 | Mouse | DSHB (F55A10) | IF/FC | 1:500/1:250 |
| PDX1 | Goat | R&D, AF2419 | IF/FC | 01:40 |
| SOX9 | Rabbit | Merck, AB5535 | IF/FC | 1:1000 |

**Table S6** List of primary antibodies for immunofluorescense (IF) and flow cytometry (FC)

| Antigen | Fluorophore | Species | Supplier, cat no | Dilution |
|---|---|---|---|---|
| Goat IgG | Alexa 647 | Donkey | Invitrogen, A21447 | 1:500 |
| Guinea pig IgG | Alexa 647 | Goat | Abcam, ab150187 | 1:500 |
| Mouse IgG | Alexa 568 | Donkey | Invitrogen, A10037 | 1:500 |
| Mouse IgG | Alexa 568 | Goat | Invitrogen, A11004 | 1:500 |
| Mouse IgG | Alexa 647 | Goat | Invitrogen, A21235 | 1:500 |
| Mouse IgG isotype | Alexa 647 | Goat | BD pharmigen, 557783 | 1:5 |
| Rabbit IgG | Alexa 488 | Donkey | Invitrogen, A21206 | 1:500 |
| Rabbit IgG | Alexa 488 | Goat | Invitrogen, A11070 | 1:500 |
| Rabbit IgG | Alexa 647 | Goat | Invitrogen, A21244 | 1:500 |
| Sheep IgG | Alexa 568 | Donkey | Invitrogen, A21099 | 1:500 |

**Table S7** List of secondary antibodies

| Stage | Gene and primer direction | Sequence |
|---|---|---|
| **Housekeeping** | *TBP F* | TATCGTGGTCACACTTGTTGAG |
| | *TBP R* | ACCATCCTCTGAGACCGTTTT |
| **PP** | *PDX1 F* | GGAGAAAGATGGACCCCTGG |
| | *PDX1 R* | CAGCCTCTACCTCGGAACAG |
| | *FOXA2 F* | ACACCACTACGCCTTCAACC |
| | *FOXA2 R* | CCTTGAGGTCCATTTTGTGG |
| | *HHEX F* | ACGGTGAACGACTACACGC |
| | *HHEX R* | CTTCTCCAGCTCGATGGTCT |
| | *TTR F* | ACTTGGCATCTCCCCATTC |
| | *TTR R* | TAGGAGTAGGGGCTCAGCAG |
| | *PTF1A F* | TTCACCGACCAGTCTTCACG |
| | *PTF1A R* | GTGGCTAAGGAACTCCACCT |
| | *NKX6.1 F* | ATTCGTTGGGGATGACAGAG |
| | *NKX6.1 R* | CGAGTCCTGCTTCTTCTTGG |
| | *AFP F* | GCAGCAGTCTGAATGTCCGTAC |
| | *AFP R* | TGCCCAGTTTGTTCAAGAAGC |
| | *CDX2 F* | GCTGGAGAAGGAGTTTCACTACAGT |
| | *CDX2 R* | AACCAGATTTTAACCTGCCTCTCA |
| **S7** | *PDX1 F* | GGAGAAAGATGGACCCCTGG |
| | *PDX1 R* | CAGCCTCTACCTCGGAACAG |
| | *NKX6.1 F* | ATTCGTTGGGGATGACAGAG |
| | *NKX6.1 R* | CGAGTCCTGCTTCTTCTTGG |
| | *GCG F* | GAGACATGCTGAAGGGACCT |
| | *GCG R* | CTTCCTCGGCCTTTCACCAG |
| | *SST F* | AGCTGCTGTCTGAACCCAAC |
| | *SST R* | GCTCAAGCCTCATTTCATCC |
| | *GK F* | AGCGTGAAGACCAAACACCA |
| | *GK R* | ATGCTTGTCCAGGAAGTCGG |
| | *MAFA F* | GCGGAGAACGGTGATTTCTA |
| | *MAFA R* | GAAGGTGGGAACGGAGAAC |
| | *INS F* | AGATCACTGTCCTTCTGCCA |
| | *INS R* | CGCACAGGTGTTGGTTCA |
| | *PC1/3 F* | TACAAGCACAGAGACGACCG |
| | *PC1/3 R* | CGCAGGGTAAGGAAGAAGCA |
| | *SLC30A8 (ZnT8) F* | GGCCGTCATGGAGTTTCTT |
| | *SLC30A8 (ZnT8) R* | CACCGGTTTCTGTTGGAGTT |

**Table S8** List of forward (F) and reverse (R) primers used for qPCR
